# Identification and *in vitro* characterization of UDP-GlcNAc-RNA cap-modifying and decapping enzymes

**DOI:** 10.1101/2023.08.17.553725

**Authors:** Frederik Weber, Nikolas Alexander Motzkus, Leona Brandl, Marvin Möhler, Andres Jäschke

## Abstract

In recent years, several noncanonical RNA caps derived from cofactors and metabolites have been identified. Purine-containing RNA caps have been extensively studied, with multiple decapping enzymes identified and efficient capture and sequencing protocols developed for nicotinamide adenine dinucleotide (NAD)-RNA, which allowed for a stepwise elucidation of capping functions. Despite being identified as an abundant noncanonical RNA-cap, UDP-sugar-capped RNA remains poorly understood, which is partly due to its complex *in vitro* preparation. Here, we describe a scalable synthesis of sugar-capped uridine-guanosine dinucleotides from readily available protected building blocks and their enzymatic conversion into several cell wall precursor-capped dinucleotides. We employed these capped dinucleotides in T7 RNA polymerase-catalyzed *in vitro* transcription reactions to efficiently generate RNAs capped with uridine diphosphate *N*-acetylglucosamine (UDP-GlcNAc), its *N*-azidoacetyl derivative UDP-GlcNAz, and various cell wall precursors. We furthermore identified four enzymes capable of processing UDP-GlcNAc-capped RNA *in vitro*: MurA, MurB and MurC from *Escherichia coli* can sequentially modify the sugar-cap structure and were used to introduce a bioorthogonal, clickable moiety, and the human Nudix hydrolase Nudt5 was shown to efficiently decap UDP-GlcNAc-RNA. Our findings underscore the importance of efficient synthetic methods for capped model RNAs and provide useful enzymatic tools for the potential use in and development of UDP-GlcNAc capture and sequencing protocols.

## INTRODUCTION

RNA 5’-capping is an important co- and post-transcriptional modification that can impact RNA stability, splicing, nuclear export and translation (1). The group of known RNA caps has recently been expanded to include various nucleotide-containing coenzymes, such as 3’-dephospho-coenzyme A (dpCoA) (2), flavin adenine dinucleotide (FAD) (3) and nicotinamide adenine dinucleotide (NAD) (4) of which the latter is best understood. Efficient capture and sequencing methods for NAD-RNAs (5-11) enabled the identification of NAD-capped RNA species in *E. coli* (5), *B. subtilis* (12), *S. aureus* (13), *S. cerevisiae* (14), *A. thaliana* (15) and *H. sapiens* (16). The incorporation of endogenous NAD as a noncanonical initiator nucleotide (NCIN) by cellular RNA polymerases (RNAP) has been identified as the predominant biosynthetic pathway to NAD-RNA (17), and decapping enzymes that remove the NAD-cap have been identified in various organisms (16,18-22). Efficient methods for the synthesis and purification of model NAD-RNAs have contributed to this rapid progress (23-25).

Part of the newly discovered noncanonical RNA cap structures is formed by the structurally diverse group of nucleotide sugars (26). So far, uridine diphosphate *N*-acetyl glucosamine (UDP-GlcNAc) and UDP-glucose (UDP-Glc) were found to be linked to cellular RNA molecules (3). *In vitro, E. coli* RNAP can incorporate these metabolites as NCINs into RNA (27). As UDP-GlcNAc is an important precursor for the biosynthesis of murein and lipopolysaccharides, its concentration in *E. coli* is very high (9.2 mM) (28), which would favor the incorporation as NCIN *in vivo,* too. And indeed, the UDP-GlcNAc-RNA modification was found by mass spectrometric detection of digested total RNA samples from a variety of organisms, and with an abundance comparable to or even surpassing that of the NAD cap (3). Despite this, a protocol for the specific identification of sugar-capped RNA sequences is still to be developed and the biological relevance of the UDP-GlcNAc modification remains elusive.

For the identification of sugar-capped RNA species, and UDP-GlcNAc-capped RNA in particular, different methods are conceivable. A well-established method for identifying and profiling sugar-conjugated proteins is metabolic labeling with sugar derivatives that bear bioorthogonal handles such as azido sugars (29). This approach has been successfully transferred to the identification of internally glycosylated RNAs in mammals (30) and may be applicable to the identification of UDP-GlcNAc-RNAs. A more direct approach would be to identify and utilize an enzyme capable of specifically modifying the UDP-GlcNAc-cap with a bioorthogonal handle, as is the case with the NAD captureSeq protocol (5,6). A third approach would require the identification of a selective decapping enzyme converting sugar-capped RNA to 5’-monophosphate-RNA, which can then be utilized for the specific ligation of a sequencing adapter. This approach, termed CapZyme-Seq, has been successfully applied to NAD-RNA (31) and recently led to the discovery of FAD-RNA (32). However, no enzymes that process the cap structure of UDP-GlcNAc-RNA have been identified thus far.

A key element for the development of a diverse toolbox for potential use in UDP-GlcNAc-RNA identification protocols is the synthetic access to model UDP-GlcNAc-RNAs. But while RNAs with purine-containing cap structures, like NAD, can be easily synthesized *in vitro* by noncanonical transcription initiation with T7 RNAP (23), the incorporation of uridine-containing caps presents a particular challenge. T7 RNAP has a strong preference for transcription initiation with guanosine, with which it interacts *via* the N-7 of the purine ring and the O-6 of guanine (33). While adenosine-containing cofactors can provide some of these interactions (N-7) and can initiate transcription (23,34), this is not the case for pyrimidine nucleotides, like UDP-GlcNAc. On the other hand, T7 RNAP does not interact with the 5’-triphosphate of the initiating nucleotide (35), and a wide variety of 5’-modified guanosine nucleotides have been incorporated by T7 RNAP as NCIN (36). These include di- and oligonucleotides carrying guanosine at their 3’-end (37-40). Thus, the poor transcription efficiency of UDP-sugars can be circumvented by using a 5’-sugar-modified uridine-guanosine (UpG) dinucleotide instead. Recently, *Depaix et al.* applied this approach to the *in vitro* transcription (IVT) of UDP-Glc- and UDP-GlcNAc-RNAs (40).

Inspired by this approach, we used a modified synthetic procedure to chemically synthesize a GlcNAc-capped UpG dinucleotide as well as a novel *N*-azidoacetyl derivative (GlcNAz) and converted both of them enzymatically into a set of novel cell wall precursor-capped dinucleotides. We further show efficient ways for the enzymatic incorporation of sugar-capped dinucleotides into RNAs of different lengths and sequences, one of which provides synthetic access to pure sugar-capped RNAs.

These sugar-capped dinucleotides and model RNA aided in the discovery of four enzymes capable of processing their cap structure: We found that enzymes of the *E. coli* murein biosynthetic pathway, namely MurA, MurB and MurC, are capable of converting and modifying sugar-capped RNA *in vitro*. We furthermore demonstrated the potential of MurC to install bioorthogonal propargyl and azido handles into sugar caps, allowing for the selective click labeling of long (80 nt) sugar-capped model RNAs. Additionally, we discovered an efficient decapping activity of the Nudix hydrolase hNudt5, which could remove sugar caps from UDP-GlcNAc-RNA and azide-labeled UDP-GlcNAz-RNA. Our work further reports the synthesis of several previously inaccessible RNAs modified with cell wall precursors and suggests the use of the discovered enzymatic tools for the development of capture and sequencing protocols for UDP-GlcNAc-RNA.

## MATERIAL AND METHODS

### Reagents

Chemicals were purchased from Sigma Aldrich (Steinheim, Germany), Biosynth (Staad, Switzerland), abcr (Karlsruhe, Germany), Thermo Fisher Scientific (Waltham, MA, USA), Carl Roth (Karlsruhe, Germany) or Jena Bioscience (Jena, Germany) and used without further purification. Dry solvents were purchased in sealed bottles over molecular sieves. Oligonucleotides were purchased from Integrated DNA Technologies (Coralville, IA, USA) and are summarized in **Table S1**. DNAzyme and ssDNA for RNA ligation were PAGE purified prior to use.

Streptavidin (N7021S), RNA 5’ pyrophosphohydrolase (M0356S), NEBuffer 2, NudC (M0607S), NEBuffer r3.1, low range ssRNA ladder (N0364S) and the Monarch RNA Cleanup Kit (T2030L) were purchased from NEB (Ipswich, MA, USA). T4 DNA ligase (EL0013), NcoI (FD0574), XhoI (FD0694) and Phusion high-fidelity DNA polymerase (F530S) were purchased from Thermo Fisher Scientific (Waltham, MA, USA). RNaseOUT (10777019) and SYBR Gold (S11494) were purchased from Invitrogen, (Carlsbad, CA, USA). DNAse I (4716728001; Roche, Basel, Switzerland) was purchased from Merck (Darmstadt, Germany). The QIAquick PCR Purification Kit (28106) was purchased from Qiagen (Venlo, Netherlands). The BIOMOL Green phosphate quantification kit (BML-AK111-0250) was purchased from Enzo Life Sciences (Farmingdale, NY, USA).

### Biological Resources

BL21 Star (DE3) and DH5α cells were purchased from Thermo Fisher Scientific (Waltham, MA, USA). The pET-28c(+) vector was purchased from Merck Millipore (Burlington, MA, USA).

### Chemical synthesis of sugar-capped dinucleotides

A detailed synthesis for GlcNAcppUpG **4** and GlcNAzppUpG **5** and the required building blocks can be found in the Supplementary Material.

#### Phosphoimidazolide coupling

The building block to be activated (protected sugar 1-phosphate or protected 5’-monophosphate-dinucleotide) (1.00 eq) and CDI (2.40 eq) were dissolved in anh. DMF under an argon atmosphere and stirred at RT for 3 h. Afterwards, it was added to an excess of the second building block (1.50 - 4.00 eq) and MgCl_2_ (6.00 eq) dissolved in anh. DMF and precooled to 0°C. The mixture was allowed to warm to RT and it was stirred overnight. The reaction was cooled to 0°C and quenched by the addition of excess EDTA in H_2_O (500 mM, pH 8) and TEAA (100 mM, pH 7).

#### Cleavage of base-labile protective groups

Protected intermediates were dissolved in a mixture of 28% NH_4_OH in H_2_O and 33% methylamine in EtOH (1 / 1) and heated at 50°C for 1.5 h. The solvent was evaporated and the residue was dissolved in TEAA (100 mM, pH 7).

#### TBDMS deprotection

Substrates were dissolved in a mixture of DMF / Et_3_N / Et_3_N*3HF (3 / 1 / 2) in a falcon tube and the reaction mixture was stirred at 50°C for 2 h. The mixture was cooled to 0°C and TEAA (100 mM, pH 7) was added.

#### Purification of sugar-capped dinucleotides and protected intermediates

After reactions were completed and quenched, Celite was added and the solvent was evaporated under reduced pressure. The crude product was purified using a PuriFlash PF420 system equipped with a 12 g PF-30C18AQ column (Interchim, Montluçon, France) and 10 mM triethylammonium acetate (TEAA) in H_2_O (pH 7) or ACN as mobile phase (for details see Supplementary Material). Fractions containing the product were lyophilized and the residue was repeatedly dissolved in Milli-Q water and lyophilized until a stoichiometric amount of triethylammonium cations was present according to ^1^H NMR measurements. For final products, if necessary, additional semi-preparative HPLC purification was conducted. Triethylammonium cations were exchanged for sodium by dissolving capped dinucleotides in Milli-Q water and passing them through a short column of Amberlite IR120 Na^+^ form. Fractions containing product were combined and lyophilized.

### Enzymatic synthesis of EP-GlcNAcppUpG 6

Compound **6** was synthesized using a modified literature procedure (41). GlcNAcppUpG **4** (23.0 mg, 22.6 µmol, 1.00 eq) in a total volume of 8 ml containing Tris-HCl (40 mM, pH 7.8), phosphoenolpyruvate (10 mM, 80.0 µmol, 3.54 eq) and MurA (0.1 mg / ml) was distributed to 1.5 ml Eppendorf tubes and incubated at 37°C, 300 rpm overnight. The reaction mixture was collected in a round-bottom flask and TEAA (100 mM, 10 ml, pH 7) was added. The product was purified as described above and EP-GlcNAcppUpG **6** was isolated as a white powder (16.2 mg, 14.6 µmol, 65%).

### Enzymatic synthesis of MurNAcppUpG 7

MurNAcppUpG **7** was synthesized using a modified literature procedure (42). EP-GlcNAcppUpG **6** (15.7 mg 14.4 µmol, 1.00 eq) in a total volume of 12 ml containing Tris-HCl (40 mM, pH 7.8), KCl (1.5 mM), DTT (1 mM), NADPH (4 mM, 48.0 µmol, 3.33 eq) and MurB (0.06 mg / ml) was distributed to 1.5 ml Eppendorf tubes and incubated at 37°C, 300 rpm overnight. Reaction mixtures were combined in a round-bottomed flask and TEAA (100 mM, 10 ml, pH 7) was added. The product was purified as described above and MurNAcppUpG **7** was isolated as white powder (8.30 mg, 7.46 µmol, 52%).

#### *In vitro* transcription

DNA sequences used for template preparation can be found in **Table S1** and prepared RNA sequences in **Table S2**. IVT reactions were conducted in a final volume of 100 µl containing Tris-HCl (40 mM, pH 8.1), spermidine (1 mM), MgCl_2_ (22 mM), Triton-x-100 (0.01%), DMSO (5%), nucleoside triphosphates (NTPs, 3 mM), dithiothreitol (DTT, 10 mM), template (5 µM) and T7 RNAP (0.05 mg/ml, lab prepared stock). Reactions were incubated at 37°C for 3-4 h before adding DNase I (0.1 U/µl) (Roche) and incubating for another 30 min at 37°C. For capped RNA, a final concentration of 6 mM NCIN was used and the GTP concentration was either reduced to 0.6 mM (for longer RNA) or GTP was absent (short RNA).

Workup was conducted depending on the used IVT system. For short RNA, reactions were diluted to 200 µl and pyrophosphate was separated by centrifugation and transferring the supernatant to a new reaction tube. The supernatant was extracted twice with P / C / I and twice with diethyl ether. The solvent was lyophilized, and the residue was dissolved in 100 µl TEAA (100 mM, pH 7). RNA was purified by RP-HPLC using VDSpher PUR 100 C18-SE 250 × 4.6 mm, 5 µm column (VDS optilab, Berlin, Germany; gradient: 1.6% buffer B for 5 min, to 14% buffer B in 35 min, flow rate 1 ml/min). Eluted RNA was collected in Eppendorf tubes, lyophilized, dissolved in Milli-Q water and analyzed by ESI-MS. Purification of longer RNA was conducted by direct addition of denaturing loading dye (2x TBE, 12 mM EDTA, BPB, XC in formamide) to the IVT reaction followed by purification using preparative 10% denaturing PAGE.

### Enzyme preparation

Enzymes used in this study have been overexpressed in *E. coli* BL21 Star (DE3) and purified by fast protein liquid chromatography (see **Fig. S1-S6**, for details see Supplementary Material). *E. coli* BL21(DE3) containing genes of MurA, MurB, NudF and hNudt5 in a Pet-28C(+) vector were kindly provided by Dr. F. Abele and Dr. K. Höfer (MurA, MurB) and Dr. A. Krause and Dr. K. Höfer (NudF, hNudt5). The gene sequence of all utilized enzymes can be seen in **Table S3**.

### MurA reaction

Substrates (1 mM), PEP (10 mM), MurA (4 µM) and Tris-HCl (40 mM, pH 7.8) in a final volume of 100 µl were incubated at 37°C. At different time points, 100 µl TEAA (200 mM, pH 7) was added, the enzyme was separated by filtration through a 10 kDa MW cut-off filter and the filtrate was analyzed by RP-HPLC and ESI-MS.

### MurA and MurB one-pot reaction

Substrates (1 mM), PEP (10 mM), MurA (4 µM) and Tris-HCl (40 mM, pH 7.8) in a final volume of 100 µl were incubated for 4 h at 37°C. Then the reaction volume was increased to 150 µl including KCl (1.5 mM), DTT (1 mM), NADPH (3.5 mM) and MurB (1 µM) and incubated overnight at 37°C. TEAA (50 µL, 400 mM, pH 7) was added, the enzymes were separated by filtration through a 10 kDa MW cut-off filter and the filtrate was analyzed by RP-HPLC and ESI-MS. If GlcNAzppUpG **5** was used as a substrate, incubation with MurA was conducted for 20 h and incubation after addition of MurB was conducted for another 7 h in the absence of DTT.

For UDP-GlcNAc-11mer RNA, 6 µg RNA was used and RNaseOUT recombinant ribonuclease inhibitor (1 µl) (Invitrogen) was added. For workup, reactions were diluted to 200 µl, extracted twice with P / C / I and twice with diethyl ether and lyophilized. The residue was dissolved in TEAA (100 µl, 100 mM, pH 7) and analyzed by RP-HPLC and ESI-MS.

### MurC reaction

MurC reactions were conducted using a modified literature procedure (43). Substrate (0.5 mM), DTT (1 mM), (NH_4_)_2_SO_4_ (25 mM), MgCl_2_ (20 mM), ATP (5 mM), Tris-HCl (100 mM, pH 8) MurC (1.6 µM) and alanine or an alanine derivative (1 mM or 10 mM) in a total volume of 50 µl were incubated for 4 h at 37°C. Reactions were diluted to 200 µl with TEAA (100 mM, pH 7) and the enzyme was separated by filtration using a 10 kDa MW cut-off filter. The filtrate was analyzed using RP-HPLC and ESI-MS.

If UDP-MurNAc-RNA was used as a substrate, alanine derivatives were used in a concentration of 10 mM and RNaseOUT recombinant ribonuclease inhibitor (1 µl) (Invitrogen) was added. Reactions with UDP-MurNAc-11mer RNA contained 6 µg RNA and were purified as described for the MurA and MurB one-pot reaction. Reactions with UDP-MurNAc-80mer RNA contained 2 µg RNA and products were purified using the Monarch RNA Cleanup Kit (NEB).

### Decapping reaction

Substrates (0.625 mM), DTT (1 mM), MgCl_2_ (10 mM), Tris-HCl (50 mM, pH 8) and NudF or hNudt5 (4 µM) in a total volume of 20 µl were incubated for 4 h at 37°C. For negative controls, Milli-Q water was added instead of enzyme or substrate. For work-up, reactions were diluted to 200 µl with TEAA (100 mM, pH 7) and the enzyme was separated by filtration through a 3 kDa MW cut-off filter. Filtrates were analyzed by RP-HPLC and ESI-MS.

For NudC reactions, substrates (0.625 mM), DTT (5 mM), NEBuffer r3.1 (1x) and NudC (0.5 µM) (NEB) in a total volume of 20 µl were incubated for 4 h at 37°C. For negative controls Milli-Q water was added instead of enzyme or substrate. Work-up was conducted as described above.

For hNudt5 decapping reactions of UDP-sugar-11mer RNA, 1 µg RNA was used as substrate. Negative controls contained Milli-Q water instead of hNudt5. For work-up, reactions were diluted to 200 µl containing EDTA (1 mM) and extracted twice with P / C / I and twice with diethyl ether. RNA was precipitated with sodium acetate and ethanol.

## RESULTS

### Sugar-capped dinucleotides for NCIN-capping of RNA are accessible by coupling of protected building blocks and enzymatic conversion

As larger amounts of sugar-capped dinucleotides are required for the incorporation into RNA and the investigation of cap-processing enzyme candidates, we applied a scalable synthesis procedure: We started the synthesis from protected building blocks, namely sugar 1-phosphate **1** or **2** and 5’-monophosphate-dinucleotide **3**, which facilitated access to starting materials, purification of intermediates and scaling, to chemically synthesize GlcNAc-capped uridine-guanosine dinculeotide (GlcNAcppUpG **4**), which corresponds to a natural UDP-GlcNAc-cap structure (3), and its *N*-azidoacetyl derivative (GlcNAzppUpG **5**), containing a clickable azido moiety (**Figure 1**).

**Figure 1:**
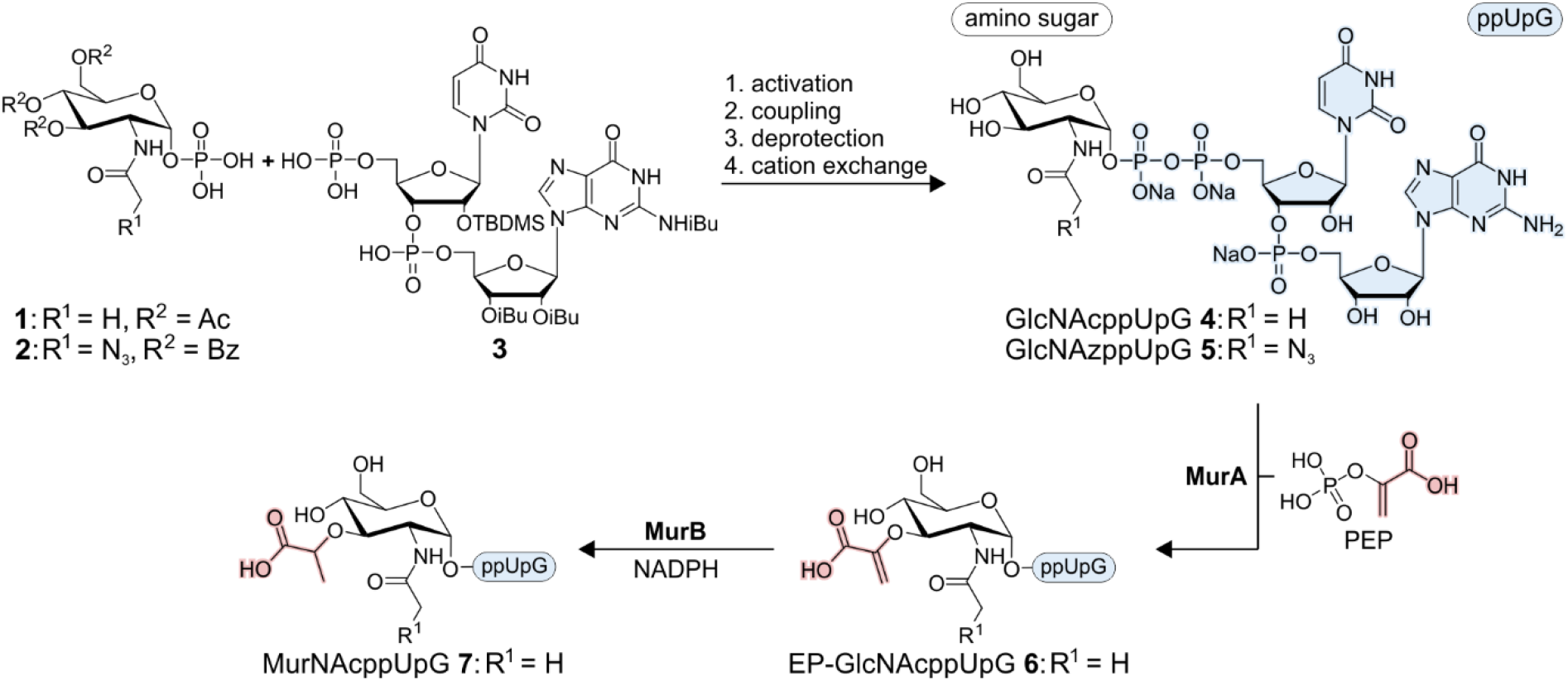
Chemical and enzymatic synthesis of sugar-capped dinucleotides. GlcNAcppUpG **4** and GlcNAzppUpG **5** were synthesized starting from sugar 1-phosphate building blocks **1** or **2** and 5’-monophosphate-dinucleotide **3**. After activation of one reactant as phosphoimidazolide, it was coupled to the second reactant leading to a fully protected intermediate. The intermediate was sequentially deprotected by treatment with a NH_4_OH / methylamine mixture and Et_3_N*3HF before cations were exchanged for sodium. GlcNAcppUpG **4** was received in an overall yield of 48% and GlcNAzppUpG **5** in an overall yield of 18%. GlcNAcppUpG **4** was further converted to EP-GlcNAcppUpG **6** by MurA in the presence of PEP in a yield of 65% and EP-GlcNAcppUpG **6** was converted to MurNAcppUpG **7** by MurB with the help of NADPH in a yield of 52%.

The coupling of the protected sugar 1-phosphate **1** or **2** and the dinucleotide-5’-phosphate **3** building block through pyrophosphate bond formation was achieved using phosphoimidazolide chemistry (24,40). For this purpose, one of the two building blocks was activated with carbonyldiimidazole (CDI) and added dropwise to an excess of the other reactant. We activated the phosphate component that was “rarer”, namely the dinucleotide **3** for the synthesis of GlcNAcppUpG **4** and the protected sugar-1-phosphate **2** for the synthesis of GlcNAzppUpG **5**. The fully protected, capped dinucleotides were deprotected sequentially using NH_4_OH / methylamine followed by treatment with Et_3_N*3HF. After a cation exchange to sodium counter ions, GlcNAcppUpG **4** was obtained with an overall yield of 48% and GlcNAzppUpG **5** with a yield of 18% compared to the input of **3** and **2**, respectively. We synthesized 67 mg GlcNAcppUpG **4** and 20 mg GlcNAzppUpG **5**, and our synthetic approach can be easily scaled up further. In addition, all reactants required for this synthesis were prepared in scalable liquid-phase syntheses described in the Supplementary Information (**Figure S7-S9**).

To be able to study enzymes of the bacterial murein biosynthetic pathway for their capability to sequentially process UDP-GlcNAc-RNA, we further reacted the 3-OH group of GlcNAcppUpG **4**. This was done by *in vitro* modifications using the enzymes MurA and MurB, to yield the corresponding enolpyruvyl (EP) derivative (EP-GlcNAcppUpG **6**) and the *N*-acetylmuramic acid (MurNAc)-capped uridine-guanosine dinucleotide (MurNAcppUpG **7**) (**Figure 1**). For this, large-scale enzymatic reactions were set up converting first GlcNAcppUpG **4** in the presence of MurA and phosphoenolpyruvate (PEP) to product **6** which was then converted in the presence of MurB and NADPH to dinucleotide **7**. Products **6** and **7** were obtained in yields of 65% and 52%, respectively.

### Sugar-capped dinucleotides are readily incorporated into RNA by T7 RNA polymerase

Our chemo-enzymatically synthesized sugar-capped dinucleotides **4, 5, 6** and **7** bearing a 3’-guanosine allow for the direct incorporation as NCINs by *in vitro* transcription. Based on T7 RNA polymerase (RNAP)-catalyzed transcription reactions, we established two routes for the *in vitro* preparation of sugar-capped RNA sequences.

A first synthesis route for sugar-capped RNA includes *in vitro* transcription with T7 RNAP from a G-initiating DNA template in the presence of sugar-capped dinucleotide and guanosine triphosphate (GTP), as recently shown in the work of *Depaix et al.* (40). As transcription initiation can occur with both, sugar dinucleotide (acting as NCIN) and GTP, this reaction typically yields a mixture of sugar-capped RNA and 5’-triphosphate RNA (ppp-RNA) (**Figure 2 A**). We used a template encoding a 79mer RNA carrying two consecutive 2’-*O*-methyl nucleotides at the 5’-end of the template strand to improve 3’-homogeneity (44), and the NCIN (6 mM) in 10-fold excess over GTP (0.6 mM) (40). Thereby, we roughly obtained 1:1 or 2:1 mixtures of 79mer ppp-RNA (GTP-initiated) and sugar-capped 80mer RNA (NCIN-initiated) for compounds **4** and **5** or **6** and **7**, respectively (**Figure 2 B, S10**). To reliably determine the incorporation efficiencies with each of them, we cleaved the mixtures of sugar-capped and ppp-RNA using a sequence-specific 8-17 DNAzyme (45) and analyzed the fragments (NCIN-22mer / ppp-21mer and 5’-OH-58mer) by 20% PAGE analysis (**Figure 2 C, S11**). Densitometric band quantification gave incorporation efficiencies of 43%, 43%, 26% and 30% for the tested NCINs of GlcNAcppUpG **4**, GlcNAzppUpG **5**, EP-GlcNAcppUpG **6** and MurNAcppUpG **7**. The results for GlcNAc-capping levels of RNA are in agreement with the results of *Depaix et al.* (40). The lower capping efficiencies with the two latter compounds suggest that modifications at the C-3 position of GlcNAc might interfere with T7 RNAP incorporation. For GlcNazppUpG **5**, which contains a clickable azide group, we additionally confirmed our DNAzyme-based results by streptavidin shift assay of the sugar-capped 80mer RNA after strain-promoted azide-alkyne cycloaddition (SPAAC) with DBCO-biotin (**Figure 2 D, S12**). Quantification of observed bands shifted due to streptavidin binding revealed a capping efficiency of 48±2%.

**Figure 2:**
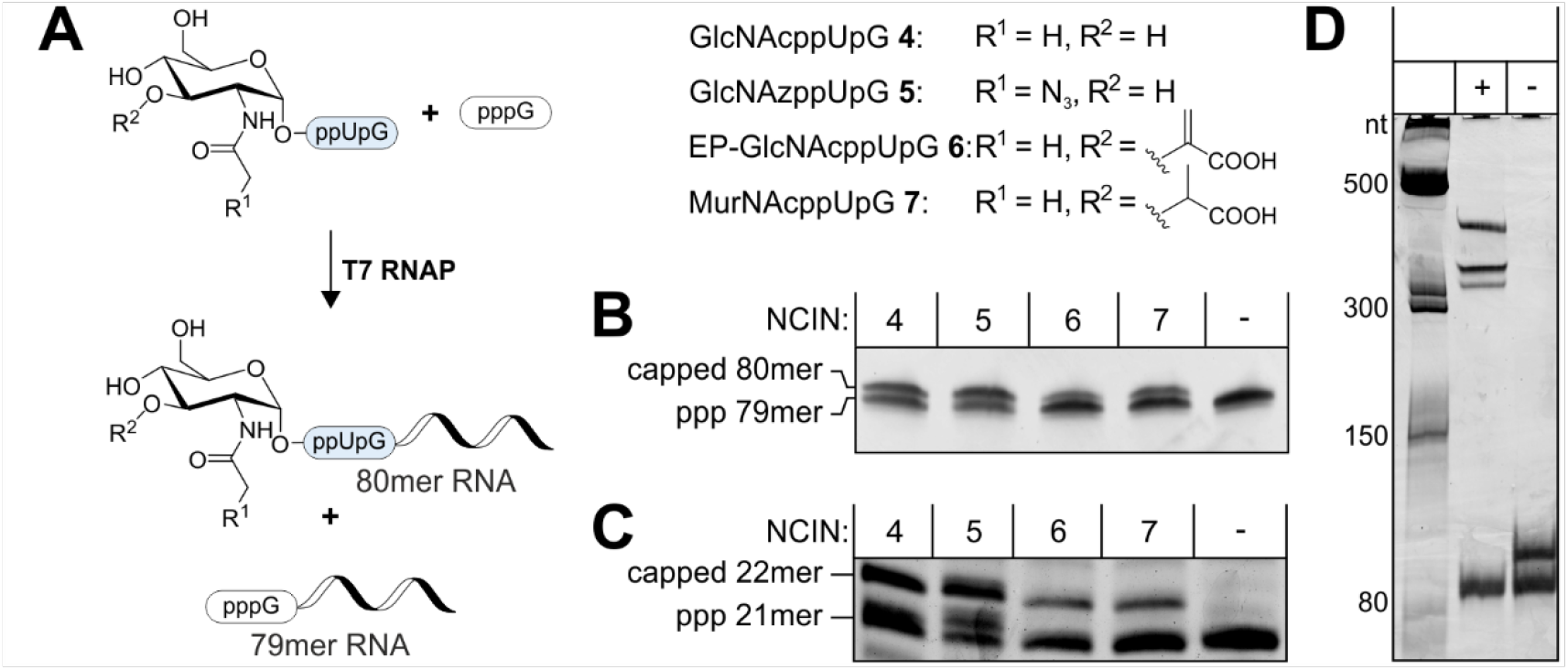
Incorporation of the NCINs GlcNAcppUpG **4**, GlcNAzppUpG **5**, EP-GlcNAcppUpG **6** or MurNAcppUpG **7** into RNA by T7 RNAP using a G-initiating template coding for a 79mer RNA and a 10-fold excess of NCIN over GTP. **A:** The NCIN competes for transcription initiation with GTP leading to a mixture of a capped 80mer RNA and ppp-79mer RNA. **B:** Section of a 10% PAGE analysis of PAGE-purified IVT products showing the transcribed RNA product. Compared to the negative control, in which no NCIN was present, an additional band can be observed in all samples. **C:** Determination of the capping efficiency by 8-17 DNAzyme cleavage of the capped RNA / ppp-RNA mixture to smaller capped 22mer / ppp-21mer fragments, separable in a 20% PAGE analysis. Densitometric band quantification revealed capping efficiencies of 43% for GlcNAcppUpG **4**, 43% for GlcNAzppUpG **5**, 26% for EP-GlcNAcppUpG **6** and 30% for MurNAcppUpG **7. D:** Determination of the capping efficiency of GlcNAzppUpG **5** by biotin-DBCO conjugation of capped 80mer RNA followed by a streptavidin shift assay. The 10% native PAGE analysis shows that due to streptavidin binding, 48±2% of RNA (n = 3, see **Figure S12**) was retarded. Multiple shifted bands were observed as streptavidin can bind multiple biotin moieties. Polyacrylamide gels were stained with SYBR Gold.

We also established a second synthesis route for sugar-capped RNA, which involves the use of a short DNA template containing a T7 class III promoter region and a single G at the transcription start site (+1 position) (**Figure 3 A, B**). In the absence of GTP, transcription can only start with the administered sugar dinucleotide and will continue until the next G within the template (in our case at the +11 position). Without the competing GTP during transcription initiation, short RNAs produced by this approach are quantitatively capped with the respective sugar dinucleotide. The full-length, sugar-capped 11mer RNA (counting the U overhang of the NCIN) could be readily purified by RP-HPLC and its identity be confirmed by ESI-MS (**Figure 3 C, S13, S14**). These short RNAs can find use in the screening of cap-binding or cap-modifying enzymes. To gain access to longer RNAs, we implemented a splinted ligation technique with our sugar-capped 11mer RNAs and 5’-monophosphate-RNA (p-RNA) using T4 DNA ligase. By ligation to 68mer p-RNA, we efficiently generated 5’-sugar-modified 79mer RNA from 11mer RNAs capped with compounds **4, 5, 6** and **7**, which was confirmed by PAGE analysis (**Figure 3C**). Therefore, the combination of short sugar-capped RNA in the absence of GTP with subsequent splinted ligation to 5’-p-RNA allows for the preparation of pure 5’-sugar-capped RNA of variable length and sequence.

**Figure 3:**
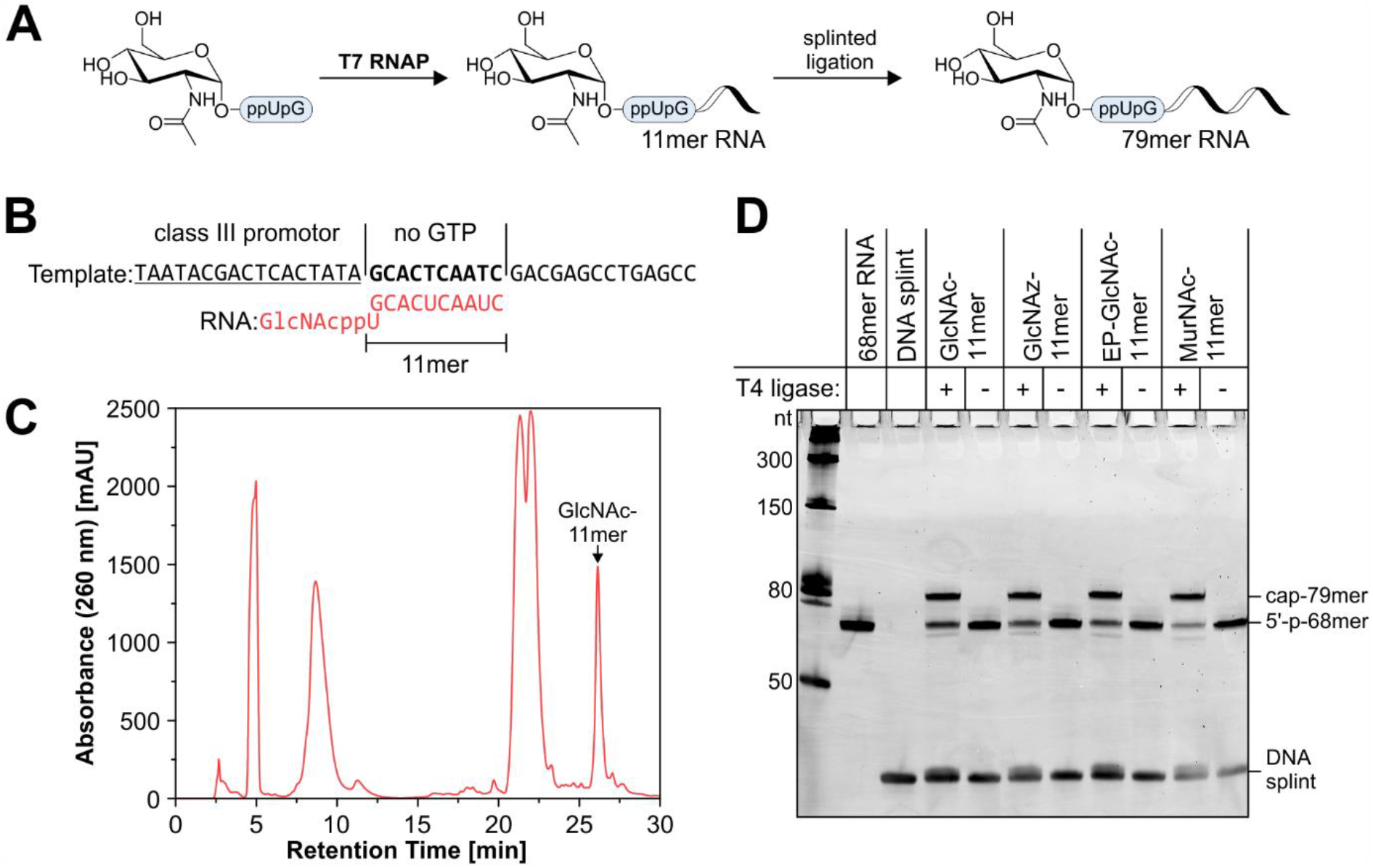
Synthesis of pure, sugar-capped RNA. **A:** Synthetic scheme exemplified for GlcNAcppUpG **4**. In the absence of GTP, transcription can only begin with the NCIN and stops at a position where a second G would be incorporated, leading to short capped 11mer RNAs, which were purified by RP-HPLC. A splinted ligation to a 5’-monophosphate-68mer RNA was used to elongate pure capped 11mers, thereby generating capped 79mer RNA. **B:** The utilized IVT template codes for G at position +1 and a second G at position +11 leading, in the presence of an NCIN, to capped 11mer RNA. **C:** HPLC chromatogram of GlcNAc-11mer RNA purification showing efficient separation of the desired RNA product from excess NTPs, dinucleotide, and abortive transcription products. **D:** Ligation reactions of pure capped 11mer RNA and 5’-monophosphate-68mer RNA analyzed by 10% PAGE. All capped 11mers could be ligated to the 68mer RNA, generating capped 79mer RNA. The polyacrylamide gel was stained with SYBR Gold.

### MurA, MurB and MurC can process sugar-capped RNA *in vitro*

The enzymes of the cytoplasmic murein biosynthetic pathway in bacteria modify the 3-OH group of UDP-GlcNAc in a sequential manner (46). In our study, we focused on the initial three enzymes of this pathway, which are MurA, MurB and MurC. MurA transfers an enolpyruvyl (EP) group from phosphoenol pyruvate (PEP) to the 3-OH of the GlcNAc moiety leading to UDP-GlcNAc-EP (47). MurB reduces the EP group through the consumption of NADPH (41) to generate UDP-*N*-acetyl muramic acid (UDP-MurNAc). Then, the UDP-*N*-acetylmuramyl:L-alanine ligase MurC preferably transfers L-alanine (Ala) to the nucleotide sugar (48,49). Following this first amino acid transfer, the murein biosynthetic pathway introduces a total of 5 amino acids to the 3-position of the sugar moiety to finally yield UDP-MurNAc-pentapeptide (46). Besides L-alanine, MurC is known to accept a variety of alanine derivatives for ligation to UDP-MurNAc, including some that contain a bioorthogonal handle (43,48). This makes Mur enzymes potentially interesting candidates for the chemo-enzymatic labeling of UDP-GlcNAc-RNA.

As indicated by our successful synthesis of sugar dinucleotides **6** and **7** from **4** using MurA and MurB (**Figure 1**), we hypothesized that Mur enzymes might generally accept 3’-elongated versions of their natural mononucleotide substrates. Therefore, we further tested the acceptance of our enzymatically synthesized sugar dinucleotide **7** by MurC using RP-HPLC and ESI-MS (**Figure S15**). Starting from MurNAcppUpG **7**, we observed quantitative conversion of the dinucleotide substrate to Ala-MurNAcppUpG **8** in a reaction setup with MurC and its natural substrate L-alanine (**Figure S15 A, B**). In addition, MurC was also able to attach unnatural amino acids like L-3-azidoalanine (Azidoala), L-propagylglycine (Propargylgly) and L-3-chloroalanine (Chloroala) to **7**, generating Azidoala-MurNAcppUpG **9**, Propargylgly-MurNAcppUpG **10** and Chloroala-MurNAcppUpG **11** in yields of roughly 91%, 97% and 96% (**Figure S15 A, C-D**). However, since the K_m_ values for some of these derivatives were reported to be higher than that of native Ala (43), these reactions were performed with 10-fold increased amino acid concentrations.

Our azide-labeled GlcNAc-derivative GlcNAzppUpG **5** was also readily converted to EP-GlcNAzppUpG **12** by MurA (**Figure S16 A, B**), with a yield of 89%, and to MurNAzppupG **13** in a one-pot reaction of MurA and MurB, with a yield of 27% (**Figure S16 C, D**).

With this, our work provides, in combination with our chemical synthesis (**Figure 1**), chemo-enzymatic access to a whole set of sugar-capped dinucleotides, including one literature-known compound (40) and nine novel sugar dinucleotide molecules (**Figure S17 A, B**). Strikingly, we found that all tested Mur enzymes accepted the 3’-elongated dinucleotide derivatives of their natural substrates *in vitro* and MurC could be used to attach unnatural amino acids to capped dinucleotides, encouraging a general synthetic use of Mur enzymes.

Comparing the reaction rate with Mur enzymes, we found a different behavior for MurA, MurB and MurC when exchanging their natural mononucleotide substrates with sugar-capped dinucleotides: With lowered enzyme concentrations, MurA reactions were monitored *via* the stoichiometric generation of orthophosphate by a malachite green phosphate assay. While UDP-GlcNAc was nearly completely converted after 45 min, only 10% conversion was observed for GlcNAcppUpG **4** (**Figure S18 A**) and the initial rate was reduced by around three orders of magnitude. However, the MurA-catalyzed reaction with **4** went to completion, when increasing the enzyme concentration and extending the reaction time (**Figure S19**). In contrast to MurA, MurB converted natural substrate UDP-GlcNAc-EP **14** and sugar-capped dinucleotide EP-GlcNAcppUpG **6** at a similar rate, which was measured by the decrease in absorbance at 340 nm due to stoichiometric NADPH consumption (**Figure S18 B**). Lastly, the reaction kinetics of MurC were examined by RP-HPLC, which revealed an around 400-fold reduction in the initial rate for the conversion of MurNAcppUpG **7**, compared to the native mononucleotide substrate UDP-MurNAc **15** in a reaction setup with L-alanine (**Figure S18 C**). In conclusion, the reactivity of sugar-capped dinucleotides with the tested Mur enzymes was partially lowered, but all dinucleotides were accepted and near-complete conversion could be achieved.

After having established the acceptance of dinucleotide substrates, we next were interested if MurA, MurB and MurC would also act as sugar-capped RNA-modifying enzymes (**Figure 4 A**). We first tested whether MurA and MurB can modify GlcNAc-11mer RNA in a one-pot reaction. Analysis of the reaction mixture by RP-HPLC and ESI-MS (**Figure 4 B**) revealed a substantial amount of MurNAc-11mer RNA, indicating that MurA and MurB both accepted GlcNAc-11mer RNA. The intermediate EP-GlcNAc-11mer RNA-product of the MurA reaction was not detected, likely due to the fast consumption by MurB, which showed faster kinetics with 3’-extended substrates in comparison to MurA. We further confirmed the acceptance of MurNAc-11mer RNA by MurC, ligating Ala or Propargylgly to the sugar-capped RNA in an almost quantitative manner (**Figure 4 C, S20 A, B**), while with 3-azidoala, the amino acid-modified MurNAc-11mer RNA was only detected to around 68% (**Figure S20 A, C-D)**.

**Figure 4:**
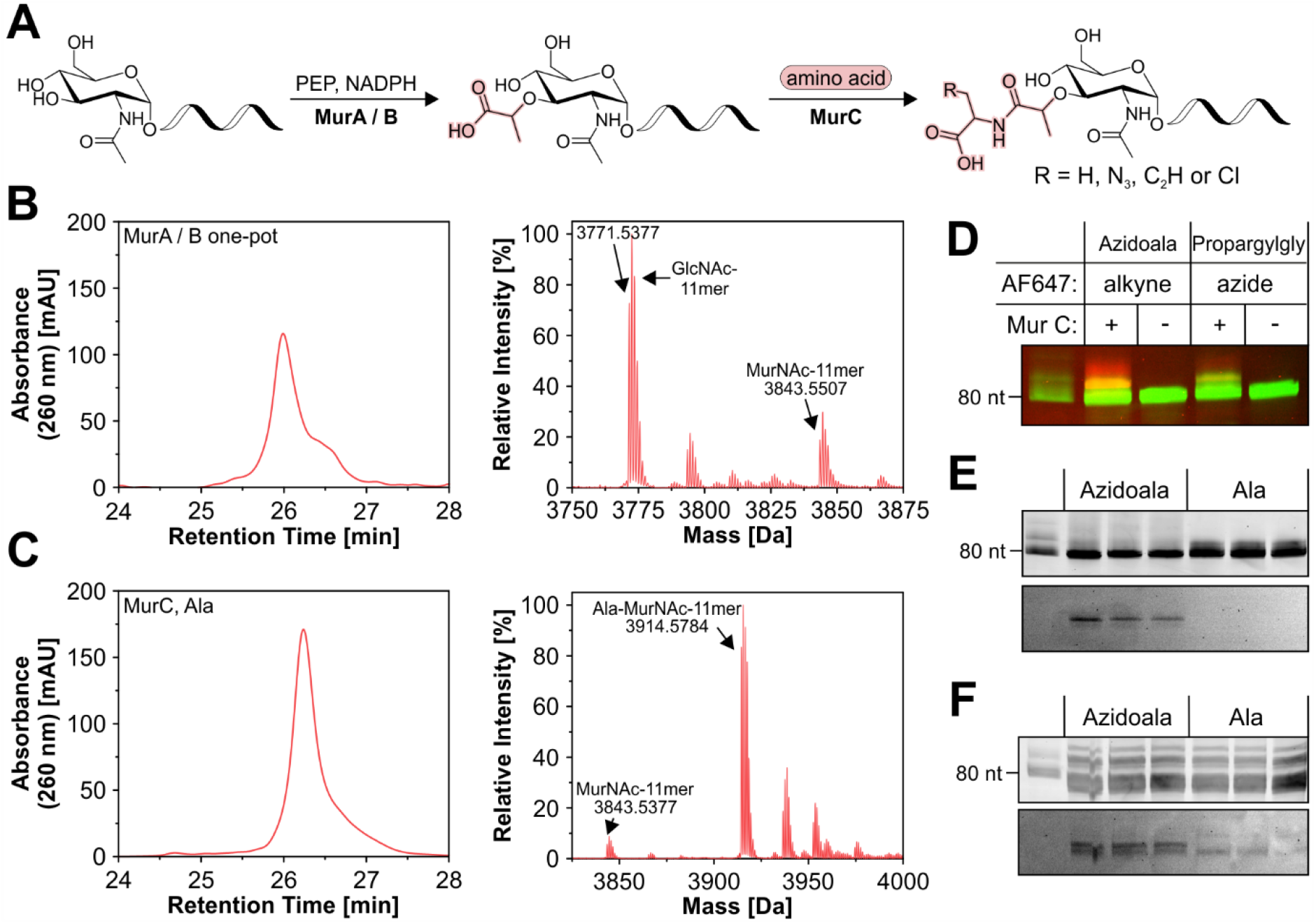
Processing of UDP-sugar RNA by Mur enzymes. **A:** GlcNAc-11mer RNA was converted by MurA and MurB in a one-pot reaction and MurNAc-11mer RNA was converted by MurC using different amino acids. **B:** RP-HPLC purification after incubating GlcNAc-11mer RNA with MurA and MurB showing capped-11mer substrate / product peak (left). The ESI-MS analysis (right) demonstrated the presence of a substantial amount of MurNAc-11mer RNA product and remaining substrate (Deconvoluted, Product [M]_calc_: 3843.5233). **C:** RP-HPLC purification after incubating MurNAc-11mer RNA with MurC and alanine showing substrate / product peaks (left). The ESI-MS analysis (right) demonstrated almost quantitative conversion to Ala-MurNAc-11mer RNA (Deconvoluted, Product [M]_calc_: 3914.5605). **D:** Section of a 10% PAGE analysis of a MurNAc-80mer / ppp-79mer RNA mixture processed by MurC in the presence of 3-azidoalanine or propargylglycine followed by CuAAC to AF647. Negative controls did not contain MurC during the enzymatic reaction. Overlay of SYBR Gold (green) and AF647 (red). **E:** Section of a 10% PAGE analysis of MurC reactions with 3-azidoalanine or alanine and MurNAc-80mer / triphosphate-79mer RNA mixture in triplicate, followed by SPAAC to DBCO-AF647 and visualized by SYBR Gold stain (top) or AF647 fluorescence readout (bottom). **F:** Section of a 10% PAGE analysis of MurC reactions with 3-azidoalanine or alanine and MurNAc-80mer / triphosphate-79mer RNA mixture spiked into *E. coli* total RNA (30 ng in 1 µg) in triplicates followed by SPAAC to DBCO-AF647 and visualized by SYBR Gold stain (top) or AF647 fluorescence readout (bottom).

We next investigated the modification of MurNAc-80mer RNA in the mixture with ppp-79mer RNA using MurC in combination with Propargylgly or Azidoala. Modification of RNA by MurC was visualized by subsequent fluorophore conjugation. As shown in the PAGE analysis, MurNAc-80mer RNA can be labeled using both amino acids and no labeling was observed in negative controls where RNA was incubated with all reagents in the absence of MurC (**Figure 4 D, S21**). In a second step, we tested the labeling of the MurNAc-80mer / ppp-79mer RNA mixture alone and spiked it into *E. coli* total RNA (**Figure 4 E, S22, S23**), using Azidoala in the MurC reaction followed by SPAAC to DBCO-AF647. Again, AF647-labeled RNA was detected only in the samples but not in the negative controls, in which Ala was used instead of Azidoala. In total RNA, some background is evident, but the samples show a clearly distinguishable band with the expected mobility. These results demonstrate that MurC can be utilized for the reproducible chemo-enzymatic labeling of UDP-MurNAc-RNA, even in complex mixtures like total RNA.

### Human Nudt5 decaps UDP-GlcNAc- and UDP-GlcNAz-RNA *in vitro*

To find a decapping enzyme capable of cleaving UDP-sugar-caps of RNA, we tested members of the Nudix (Nucleotide diphosphate X) hydrolase family. Nudix enzymes cleave phosphoanhydride bonds between a nucleotide and a moiety X, which can be a nucleotide or another biomolecule such as a sugar (50). While some Nudix enzymes cleave only mononucleotide metabolites, others have been identified to also act in RNA decapping (19-21). The human Nudix enzyme Nudt5 (hNudt5) and its bacterial counterpart NudF from *E. coli* are known mainly for the hydrolysis of ADP-sugars - particularly ADP-ribose, -mannose and -glucose (51-54). However, hNudt5 was reported to cleave several other substrates with greatly reduced activity, including the mononucleotide UDP-GlcNAc (54), and we were interested, if it would process GlcNAc-capped RNA in a manner as visualized in **Figure 5 A**.

**Figure 5:**
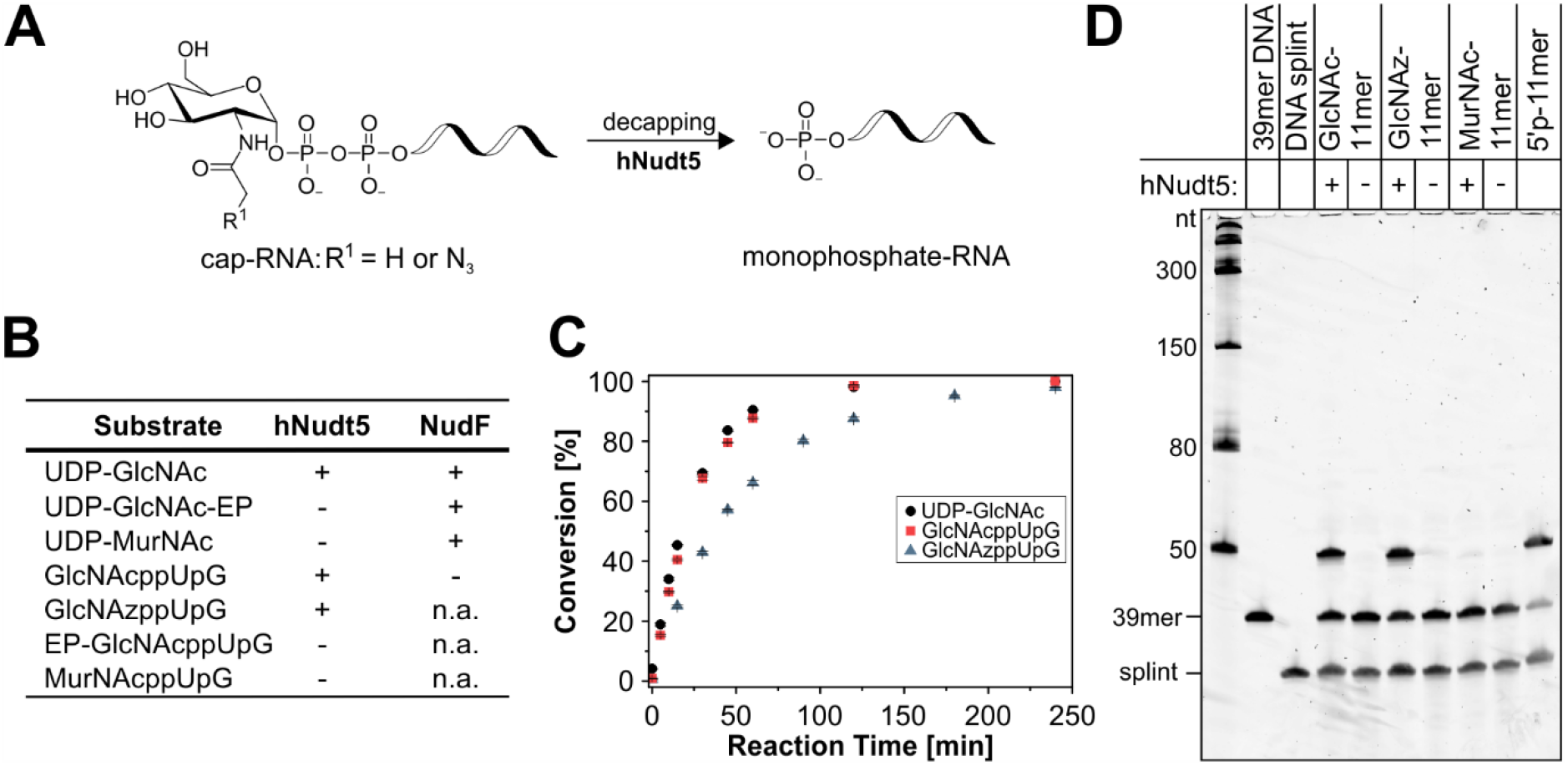
Cleavage of UDP-sugars, sugar-capped dinucleotides and sugar-capped RNA by the Nudix hydrolases hNudt5 and NudF. **A:** The Nudix enzyme hNudt5 was found to decap UDP-GlcNAc- and UDP-GlcNAz-RNA, leading to 5’-monophosphate-RNA by hydrolyzing the pyrophosphate bridge to the sugar cap. **B:** Mononucleotide and dinucleotide substrates tested and the ability of hNudt5 and NudF to hydrolyze them. **C:** Conversion of UDP-GlcNAc, GlcNAcppUpG **4** and GlcNAzppUpG **5** by hNudt5 (n = 3, data points show mean ± S.D.). Hydrolysis was analyzed by RP-HPLC. **D:** 10% PAGE analysis of a PABLO assay conducted with hNudt5-treated sugar-capped 11mer RNA (SYBR Gold stain). If the enzyme hydrolyzes the cap, generated 5’-monophosphate-RNA can be ligated to a 39mer DNA leading to a chimeric 50mer. A 50-nucleotide long ligation product can only be detected for hNudt5-treated GlcNAc-11mer and GlcNAz-11mer RNA as well as for 5’-monophosphate-11mer RNA positive controls. The experiment was repeated in triplicate using GlcNAc-11mer RNA and GlcNAz-11mer RNA (**Figure S27**).

We started our investigation at the mononucleotide level and tested the capabilities of NudF and hNudt5 in hydrolyzing the mononucleotides UDP-GlcNAc, UDP-GlcNAc-EP **14** and UDP-MurNAc **15**, which was analyzed by RP-HPLC and ESI-MS. Analysis of the reaction mixtures revealed that both enzymes cleaved the pyrophosphate linkage between the sugar and the nucleotide moiety of UDP-GlcNAc (**Figure S24**). While NudF was able to cleave UDP-GlcNAc-EP and UDP-MurNAc with reduced efficiency, the 3-OH modification of the GlcNAc moiety seemed to completely impede cleavage by hNudt5 (**Figure 5 B, S24**).

For the cleavage of capped dinucleotides (**Figure S25**), we detected the complete inability of NudF to decap even the dinucleotide closest to its natural substrate (GlcNAcppUpG **4**), while hNudt5 hydrolyzed both, GlcNAcppUpG **4** and GlcNAzppUpG **5**, but not the MurA- and MurB-products EP-GlcNAcppUpG **6** and MurNAcppUpG **7** (**Figure 5 B, S25**). Comparing the conversion of mononucleotide and dinucleotide cleavage by hNudt5, we found that GlcNAcppUpG **4** was hydrolyzed almost as fast as UDP-GlcNAc, reaching 98% conversion after 2 h, while GlcNAzppUpG **5** was converted slightly slower (**Figure 5 C**).

Consequently, we also confirmed the ability of hNudt5 in decapping reactions of sugar-capped 11mer RNA. As GlcNAc-capping of RNA did not result in a clear shift in PAGE or APB-PAGE analysis, we used an adapted version of the phosphorylation assay by ligation of oligonucleotides (PABLO) (55) to visualize the 5’-monophosphate-RNA generated during the decapping process (**Figure 5 D**). For this purpose, the RNA was incubated with hNudt5 and in a second step, a splinted ligation of the generated 5’-monophosphate-RNA to a 39mer DNA oligonucleotide was conducted. The resulting chimeric 50mer oligonucleotide could be easily distinguished on a PAGE gel. 50mers were detected for GlcNAc-11mer and GlcNAz-11mer RNA (and for the synthetic 5’-monophosphate-RNA positive control) and no additional band was detected for MurNAc-11mer RNA and in negative controls in which hNudt5 or the T4 DNA ligase (**Figure 5 D, S26**) were absent during the respective steps. We repeated the experiment for GlcNAc-11mer and GlcNAz-11mer RNA in triplicate (**Figure S27 A, B**) confirming the reliable decapping by hNudt5.

## DISCUSSION

Using liquid chromatography-mass spectrometry (LC-MS) techniques after digestion of total RNA, UDP-GlcNAc modification of RNA has been confirmed to occur at significant levels in a variety of organisms (3). Despite this apparent abundance, UDP-GlcNAc-caps are still poorly understood. In our opinion, this is partly due to the challenging *in vitro* synthesis of UDP-GlcNAc-RNA, which complicates the search for (e.g., enzymatic) tools for their analysis and, hence, the design and evaluation of techniques for the identification of GlcNAc-capped RNA sequences. In this work, we have presented chemical and enzymatic synthesis routes for one known GlcNAc-capped UpG dinucleotide and several novel dinucleotides capped with GlcNAc derivatives. These sugar-capped dinucleotides allowed for the *ab initio* incorporation of sugar-caps into RNA using T7 RNA polymerase. In addition to a preparation approach leading to a mixture of RNA with a 5’-sugar-cap and 5’-triphosphate, we presented an efficient way for the generation of pure GlcNAc- and GlcNAc derivative-capped RNAs: the highlighted combination of T7 RNAP-catalyzed *in vitro* transcription of short RNA initiating with our sugar-capped dinucleotides with subsequent splinted ligation to 5’-monophosphate-RNA allows for the generation of pure sugar-capped RNAs of varying length and sequence.

Our approaches made sugar-capped dinucleotides and RNA readily accessible for the extensive study of possible cap-processing enzymes: In MurA, MurB and MurC, as well as hNudt5, we were able to identify what, to our knowledge, are the first described modifying and decapping enzymes of GlcNAc-capped RNA.

For MurA, MurB and MurC, the initial enzymes of the murein biosynthesis pathway, we demonstrated the acceptance of not only sugar-capped dinucleotides but also sugar-capped RNAs, which enabled us to generate cell wall precursor-capped RNAs post-transcriptionally. This finding first let us speculate about the existence of cell wall precursor-capped RNA *in vivo*. The *ab initio* capping of RNA with UDP-GlcNAc as NCIN was demonstrated *in vitro* (27) and its intracellular concentration in *E. coli* (28) exceeds the K_m_ value determined for the bacterial RNAP (27). This suggests *ab initio* capping to be a major source of the UDP-GlcNAc-modification levels or RNA determined in several organisms (3). The intracellular concentrations for cell wall precursors, however, appear to be one to two orders of magnitude lower (56,57), making their *ab initio* incorporation into RNA less likely. In addition, the only cell wall precursor present in higher concentration, UDP-GlcNAc-pentapeptide (56), was described to not be incorporated into RNA by the *E. coli* wild-type RNAP *in vitro* (27).

Our findings of Mur enzymes readily processing sugar-capped RNAs *in vitro* point towards a possible post-transcriptional conversion of UDP-GlcNAc-caps to cell wall precursor-capped RNA. However, comparing the initial rate of mono- and dinucleotide substrates of MurA-C demonstrated a reduced rate for MurA and MurC, while MurB seemed to be more permissive. The different discrimination between both substrates by the Mur enzymes can be explained based on molecular structures. A crystal structure of MurA in complex with UDP-GlcNAc (58) shows a water-filled channel between the ribose and the protein surface that could allow the accommodation of 3’-elongated UDP-GlcNAc derivatives (**Figure S28 A**). However, there is also a hydrogen bond between a valine backbone and the 3’-OH group of the ribose moiety. This 3’-OH group has become part of a phosphodiester bond in the dinucleotide substrate, which may explain its slower conversion. Interactions with the uridine ribose moiety of the natural substrate are also indicated for MurC. A complex of *Haemophilus influenza* MurC with UDP-MurNAc **15** showed hydrogen bonding between both hydroxyl groups of the ribose part and a single aspartate side chain (**Figure S28 C**) (59). Comparison with the *E. coli* MurC crystal structure suggests that this enzyme binds its substrate analogously (60). In contrast, MurB interacts with the uridine ribose moiety of its substrate UDP-GlcNAc-EP **14** only *via* its uracil ring (**Figure S28 B**) (61,62). The 3’-OH group of the ribose does not take part in bonding interactions and points out of the enzymatic active site, which explains the readily acceptance of 3’-extended substrates.

The significantly reduced reaction rate of 3’-elongated substrates by MurA, the first enzyme of the biosynthetic pathway, renders a possible function of Mur enzymes in post-transcriptional UDP-GlcNAc-RNA modification debatable. Nevertheless, we consider the Mur enzymes to be an interesting *in vitro* tool. Using enzymes and other substrates in excess enabled us to generate a multitude of previously inaccessible capped dinucleotides with high yields, as well as sugar-capped RNAs. In addition, we demonstrated the incorporation of bioorthogonal alkyne and azide handles to MurNAc-80mer RNA, allowing conjugation by copper-catalyzed and strain-promoted click chemistry. Therefore, the enzymes can be interesting for the chemo-enzymatic labeling of capped RNA and might aid researchers in the development of a chemo-enzymatic capture approach of UDP-GlcNAc-RNA. Even though this would require the conversion by three enzymes, the possibility to conduct one-pot reactions with Mur enzymes could make this approach practical. The availability of crystal structures in complex with their natural substrates can further be a starting point for engineering the processing capabilities and reaction kinetics with 3’-elongated substrates.

In addition, we identified the Nudix hydrolase hNudt5 as a decapping enzyme for GlcNAc-RNA, as it was able to act on UDP-GlcNAc-RNA *in vitro*, while the investigated bacterial counterpart, NudF, was found to only cleave mononucleotide substrates. As hNudt5 is much better in hydrolyzing ADP-sugars than UDP-sugars (53,54), the decapping activity towards UDP-GlcNAc-RNA might only be a niche role of hNudt5 *in vivo*. The enzyme-substrate interactions derived from crystal structures can explain the preference for ADP substrates and differences between hNudt5 and NudF in terms of accepting 3’-extended UDP-GlcNAc analogues: The structures of both enzymes in complex with ADP-ribose (**Figure S28 D and E**) show that the enzymes recognize N-1 and N-6 of the adenine ring (63,64). However, while NudF additionally forms a hydrogen bond with the 3’-OH group of the adenosyl ribose(63), hNudt5 does not directly interact at this position (64).

As hNudt5 is, to the best of our knowledge the first described enzyme capable of cleaving UDP-GlcNAc-caps, we would like to promote it as an interesting tool for *in vitro* applications. The potential use of hNudt5 in a decapping protocol, such as CapZyme-Seq (31), for the identification of GlcNAc-capped RNA sequences, could be especially interesting. As hNudt5 cleaves the GlcNAc-cap structure under the release of 5’-monophosphate-RNA, it could be readily implemented into the CapZyme-Seq approach (31). Additional adaptations might be necessary to cover the wider substrate field of hNudt5, such as NAD (**Figure S29 A** and Ref. (54)), which is a prominent noncanonical 5’-cap structure of RNA (3). However, UDP-GlcNAc is stable in the presence of well-studied decapping enzymes like the Nudix hydrolase NudC (**Figure S29 B**), which cleaves NAD- and dpCoA-caps (21,65). We would therefore suggest a combinatorial approach of decapping enzymes to deplete such RNA caps before addressing the GlcNAc modification for the specific detection of UDP-GlcNAc-RNAs following the CapZyme-Seq methodologies.

The decapping enzyme hNudt5 could further aid researchers aiming to identify UDP-GlcNAc-RNA following a metabolic labeling approach using azide derivatives of GlcNAc as a chemical reporter (29). In the process of metabolic labeling, an azide derivative of GlcNAc would be added to the investigated biological system and incorporated into glycoconjugates by its biosynthetic machinery. The activated biosynthetic intermediate UDP-GlcNAz could then be incorporated into RNA by *ab initio* capping (27). Negative controls are usually generated by growing cells in the absence of the supplemented azido sugar. These differences in growth conditions, however, might lead to changes in the transcriptome between the sample and control, thereby introducing a bias. Our identification of hNudt5 as being capable of cleaving azide-bearing GlcNAz caps can be a solution to this problem. The decapping enzyme can be used for the generation of negative controls by efficiently cleaving the azide-bearing cap structures before capturing GlcNAz-capped RNAs using copper-catalyzed or strain-promoted click chemistry, as adapted in a variety of NAD-RNA capture protocols, such as NAD captureSeq (5,6) and SPAAC-NAD-seq (8). The acceptance of GlcNAzppUpG **5** by MurA and B further indicates potential metabolic intersections, converting UDP-GlcNAz caps to azide-bearing cell wall precursor caps, if the latter exist *in vivo*. Our finding that hNudt5 is unable to hydrolyze cell wall precursor caps could additionally aid to distinguish UDP-GlcNAz caps from products of described metabolic intersections.

In summary, our work provides solutions for the need for efficient synthesis strategies of UDP-sugar-capped RNAs, and UDP-GlcNAc-RNAs in particular. We generated novel capped dinucleotides and RNA, which we successfully employed for the identification of previously unknown UDP-GlcNAc-cap and cell wall precursor cap-modifying activities of different enzymes from the Mur family, as well as a UDP-GlcNAc-RNA decapping activity of hNudt5. We expect these first additions of sugar-cap processing enzymes to the toolbox for the investigation of UDP-GlcNAc capping to aid in future research examining this, despite of its abundance, still poorly understood RNA modification.

## ACKNOWLEDGEMENT

We thank Dr. K. Höfer, Dr. F. Abele and Dr. A. Krause for providing *E. coli* BL21(DE3) containing Pet-28C(+) with genes of NudF, hNudt5, MurA and MurB. We further thank Dr. F. Abele for pioneering work of MurA and MurB with 3’ elongated substrates. We thank M. Kintzel for assistance in synthesis and H. Rudy and T. Timmermann for technical support.

## REFERENCES

1. Furuichi, Y. (2015) Discovery of m^7^G-cap in eukaryotic mRNAs. Proc. Jpn. Acad., Ser. B, Phys. Biol. Sci., 91, 394–409.

2. Kowtoniuk, W.E., Shen, Y.H., Heemstra, J.M., Agarwal, I. and Liu, D.R. (2009) A chemical screen for biological small molecule-RNA conjugates reveals CoA-linked RNA. Proc. Natl. Acad. Sci. U.S.A., 106, 7768–7773.

3. Wang, J., Chew, B.L.A., Lai, Y., Dong, H.P., Xu, L., Balamkundu, S., Cai, W.M., Cui, L., Liu, C.F., Fu, X.Y. et al. (2019) Quantifying the RNA cap epitranscriptome reveals novel caps in cellular and viral RNA. Nucleic Acids Res., 47, e130.

4. Chen, Y.G., Kowtoniuk, W.E., Agarwal, I., Shen, Y.H. and Liu, D.R. (2009) LC/MS analysis of cellular RNA reveals NAD-linked RNA. Nat. Chem. Biol., 5, 879–881.

5. Cahová, H., Winz, M.L., Höfer, K., Nübel, G. and Jäschke, A. (2015) NAD captureSeq indicates NAD as a bacterial cap for a subset of regulatory RNAs. Nature, 519, 374–377.

6. Winz, M.L., Cahová, H., Nübel, G., Frindert, J., Höfer, K. and Jäschke, A. (2017) Capture and sequencing of NAD-capped RNA sequences with NAD captureSeq. Nat. Protoc., 12, 122–149.

7. Zhang, H.L., Zhong, H., Zhang, S.D., Shao, X.J., Ni, M., Cai, Z.W., Chen, X.M. and Xia, Y.J. (2019) NAD tagSeq reveals that NAD^+^-capped RNAs are mostly produced from a large number of protein-coding genes in Arabidopsis. Proc. Natl. Acad. Sci. U.S.A., 116, 12072–12077.

8. Hu, H., Flynn, N., Zhang, H.L., You, C.J., Hang, R.L., Wang, X.F., Zhong, H., Chan, Z.L., Xia, Y.J. and Chen, X.M. (2021) SPAAC-NAD-seq, a sensitive and accurate method to profile NAD^+^-capped transcripts. Proc. Natl. Acad. Sci. U.S.A., 118, e2025595118.

9. Zhang, H.L., Zhong, H., Wang, X.F., Zhang, S.D., Shao, X.J., Hu, H., Yu, Z.L., Cai, Z.W., Chen, X.M. and Xia, Y.J. (2021) Use of NAD tagSeq II to identify growth phase-dependent alterations in E. coli RNA NAD^+^ capping. Proc. Natl. Acad. Sci. U.S.A., 118, e2026183118.

10. Niu, K.Y., Zhang, J.Y., Ge, S.W., Li, D., Sun, K.F., You, Y.N., Qiu, J.Q., Wang, K., Wang, X.T., Liu, R. et al. (2023) ONE-seq: epitranscriptome and gene-specific profiling of NAD-capped RNA. Nucleic Acids Res., 51, e12.

11. Sharma, S., Yang, J., Favate, J., Shah, P.M. and Kiledjian, M. (2023) NADcapPro and circNC: Methods for accurate profiling of NAD and non-canonical RNA caps in eukaryotes. Commun. Biol., 6, 406.

12. Frindert, J., Zhang, Y.Q., Nübel, G., Kahloon, M., Kolmar, L., Hotz-Wagenblatt, A., Burhenne, J., Haefeli, W.E. and Jäschke, A. (2018) Identification, biosynthesis, and decapping of NAD-capped RNAs in B. subtilis. Cell Rep., 24, 1890–1901.

13. Morales-Filloy, H.G., Zhang, Y.Q., Nübel, G., George, S.E., Korn, N., Wolz, C. and Jäschke, A. (2020) The 5 ‘ NAD cap of RNAIII modulates toxin production in Staphylococcus aureus isolates. J. Bacteriol., 202, e00591–19.

14. Walters, R.W., Matheny, T., Mizoue, L.S., Rao, B.S., Muhlrad, D. and Parker, R. (2017) Identification of NAD^+^ capped mRNAs in Saccharomyces cerevisiae. Proc. Natl. Acad. Sci. U.S.A., 114, 480–485.

15. Wang, Y., Li, S.F., Zhao, Y.H., You, C.J., Le, B., Gong, Z.Z., Mo, B.X., Xia, Y.J. and Chen, X.M. (2019) NAD^+^-capped RNAs are widespread in the Arabidopsis transcriptome and can probably be translated. Proc. Natl. Acad. Sci. U.S.A., 116, 12094–12102.

16. Jiao, X., Doamekpor, S.K., Bird, J.G., Nickels, B.E., Tong, L., Hart, R.P. and Kiledjian, M. (2017) 5’ End nicotinamide adenine dinucleotide cap in human cells promotes RNA decay through DXO-mediated deNADding. Cell, 168, 1015–1027.

17. Bird, J.G., Zhang, Y., Tian, Y., Panova, N., Barvik, I., Greene, L., Liu, M., Buckley, B., Krásný, L., Lee, J.K. et al. (2016) The mechanism of RNA 5’ capping with NAD^+^, NADH and desphospho-CoA. Nature, 535, 444–447.

18. Zhang, Y.Q., Kuster, D., Schmidt, T., Kirrmaier, D., Nübel, G., Ibberson, D., Benes, V., Hombauer, H., Knop, M. and Jäschke, A. (2020) Extensive 5 ‘-surveillance guards against non-canonical NAD-caps of nuclear mRNAs in yeast. Nat. Commun., 11, 5508.

19. Grudzien-Nogalska, E., Wu, Y.X., Jiao, X.F., Cui, H.J., Mateyak, M.K., Hart, R.P., Tong, L. and Kiledjian, M. (2019) Structural and mechanistic basis of mammalian Nudt12 RNA deNADding. Nat. Chem. Biol., 15, 575–582.

20. Sharma, S., Grudzien-Nogalska, E., Hamilton, K., Jiao, X.F., Yang, J., Tong, L. and Kiledjian, M. (2020) Mammalian Nudix proteins cleave nucleotide metabolite caps on RNAs. Nucleic Acids Res., 48, 6788–6798.

21. Höfer, K., Li, S.S., Abele, F., Frindert, J., Schlotthauer, J., Grawenhoff, J., Du, J.M., Patel, D.J. and Jäschke, A. (2016) Structure and function of the bacterial decapping enzyme NudC. Nat. Chem. Biol., 12, 730–734.

22. Sharma, S., Yang, J., Grudzien-Nogalska, E., Shivas, J., Kwan, K.Y. and Kiledjian, M. (2022) Xrn1 is a deNADding enzyme modulating mitochondrial NAD-capped RNA. Nat. Commun., 13, 889.

23. Huang, F. (2003) Efficient incorporation of CoA, NAD and FAD into RNA by in vitro transcription. Nucleic Acids Res., 31, e8.

24. Höfer, K., Abele, F., Schlotthauer, J. and Jäschke, A. (2016) Synthesis of 5’-NAD-capped RNA. Bioconjugate Chem., 27, 874–877.

25. Nübel, G., Sorgenfrei, F.A. and Jäschke, A. (2017) Boronate affinity electrophoresis for the purification and analysis of cofactor-modified RNAs. Methods, 117, 14–20.

26. Mikkola, S. (2020) Nucleotide sugars in chemistry and biology. Molecules, 25, 5755.

27. Julius, C. and Yuzenkova, Y. (2017) Bacterial RNA polymerase caps RNA with various cofactors and cell wall precursors. Nucleic Acids Res., 45, 8282–8290.

28. Bennett, B.D., Kimball, E.H., Gao, M., Osterhout, R., Van Dien, S.J. and Rabinowitz, J.D. (2009) Absolute metabolite concentrations and implied enzyme active site occupancy in Escherichia coli. Nat. Chem. Biol., 5, 593–599.

29. Laughlin, S.T. and Bertozzi, C.R. (2007) Metabolic labeling of glycans with azido sugars and subsequent glycan-profiling and visualization via Staudinger ligation. Nat. Protoc., 2, 2930–2944.

30. Flynn, R.A., Pedram, K., Malaker, S.A., Batista, P.J., Smith, B.A.H., Johnson, A.G., George, B.M., Majzoub, K., Villalta, P.W., Carette, J.E. et al. (2021) Small RNAs are modified with N-glycans and displayed on the surface of living cells. Cell, 184, 3109–3124.

31. Vvedenskaya, I.O., Bird, J.G., Zhang, Y.C., Zhang, Y., Jiao, X.F., Barvík, I., Krásný, L., Kiledjian, M., Taylor, D.M., Ebright, R.H. et al. (2018) CapZyme-Seq comprehensively defines promoter-sequence determinants for RNA 5’ capping with NAD^+^. Mol. Cell, 70, 553–564.

32. Sherwood, A.V., Rivera-Rangel, L.R., Ryberg, L.A., Larsen, H.S., Anker, K.M., Costa, R., Vågbø, C.B., Jakljevič, E., Pham, L.V., Fernandez-Antunez, C. et al. (2023) Hepatitis C virus RNA is 5’-capped with flavin adenine dinucleotide. Nature, 619, 811–818.

33. Kuzmine, I., Gottlieb, P.A. and Martin, C.T. (2003) Binding of the priming nucleotide in the initiation of transcription by T7 RNA polymerase. J. Biol. Chem., 278, 2819–2823.

34. Möhler, M., Höfer, K. and Jäschke, A. (2020) Synthesis of 5’-thiamine-capped RNA. Molecules, 25, 5492.

35. Martin, C.T. and Coleman, J.E. (1989) T7 RNA-polymerase does not interact with the 5’-phosphate of the initiating nucleotide. Biochemistry, 28, 2760–2762.

36. Seelig, B. and Jäschke, A. (1997) Site-specific modification of enzymatically synthesized RNA: Transcription initiation and Diels-Alder reaction. Tetrahedron Lett., 38, 7729–7732.

37. Samanta, A., Krause, A. and Jäschke, A. (2014) A modified dinucleotide for site-specific RNA-labelling by transcription priming and click chemistry. Chem. Commun., 50, 1313–1316.

38. Krause, A., Hertl, A., Muttach, F. and Jäschke, A. (2014) Phosphine-free Stille-Migita chemistry for the mild and orthogonal modification of DNA and RNA. Chem. Eur. J., 20, 16613–16619.

39. Pitulle, C., Kleineidam, R.G., Sproat, B. and Krupp, G. (1992) Initiator oligonucleotides for the combination of chemical and enzymatic RNA-synthesis. Gene, 112, 101–105.

40. Depaix, A., Grudzien-Nogalska, E., Fedorczyk, B., Kiledjian, M., Jemielity, J. and Kowalska, J. (2022) Preparation of RNAs with non-canonical 5’ ends using novel di- and trinucleotide reagents for co-transcriptional capping. Front. Mol. Biosci., 9, 854170.

41. Benson, T.E., Marquardt, J.L., Marquardt, A.C., Etzkorn, F.A. and Walsh, C.T. (1993) Overexpression, purification, and mechanistic study of UDP-N-acetylenolpyruvylglucosamine reductase. Biochemistry, 32, 2024–2030.

42. Mizyed, S., Oddone, A., Byczynski, B., Hughes, D.W. and Berti, P.J. (2005) UDP-N-acetylmuramic acid (UDP-MurNAc) is a potent inhibitor of MurA (enolpyruvyl-UDP-GlcNAc synthase). Biochemistry, 44, 4011–4017.

43. Emanuele, J.J., Jin, H.Y., Jacobson, B.L., Chang, C.Y.Y., Einspahr, H.M. and Villafranca, J.J. (1996) Kinetic and crystallographic studies of Escherichia coli UDP-N-acetylmuramate:L-alanine ligase. Protein Sci., 5, 2566–2574.

44. Kao, C., Zheng, M. and Rüdisser, S. (1999) A simple and efficient method to reduce nontemplated nucleotide addition at the 3’ terminus of RNAs transcribed by T7 RNA polymerase. RNA, 5, 1268–1272.

45. Santoro, S.W. and Joyce, G.F. (1997) A general purpose RNA-cleaving DNA enzyme. Proc. Natl. Acad. Sci. U.S.A., 94, 4262–4266.

46. El Zoeiby, A., Sanschagrin, F. and Levesque, R.C. (2003) Structure and function of the Mur enzymes: development of novel inhibitors. Mol. Microbiol., 47, 1–12.

47. Zemell, R.I. and Anwar, R.A. (1975) Mechanism of pyruvate-uridine diphospho-N-acetylglucosamine transferase - evidence for an enzyme-enolpyruvate intermediate. J. Biol. Chem., 250, 4959–4964.

48. Gubler, M., Appoldt, Y. and Keck, W. (1996) Overexpression, purification, and characterization of UDP-N-acetylmuramyl:L-alanine ligase from Escherichia coli. J. Bacteriol., 178, 906–910.

49. Liger, D., Masson, A., Blanot, D., Vanheijenoort, J. and Parquet, C. (1995) Over-production, purification and properties of the uridine-diphosphate-N-acetylmuramate:L-alanine ligase from Escherichia coli. Eur. J. Biochem., 230, 80–87.

50. McLennan, A.G. (2006) The Nudix hydrolase superfamily. Cell. Mol. Life. Sci., 63, 123–143.

51. Dunn, C.A., O’Handley, S.F., Frick, D.N. and Bessman, M.J. (1999) Studies on the ADP-ribose pyrophosphatase subfamily of the Nudix hydrolases and tentative identification of trgB, a gene associated with tellurite resistance. J. Biol. Chem., 274, 32318–32324.

52. Moreno-Bruna, B., Baroja-Fernández, E., Muñoz, F.J., Bastarrica-Berasategui, A., Zandueta-Criado, A., Rodriguez-López, M., Lasa, I., Akazawa, T. and Pozueta-Romero, J. (2001) Adenosine diphosphate sugar pyrophosphatase prevents glycogen biosynthesis in Escherichia coli. Proc. Natl. Acad. Sci. U.S.A., 98, 8128–8132.

53. Yang, H., Slupska, M.M., Wei, Y.F., Tai, J.H., Luther, W.M., Xia, Y.R., Shih, D.M., Chiang, J.H., Baikalov, C., Fitz-Gibbon, S. et al. (2000) Cloning and characterization of a new member of the Nudix hydrolases from human and mouse. J. Biol. Chem., 275, 8844–8853.

54. Gasmi, L., Cartwright, J.L. and McLennan, A.G. (1999) Cloning, expression and characterization of YSA1H, a human adenosine 5 ‘-diphosphosugar pyrophosphatase possessing a MutT motif. Biochem J., 344, 331–337.

55. Celesnik, H., Deana, A. and Belasco, J.G. (2008) PABLO analysis of RNA: 5’-phosphorylation state and 5’-end mapping. Methods Enzymol., 447, 83–98.

56. Mengin-Lecreulx, D., Flouret, B. and Van Heijenoort, J. (1982) Cytoplasmic steps of peptidoglycan synthesis in Escherichia coli. J. Bacteriol., 151, 1109–1117.

57. Mengin-Lecreulx, D., Flouret, B. and Van Heijenoort, J. (1983) Pool levels of UDP N-acetylglucosamine and UDP N-acetylglucosamine-enolpyruvate in Escherichia Coli and correlation with peptidoglycan synthesis. J. Bacteriol., 154, 1284–1290.

58. Skarzynski, T., Mistry, A., Wonacott, A., Hutchinson, S.E., Kelly, V.A. and Duncan, K. (1996) Structure of UDP-N-acetylglucosamine enolpyruvyl transferase, an enzyme essential for the synthesis of bacterial peptidoglycan, complexed with substrate UDP-N-acetylglucosamine and the drug fosfomycin. Structure, 4, 1465–1474.

59. Mol, C.D., Brooun, A., Dougan, D.R., Hilgers, M.T., Tari, L.W., Wijnands, R.A., Knuth, M.W., McRee, D.E. and Swanson, R.V. (2003) Crystal structures of active fully assembled substrate- and product-bound complexes of UDP-N-acetylmuramic acid:L-alanine ligase (MurC) from Haemophilus influenzae. J. Bacteriol., 185, 4152–4162.

60. Deva, T., Baker, E.N., Squire, C.J. and Smith, C.A. (2006) Structure of Escherichia coli UDP-N-acetylmuramoyl:L-alanine ligase (MurC). Acta Crystallogr. D, 62, 1466–1474.

61. Benson, T.E., Walsh, C.T. and Hogle, J.M. (1997) X-ray crystal structures of the S229A mutant and wild-type MurB in the presence of the substrate enolpyruvyl-UDP-N-acetylglucosamine at 1.8-Å resolution. Biochemistry, 36, 806–811.

62. Benson, T.E., Filman, D.J., Walsh, C.T. and Hogle, J.M. (1995) An enzyme-substrate complex involved in bacterial cell wall biosynthesis. Nat. Struct. Biol., 2, 644–653.

63. Gabelli, S.B., Bianchet, M.A., Bessman, M.J. and Amzel, L.M. (2001) The structure of ADP-ribose pyrophosphatase reveals the structural basis for the versatility of the Nudix family. Nat. Struct. Biol., 8, 467–472.

64. Zha, M., Zhong, C., Peng, Y., Hu, H. and Ding, J. (2006) Crystal structures of human NUDT5 reveal insights into the structural basis of the substrate specificity. J. Mol. Biol., 364, 1021–1033.

65. Zhou, W., Guan, Z.Y., Zhao, F., Ye, Y.G., Yang, F., Yin, P. and Zhang, D.L. (2021) Structural insights into dpCoA-RNA decapping by NudC. RNA Biol., 18, 244–253.

