## Supplementary_Material for "Identification and *in vitro* characterization of UDP-GlcNAc-RNA cap-modifying and decapping enzymes"

### Supplementary Information

#### 1 Supporting Tables and Figures

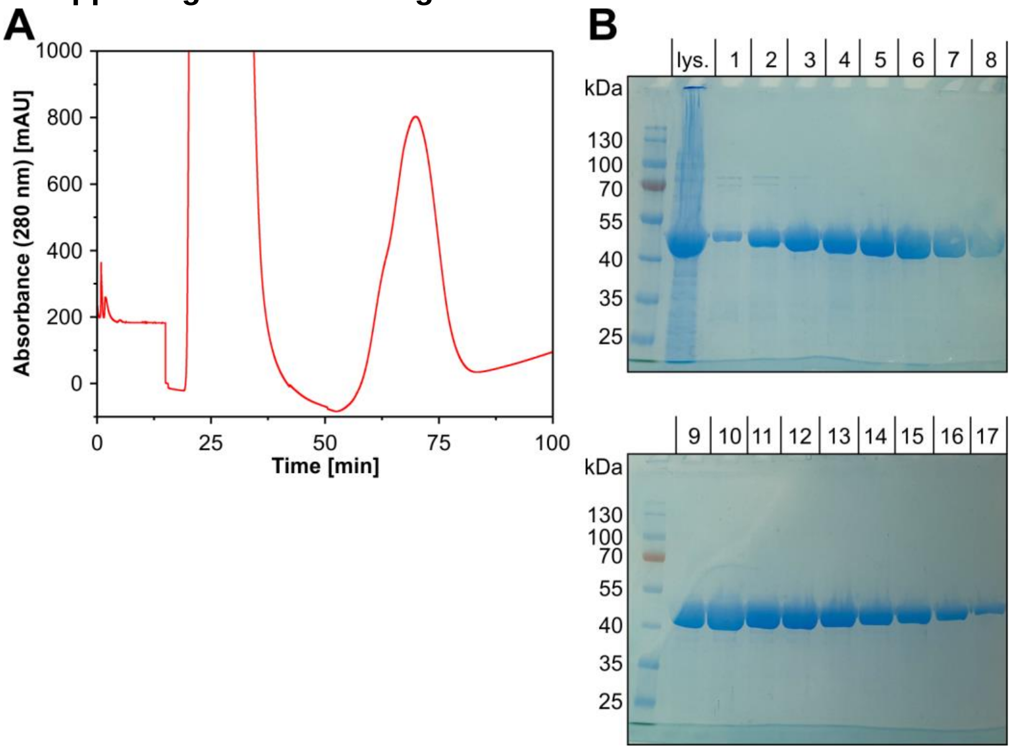

**Figure S 1:** Purification of MurA from *E. coli* BL21(DE3). **A:** Chromatogram of IMAC purification of the protein from cell lysate. **B:** 10% SDS-PAGE analysis of individual eluted fractions from IMAC stained with READYBLUE PROTEIN GEL STAIN. The protein size is in accordance with the literature (~44 kDa) (1). Lys.: Cell lysate.

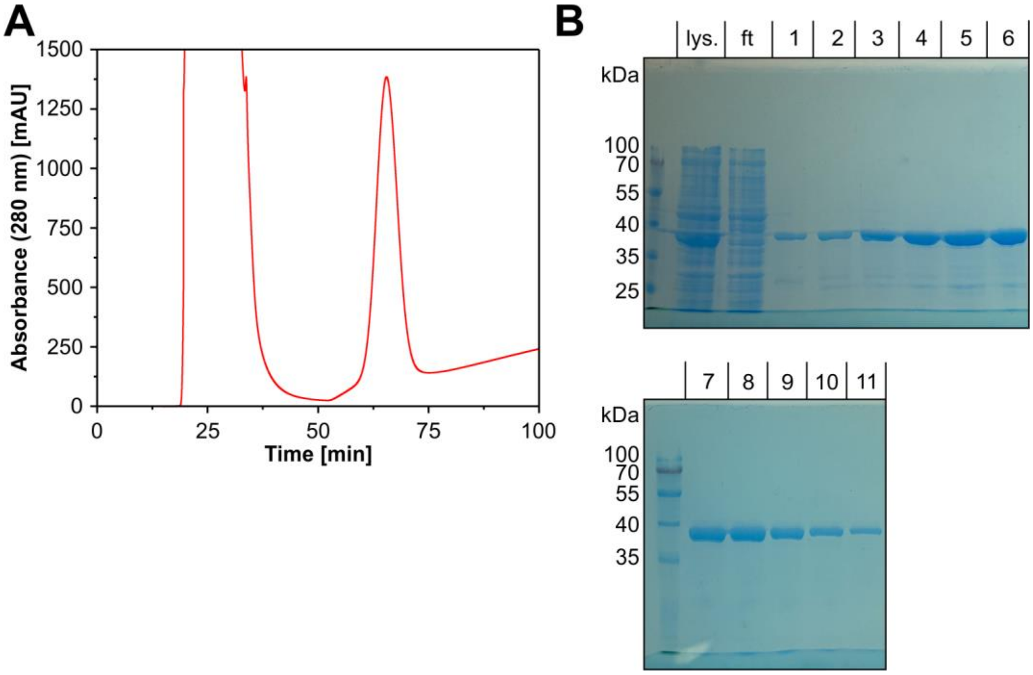

**Figure S 2:** Purification of MurB from *E. coli* BL21(DE3). **A:** Chromatogram of IMAC purification of the protein from cell lysate. **B:** 10% SDS-PAGE analysis of individual eluted fractions from IMAC stained with READYBLUE PROTEIN GEL STAIN. The protein size is in accordance with the literature (~38 kDa) (2). Lys.: Cell lysate; ft: Flow through.

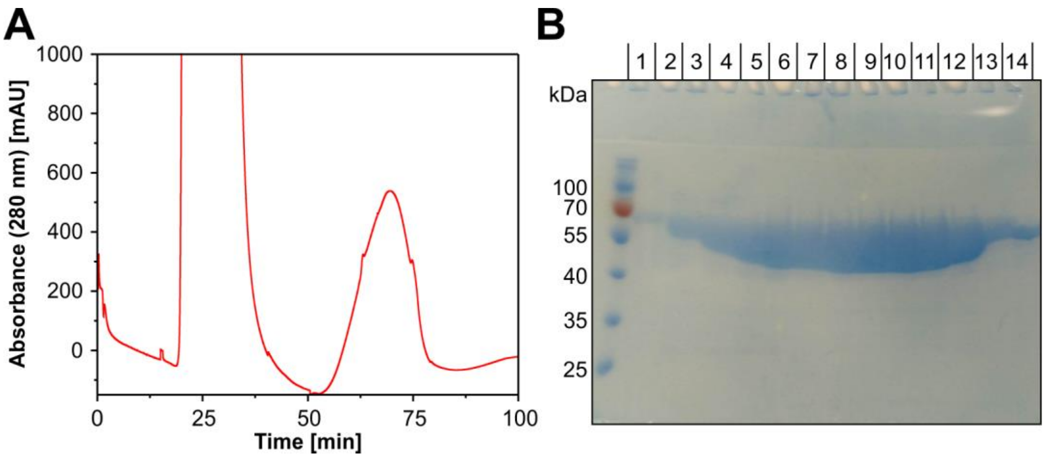

**Figure S 3:** Purification of MurC from *E. coli* BL21(DE3). **A:** Chromatogram of IMAC purification of the protein from cell lysate. **B:** 10% SDS-PAGE analysis of individual eluted fractions from IMAC stained with READYBLUE PROTEIN GEL STAIN. The protein size is in accordance with the literature (~54 kDa) (3).

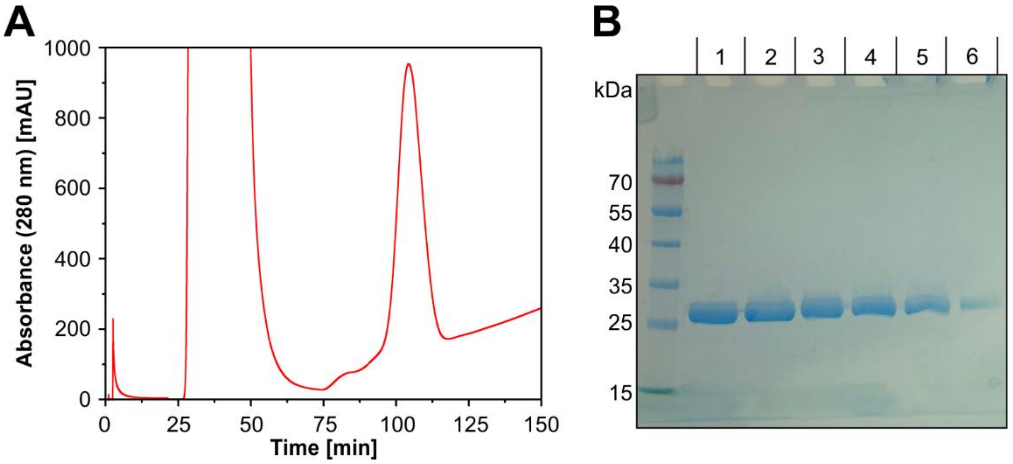

**Figure S 4:** Purification of NudF from *E. coli* BL21(DE3). **A:** Chromatogram of IMAC purification of the protein from cell lysate. **B:** 10% SDS-PAGE analysis of individual eluted fractions from IMAC stained with READYBLUE PROTEIN GEL STAIN. The protein size is in accordance with literature (~26 kDa) (4).

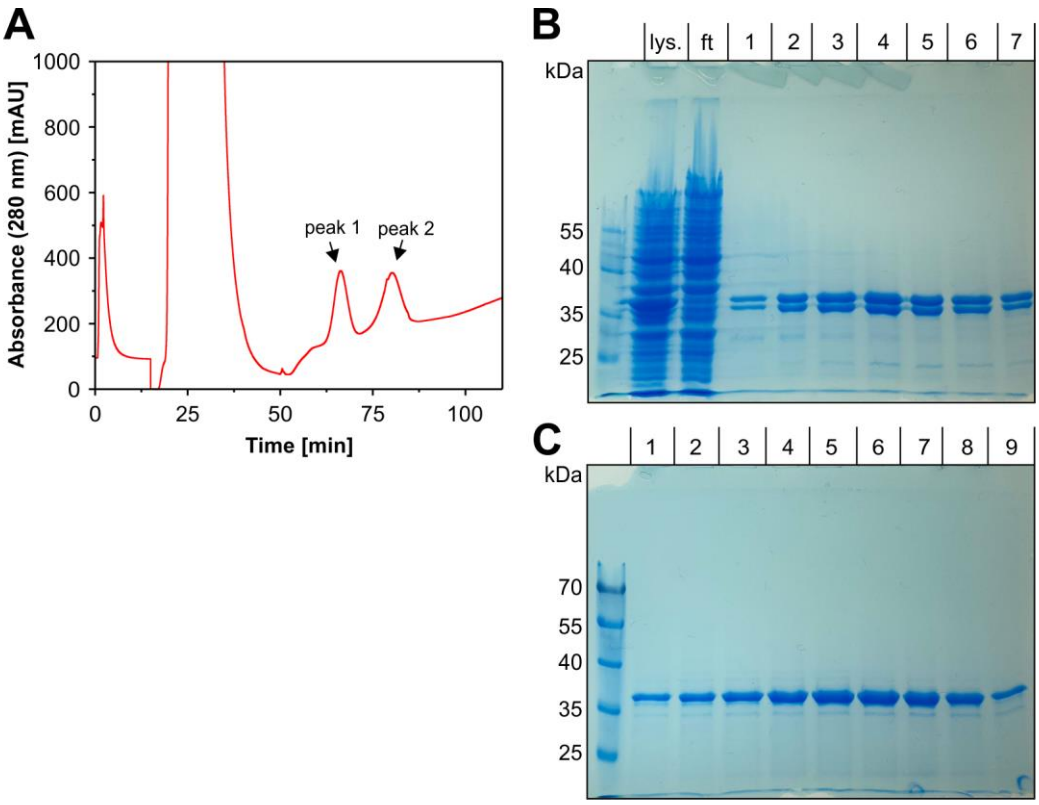

**Figure S 5:** Purification of recombinant human Nudt5 from *E. coli* BL21(DE3). **A:** Chromatogram of IMAC purification of the protein from cell lysate. **B and C:** 10% SDS-PAGE analysis of individual eluted fractions from IMAC stained with READYBLUE PROTEIN GEL STAIN. The chromatogram showed two peaks which both contained active protein (B shows fractions from peak 1; C shows fractions from peak 2). The larger molecular mass indicated by SDS-PAGE analysis compared to the theoretical value of 26 kDa is in accordance to the literature (5,6). Furthermore, the appearance of a doublet in SDS-PAGE analysis and unusual tight binding to $Ni^{2+}$-NTA has been reported (5). Lys.: Cell lysate; ft: Flow through.

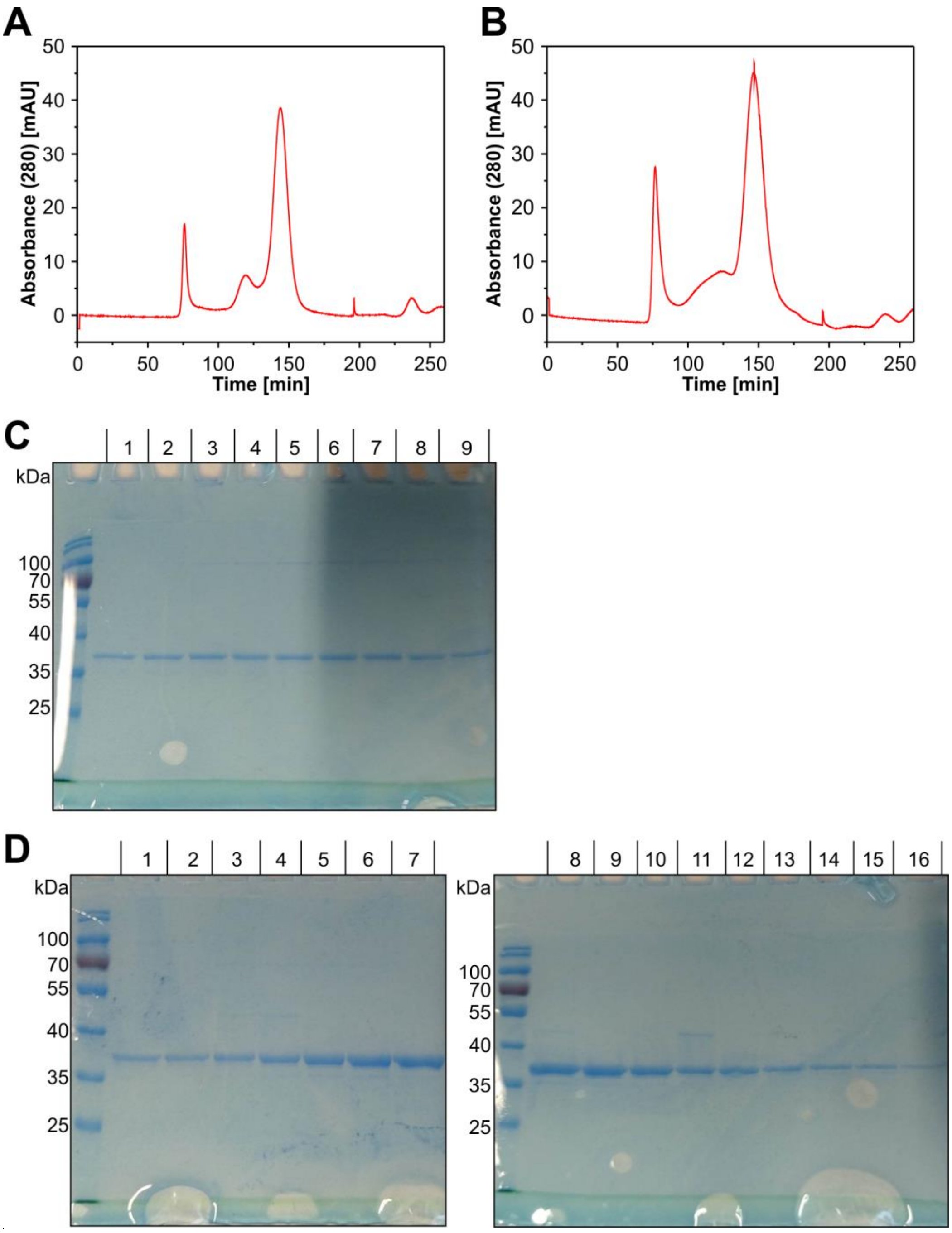

**Figure S 6:** Purification of recombinant human Nudt5 by SEC. **A:** Chromatogram of SEC using eluted protein from IMAC purification (first peak, see supplementary **Figure. S5**). **B:** Chromatogram of SEC using eluted protein from IMAC purification (second peak, see supplementary **Figure S5**). **C:** 10% SDS-PAGE analysis of individual eluted fractions of the main peak from SEC seen in A. **D:** 10% SDS-PAGE analysis of individual eluted fractions of the main peak from SEC seen in B. Gels were stained with READYBLUE PROTEIN GEL STAIN. The larger molecular mass indicated by SDS-PAGE analysis compared to the theoretical value of 26 kDa is in accordance with the literature (5,6).

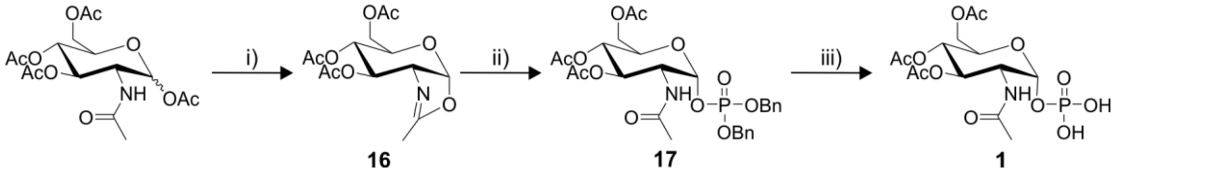

**Figure S 7:** Chemical synthesis of $Ac_3$GlcNAc 1-phosphate **1** building block. i) TMSOTf, DCE (anh.), 60°C, overnight, then $Et_3N$, 0°C, 96%. ii) Dibenzylphosphate, DCE (anh.), 5 d, 44%. iii) 10% Pd/C, $H_2$, MeOH, RT, overnight, 97%.

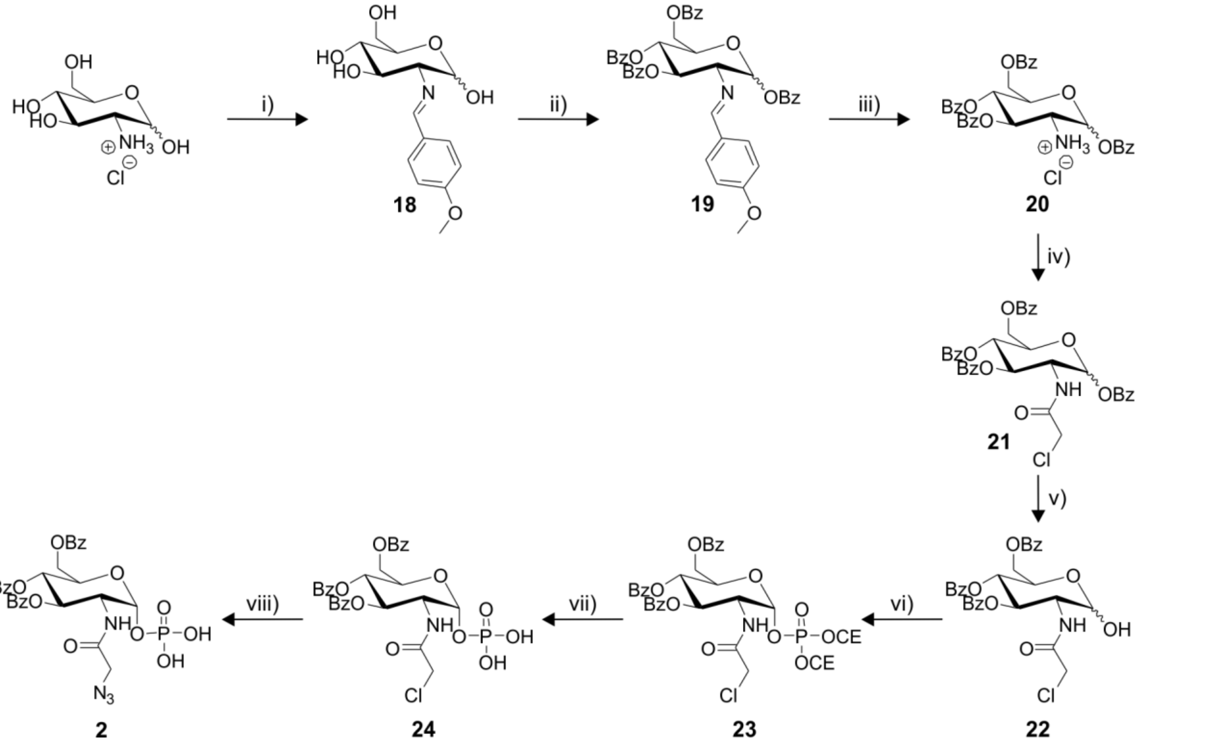

**Figure S 8:** Chemical synthesis of $Bz_3$GlcNAz 1-phosphate **2** building block. i) NaOH, 4-anisaldehyde, RT, 69%. ii) Benzoyl chloride, pyridine (anh.), RT, 3 h, 83%. iii) HCl, acetone, reflux, 53%. iv) Chloroacetic anhydride, $Et_3N$, DCM, RT, 88%. v) HBr in AcOH (30%), DCM, RT, 3 h, then $Ag_2CO_3$, acetone, $H_2O$, RT, 2 h, 43%. vi) Bis(2-cyanoethyl)-N,N-diisopropyl phosphoramidite, tetrazole, ACN (anh.), 40°C, then $H_2O_2$, THF, -78°C, 1.5 h, 60%. vii) DBU, BSA, ACN (anh.), RT, 1 h, 39%. viii) $NaN_3$, MeOH, 70°C, overnight, 71%.

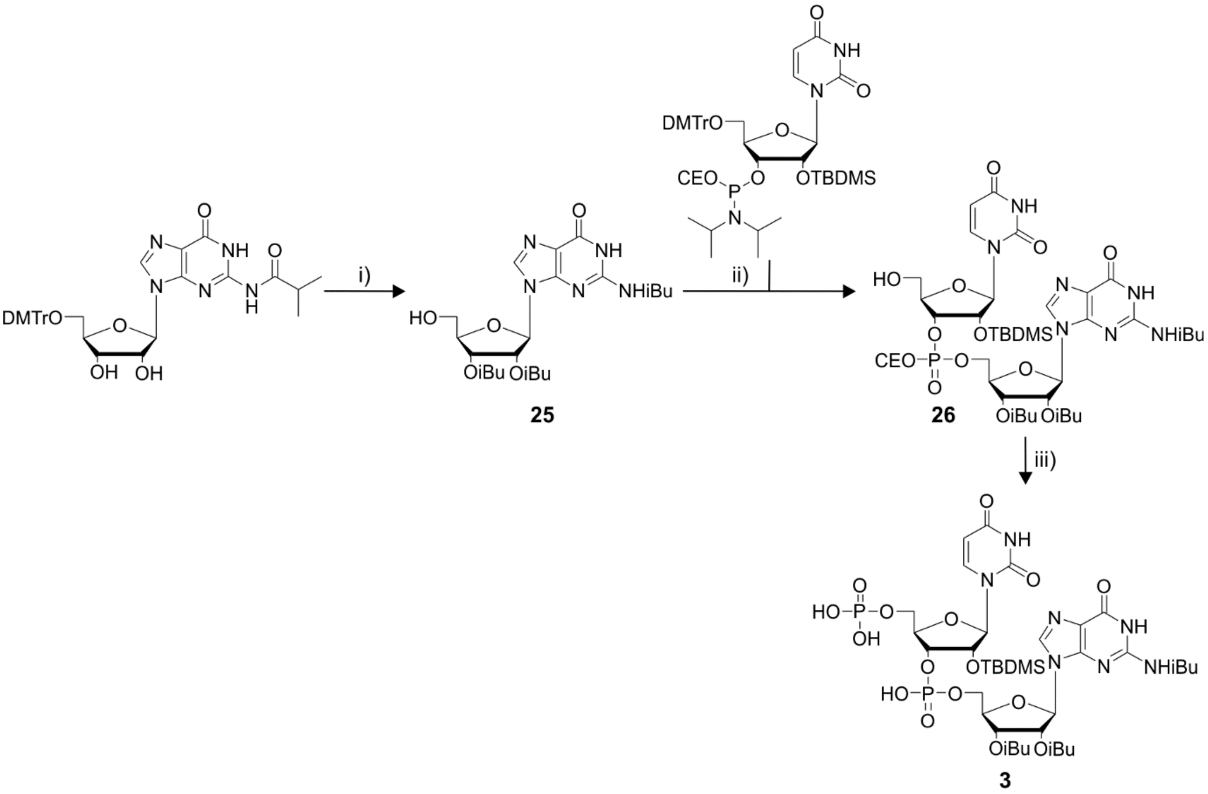

**Figure S 9:** Chemical synthesis of pUpG building block **3**. i) 1. Isobutyric anhydride, pyridine (anh.), RT, overnight; 2. TFA, MeOH, DCM, RT, 91%. ii) 1. DMT-2′O-TBDMS-rU-phosphoramidite, tetrazole, ACN (anh.), RT, then $I_2$, pyridine, THF, RT; 2. TFA, MeOH, DCM, RT, 78%. iii) 1. bis(2-cyanoethyl)-*N*,*N*-diisopropyl phosphoramidite, tetrazole, ACN (anh.), RT, then $I_2$, pyridine, THF, RT; 2. DBU, BSA, ACN (anh.), RT, 1 h, 65%.

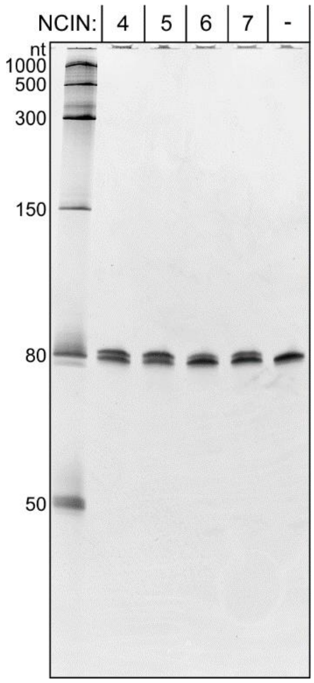

**Figure S 10:** Incorporation of the NCINs GlcNAcppUpG **4**, GlcNAzppUpG **5**, EP-GlcNAcppUpG **6** or MurNAcppUpG **7** into RNA by the T7 RNAP using a G initiating template coding for a 79mer RNA and a 10-fold excess of NCIN over GTP. 10% PAGE analysis of purified products being a mixture of capped 80mer RNA and ppp-79mer RNA. The gel was stained with SYBR Gold.

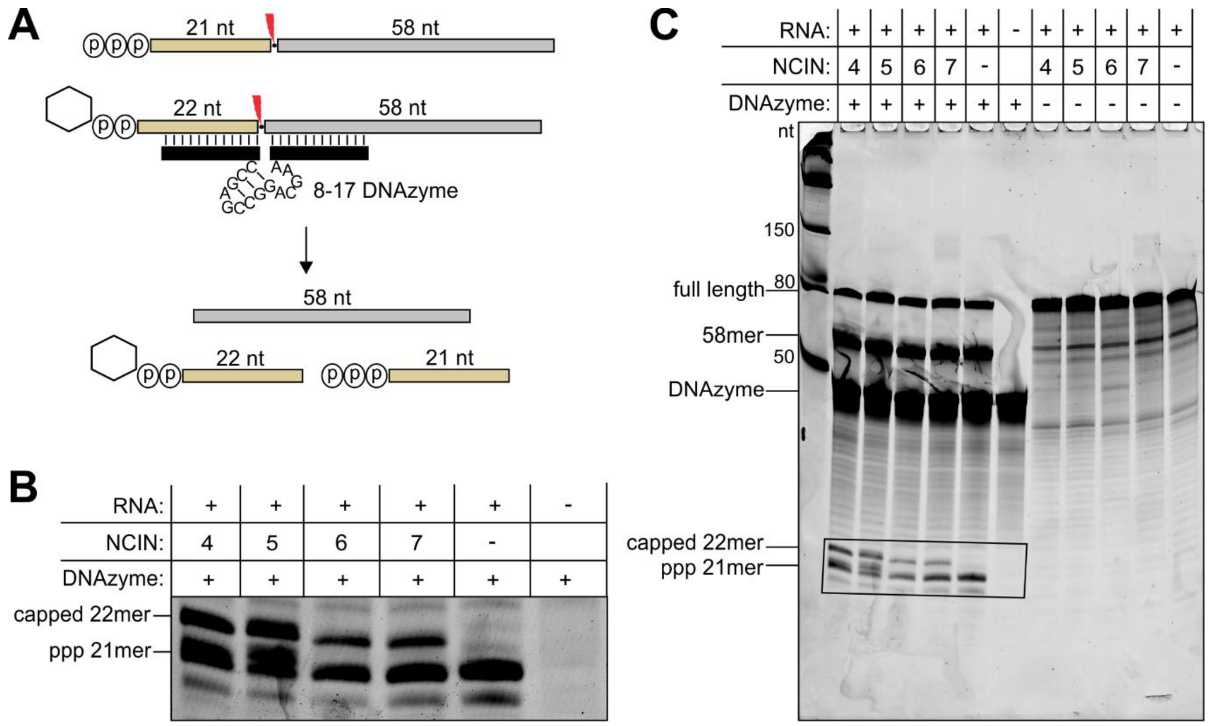

**Figure S 11:** Cleavage of UDP-sugar-capped 80mer / triphosphate 79mer mixtures by a sequence-specific 8-17 DNAzyme. The capped / uncapped RNA mixture was generated in T7 RNAP-catalyzed IVT reactions using a 10-fold excess of the NCIN GlcNAcppUpG **4**, GlcNAzppUpG **5**, EP-GlcNAcppUpG **6** or MurNAcppUpG **7** compared to GTP. **A:** Depiction of the DNAzyme mediated cleavage of 80mer / 79mer RNA to capped 22mer / triphosphate-21mer RNA and 5'-OH-58mer RNA. **B:** 20% PAGE analysis of reaction mixtures. 22mer and 21mer RNA can be separated (marked box). **C:** RNA from IVT reactions in which an NCIN was present show an additional small cleavage product corresponding to capped-22mer RNA. This band is absent if no NCIN was present in the IVT reaction. Densitometric quantification of bands corresponding to capped 22mer RNA and triphosphate-21mer RNA revealed capping efficiencies of 43% for GlcNAcppUpG **4** and GlcNAzppUpG **5**, 30% for EP-GlcNAcppUpG **6** and 26% for MurNAcppUpG **7**.

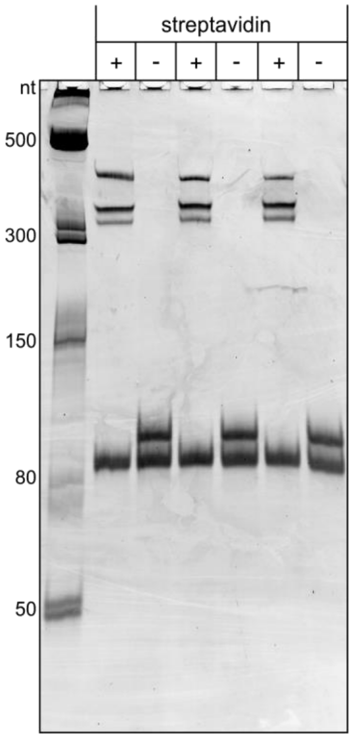

**Figure S 12:** 10% native PAGE analysis of streptavidin shift using GlcNAz-80mer / triphosphate-79mer RNA mixture after conjugation to DBCO-PEG$_4$-biotin in triplicates. The RNA 80mer / 79mer mixture was generated in a T7 RNAP-catalyzed IVT reaction with a 10-fold excess of GlcNAzppUpG **5** as NCIN compared to GTP. Due to streptavidin binding, 48±2% of RNA was retarded. Multiple shifted bands were observed as streptavidin can bind multiple biotin moieties.

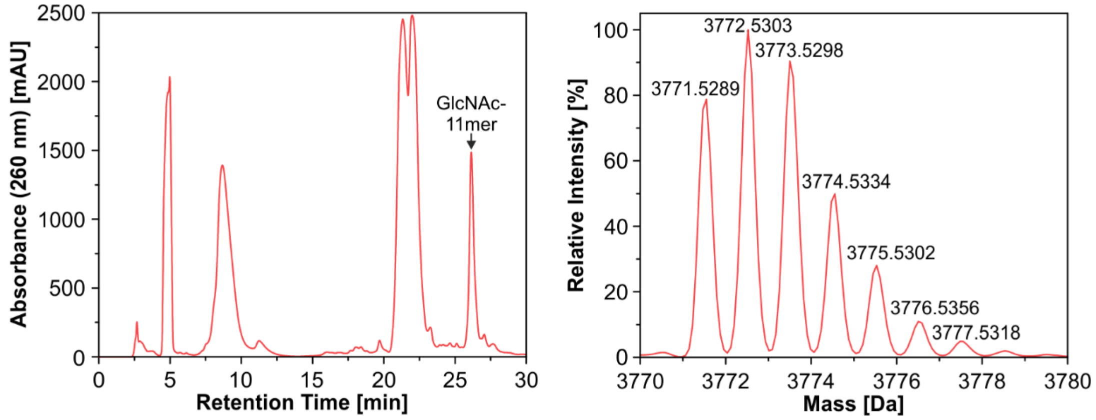

**Figure S 13:** Purification of GlcNAc-11mer RNA after using GlcNAcppUpG **4** in a GTP-free, T7 RNAP catalyzed *in vitro* transcription. Left: HPLC chromatogram of GlcNAc-11mer RNA purification showing efficient separation of capped 11mer RNA products from excess NTPs, dinucleotide, and abortive transcription products. Right: ESI-MS analysis of the purified product (Deconvoluted, $[M]_{calc}$: 3771.5026).

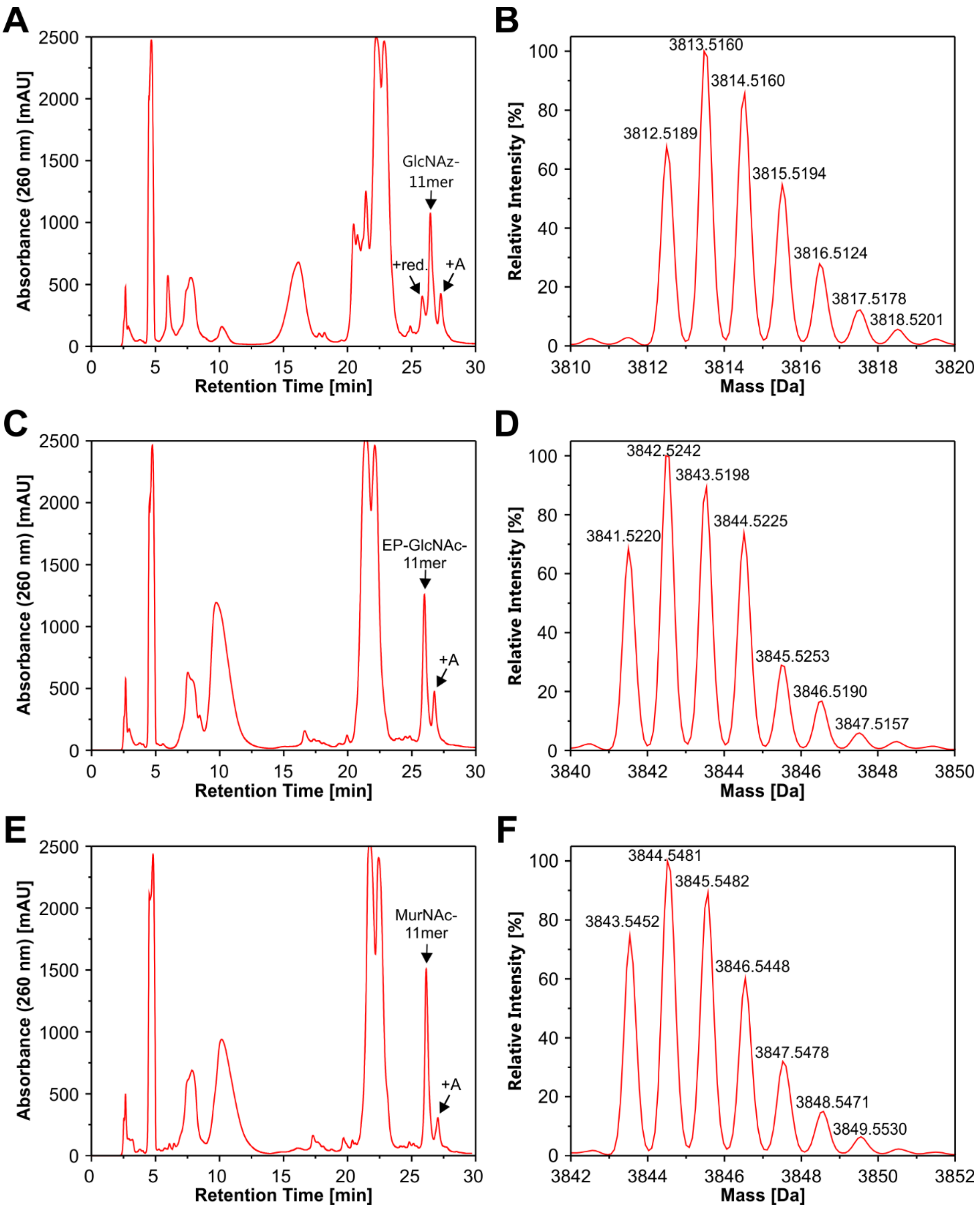

**Figure S 14:** Purification of sugar-11mer RNA after using capped dinucleotides as NCIN in T7 RNAP catalyzed *in vitro* transcription. "+A" depicts a product elongated by one adenosine. "red." depicts a GlcNAz-11mer byproduct in which the azide was reduced to the corresponding amine, probably caused by the presence of DTT during the IVT reaction. **A:** GlcNAzppUpG **5** was used as NCIN. RP-HPLC chromatogram of GlcNAz-11mer RNA purification. **B:** ESI-MS analysis of GlcNAz-11mer RNA (Deconvoluted, $[M]_{calc}$: 3812.5036). **C:** EP-GlcNAcppUpG **6** was used as NCIN. RP-HPLC chromatogram of EP-GlcNAcP-11mer RNA purification. **D:** ESI-MS analysis of EP-GlcNAc-11mer RNA (Deconvoluted, $[M]_{calc}$: 3841.5076). **E:** MurNAcppUpG **7** was used as NCIN. RP-HPLC chromatogram of MurNAc-11mer RNA purification. **F:** ESI-MS analysis of MurNAc-11mer RNA (Deconvoluted, $[M]_{calc}$: 3843.5233).

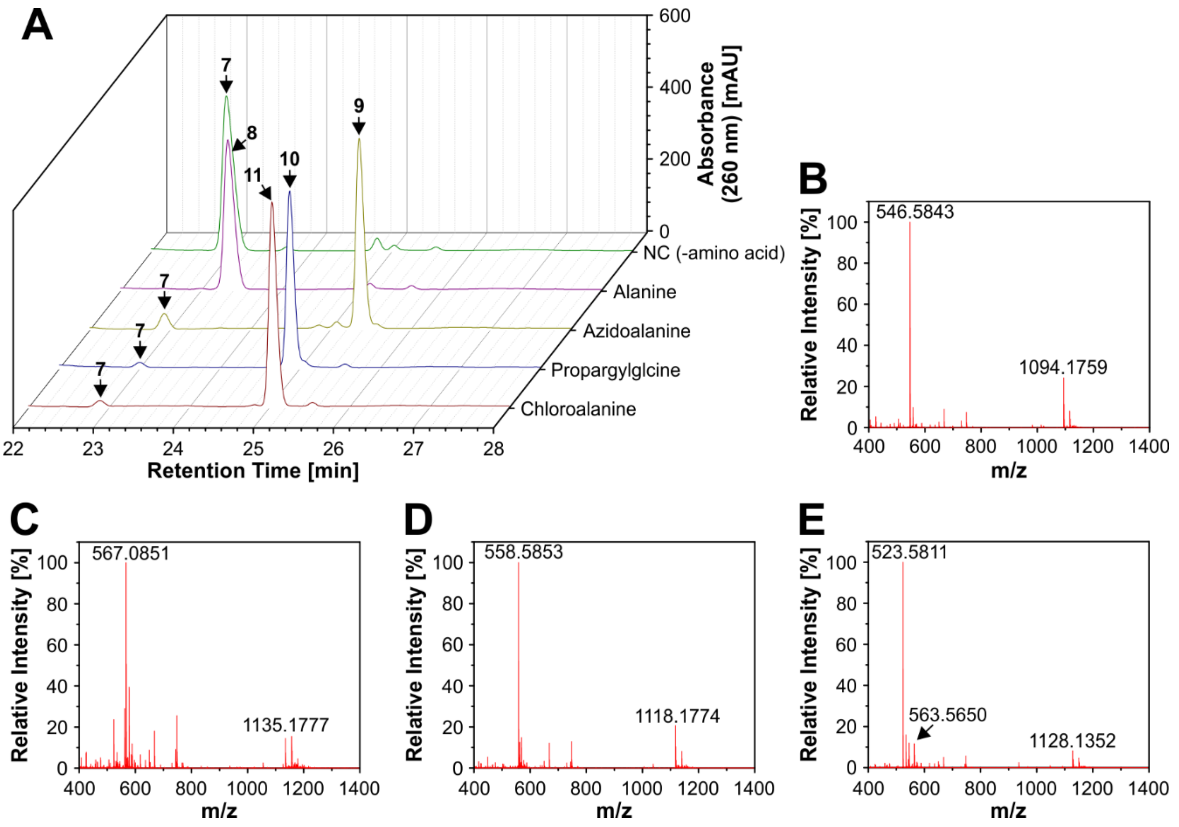

**Figure S 15:** Conversion of MurNAcppUpG **7** by MurC using different alanine derivatives and leading to Ala-MurNAcppUpG **8**, Azidoala-MurNAcppUpG **9**, Propargylgly-MurNAcppUpG **10** or Chloroala-MurNAcppUpG **11**. In the negative control, no amino acid was added to the reaction. **A:** RP-HPLC chromatograms of all reactions showing substrate and product peaks. **B:** ESI-MS analysis of RP-HPLC-purified Ala-MurNAcppUpG **8** ($[M-H]^{-}_{calc}$: 1094.1800; $[M-2H]^{2-}_{calc}$: 546.5864). **C:** ESI-MS analysis of RP-HPLC-purified Azidoala-MurNAcppUpG **9** ($[M-H]^{-}_{calc}$: 1135.1814; $[M-2H]^{2-}_{calc}$: 567.0871). **D:** ESI-MS analysis of RP-HPLC-purified Propargylgly-MurNAcppUpG **10** ($[M-H]^{-}_{calc}$: 1118.1800; $[M-2H]^{2-}_{calc}$: 558.5864). **E:** ESI-MS analysis of RP-HPLC-purified Chloroala-MurNAcppUpG **11** ($[M-H]^{-}_{calc}$: 1128.1410; $[M-2H]^{2-}_{calc}$: 563.5669), main mass peak corresponds to a species which probably underwent decarboxylation and β-elimination of chloride ($[M-2H]^{2-}_{calc}$: 523.5836) which might occur during the ionization process.

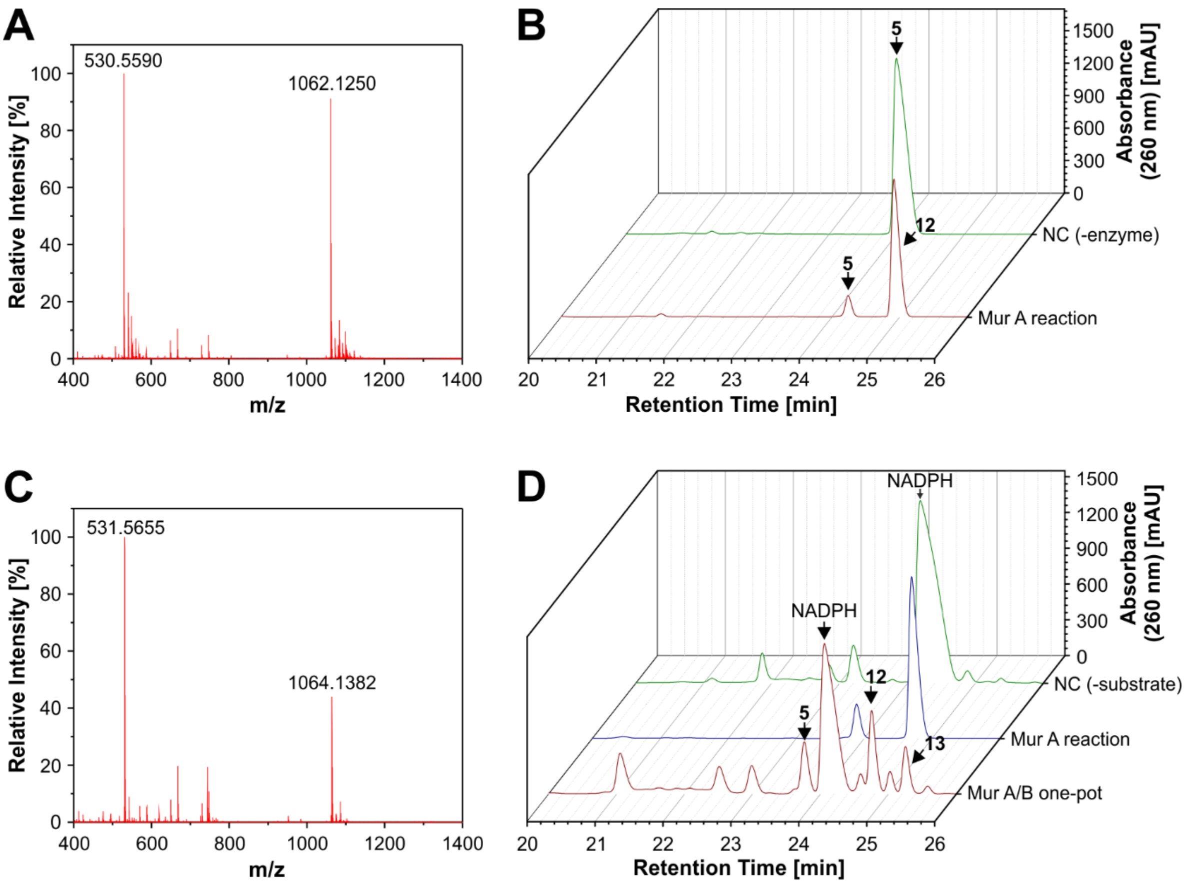

**Figure S 16:** Conversion of GlcNAzppUpG **5** by MurA or MurA and B in a one-pot reaction. **A:** ESI-MS analysis of the purified product after conversion of GlcNAzppUpG **5** by MurA leading to EP-GlcNAzppUpG **12** ($[M\text{-}H]^{-}_{calc}$: 1062.1286; $[M\text{-}2H]^{2-}_{calc}$: 530.5607). **B:** RP-HPLC purification of the reaction mixture after incubating GlcNAzppUpG **5** with MurA and PEP for 24 h. **C:** ESI-MS analysis of the product after conversion of GlcNAzppUpG **5** by MurA and MurB to MurNAzppUpG **13** ($[M\text{-}H]^{-}_{calc}$: 1064.1443; $[M\text{-}2H]^{2-}_{calc}$: 531.5685). **D:** RP-HPLC purification of the reaction mixture after incubating GlcNAzppUpG **5** with MurA and PEP for 20 h followed by the addition of MurB and NADPH and incubation for another 7 h. The identity of all peaks was confirmed by ESI-MS.

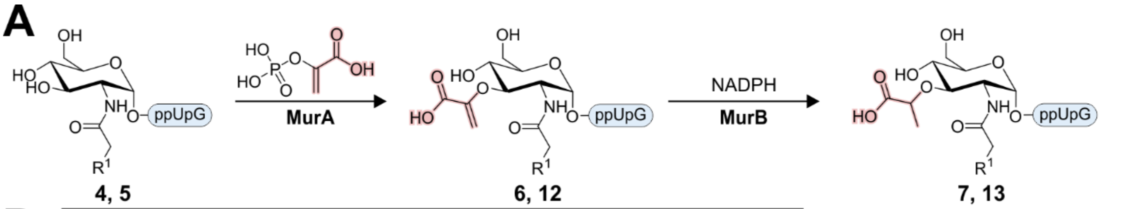

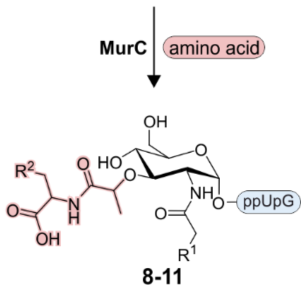

**B**

| No. | Name | R[1] | R[2] | Yield / Conversion |
|---|---|---|---|---|
| **4** | GlcNAcppUpG | H | - | 48%[a] |
| **5** | GlcNAzppUpG | $N_3$ | - | 18%[a] |
| **6** | EP-GlcNAcppUpG | H | - | 65%[b] |
| **7** | MurNAcppUpG | H | - | 52%[b] |
| **8** | Ala-MurNAcppUpG | H | H | quant.[c] |
| **9** | Azidoala-MurNAcppUpG | H | $N_3$ | 91±0.6%[c] |
| **10** | Propargylgly-MurNAcppUpG | H | $C_2H$ | 97±0.2%[c] |
| **11** | Chloroala-MurNAcppUpG | H | Cl | 96±0.3%[c] |
| **12** | EP-GlcNAzppUpG | $N_3$ | - | 89±2.3%[c] |
| **13** | MurNAzppUpG | $N_3$ | - | 27±3.4%[c, d] |

**Figure S 17: A:** Enzymatic synthesis of cell wall precursor-capped dinucleotides using MurA, MurB or MurC. **B:** Summary of all capped dinucleotides chemically or enzymatically synthesized in this study and corresponding yield or conversion. [a] Overall yield of isolated compounds from chemical synthesis starting from building blocks **1** or **2** and **3**. [b] Yield for isolated compounds after enzymatic conversions. [c] Conversion calculated based on RP-HPLC analysis, mean value of three separate reactions. [d] Conversion over two enzyme-catalyzed steps (MurA and B one-pot).

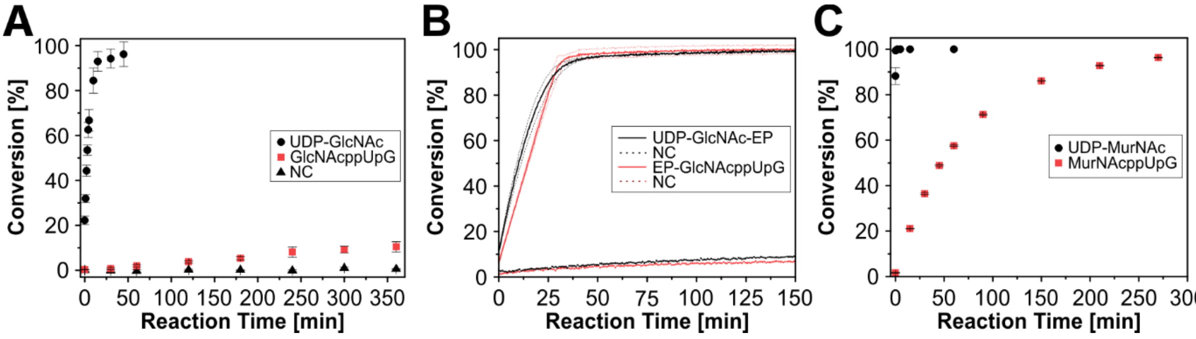

**Figure S 18: A:** Comparison of the conversion of UDP-GlcNAc with the conversion of GlcNAcppUpG **4** by MurA (n = 3, data points show mean ± S.D.). **B:** Comparison of the conversion of UDP-GlcNAc-EP **14** with the conversion of EP-GlcNAcppUpG **6** by MurB (n = 3, curves show mean ± S.D.). **C:** Conversion of UDP-MurNAc **15** and MurNAcppUpG **7** by MurC in the presence of L-alanine (n = 3, data points show mean ± S.D.).

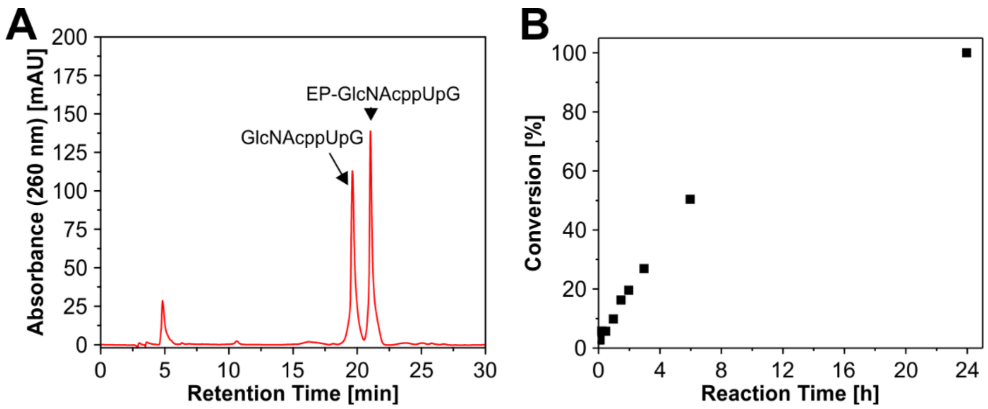

**Figure S 19:** Analysis of the conversion of GlcNAcppUpG **4** by MurA to EP-GlcNAcppUpG **6** on a preparative scale by RP-HPLC. **A:** RP-HPLC chromatogram after a reaction time of 6 h. The identity of peaks was confirmed by ESI-MS. **B:** Conversion of GlcNAcppUpG **4** as a function of time. The reaction was completed after 24 h.

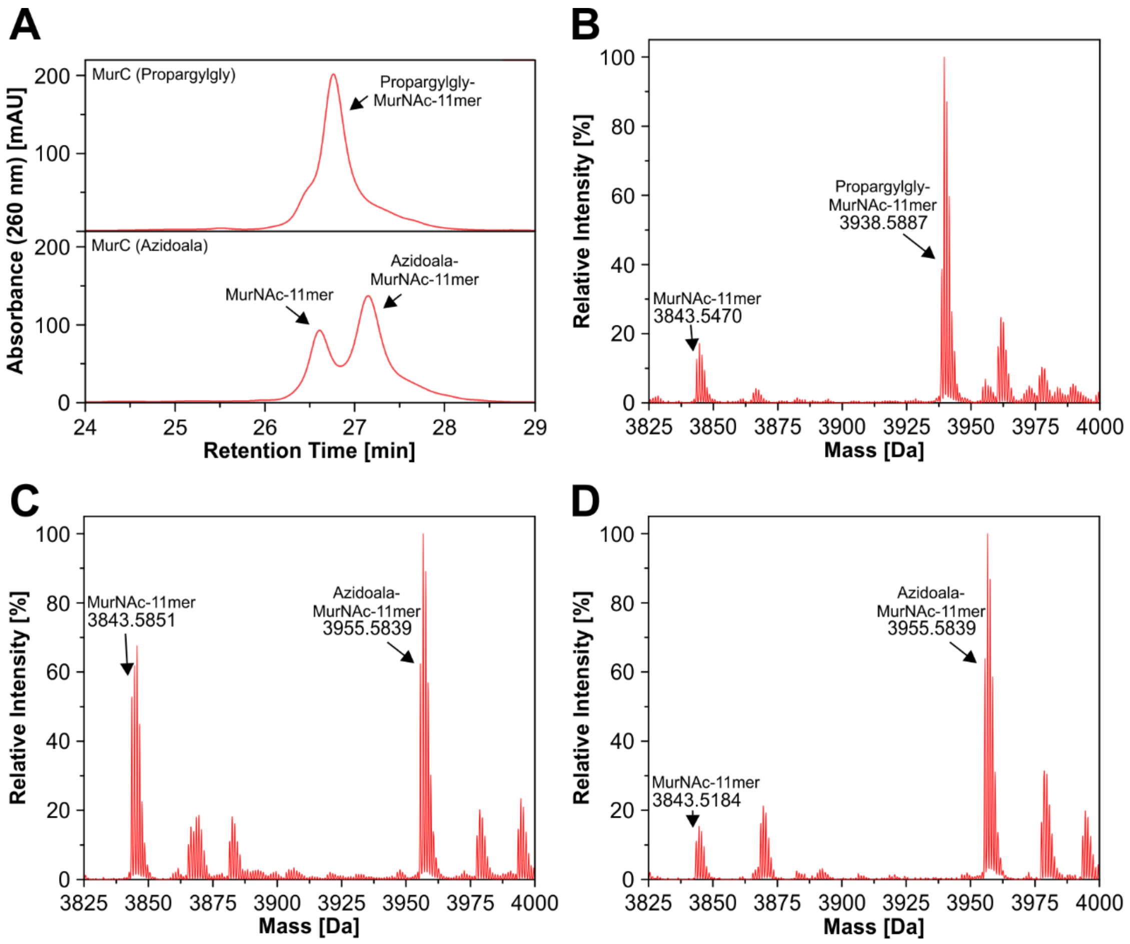

**Figure S 20: A:** RP-HPLC purification of MurNAc-capped 11mer RNA ligated to Propargylgly (top) or Azidoala (bottom) by MurC. The product and starting material could be partially separated by RP-HPLC after ligation. **B:** ESI-MS analysis of purified 11mer-RNA after the MurC reaction in the presence of Propargylgly. Ligation to Propargylgly led to a nearly complete conversion of MurNAc-11mer RNA to Propargylgly-MurNAc-11mer RNA (Deconvoluted, $[M]_{calc}$: 3938.5600). **C:** ESI-MS analysis of purified 11mer-RNA (first peak) after the MurC reaction in the presence of Azidoala. This fraction contained MurNAc-11mer RNA (Deconvoluted, $[M]_{calc}$: 3843.5233) and Azidoala-MurNAc-11mer RNA (Deconvoluted, $[M]_{calc}$: 3955.5620). **D:** ESI-MS analysis of purified 11mer-RNA (second peak) after the MurC reaction in the presence of Azidoala. This fraction contained mainly Azidoala-MurNAc-11mer RNA (Deconvoluted, $[M]_{calc}$: 3955.5620).

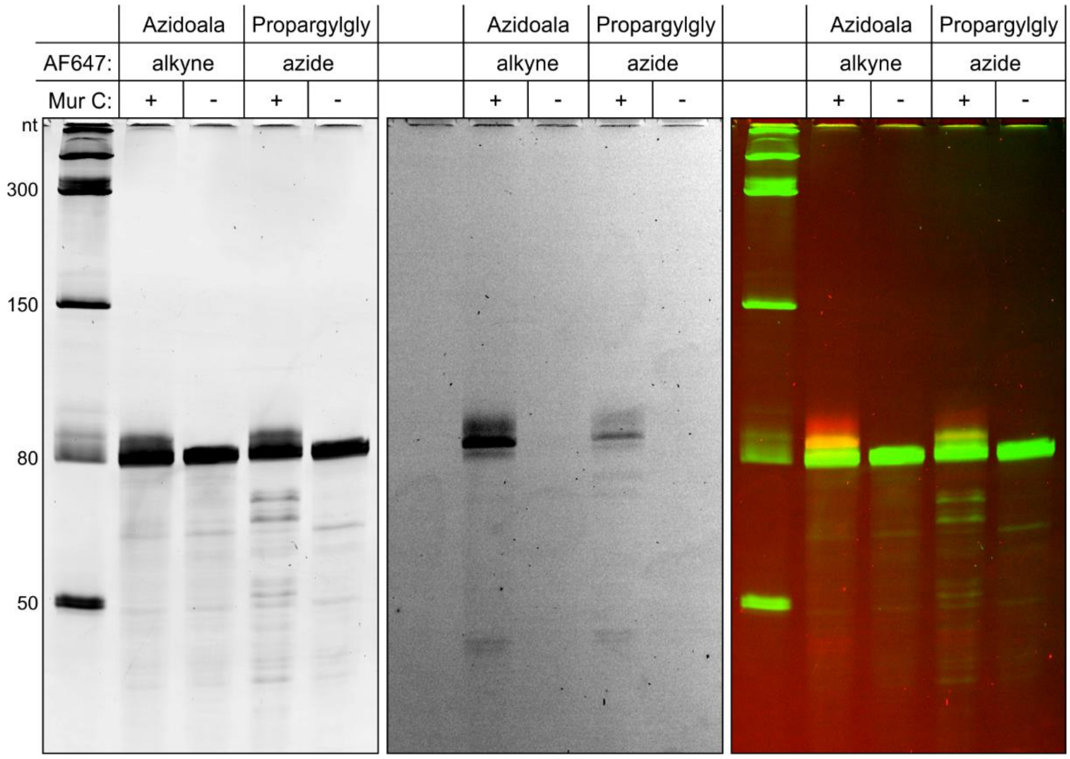

**Figure S 21:** 10% PAGE analysis of MurNAc-80mer / triphosphate-79mer RNA mixture after ligation to Azidoala or Propargylgly by MurC and conjugation to AF647-alkyne or AF647-azide by CuAAC. Left: SYBR Gold stain; center: AF647; right: overlay. In negative controls, RNA was incubated with alanine derivatives in the absence of MurC. MurNAc-capped RNA could be labeled with both amino acid derivatives.

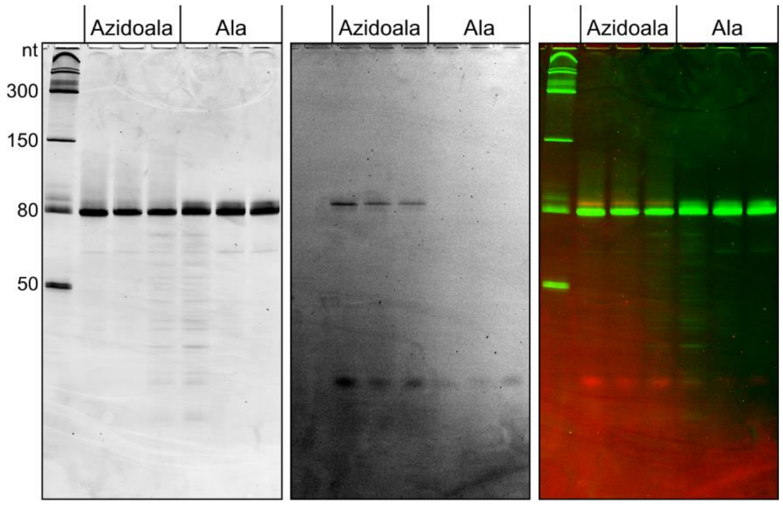

**Figure S 22:** 10% PAGE analysis of MurNAc-80mer / triphosphate-79mer RNA mixture after ligation to Azidoala by MurC and conjugation to DBCO-AF647 using SPAAC. Left: Sybr Gold stain; center: AF647; right: overlay. Ligation was conducted in triplicates, negative controls contained MurC during the ligation step, but Ala instead of Azidoala.

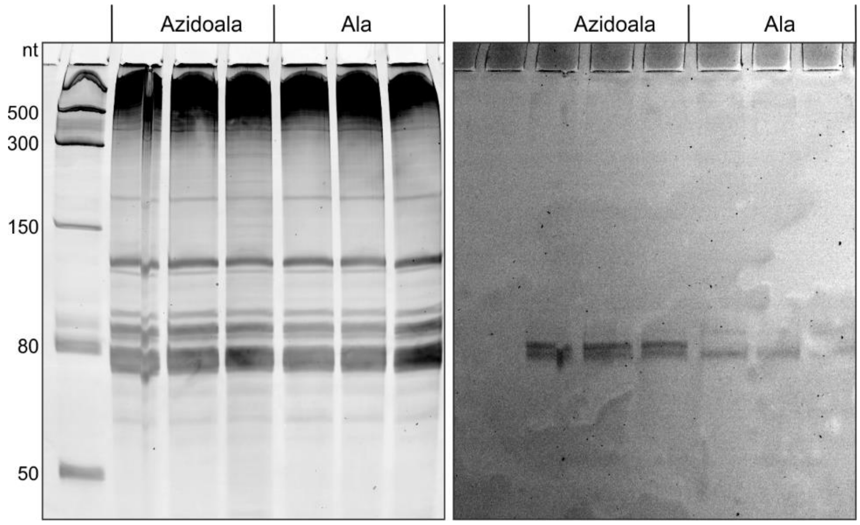

**Figure S 23:** 10% PAGE analysis of MurNAc-80mer / triphosphate-79mer RNA mixture spiked into *E. coli* total RNA (30 ng per 1 µg total RNA) prior to ligation to Azidoala by MurC and conjugation to DBCO-AF647 using SPAAC. Left: Sybr Gold stain; right: AF647. Ligation was conducted in triplicates, negative controls contained MurC during the ligation step, but Ala instead of Azidoala.

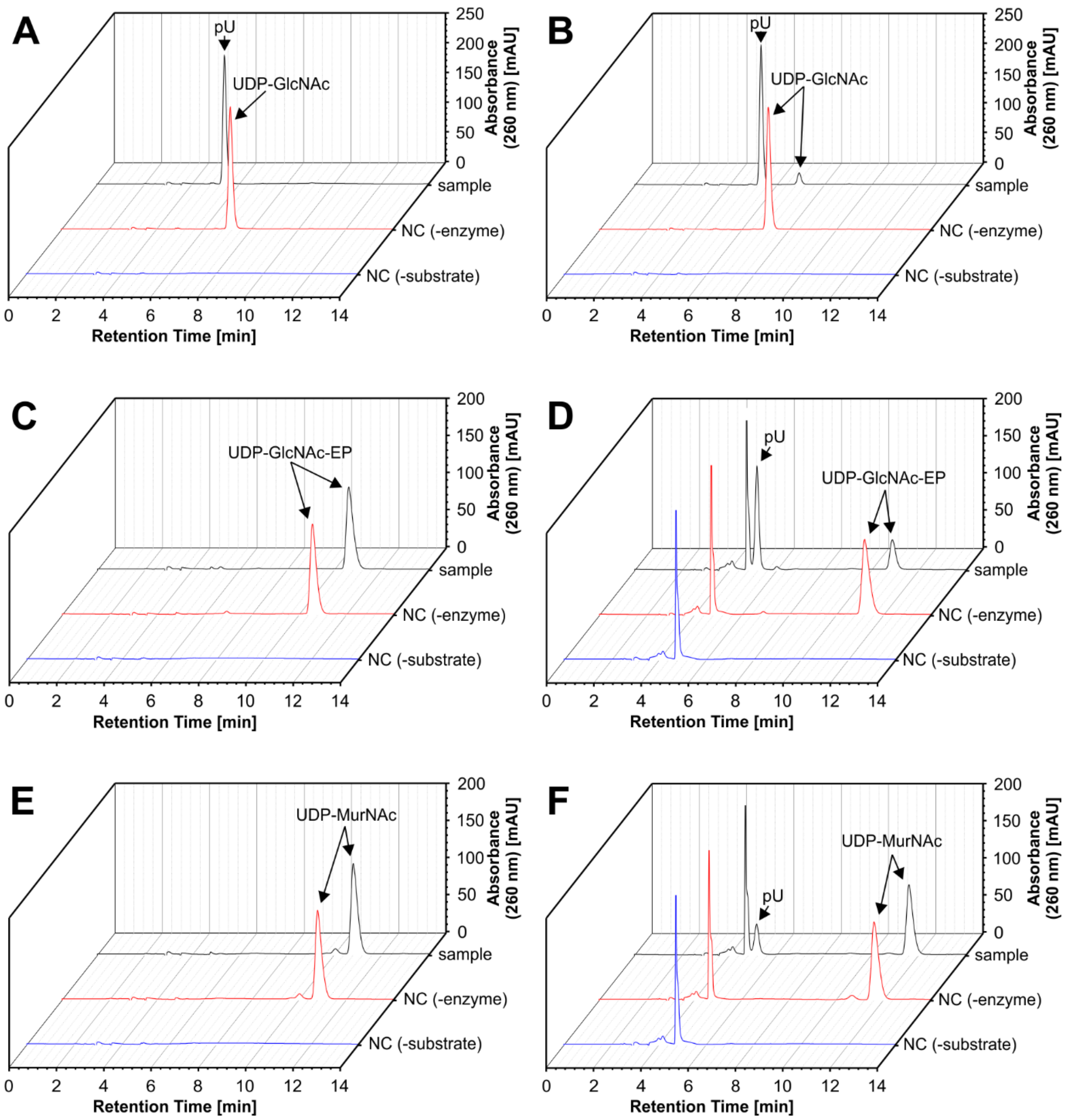

**Figure S 24:** RP-HPLC analysis after treatment of mononucleotide-sugars with hNudt5 or NudF for 4 h. Identity of peaks was confirmed by ESI-MS analysis. Negative control (NC) reactions were set up in the absence of either enzyme or mononucleotide-sugar substrate. **A:** Treatment of UDP-GlcNAc with hNudt5. The substrate was completely cleaved to uridine monophosphate (pU). **B:** Treatment of UDP-GlcNAc with NudF. Cleavage to pU was nearly completed. **C:** Treatment of UDP-GlcNAc-EP **14** with hNudt5. No cleavage product could be detected. **D:** Treatment of UDP-GlcNAc-EP **14** with NudF. Most of the substrate was converted. **E:** Treatment of UDP-MurNAc **15** with hNudt5. No cleavage product could be detected. **F:** Treatment of UDP-MurNAc **15** with NudF. The substrate was partially converted.

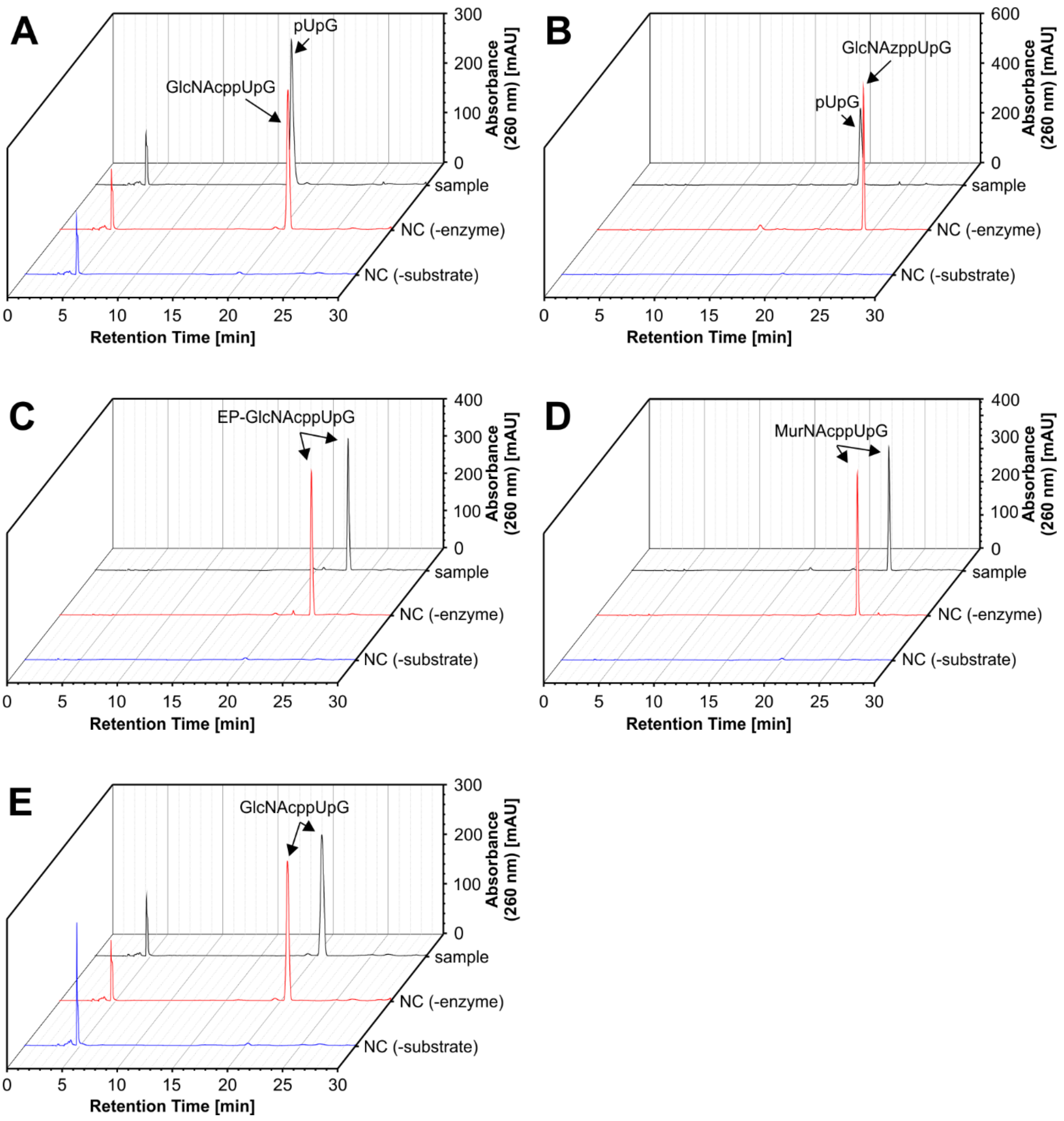

**Figure S 25:** RP-HPLC analysis after treatment of sugar-dinucleotide with hNudt5 or NudF for 4 h. Identity of peaks was confirmed by ESI-MS analysis. Negative control (NC) reactions were set up in the absence of either enzyme or sugar-dinucleotide substrate. **A:** Treatment of GlcNAcppUpG **4** with hNudt5. The substrate was completely cleaved to monophosphate uridine-guanosine dinucleotide (pUpG). **B:** Treatment of GlcNAzppUpG **5** with hNudt5. The substrate was completely cleaved to pUpG. **C:** Treatment of EP-GlcNAcppUpG **6** with hNudt5. No cleavage product could be detected. **D:** Treatment of MurNAcppUpG **7** with hNudt5. No cleavage product could be detected. **E:** Treatment of GlcNAcppUpG **4** with NudF. No cleavage product could be detected.

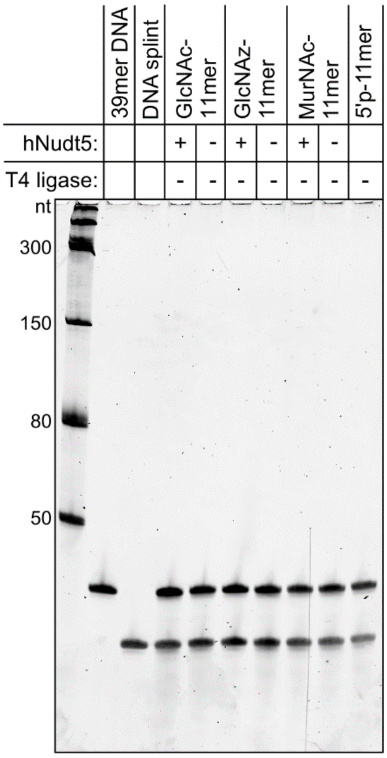

**Figure S 26:** 10% PAGE analysis of PABLO assay carried out with hNudt5-treated 11mer RNA carrying different sugar caps (SYBR Gold stain). Negative controls in which T4 DNA ligase was absent during the ligation step. In the absence of T4 DNA ligase, no chimeric 50mer ligation product due to the ligation of decapped 11mer RNA to 39mer DNA can be observed.

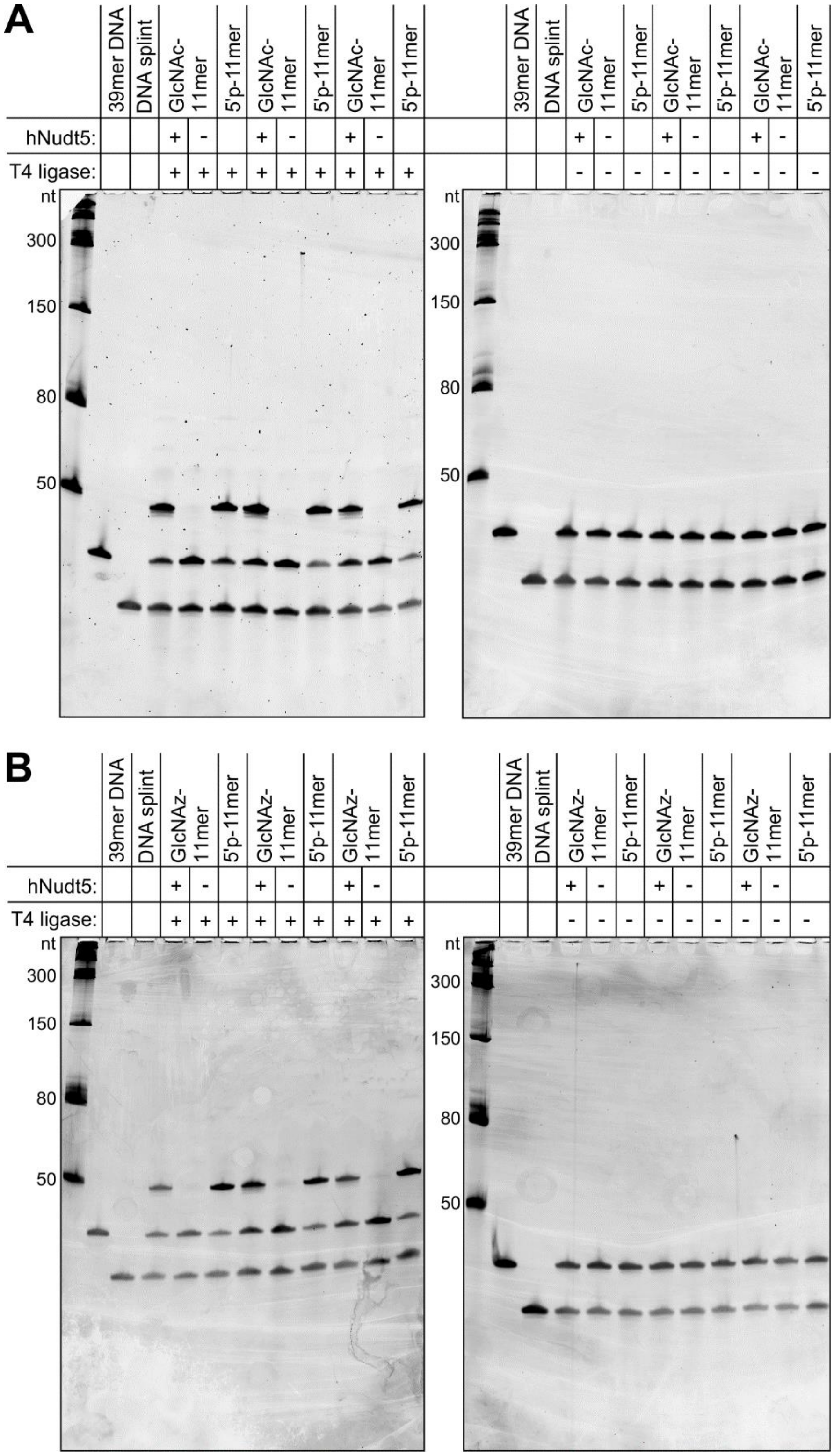

**Figure S 27:** 10% PAGE analysis of PABLO assay conducted with hNudt5-treated capped 11mer RNA (SYBR Gold stain). **A:** PABLO assay of hNudt5-treated GlcNAc-capped RNA in triplicates. **B:** PABLO assay of hNudt5-treated GlcNAz-capped RNA in triplicates. The 50 nucleotide-long chimeric ligation product formed due to the ligation of decapped 11mer RNA to 39mer DNA can only be observed if hNudt5 was present during the decapping step and T4 DNA ligase during the ligation step.

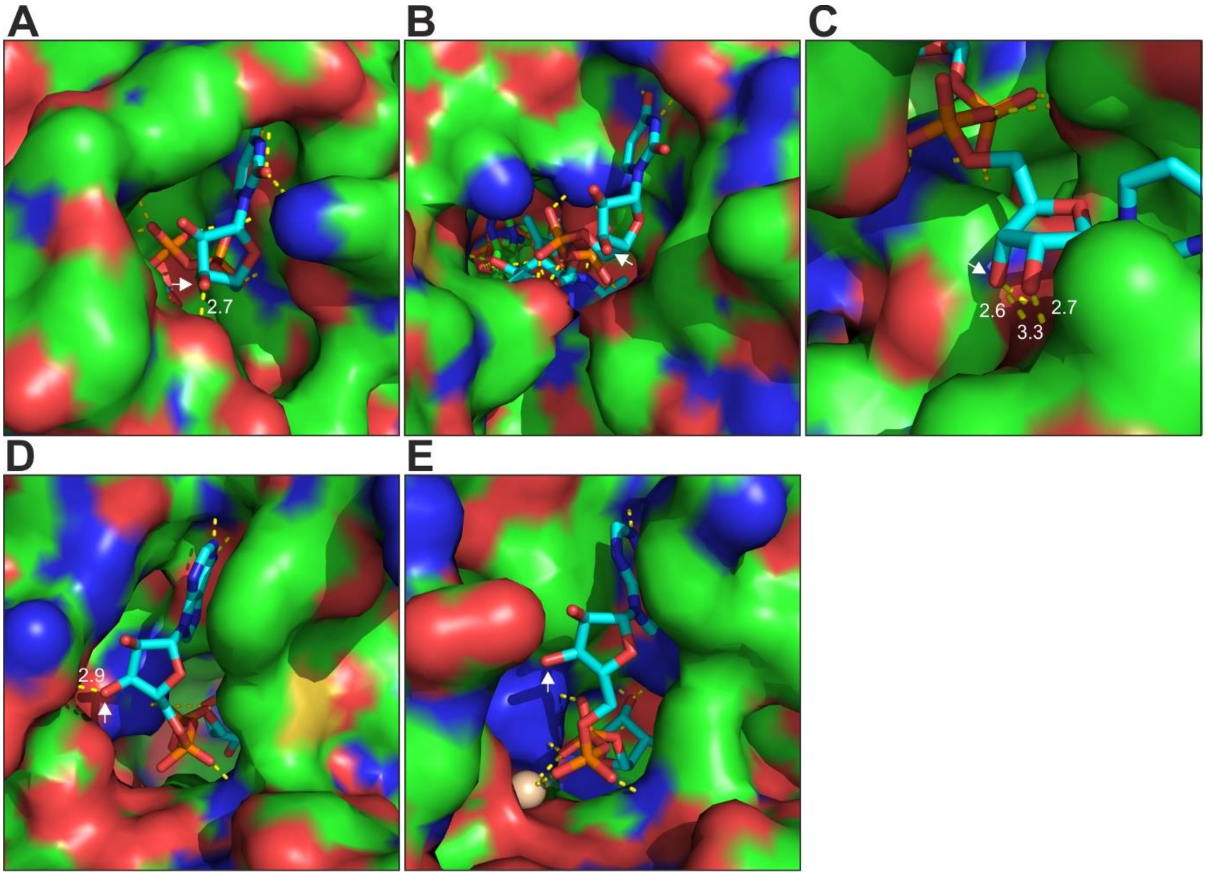

**Figure S 28:** Investigated enzymes in complex with their respective mononucleotide substrates. The white arrow points towards the 3’ OH group of the ribose which is involved in the phosphodiester bond in the dinucleotide analog. Yellow dashed lines indicate hydrogen bonds, and white numbers indicate the length of hydrogen bonds of the 2’-OH or 3’-OH group of ribose in Å. Structures were visualized using PyMOL Molecular Graphics System (Version 2.6.0a0, Schrödinger). **A:** *E. coli* MurA in complex with UDP-GlcNAc (PDB ID: 1UAE) (7). **B:** *E. coli* MurB in complex with UDP-GlcNAc-EP (PDB ID: 2MBR) (8). **C:** *H. influenzae* MurC in complex with UDP-MurNAc (PDB ID: 1P31) (9). **D:** *E. coli* NudF in complex with ADP-ribose (PDB ID: 1G9Q) (10). **E:** *H. sapiens* Nudt5 in complex with ADP-ribose (PDB ID: 2DSC) (11).

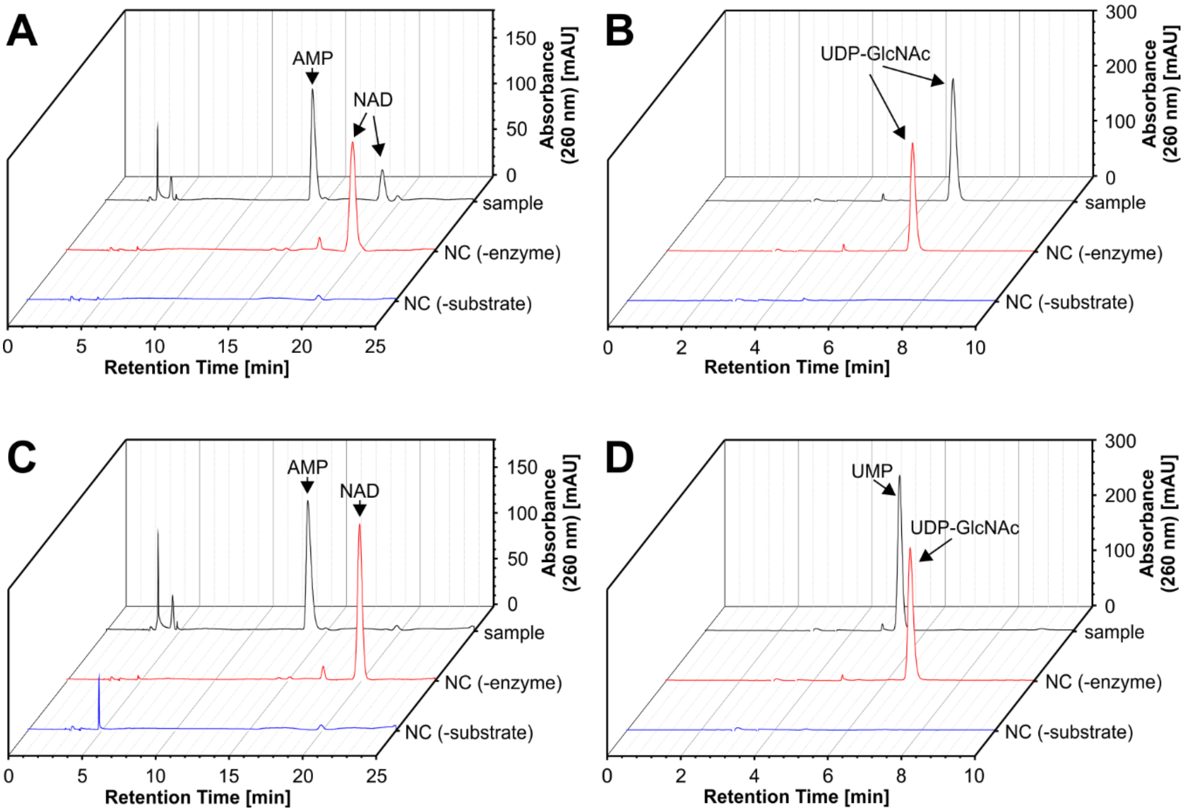

**Figure S 29:** RP-HPLC analysis after treatment of NAD or UDP-GlcNAc with NudC or hNudt5 for 4 h. Identity of peaks was confirmed by ESI-MS analysis. **A:** Treatment of NAD with hNudt5. ~78% cleavage to AMP was detected. **B:** Treatment of UDP-GlcNAc with NudC. No cleavage could be observed. **C:** Treatment of NAD with NudC. The substrate was completely cleaved to AMP. **D:** Treatment of UDP-GlcNAc with hNudt5. The substrate was completely cleaved to UMP.

**Table S 1:** Sequences of used DNA oligonucleotides. “mN” (N = A, C, G or U) depicts 2’-O-methyl RNA nucleotides.

| Description | Sequence |
|---|---|
| 11 mer FWD | TAATACGACTCACTATAGCACTCAATCGACGAGCCTGAGCC |
| 11mer REV | GGCTCAGGCTCGTCGATTGAGTGCTATAGTGAGTCGTATTA |
| 79mer FWD | TAATACGACTCACTATAGGATCCTCTGTAGTTCAGTCGGTAG AACGGCGGACTGTTAATC |
| 79mer REV | mUmGGCTCCTCTGACTGGACTCGAACCAGTGACATACGGAT TAACAGTCCGCCGTT |
| 68mer FWD | TAATACGACTCACTATAGGATCCTCTGTAGTTCAGTCGGTAG AACGGCGGACTGTT |
| 68mer REV | mAmCTGGACTCGAACCAGTGACATACGGATTAACAGTCCGC CGTTCTACCG |
| PABLO assay 39mer | GGAGTGCTACCACAACTTTAGCCATAATGTCACTTCTGC |
| Splint (PABLO assay) | GATTGAGTGCAGCAGAAGTGACATTATGGC |
| Splint (ligation capped 11mer) | CCGACTGAACTACAGAGGATCCGATTGAGTGCA |
| MurC primer FWD | CGACCCATGGCGAATACACAACAATTGGC |
| MurC primer Rev | TGGTGCTCGAGGCTGCCGCGCGGCACCAGGTCATGTTGTTC TTCCTCCGG |
| DNAzyme 5 | TAACAGTCCGCCGTTCTATCCGAGCCGGACGAAGACTGAAC TACAGAGGA |

**Table S 2:** Sequences of used RNA oligonucleotides.

| Description | Sequence |
|---|---|
| 11mer | UGCACUCAAUC |
| 68mer | GGAUCCUCUGUAGUUCAGUCGGUAGAACGGCGGACUGUUAA UCCGUAUGUCACUGGUUCGAGUCCAGU |
| 79mer | GGAUCCUCUGUAGUUCAGUCGGUAGAACGGCGGACUGUUAA UCCGUAUGUCACUGGUUCGAGUCCAGUCAGAGGAGCCA |

**Table S 3:** DNA sequences of genes in Pet-28C(+) vectors used for protein expression. Used restriction sites are underlined.

| Enzyme | DNA sequence |
|---|---|
| MurA | CCATGGATAAATTTCGTGTTCAGGGGCCAACGAAGCTCCAGGGCGAAGTC<br>ACAATTTCCGGCGCTAAAAATGCTGCTCTGCCTATCCTTTTTGCCGCACTA<br>CTGGCGGAAGAACCGGTAGAGATCCAGAACGTCCCGAAACTGAAAGACG<br>TCGATACATCAATGAAGCTGCTAAGCCAGCTGGGTGCGAAAGTAGAACGT<br>AATGGTTCTGTGCATATTGATGCCCGCGACGTTAATGTATTCTGCGCACCT<br>TACGATCTGGTTAAAACCATGCGTGCTTCTATCTGGGCGCTGGGGCCGCT<br>GGTAGCGCGCTTTGGTCAGGGGCAAGTTTCACTACCTGGCGGTTGTACG<br>ATCGGTGCGCGTCCGGTTGATCTACACATTTCTGGCCTCGAACAATTAGG<br>CGCGACCATCAAACTGGAAGAAGGTTACGTTAAAGCTTCCGTCGATGGTC<br>GTTTGAAAGGTGCACATATCGTGATGGATAAAGTCAGCGTTGGCGCAACG<br>GTGACCATCATGTGTGCTGCAACCCTGGCGGAAGGCACCACGATTATTGA<br>AAACGCAGCGCGTGAACCGGAAATCGTCGATACCGCGAACTTCCTGATTA<br>CGCTGGGTGCGAAAATTAGCGGTCAGGGCACCGATCGTATCGTCATCGA<br>AGGTGTGGAACGTTTAGGCGGCGGTGTCTATCGCGTTCTGCCGGATCGT<br>ATCGAAACCGGTACTTTCCTGGTGGCGGCGGCGATTTCTCGCGGCAAAAT<br>TATCTGCCGTAACGCGCAGCCAGATACTCTCGACGCCGTGCTGGCGAAA<br>CTGCGTGACGCTGGAGCGGACATCGAAGTCGGCGAAGACTGGATTAGCC<br>TGGATATGCATGGCAAACGTCCGAAGGCTGTTAACGTACGTACCGCGCC<br>GCATCCGGCATTCCCGACCGATATGCAGGCCCAGTTCACGCTGTTGAACC<br>TGGTGGCAGAAGGGACCGGGTTTATCACCGAAACGGTCTTTGAAAACCG<br>CTTTATGCATGTGCCAGAGCTGAGCCGTATGGGCGCGCACGCCGAAATC<br>GAAAGCAATACCGTTATTTGTCACGGTGTTGAAAAACTTTCTGGCGCACAG<br>GTTATGGCAACCGATCTGCGTGCATCAGCAAGCCTGGTGCTGGCTGGCT<br>GTATTGCGGAAGGGACGACGGTGGTTGATCGTATTTATCACATCGATCGT<br>GGCTACGAACGCATTGAAGACAAACTGCGCGCTTTAGGTGCAAATATTGA<br>GCGTGTGAAAGGCGAACTGGTGCCGCGCGGCAGCCTCGAG |
| MurB | CCATGGACCACTCCTTAAAACCCTGGAACACATTTGGCATTGATCATAATG<br>CTCAGCACATTGTATGTGCCGAAGACGAACAACAATTACTCAATGCCTGG<br>CAGTATGCAACCGCAGAAGGACAACCCGTTCTTATTCTGGGTGAAGGAAG<br>TAATGTACTTTTTCTGGAGGACTATCGCGGCACGGTGATCATCAACCGGA<br>TCAAAGGTATCGAAATTCATGATGAACCTGATGCGTGGTATTTACATGTAG<br>GAGCCGGAGAAAACTGGCATCGTCTGGTAAAATACACTTTGCAGGAAGGT<br>ATGCCTGGTCTGGAAAATCTGGCATTAATTCCTGGTTGTGTCGGCTCATCA<br>CCTATCCAGAATATTGGTGCTTATGGCGTAGAATTACAGCGAGTTTGCGCT<br>TATGTTGATTCTGTTGAACTGGCGACAGGCAAGCAAGTGCGCTTAACTGC<br>CAAAGAGTGCCGTTTTGGCTATCGCGACAGTATTTTTAAACATGAATACCA<br>GGACCGCTTCGCTATTGTAGCCGTAGGTCTGCGTCTGCCAAAAGAGTGG<br>CAACCTGTACTAACGTATGGTGACTTAACTCGTCTGGATCCTACAACAGTA<br>ACGCCACAGCAAGTATTTAATGCGGTGTGTCATATGCGCACCACCAAACT<br>CCCTGATCCAAAAGTGAATGGCAATGCCGGTAGTTTCTTCAAAAACCCTGT<br>TGTATCTGCCGAAACGGCTAAAGCATTACTGTCACAATTTCCAACAGCACC<br>AAATTACCCCCAGGCGGATGGTTCAGTAAAACTGGCAGCAGGTTGGCTTA<br>TCGATCAGTGCCAGCTAAAAGGGATGCAAATAGGTGGGGCTGCGGTGCA<br>CCGTCAACAGGCGTTAGTTCTCATTAATGAAGACAATGCAAAAAGCGAAG<br>ATGTTGTACAGCTGGCGCATCATGTAAGACAGAAAGTTGGTGAAAAATTTA<br>ATGTCTGGCTTGAGCCTGAAGTCCGCTTTATTGGTGCATCAGGTGAAGTG<br>AGCGCAGTGGAGACAATTTCACTGGTGCCGCGCGGCAGCCTCGAG |
| MurC | CCATGGCGAATACACAACAATTGGCAAAACTGCGTTCCATCGTGCCCGAA<br>ATGCGTCGCGTTCGGCACATACATTTTGTCGGCATTGGTGGTGCCGGTAT<br>GGGCGGTATTGCCGAAGTTCTGGCCAATGAAGGTTATCAGATCAGTGGTT |

| | |
|---|---|
| | CCGATTTAGCGCCAAATCCGGTCACGCAGCAGTTAATGAATCTGGGTGCG<br>ACGATTTATTTCAACCATCGCCCGGAAAACGTACGTGATGCCAGCGTGGT<br>CGTTGTTTCCAGCGCGATTTCTGCCGATAACCCGGAAATTGTCGCCGCTC<br>ATGAAGCGCGTATTCCGGTGATCCGTCGTGCCGAAATGCTGGCTGAGTTA<br>ATGCGTTTTCGTCATGGCATCGCCATTGCCGGAACGCACGGCAAAACGAC<br>AACCACCGCGATGGTTTCCAGCATCTACGCAGAAGCGGGGCTCGACCCA<br>ACCTTCGTTAACGGCGGGCTGGTAAAAGCGGCGGGGGTTCATGCGCGTT<br>TGGGGCATGGTCGGTACCTGATTGCCGAAGCAGATGAGAGTGATGCATC<br>GTTCCTGCATCTGCAACCGATGGTGGCGATTGTCACCAATATCGAAGCCG<br>ACCACATGGATACCTACCAGGGCGACTTTGAGAATTTAAAACAGACTTTTA<br>TTAATTTTCTGCACAACCTGCCGTTTTACGGTCGTGCGGTGATGTGTGTTG<br>ATGATCCGGTGATCCGCGAATTGTTACCGCGAGTGGGGCGTCAGACCAC<br>GACTTACGGCTTCAGCGAAGATGCCGACGTGCGTGTAGAAGATTATCAGC<br>AGATTGGCCCGCAGGGGCACTTTACGCTGCTGCGCCAGGACAAAGAGCC<br>GATGCGCGTCACCCTGAATGCGCCAGGTCGTCATAACGCGCTGAACGCC<br>GCAGCTGCGGTTGCGGTTGCTACGGAAGAGGGCATTGACGACGAGGCTA<br>TTTTGCGGGCGCTTGAAAGCTTCCAGGGGACTGGTCGCCGTTTTGATTTC<br>CTCGGTGAATTCCCGCTGGAGCCAGTGAATGGTAAAAGCGGTACGGCAA<br>TGCTGGTCGATGACTACGGCCACCACCCGACGGAAGTGGACGCCACCAT<br>TAAAGCGGCGCGCGCAGGCTGGCCGGATAAAAACCTGGTAATGCTGTTT<br>CAGCCGCACCGTTTTACCCGTACGCGCGACCTGTATGATGATTTCGCCAA<br>TGTGCTGACGCAGGTTGATACCCTGTTGATGCTGGAAGTGTATCCGGCTG<br>GCGAAGCGCCAATTCCGGGAGCGGACAGCCGTTCGCTGTGTCGCACAAT<br>TCGTGGACGTGGGAAAATTGATCCCATTCTGGTGCCGGATCCGGCGCGG<br>GTAGCCGAGATGCTGGCACCGGTATTAACCGGTAACGACCTGATTCTCGT<br>TCAGGGGGCTGGTAATATTGGAAAAATTGCCCGTTCTTTAGCTGAAATCAA<br>ACTGAAGCCGCAAACTCCGGAGGAAGAACAACATGACCTGGTGCCGCGC<br>GGCAGCCTCGAG |
| NudF | TCTAGAAATAATTTTGTTTAACTTTAAGAAGGAGATATACCATGCTTAAGCC<br>AGACAACCTGCCCGTTACATTTGGCAAAAACGATGTAGAAATTATTGCACG<br>AGAAACACTTTATCGCGGCTTTTTTTCATTAGATCTTTATAGATTTCGTCAT<br>CGTCTATTCAACGGGCAAATGAGTCATGAGGTGCGGCGGGAAATTTTTGA<br>GCGCGGTCACGCCGCAGTCTTGCTACCCTTTGACCCAGTGCGTGATGAA<br>GTTGTGCTGATTGAGCAGATTCGGATTGCCGCGTACGACACCAGCGAAAC<br>CCCTTGGCTACTGGAGATGGTTGCCGGGATGATTGAAGAGGGTGAAAGT<br>GTGGAAGATGTTGCCCGTCGCGAAGCGATTGAAGAGGCGGGACTGATAG<br>TCAAACGGACCAAACCGGTGTTAAGTTTCCTGGCAAGCCCGGGGGGCAC<br>CAGTGAGCGTTCGTCAATTATGGTGGGCGAAGTGGACGCCACGACCGCA<br>AGCGGTATTCACGGTCTGGCTGATGAAAACGAAGATATTCGCGTTCATGT<br>GGTAAGCCGGGAACAGGCATACCAGTGGGTAGAAGAGGGGAAAATCGAC<br>AACGCAGCGTCGGTCATCGCTTTGCAATGGCTGCAGCTGCATCATCAAGC<br>GTTAAAAAATGAGTGGGCACTGGTGCCGCGCGGCAGCCTCGAG |
| hNudt5 | CCATGGAGTCCCAGGAGCCGACCGAGTCGTCGCAGAACGGCAAACAATA<br>CATTATCAGTGAAGAACTGATTAGTGAAGGCAAATGGGTAAAACTGGAAA<br>AAACCACTTATATGGATCCTACGGGCAAAACGCGCACCTGGGAAAGCGTG<br>AAGCGCACCACGCGCAAAGAACAGACCGCGGATGGAGTGGCTGTAATCC<br>CAGTGCTGCAGCGTACGTTACATTACGAATGTATTGTGCTTGTCAAGCAGT<br>TCCGCCCGCCTATGGGAGGATATTGCATTGAATTCCCTGCAGGCTTAATC<br>GATGACGGAGAGACCCCAGAAGCGGCAGCCTTGCGTGAGCTGGAAGAA<br>GAAACAGGTTACAAGGGCGATATTGCTGAATGTTCACCGGCGGTGTGTAT<br>GGACCCAGGTTTGAGTAATTGCACCATTCATATTGTAACGGTAACCATCAA<br>CGGAGATGATGCCGAGAATGCGCGGCCGAAACCGAAACCGGGTGATGG<br>AGAATTCGTCGAAGTTATTTCGCTGCCAAAAAATGATTTGCTGCAGCGCCT |

| | |
|---|---|
| | GGACGCCCTGGTAGCGGAAGAACATTTAACGGTCGATGCGCGCGTATATT CCTATGCGCTGGCGCTCAAACATGCAAATGCAAAACCATTTGAAGTCCCT TTTCTGAAGTTTCTGGTGCCGCGCGGCAGCCTCGAG |

#### 2 General Information

Chemical synthesis under argon was performed using the Schlenk technique. For thin layer chromatography (TLC), Polygram Sil G/UV254 pre-coated polyester sheets (Macherey Nagel, Düren, Germany) were used and evaluated using a UV hand-lamp from Krüss Optronic (Hamburg, Germany) or stained using Hanessian's stain (235 ml $H_2O$, 15 ml sulfuric acid, 12 g $(NH_4)_6Mo_7O_{24}*4H_2O$, 0.5 g $Ce(NH_4)_4(SO_4)_4*2H_2O$). Flash column chromatography was conducted using silica gel (60 Å, 40–63 µm; Sigma Aldrich, Steinheim, Germany). Deionized water was filtered using a MilliQ purification system (Merck Millipore, Burlington, MA, USA).

Analytical HPLC was conducted using the 1200 series from Agilent Technologies (Santa Clara, CA, USA) equipped with a VDSpher PUR 100 C18-SE 250 x 4.6 mm, 5 µm column (VDS optilab, Berlin, Germany). For semi-preparative HPLC, an 1100 Agilent technologies system equipped with a Luna 5µm C18(2) 100 Å 250 x 21.2 mm column (Phenomenex, Torrance, CA, USA) was used. Preparative reverse phase (RP) chromatography was conducted using a PuriFlash PF420 system (Interchim, Montluçon, France).

Nuclear magnetic resonance (NMR) spectra were recorded on a Mercury plus 300 MHz or Mercury plus 500 MHz spectrometer from Varian (Crawley, UK). Chemical shifts (δ) are indicated in parts per million (ppm) to TMS or phosphate and referenced to used solvents. Abbreviations for signal multiplicity are s = singlet, d = doublet, t = triplet, q = quartet, br = broad signal, m = multiplet. $^{13}C$ signals only obtained from HSQC experiments are marked with "*", signals corresponding to triethylammonium counterions are marked with "#".

High-resolution mass spectrometry (HR-MS) spectra were recorded on a Bruker microTOF-QII system (Bruker, Billerica, MA, USA) operated in electron spray ionization (ESI) positive or negative mode. UV-Vis spectra were recorded on a Cary 50 or a Cary 100 Bio UV-VIS spectrophotometer (Varian Medical Systems, Palo Alto, CA, USA).

Gel electrophoresis of oligonucleotides was performed using Tris-borate EDTA (TBE) buffer (0.1 M Tris-borate, 20 mM EDTA, pH 8.3) and a power of 25 W. Denaturing polyacrylamide gels (8.3 M urea, 0.1 M Tris-borate, 20 mM EDTA, pH 8.3) were prepared by diluting Rotiphorese Sequencing gel concentrate (25%, 19:1; Carl Roth, Karlsruhe Germany). Native polyacrylamide gels (0.1 M Tris-borate, 20 mM EDTA, 40 mM NaCl, pH 8.3) were prepared by diluting Rotiphorese gel 40 (40%, 19:1; Carl Roth, Karlsruhe Germany). Gels were stained with SYBR Gold (Invitrogen, Carlsbad, CA, USA) and scanned on a Typhoon FLA 9500 Biomolecular Imager (Cytiva, Marlborough, MA, USA) or bands were identified by UV shadowing over TLC Silica gel 60 $F_{254}$ aluminum sheets (Merck Millipore, Burlington, MA, USA). For size comparison, low range ssRNA Ladder (50 – 1,000 bases, NEB, Ipswich, MA, USA) was used. Densitometric quantifications were conducted using Image Quant TL 1D (version 8.1, GE Healthcare, Chicago, IL, USA). Protein structures were visualized using PyMOL Molecular Graphics System (Version 2.6.0a0, Schrödinger).

Bacterial cultures for protein expression were grown in a MaxQ 8000 incubator (Thermo Fischer Scientific, Waltham, MA, USA). Fast protein liquid chromatography (FPLC) was conducted using an NGC 10 Medium-Pressure Chromatography System (Bio-Rad, Hercules, CA, USA). Protein and nucleic acid concentrations have been determined using Nanodrop One (Thermo Fischer Scientific, Waltham, MA, USA).

### 3 Chemical Synthesis

#### 3.1 Reverse phase (RP) chromatography and RP-HPLC

##### 3.1.1. Mobile phase

Reverse phase (RP) chromatography was either conducted using water / acetonitrile (ACN) as mobile phase or a triethylammonium acetate (TEAA) buffer system. The latter was prepared by diluting a 1 M TEAA stock solution (pH 7) either with Milli-Q water (buffer A) or ACN (buffer B). While used buffer concentration for RP chromatography was rather low (10 mM) to reduce lyophilization cycles after purification, a higher concentration (100 mM) of buffer A was added to samples before purification to ensure proper pH adjustment. Chromatograms were recorded at 260 nm.

##### 3.1.2 General procedure for capped dinucleotide purification

After reactions were completed and quenched, Celite was added and the solvent was evaporated under reduced pressure. The crude product was purified using a PuriFlash PF420 (Interchim) system equipped with a 12 g PF-30C18AQ column (Interchim) and buffers A and B as mobile phase. Fractions containing the product were lyophilized and the residue was repeatedly dissolved in Milli-Q water and lyophilized until a stoichiometric amount of triethylammonium cations was present according to $^{1}$H NMR measurements.

For final products, if necessary, additional semi-preparative HPLC purification was conducted (1% buffer B for 5 min, to 5.5% buffer B in 25 min, flow rate: 5 ml/min). Furthermore, triethylammonium cations were exchanged for sodium by dissolving capped dinucleotides in Milli-Q water and passing them through a short column of Amberlite IR120 $Na^{+}$ form. Fractions containing the product were identified by NanoDrop measurement, combined and lyophilized.

#### 3.3 Synthetic procedures

##### 2-Acetamido-3,4,6-tri-*O*-acetyl-2-deoxy-α-D-glucopyranose 1-phosphate (1)

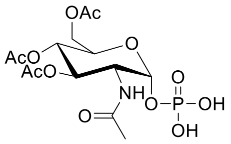

Compound **1** was synthesized using a modified literature procedure (12,13). Compound **17** (4.40 g, 7.24 mmol, 1.00 eq) was dissolved in anh. MeOH (150 ml) under an argon atmosphere. 10% Palladium on charcoal was added (1.20 g) and the atmosphere was exchanged with hydrogen. The mixture was stirred overnight and filtered through a plug of silica gel. The plug was washed with EtOAc / MeOH (3 / 1) and the combined organic layer was evaporated *in vacuo* giving compound **1** as white crystals (3.00 g, 7.03 mmol, 97%).

**$^{1}$H NMR** (500 MHz, DMSO-$d_6$) δ 7.97 (d, J = 8.5 Hz, 1H), 5.41 (dd, J = 6.7, 3.2 Hz, 1H), 5.16 (dd, J = 11.1, 9.3 Hz, 1H), 4.95 (t, J = 9.7 Hz, 1H), 4.17 (dd, J = 12.2, 3.9 Hz, 1H), 4.14 – 3.98 (m, 3H), 2.02 (s, 3H), 1.98 (s, 3H), 1.93 (s, 3H), 1.82 (s, 3H).

**$^{13}$C NMR** (126 MHz, DMSO-$d_6$) δ 170.05, 169.96, 169.80, 169.24, 93.56 (d, J = 6.0 Hz), 69.51, 68.38, 68.21, 61.26, 51.12 (d, J = 8.2 Hz), 22.30, 20.50, 20.45, 20.39.

**$^{31}$P NMR** (202 MHz, DMSO-$d_6$) δ -2.71.

**HRMS** (ESI, negative): Calculated for $C_{14}H_{21}NO_{12}P^{-}$ $[M-H]^{-}$ 426.0807; found: 426.0807.

**2-(2-Azidoacetamido)-3,4,6-tri-*O*-benzoyl-2-deoxy-α-D-glucopyranose 1-phosphate (2)**

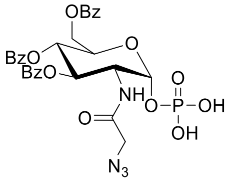

Compound **24** (as mono(triethylammonium) salt) (100 mg, 134 µmol, 1.00 eq) and sodium azide (50.2 mg, 772 µmol, 5.00 eq) were suspended in MeOH (10 ml) and the mixture was heated to 70°C overnight. Excess sodium azide was filtered off, Celite was added and the solvent was removed under reduced pressure. The residue was purified by reverse phase chromatography using a 12 g PF-30C18AQ column (interchim) and 10 mM buffer A / B as mobile phase (gradient: 5% buffer B for 10 min, to 80% buffer B in 50 min, flow rate 15 ml/min). The mono(triethylammonium) salt of **2** was obtained as a white powder (71.5 mg, 94.6 µmol, 71%).

**$^1$H NMR** (500 MHz, Methanol-*d*$_4$) δ 8.05 – 8.00 (m, 2H), 7.90 – 7.84 (m, 4H), 7.57 (t, 1H), 7.54 – 7.48 (m, 2H), 7.43 (t, J = 7.6 Hz, 2H), 7.36 (t, J = 7.7 Hz, 4H), 5.85 (t, J = 10.1 Hz, 1H), 5.75 – 5.67 (m, 2H), 4.73 (d, J = 10.1 Hz, 1H), 4.64 – 4.58 (m, 2H), 4.49 – 4.43 (m, 1H), 3.84 (d, J = 15.8 Hz, 1H), 3.71 (d, J = 15.8 Hz, 1H), 3.17 (q, J = 7.3 Hz, 6H)#, 1.30 (t, J = 7.3 Hz, 9H)#.

**$^{13}$C NMR** (126 MHz, Methanol-*d*$_4$) δ 170.63, 167.58, 167.57, 166.75, 134.57, 134.41, 134.26, 131.06, 130.74, 130.72, 130.69, 130.55, 130.42, 129.54, 129.51, 129.48, 95.14 (d, J = 4.2 Hz), 73.31, 71.17, 69.85, 64.02, 53.81 (d, J = 6.7 Hz), 52.57, 47.62#, 9.21#.

**$^{31}$P NMR** (202 MHz, Methanol-*d*$_4$) δ -0.97.

**HRMS** (ESI, negative): Calculated for $C_{29}H_{26}N_4O_{12}P^-$ $[M-H]^-$ 653.1290; found: 653.1303.

**Protected pUpG building block (3)**

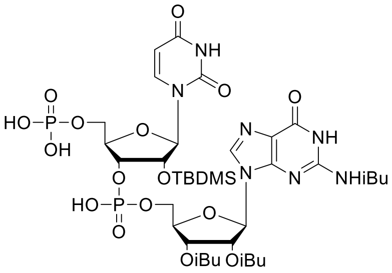

The synthetic procedure towards protected pUpG building block **3** was inspired by literature (14). Therefore, protected UpG building block **26** (1.52 g, 1.57 mmol, 1.00 eq) was dissolved in anh. ACN (10 ml) under an argon atmosphere. A tetrazole solution in anh. ACN (0.45 mM, 7.00 ml, 3.14 mmol, 2.00 eq) and bis(2-cyanoethyl)-*N*,*N*-diisopropyl phosphoramidite (0.49 ml, 512 mg, 1.89 mmol, 1.2 eq) was added dropwise. The mixture was heated to 40°C until full conversion according to TLC (MeOH / DCM 1 / 9).

An $I_2$ solution (0.1 M) in THF / pyridine / $H_2O$ (2 / 2 / 1) was added dropwise until the brownish color persisted. Excess $I_2$ was quenched by the addition of a $Na_2S_2O_3$ solution (1 M, 50 ml). The aqueous layer was extracted with DCM (3 x 50 ml) and the combined organic layers were dried over $MgSO_4$ and filtered. The solvent was removed under reduced pressure giving the intermediate product as a yellowish foam.

The intermediate was dissolved in anh. ACN (10 ml) and cooled to 0°C. DBU (0.70 ml, 717 mg, 4.71 mmol, 3.00 eq) and bis(trimethylsilyl)acetamide (BSA) (8.64 ml, 7.18 g, 35.3 mmol, 22.5 eq) were added dropwise. The reaction was warmed to RT and stirred for 1 h. It was cooled to 0°C and a mixture of TFA in MeOH (1 / 3; 3 ml) was added. The mixture was stirred for 10 min at RT, cooled again to 0°C and buffer A was added (100 mM, 30 ml). Celite was added and the solvent was evaporated under reduced pressure. The product was purified using two C18 23 g flash columns (Telos) and buffer A / B (10 mM) as mobile phase (gradient: 5% buffer B for 10 min, to 70 % B in 50 min, flow rate 15 ml/min). Fractions containing the product were lyophilized and excess buffer was removed by repeated dissolving in Milli-Q water and lyophilization. Compound **3** was isolated as tris(triethylammonium) salt. The product was a white foam (1.32 g, 1.03 mmol. 65%).

**$^1$H NMR** (500 MHz, Deuterium Oxide) δ 8.19 (s, 1H), 7.91 (d, J = 8.1 Hz, 1H), 6.23 (d, J = 5.1 Hz, 1H), 6.07 (t, J = 5.5 Hz, 1H), 5.98 (d, J = 8.1 Hz, 1H), 5.89 (d, J = 7.2 Hz, 1H), 5.76 (t, J = 5.3 Hz, 1H), 4.64 – 4.55 (m, 2H), 4.39 – 4.32 (m, 2H), 4.24 (t, J = 4.4 Hz, 2H), 3.94 – 3.89 (m, 1H), 3.74 – 3.68 (m, 1H), 3.21 (q, J = 7.3 Hz, 18H)#, 2.86 (hept, J = 6.9 Hz, 1H), 2.76 (hept, J = 7.0 Hz, 1H), 2.64 (hept, J = 7.0 Hz, 1H), 1.29 (t, J = 7.3 Hz, 27H)#, 1.25 (d, J = 7.1 Hz, 6H), 1.22 (d, J = 6.7 Hz, 6H), 1.11 (d, J = 7.0 Hz, 3H), 1.09 (d, J = 7.0 Hz, 3H), 0.77 (s, 9H), 0.02 (s, 6H).

**$^{13}$C NMR** (126 MHz, Deuterium Oxide) δ 182.47, 178.70, 178.36, 166.04, 157.27, 151.52, 149.25, 148.06, 141.59, 140.76, 120.63, 103.15, 87.13, 86.90, 83.25 (d, J = 9.2 Hz), 81.43 (d, J = 9.5 Hz), 75.23 (d, J = 5.5 Hz), 74.97 (d, J = 7.1 Hz), 72.87, 70.99, 65.40 (d, J = 4.9 Hz), 64.31 (d, J = 4.8 Hz), 46.73#, 35.70, 33.80, 33.65, 24.98, 18.43, 18.27, 18.25, 18.08, 18.04, 18.02, 17.39, 8.30#, -5.52, -6.08.

**$^{31}$P NMR** (202 MHz, Deuterium Oxide) δ 0.67, -1.05.

**HRMS** (ESI, negative): Calculated for $C_{37}H_{56}N_7O_{19}P_2Si^-$ $[M-H]^-$ 992.2881; found: 992.2889.

**GlcNAcppUpG (4)**

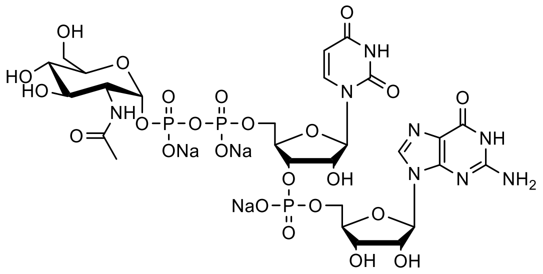

Protected pUpG building block **3** (as tris(triethylammonium) salt) (200 mg, 0.15 mmol, 1.00 eq) and CDI (60.0 mg, 0.37 mmol, 2.40 eq) were dissolved in anh. DMF (3 ml) under an argon atmosphere. The reaction progress was monitored using RP-HPLC. After approximately 3 h, the reaction was completed.

The mixture was directly added to a mixture of compound **1** (294 mg, 0.62 mmol, 4.00 eq) and $MgCl_2$ (88.0 mg, 0.93 mmol, 6.00 eq) dissolved in anh. DMF (3 ml) at 0°C. The reaction was allowed to warm to RT and it was stirred overnight. It was cooled to 0°C and quenched by the addition of an EDTA solution (500 mM, pH 8, 2 ml, 1 mmol) and buffer A (10 ml, 100 mM). The first intermediate was purified using the general procedure for capped dinucleotide purification (gradient: 0% buffer B for 10 min, to 80% buffer B in 50 min, flow rate 15 ml/min). The protected intermediate was isolated as mono(triethylammonium) salt and was a white solid (132 mg, 87.7 µmol, 58%).

**$^1$H NMR** (500 MHz, Deuterium Oxide) δ 8.24 (s, 1H), 7.94 (d, J = 8.1 Hz, 1H), 6.17 (d, J = 5.0 Hz, 1H), 5.97 (d, J = 8.0 Hz, 1H), 5.92 (t, J = 5.3 Hz, 1H), 5.86 (d, J = 5.0 Hz, 1H), 5.69 – 5.66 (m, 1H), 5.62 (dd, J = 7.0, 3.3 Hz, 1H), 5.22 (t, J = 9.9 Hz, 1H), 5.10 (t, J = 9.7 Hz, 1H), 4.58 – 4.52 (m, 2H), 4.46 (s, 1H), 4.42 – 4.31 (m, 4H), 4.29 – 4.11 (m, 5H), 3.21 (q, J = 7.3 Hz, 7H)#, 2.89 – 2.81 (m, 1H), 2.73 – 2.64 (m, 1H), 2.62 – 2.53 (m, 1H), 2.11 (s, 3H), 2.05 (s, 3H), 2.01 (s, 3H), 2.01 (s, 3H), 1.28 (t, J = 7.3 Hz, 11H)#, 1.22 – 1.15 (m, 12H), 1.09 – 1.03 (m, 6H), 0.76 (s, 9H), 0.03 (s, 3H), 0.02 (s, 3H).

**$^{13}$C NMR** (126 MHz, Deuterium Oxide) δ 182.04, 177.86, 177.49, 174.40, 173.51, 172.80, 172.58, 165.87, 156.98, 151.24, 149.17, 148.11, 141.56, 120.37, 102.67, 94.15, 88.37, 86.81, 82.18, 81.67, 74.78, 73.23, 71.60, 70.95, 68.36, 68.21, 65.57, 65.19, 61.70, 51.55 (d, J = 8.9 Hz), 46.73#, 35.66, 33.76, 33.61, 25.19, 22.02, 20.21, 20.18, 20.09, 18.48, 18.45, 18.22, 18.16, 17.53, 8.32#, -5.32, -5.73.

**$^{31}$P NMR** (202 MHz, Deuterium Oxide) δ -1.17, -12.01 (d, J = 18.0 Hz), -13.61 (d, J = 18.3 Hz).

**HRMS** (ESI, negative): Calculated for $C_{51}H_{75}N_8O_{30}P_3Si^{2-}$ $[M-2H]^{2-}$ 700.1791; found: 700.1819.

The intermediate was dissolved in 28% $NH_4OH$ in $H_2O$ (2 ml) and 33% methylamine in EtOH (2 ml). The mixture was heated at 50°C for 1.5 h and the solvent was removed under reduced pressure. The residue was dissolved in buffer A (100 mM, 2 ml) and purified using the general procedure for capped dinucleotide purification (gradient: 0% buffer B for 10 min, to 50% buffer B in 50 min, flow rate 15 ml/min). The TBDMS-protected intermediate GlcNAcppU(TBDMS)pG was isolated as tris(triethylammonium) salt and was a white powder (114 mg, 83.2 µmol, 95%).

**$^{1}$H NMR** (500 MHz, Deuterium Oxide) δ 8.02 (s, 1H), 7.88 (d, J = 8.1 Hz, 1H), 5.99 (d, J = 8.1 Hz, 1H), 5.94 (d, J = 7.1 Hz, 1H), 5.89 (d, J = 5.4 Hz, 1H), 5.54 (dd, J = 7.2, 3.3 Hz, 1H), 4.92 (t, J = 5.3 Hz, 1H), 4.60 – 4.56 (m, 1H), 4.55 – 4.51 (m, 1H), 4.46 – 4.43 (m, 1H), 4.35 – 4.30 (m, 2H), 4.20 – 4.11 (m, 3H), 4.07 (ddd, J = 11.7, 4.9, 2.5 Hz, 1H), 3.99 (dt, J = 10.5, 3.1 Hz, 1H), 3.96 (ddd, J = 10.1, 4.4, 2.3 Hz, 1H), 3.88 (dd, J = 12.5, 2.3 Hz, 1H), 3.85 – 3.79 (m, 2H), 3.56 (dd, J = 10.1, 9.1 Hz, 1H), 3.20 (q, J = 7.3 Hz, 18H)#, 2.07 (s, 3H), 1.28 (t, J = 7.3 Hz, 27H)#, 0.77 (s, 9H), 0.03 (s, 3H), 0.01 (s, 3H).

**$^{13}$C NMR** (126 MHz, Deuterium Oxide) δ 174.79, 165.98, 158.92, 153.82, 151.88, 151.64, 141.37, 137.97*, 116.65, 103.31, 94.61 (d, J = 6.3 Hz), 87.61, 86.84, 83.61 (d, J = 9.1 Hz), 82.99 (d, J = 9.1 Hz), 74.88 (m), 73.11, 73.00, 70.97, 70.65, 69.60, 65.80 (d, J = 5.2 Hz), 65.56 (d, J = 5.0 Hz), 60.39, 53.76 (d, J = 8.6 Hz), 46.74#, 24.98, 23.31#, 22.19, 17.41, 8.30#, -5.55, -6.08.

**$^{31}$P NMR** (202 MHz, Deuterium Oxide) δ -0.98, -11.92 (d, J = 18.9 Hz), -13.05 (d, J = 19.1 Hz).

**HRMS** (ESI, negative): Calculated for $C_{33}H_{52}N_8O_{24}P_3Si^{-}$ $[M-H]^{-}$ 1065,2082; found: 1065.2083.

GlcNAcppU(TBDMS)pG (as tris(triethylammonium) salt) (104 mg, 75.9 µmol, 1.00 eq) was dissolved in DMF (3 ml), $Et_3N$ (1 ml) and $Et_3N$*3HF (2 ml) in a falcon tube. The reaction was stirred for 2 h at 50°C. The mixture was cooled to 0°C and buffer A (100 mM, 20 ml) was added. The product was purified according to the general procedure for capped dinucleotide purification (gradient: 0% buffer B for 20 min, to 30% buffer B in 40 min, flow rate 15 ml/min) and the sodium salt of GlcNAcppUpG **4** was isolated as a white powder (67.6 mg, 66.4 µmol, 87%).

**$^{1}$H NMR** (500 MHz, Deuterium Oxide) δ 8.03 (s, 1H), 7.88 (d, J = 8.1 Hz, 1H), 5.94 (d, J = 8.2 Hz, 1H), 5.92 – 5.89 (m, 2H), 5.53 (dd, J = 7.3, 3.3 Hz, 1H), 4.83 (t, J = 5.4 Hz, 1H), 4.70 – 4.63 (m, 1H), 4.55 – 4.51 (m, 1H), 4.42 – 4.34 (m, 3H), 4.25 – 4.12 (m, 4H), 4.01 (dt, J = 10.5, 3.0 Hz, 1H), 3.96 (ddd, J = 10.1, 4.4, 2.3 Hz, 1H), 3.89 (dd, J = 12.5, 2.3 Hz, 1H), 3.86 – 3.79 (m, 2H), 3.56 (t, J = 9.6 Hz, 1H), 2.08 (s, 3H).

**$^{13}$C NMR** (126 MHz, Deuterium Oxide) δ 174.83, 166.07, 158.90, 153.87, 151.76, 151.75, 141.25, 137.81, 116.58, 102.91, 94.58 (d, J = 6.2 Hz), 87.80, 87.48, 83.68 (d, J = 9.1 Hz), 82.35 (dd, J = 8.9, 3.6 Hz), 74.43 (d, J = 5.4 Hz), 73.36, 73.18 (d, J = 5.0 Hz), 73.09, 71.04, 70.47, 69.62, 65.46 (d, J = 5.2 Hz), 65.18 (d, J = 5.6 Hz), 60.39, 53.76 (d, J = 8.6 Hz), 22.17.

**$^{31}$P NMR** (202 MHz, Deuterium Oxide) δ -0.59, -11.37 (d, J = 20.3 Hz), -12.87 (d, J = 20.5 Hz).

**HRMS** (ESI, negative): Calculated for $C_{27}H_{37}N_8O_{24}P_3^{2-}$ $[M-2H]^{2-}$ 475.0572; found: 475.0585.

**GlcNAzppUpG (5)**

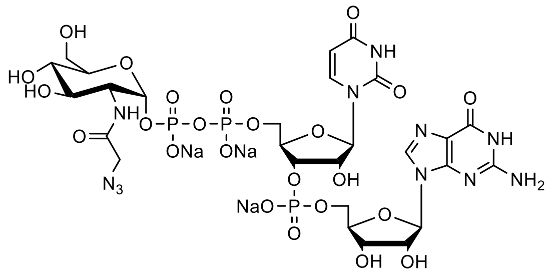

Compound **2** (as bis(triethylammonium) salt) (50.0 mg, 58.4 µmol, 1.00 eq) and CDI (22.7 mg, 149 µmol, 2.40 eq) were dissolved in anh. DMF (3 ml) under an argon atmosphere. The mixture was stirred at RT for 3 h and directly added to a mixture of protected pUpG building block **3** (as tris(triethylammonium) salt) (114 mg, 87.5 µmol, 1.50 eq) and $MgCl_2$ (33.3 mg, 350 µmol, 6.00 eq) in anh. DMF (3 ml) at 0°C. The reaction was allowed to warm to RT and it was stirred overnight. The reaction was cooled to 0°C and quenched by the addition of an EDTA solution (500 mM, pH 8, 2.00 ml, 1.00 mmol) and buffer A (10 ml, 100 mM). The intermediate product was partially purified using the general procedure for capped dinucleotide purification (gradient: 0% buffer B for 10 min, to 80% buffer B in 50 min, flow rate 15 ml/min).

The intermediate was dissolved in 28% $NH_4OH$ in $H_2O$ (2 ml) and 33% methylamine in EtOH (2 ml). The mixture was heated to 50°C for 1.5 h and the solvent was removed under reduced pressure. The residue was dissolved in buffer A (100 mM, 2 ml) and purified using the general procedure for capped dinucleotide purification (gradient: 0% buffer B for 10 min, to 50% Buffer B in 50 min, flow rate 15 ml/min). The TBDMS-protected intermediate GlcNAzppU(TBDMS)pG was isolated as the bis(triethylammonium) salt and was a white powder (15.7 mg, 12.0 µmol, 21%).

**$^1$H NMR** (500 MHz, Deuterium Oxide) δ 8.02 (s, 1H), 7.91 (d, J = 8.1 Hz, 1H), 6.01 (d, J = 8.1 Hz, 1H), 5.96 (d, J = 7.2 Hz, 1H), 5.90 (d, J = 5.5 Hz, 1H), 5.56 (br, 1H), 4.95 (t, J = 5.4 Hz, 1H), 4.58 (br, 1H), 4.55 – 4.51 (m, 1H), 4.47 (s, 1H), 4.37 – 4.30 (m, 2H), 4.21 – 4.04 (m, 7H), 4.01 – 3.94 (m, 1H), 3.93 – 3.80 (m, 3H), 3.57 (t, J = 9.6 Hz, 1H), 3.21 (q, J = 7.4 Hz, 13H)#, 1.29 (t, J = 7.3 Hz, 20H)#, 0.78 (s, 9H), 0.04 (s, 3H), 0.02 (s, 3H).

**$^{13}$C NMR** (126 MHz, Deuterium Oxide) δ 171.02, 166.01, 153.83, 151.67, 141.43, 103.36, 94.53 (d, J = 5.2 Hz), 87.54, 86.80, 83.69 (d, J = 8.8 Hz), 83.05 (d, J = 7.7 Hz), 74.98, 74.94, 73.13, 72.90, 71.02, 70.72, 69.58, 65.81 (d, J = 5.1 Hz), 65.61, 60.40, 53.84 (d, J = 7.8 Hz), 51.66, 46.74#, 24.99, 17.42, 8.30#, -5.55, -6.10.

**$^{31}$P NMR** (202 MHz, Deuterium Oxide) δ -0.87, -11.72, -12.95.

**HRMS** (ESI, negative): Calculated for $C_{33}H_{51}N_{11}O_{24}P_3Si^-$ $[M-H]^-$ 1106.2096; found: 1106.2100.

GlcNAzppU(TBDMS)pG (as bis(triethylammonium) salt) (29.2 mg, 22.3 µmol, 1.00 eq) was dissolved in DMF (1.5 ml), $Et_3N$ (0.5 ml) and $Et_3N$*3HF (1 ml) in a falcon tube. The mixture was stirred for 2 h at 50°C. It was cooled to 0°C and buffer A (100 mM, 20 ml) was added. The product was purified according to the general procedure for capped dinucleotide purification (gradient: 0% Buffer B for 20 min, to 30% Buffer B in 40 min, flow rate 15 ml/min). The sodium salt of GlcNAzppUpG **5** was isolated as a white powder (20.2 mg, 19.1 µmol, 86%).

**$^1$H NMR** (500 MHz, Deuterium Oxide) δ 8.03 (s, 1H), 7.88 (d, J = 8.1 Hz, 1H), 5.94 (d, J = 8.1 Hz, 1H), 5.92 – 5.89 (m, 2H), 5.56 (dd, J = 7.2, 3.3 Hz, 1H), 4.83 (t, J = 5.4 Hz, 1H), 4.69 – 4.62 (m, 1H), 4.56 – 4.50 (m, 1H), 4.42 – 4.37 (m, 2H), 4.37 – 4.32 (m, 1H), 4.26 – 4.03 (m, 7H), 3.99 – 3.93 (m, 1H), 3.92 – 3.79 (m, 3H), 3.60 – 3.54 (m, 1H).

**$^{13}$C NMR** (126 MHz, Deuterium Oxide) δ 171.01, 166.15, 158.95, 153.91, 151.80, 151.77, 141.26, 137.74*, 116.59, 102.91, 94.47 (d, J = 6.2 Hz), 87.87, 87.47, 83.68 (d, J = 9.1 Hz), 82.33 (dd, J = 9.3, 3.8 Hz), 74.39 (d, J = 5.5 Hz), 73.37, 73.17 (d, J = 4.9 Hz), 73.12, 70.98, 70.46, 69.58, 65.45 (d, J = 5.3 Hz), 65.22 (d, J = 5.5 Hz), 60.38, 53.81 (d, J = 8.6 Hz), 51.68.

**$^{31}$P NMR** (202 MHz, Deuterium Oxide) δ -0.59, -11.35 (d, J = 20.4 Hz), -12.90 (d, J = 20.3 Hz).

**HRMS** (ESI, negative): Calculated for $C_{27}H_{37}N_{11}O_{24}P_3^-$ [M-H]$^-$ 992.1231; found: 992.1271.

**EP-GlcNAcppUpG (6)**

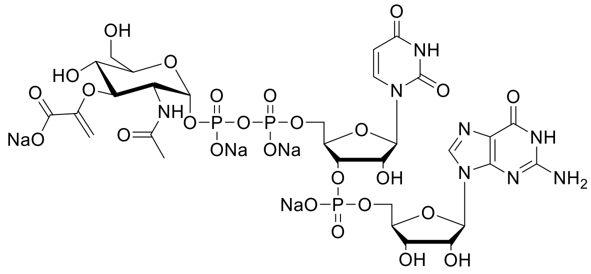

EP-GlcNAcppUpG **6** was synthesized using a modified literature procedure (2). GlcNAcppUpG **4** (23.0 mg, 22.6 µmol, 1.00 eq) was dissolved in Milli-Q water (6.08 ml). Tris-HCl (1 M, pH 7.8, 0.32 ml) and phosphoenolpyruvate (100 mM, 0.80 ml, 80.0 µmol, 3.54 eq) were added. It was mixed and the mixture was distributed to eight 1.5 ml Eppendorf tubes (900 µl each). MurA (1 mg / ml, 100 µl) was added to each Eppendorf tube, it was mixed by pipetting and incubated at 37°C, 300 rpm overnight.

The reaction mixture was collected in a round-bottom flask and buffer A (100 mM, 10 ml) was added. The product was purified according to the general procedure for capped dinucleotide purification (gradient: 0% buffer B for 20 min, to 30% buffer B in 40 min, flow rate 15 ml/min) and the sodium salt of EP-GlcNAcppUpG **6** was isolated as white powder (16.2 mg, 14.6 µmol, 65%).

**$^{1}$H NMR** (500 MHz, Deuterium Oxide) δ 8.03 (s, 1H), 7.88 (d, J = 8.2 Hz, 1H), 5.94 (d, J = 8.1 Hz, 1H), 5.92 – 5.88 (m, 2H), 5.55 (dd, J = 7.2, 3.3 Hz, 1H), 5.24 (d, J = 2.7 Hz, 1H), 4.90 (d, J = 2.7 Hz, 1H), 4.83 (t, J = 5.4 Hz, 1H), 4.69 – 4.62 (m, 1H), 4.56 – 4.50 (m, 1H), 4.45 – 4.33 (m, 4H), 4.28 – 4.12 (m, 5H), 4.06 – 3.99 (m, 1H), 3.93 – 3.77 (m, 3H), 1.99 (s, 3H).

**$^{13}$C NMR** (126 MHz, Deuterium Oxide) δ 174.57, 170.70, 166.11, 158.96, 155.85, 153.92, 151.79, 151.77, 141.25, 138.40*, 116.59, 102.90, 94.83 (d, J = 6.2 Hz), 94.45, 87.88, 87.45, 83.70 (d, J = 9.1 Hz), 82.29 (dd, J = 9.0, 3.6 Hz), 78.54, 74.36 (d, J = 5.6 Hz), 73.36, 73.17 (d, J = 4.8 Hz), 72.91, 70.48, 68.65, 65.44 (d, J = 5.3 Hz), 65.20 (d, J = 5.6 Hz), 60.32, 53.01 (d, J = 8.7 Hz), 22.14.

**$^{31}$P NMR** (202 MHz, Deuterium Oxide) δ -0.74, -11.32 (d, J = 20.4 Hz), -13.12 (d, J = 20.3 Hz).

**HRMS** (ESI, negative): Calculated for $C_{30}H_{40}N_8O_{26}P_3^-$ [M-H]$^-$ 1021.1272; found: 1021.1261.

**MurNAcppUpG (7)**

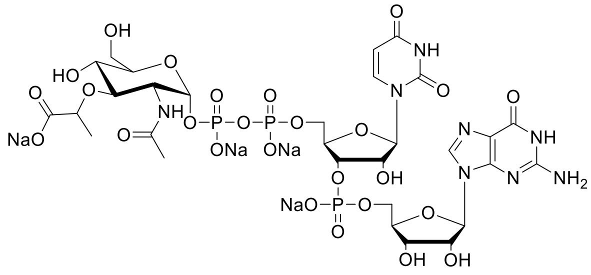

MurNAcppUpG **7** was synthesized using a modified literature procedure (15). EP-GlcNAcppUpG **6** (15.7 mg 14.4 µmol, 1.00 eq) was dissolved in Milli-Q water (9.60 ml). Tris-HCl (1 M, pH 7.8, 0.48 ml), KCl (50 mM, 0.36 ml), DTT (100 mM, 0.12 ml) and NADPH (50 mM, 0.96 ml, 48.0 µmol, 3.30 eq) was added and the mixture was distributed to eight 1.5 ml

Eppendorf tubes (1.44 ml each). MurB (1 mg/ml, 0.06 ml) was added to each Eppendorf tube, it was mixed by pipetting and incubated at 37°C, 300 rpm overnight.

Reaction mixtures were combined in a round-bottomed flask and buffer A (100 mM, 10 ml) was added. The product was purified according to the general procedure for capped dinucleotide purification (gradient: 0% buffer B for 20 min, to 30% buffer B in 40 min, flow rate 15 ml/min) and the sodium salt of MurNAcppUpG **7** was isolated as white powder (8.30 mg, 7.46 µmol, 52%).

**$^{1}$H NMR** (500 MHz, Deuterium Oxide) δ 7.89 (s, 1H), 7.76 (d, J = 8.2 Hz, 1H), 5.80 (d, J = 8.1 Hz, 1H), 5.77 – 5.75 (m, 2H), 5.50 (dd, J = 7.2, 3.2 Hz, 1H), 4.70 (t, J = 5.5 Hz, 1H), 4.54 – 4.49 (m, 1H), 4.38 (dd, J = 5.2, 3.7 Hz, 1H), 4.28 – 4.19 (m, 3H), 4.14 (q, J = 6.9 Hz, 1H), 4.10 – 3.96 (m, 4H), 3.83 – 3.75 (m, 2H), 3.74 – 3.64 (m, 3H), 3.52 – 3.44 (m, 1H), 1.93 (s, 3H), 1.21 (d, J = 6.9 Hz, 3H).

**$^{13}$C NMR** (126 MHz, Deuterium Oxide) δ 181.41, 174.33, 165.96, 158.85, 153.78, 151.64, 141.18, 138.21*, 116.48, 102.78, 93.94 (d, J = 6.2 Hz), 87.63, 87.28, 83.64 (d, J = 9.0 Hz), 82.19 (dd, J = 8.6, 3.9 Hz), 77.51, 77.18, 74.19 (d, J = 5.4 Hz), 73.17, 73.02 (d, J = 4.7 Hz), 72.90, 70.38, 68.95, 65.29 (d, J = 5.2 Hz), 64.90 (d, J = 5.2 Hz), 60.15, 53.01 (d, J = 8.7 Hz), 22.09, 18.72.

**$^{31}$P NMR** (202 MHz, Deuterium Oxide) δ -0.73, -11.61 (d, J = 19.6 Hz), -13.06 (d, J = 19.5 Hz).

**HRMS** (ESI, negative): Calculated for $C_{30}H_{42}N_8O_{26}P_3^-$ [M-H]$^-$ 1023.1429; found: 1023.1410.

**UDP-GlcNAc-EP (14)**

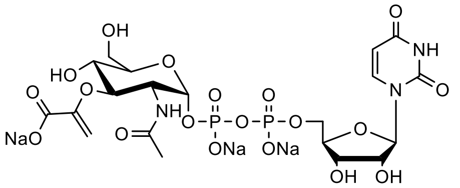

UDP-GlcNAc-EP **14** was synthesized using a modified literature procedure (2). UDP-GlcNAc sodium salt (5.00 mg, 7.68 µmol, 1.00 eq) was dissolved in Milli-Q water (2.085 ml). Tris-HCl (1 M, pH 7.8, 0.3 ml) and phosphoenolpyruvate (100 mM, 315 µl, 31.5 µmol, 4.1 eq) was added. It was mixed and the solution was distributed to three 1.5 ml Eppendorf tubes (900 µl each). MurA (1 mg / ml, 100 µl) was added to each Eppendorf tube, it was mixed by pipetting and incubated at 37°C, 300 rpm overnight. The enzyme was removed by filtration using a 10 kDa MW cut-off filter and the filtrate was purified by semi-preparative HPLC using a Luna 5µm C18(2) 100 Å 250 x 21.2 mm column (gradient: 1% buffer B for 5 min, to 5% buffer B in 60 min, flow rate 5 ml/min).

The product was lyophilized, and excess buffer was removed by repeated dissolution in Milli-Q water and lyophilization. The product was dissolved in Milli-Q water (1 ml) and triethylammonium cations were exchanged for sodium by passing it through a short column of Amberlite IR120 Na$^+$ form. Fractions containing the product were identified using NanoDrop measurement, combined and lyophilized. The product was isolated as a white powder (3.2 mg, 4.30 µmol, 56%) and was analyzed by RP-HPLC and ESI-MS.

**HRMS** (ESI, negative): Calculated for $C_{20}H_{28}N_3O_{19}P_2^-$ [M-H]$^-$ 676.0798; found: 676.0786.

**UDP-MurNAc (15)**

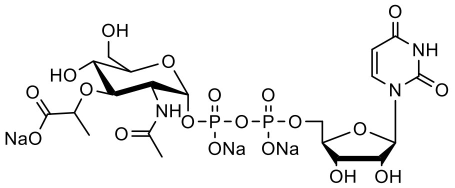

UDP-MurNAc **15** was synthesized using a modified literature procedure (15). UDP-GlcNAc sodium salt (10.0 mg 15.4 µmol, 1.00 eq) was dissolved in Milli-Q water (3.475 ml). Tris-HCl (1 M, pH 7.8, 0.5 ml) and phosphoenolpyruvate (100 mM, 0.525 ml, 52.5 µmol, 3.42 eq) were added. It was mixed and the solution was distributed to five 1.5 ml Eppendorf tubes (900 µl each). MurA (1 mg/ml, 100 µl) was added to each Eppendorf tube, it was mixed by pipetting and incubated at 37°C, 300 rpm for 4 h.

Milli-Q water (182 µl), KCl (50 mM, 45.0 µl), DTT (100 mM, 15.0 µl), NADPH (50 mM, 198 µl, 49.5 µmol (total), 3.22 eq) and MurB (1 mg/ml, 0.06 ml) was added to each Eppendorf tube. After mixing by pipetting, it was incubated at 37°C, 300 rpm overnight. The enzymes were removed by filtration using a 10 kDa MW cut-off filter and the filtrate was purified as described for UDP-GlcNAc-EP **14**. The sodium salt of UDP-MurNAc **15** was isolated as a white powder (5.10 mg, 6.84 µmol, 45%) and was analyzed using RP-HPLC and ESI-MS.

**HRMS** (ESI, negative): Calculated for $C_{20}H_{30}N_3O_{19}P_2^-$ [M-H]$^-$ 678.0954; found: 678.0949.

**2-Methyl-(3,4,6-tri-*O*-acetyl-1,2-dideoxy-α-D-glucopyrano)[1,2-*d*]-oxazoline (16)**

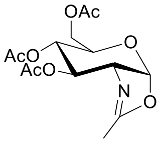

Compound **16** was synthesized using a modified literature procedure (12). Pentaacetyl-β-D-glucosamin (10.0 g, 25.7 mmol, 1.00 eq) was dissolved in anh. 1,2-dichloroethane (DCE) (200 ml) under an argon atmosphere. TMSOTf (6.28 g, 5.10 ml, 28.3 mmol, 1.10 eq) was added dropwise and the mixture was heated at 60°C overnight.

The reaction was cooled on ice and triethylamine (7.80 g, 10.7 ml, 77.1 mmol, 3.00 eq) was added dropwise. It was stirred for another 10 min at RT and filtered through a small plug of silica gel. The solvent was evaporated under reduced pressure and the residue was purified by silica gel chromatography using EtOAc / cyclohexane / $Et_3N$ (80 / 20 / 1) as a mobile phase. Compound **16** was isolated as a yellowish oil (8.15 g, 24.7 mmol, 96%).

**$^1$H NMR** (500 MHz, Chloroform-*d*) δ 5.96 (d, J = 7.4 Hz, 1H), 5.26 (t, J = 2.4 Hz, 1H), 4.93 (ddd, J = 9.3, 2.2, 1.3 Hz, 1H), 4.18 – 4.16 (m, 2H), 4.16 – 4.12 (m, 1H), 3.63 – 3.58 (m, 1H), 2.11 (s, 3H), 2.10 (s, 3H), 2.09 (d, J = 1.9 Hz, 3H), 2.08 (s, 3H).

**$^{13}$C NMR** (126 MHz, Chloroform-*d*) δ 170.75, 169.69, 169.36, 166.80, 99.55, 70.54, 68.53, 67.69, 65.14, 63.49, 21.06, 20.99, 20.90, 14.11.

**HRMS** (ESI, positive): Calculated for $C_{14}H_{19}NO_8Na$ [M+Na]$^+$ 352.1003; found: 352.1013.

**2-Acetamido-3,4,6-tri-*O*-acetyl-2-deoxy-α-D-glucopyranose 1-dibenzylphosphate (17)**

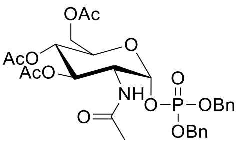

Compound **17** was synthesized using a modified literature procedure (12,13). Compound **16** (8.15 g, 24.7 mmol, 1.00 eq) was dissolved in anh. DCE (80 ml) under an argon atmosphere. Dibenzyl phosphate (8.26 g, 29.7 mmol, 1.20 eq) dissolved in anh. DCE (20 ml) was added dropwise, and the mixture was stirred for 5 days at RT. DCM (100 ml) was added and the mixture was washed with 5% $NaHCO_3$ (200 ml) and $H_2O$ (2 x 200 ml). The organic layer was dried over $MgSO_4$, filtrated, and evaporated *in vacuo*. The crude product was purified by silica gel chromatography using EtOAc as the mobile phase. Compound **17** was isolated as a white foam (6.67 g, 11.0 mmol, 44%).

**$^1$H NMR** (500 MHz, Chloroform-*d*) δ 7.42 – 7.33 (m, 10H), 5.67 (dd, J = 6.0, 3.3 Hz, 1H), 5.63 (d, J = 9.3 Hz, 1H), 5.18 – 5.02 (m, 6H), 4.41 – 4.32 (m, 1H), 4.13 (dd, J = 12.5, 4.0 Hz, 1H), 4.01 – 3.96 (m, 1H), 3.91 (dd, J = 12.5, 2.3 Hz, 1H), 2.03 (s, 3H), 2.01 (s, 3H), 2.01 (s, 3H), 1.71 (s, 3H).

**$^{13}$C NMR** (126 MHz, Chloroform-*d*) δ 171.32, 170.70, 170.29, 169.25, 135.47 (d, J = 6.3 Hz), 135.34 (d, J = 6.4 Hz), 129.12, 129.00, 128.95, 128.28, 128.23, 96.39 (d, J = 6.5 Hz), 70.26, 70.15 (d, J = 5.5 Hz), 70.10 (d, J = 5.4 Hz), 69.77, 67.41, 61.40, 51.94 (d, J = 7.7 Hz), 22.91, 20.79, 20.76, 20.69.

**$^{31}$P NMR** (202 MHz, Chloroform-*d*) δ -2.38.

**HRMS** (ESI, positive): Calculated for $C_{28}H_{34}NNaO_{12}P^+$ $[M+Na]^+$ 630.1711; found: 630.1743.

##### 2-Amino-2-deoxy-*N*-(4-methoxybenzylidene)-D-glucopyranose (18)

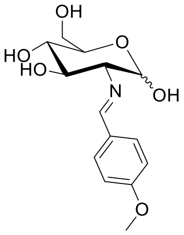

Compound **18** was synthesized using an adapted literature procedure (16). Glucosamine hydrochloride (42.0 g, 195 mmol, 1.00 eq) was dissolved in NaOH (1 M, 200 ml). The mixture was stirred vigorously and 4-anisaldehyde (39.8 g, 292 mmol, 1.50 eq) was added dropwise. The white precipitate was filtered off and the solid was washed with water (8 x 200 ml) and MeOH / EtOAc (1 / 1; 8 x 200 ml). Compound **18** was dried *in vacuo* (40.0 g, 135 mmol, 69%).

**$^1$H NMR** (500 MHz, DMSO-$d_6$) δ 8.11 (s, 1H), 7.68 (d, J = 8.7 Hz, 2H), 6.98 (d, J = 8.8 Hz, 2H), 6.53 (s, 1H), 4.93 (m, 1H), 4.82 (m, 1H), 4.69 (d, J = 7.7 Hz, 1H), 4.56 (br, 1H), 3.79 (s, 3H), 3.77 – 3.68 (m, 1H), 3.52 – 3.46 (m, 1H), 3.45 – 3.40 (m, 1H), 3.26 – 3.20 (m, 1H), 3.18 – 3.11 (m, 1H), 2.79 (dd, J = 9.3, 7.7 Hz, 1H).

**$^{13}$C NMR** (126 MHz, DMSO-$d_6$) δ 161.32, 161.08, 129.66, 129.12, 113.93, 95.65, 78.20, 76.88, 74.61, 70.39, 61.29, 55.30.

##### 2-Amino-2-deoxy-*N*-(4-methoxybenzylidene)-1,3,4,6-tetra-*O*-benzoyl-D-glucopyranose (19)

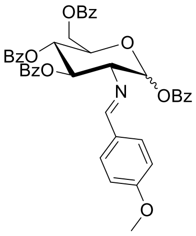

Compound **19** was synthesized using an adapted literature procedure (16). Compound **18** (20.0 g, 67.3 mmol, 1.00 eq) was dissolved in anh. pyridine (200 ml) under an argon atmosphere. The mixture was cooled to 0°C and benzoyl chloride (47.3 g, 39.0 ml, 336 mmol, 5.00 eq) was added dropwise. The reaction was stirred for 15 min, warmed to RT, and stirred for another 3 h. The reaction was cooled to 0°C and quenched by the addition of MeOH (40 ml). The solvent was removed under reduced pressure and the residue was dissolved in EtOAc (400 ml). The organic layer was washed with HCl (1 M, 2 x 400 ml), $NaHCO_3$ (sat., 400 ml) and brine (400 ml). It was dried over $MgSO_4$, filtered, and evaporated *in vacuo*. The product was recrystallized from EtOH giving white crystals (40.0 g, 56.0 mmol, 83%).

**$^1$H NMR** (500 MHz, Chloroform-*d*) δ 8.29 (s, 1H), 8.06 – 7.98 (m, 4H), 7.96 – 7.90 (m, 2H), 7.86 – 7.81 (m, 2H), 7.57 – 7.27 (m, 14H), 6.81 – 6.77 (m, 2H), 6.41 (d, J = 8.0 Hz, 1H), 6.03 (t, J = 9.5 Hz, 1H), 5.78 (t, J = 9.7 Hz, 1H), 4.66 (dd, J = 12.2, 2.9 Hz, 1H), 4.52 (dd, J = 12.2, 5.0 Hz, 1H), 4.48 – 4.43 (m, 1H), 3.93 (dd, J = 9.5, 8.0 Hz, 1H), 3.75 (s, 3H).

**$^{13}$C NMR** (126 MHz, Chloroform-*d*) δ 166.29, 165.61, 165.34, 164.54, 164.51, 162.23, 133.64, 133.50, 133.16, 133.10, 130.31, 130.09, 129.99, 129.95, 129.86, 129.68, 129.43, 129.26, 129.07, 128.53, 128.48, 128.42, 128.39, 114.00, 94.15, 73.97, 73.67, 73.22, 69.54, 63.29, 55.43.

**HRMS** (ESI, positive): Calculated for $C_{42}H_{36}NO_{10}^+$ $[M+H]^+$ 714.2334; found: 714.2346.

##### 1,3,4,6-Tetra-*O*-benzoyl-2-deoxy-2-amino-D-glucopyranose hydrochloride (20)

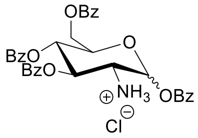

Compound **20** was synthesized using an adapted procedure (16). Compound **19** (2.66 g, 3.73 mmol, 1.00 eq.) was suspended in acetone (15 mL) and the mixture was refluxed. After the substrate was fully dissolved, HCl (5 M, 1.0 ml) was added dropwise to the refluxing solution and a white precipitate formed. The reaction mixture was stirred for 10 min and then allowed to cool to RT. The precipitate was filtered off, washed with acetone (1 x 30 mL) and diethyl ether (2 x 40 mL) and dried *in vacuo* (1.24 g, 1.96 mmol, 53%).

**$^1$H NMR** (500 MHz, DMSO-*d*$_6$) δ 8.12 – 8.07 (m, 2H), 7.94 – 7.87 (m, 4H), 7.84 – 7.80 (m, 2H), 7.73 – 7.69 (m, 1H), 7.67 – 7.54 (m, 5H), 7.52 – 7.42 (m, 6H), 6.02 (d, J = 8.3 Hz, 1H), 5.65 (t, J = 9.7 Hz, 1H), 5.50 (t, J = 9.4 Hz, 1H), 4.51 – 4.43 (m, 3H), 3.30 – 3.21 (br, 1H).

**$^{13}$C NMR** (126 MHz, DMSO-*d*$_6$) δ 165.47, 165.32, 164.84, 164.46, 133.98, 133.65, 133.42, 133.38, 129.71, 129.34, 129.28, 129.23, 129.18, 129.17, 128.83, 128.78, 128.78, 128.69, 128.60, 95.85, 75.43, 71.53, 69.57, 62.63, 55.55.

**HRMS** (ESI, positive): Calculated for $C_{34}H_{30}NO_9^+$ $[M+H]^+$ 596.1915; found: 596.1917.

##### 2-(2-Chloroacetamido)-1,3,4,6-tetra-*O*-benzoyl-2-deoxy-D-glucopyranose (21)

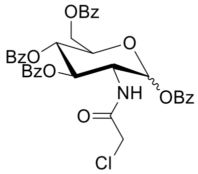

Compound **20** (3.20 g, 5.06 mmol, 1.00 eq) was dissolved in anh. DCM (50 ml) under an argon atmosphere. $Et_3N$ (5.08 g, 7.00 ml, 40.2 mmol, 7.95 eq) was added and the solution was stirred at RT for 10 min. Chloroacetic anhydride (2.60 g, 15.21 mmol, 3.00 eq) was added portionwise and the reaction was stirred at RT for 4 h. It was diluted with DCM (50 ml) and the organic layer was washed with water (100 ml) and brine (100 ml). The organic layer was dried over $MgSO_4$, filtered and the solvent was removed under reduced pressure. The residue was purified by silica gel chromatography using EtOAc / cyclohexane (2 / 3) yielding compound **21** as a white foam (3.01 g, 4.48 mmol, 88%).

**$^1$H NMR** (500 MHz, Chloroform-*d*) δ 8.09 (d, J = 7.2 Hz, 2H), 8.03 (d, J = 7.2 Hz, 2H), 7.96 (d, J = 7.2 Hz, 2H), 7.89 (d, J = 7.2 Hz, 2H), 7.64 – 7.30 (m, 12H), 6.92 (d, J = 9.3 Hz, 1H), 6.19 (d, J = 8.6 Hz, 1H), 5.85 – 5.76 (m, 2H), 4.79 – 4.71 (m, 1H), 4.64 (dd, J = 12.4, 2.9 Hz, 1H), 4.49 (dd, J = 12.3, 4.6 Hz, 1H), 4.37 – 4.28 (m, 1H), 3.87 (d, J = 15.3 Hz, 1H), 3.83 (d, J = 15.2 Hz,1H).

**$^{13}$C NMR** (126 MHz, Chloroform-*d*) δ 166.86, 166.76, 166.26, 165.18, 165.07, 134.08, 133.89, 133.73, 133.28, 130.47, 130.15, 129.98, 129.92, 129.65, 128.77, 128.69, 128.62, 128.56, 128.51, 93.40, 73.38, 72.63, 68.83, 62.79, 53.94, 42.41.

**HRMS** (ESI, positive): Calculated for $C_{36}H_{30}ClNNaO_{10}^+$ $[M+Na]^+$ 694.1450; found: 694.1456.

**2-(2-Chloroacetamido)-3,4,6-tri-*O*-benzoyl-2-deoxy-D-glucopyranose (22)**

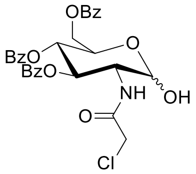

For the synthesis of compound **22** a literature procedure was adapted (17). Compound **21** (7.60 g, 11.31 mmol, 1.00 eq) was dissolved in anhydrous DCM (40 ml) under an argon atmosphere. HBr in acetic acid (30%, 4 ml) was added and the solution was stirred for 3 h at RT. The organic layer was washed with sat. $NaHCO_3$ (2 x 40 ml) and $H_2O$ (40 ml). It was dried over $MgSO_4$, filtered and the solvent was removed under reduced pressure. The resulting foam was dissolved in acetone (20 ml) and $H_2O$ (1 ml). $Ag_2CO_3$ (3.12 g, 11.31 mmol, 1.00 eq) was added and it was stirred for 2 h. The solution was filtered over a plug of Celite and the solvent was removed under reduced pressure.

The residue was purified by RP-column chromatography using two C18 23 g flash columns (Telos) and water / ACN as mobile phase (gradient: 30% ACN for 10 min, to 90% ACN in 50 min, flow rate: 15 ml/min). Fractions containing the product were lyophilized giving a yellowish powder. The product was a mixture of the *N*-chloroacetyl and *N*-bromoacetyl (Chemical shifts depicted with “Br”) product (2.77 g, 4.88 mmol, 43%).

**$^1$H NMR** (500 MHz, Chloroform-*d*) δ 8.08 – 8.01 (m, 2H), 7.95 – 7.86 (m, 4H), 7.59 – 7.32 (m, 9H), 7.05 (d, J = 9.2 Hz, 0.7H), 6.89 (d, J = 9.1 Hz, 0.3H)$^{Br}$, 5.89 – 5.80 (m, 1H), 5.76 – 5.68 (m, 1H), 5.47 – 5.42 (br, 1H), 4.68 – 4.58 (m, 2H), 4.57 – 4.48 (m, 1H), 4.41 (m, 1H), 3.96 (d, J = 15.3 Hz, 0.7H), 3.85 (d, J = 15.2 Hz, 0.7H), 3.76 (d, J = 13.4 Hz, 0.3H)$^{Br}$, 3.68 (d, J = 13.4 Hz, 0.3H)$^{Br}$, 3.44 (br, 1H).

**$^{13}$C NMR** (126 MHz, Chloroform-*d*) δ 166.99, 166.50, 166.45, 166.15$^{Br}$, 165.33, 133.60, 133.58, 133.34, 130.06, 130.05, 129.96, 129.74, 129.04, 128.95, 128.59, 128.57, 91.76, 71.15, 71.11$^{Br}$, 69.30, 68.35, 62.89, 53.48$^{Br}$, 53.24, 42.41, 28.50$^{Br}$.

**HRMS** (ESI, positive): Calculated for $C_{29}H_{26}ClNNaO_9^+$ $[M+Na]^+$ 590.1188; found: 590.1203.

**HRMS** (ESI, positive): Calculated for $C_{36}H_{30}BrNNaO_{10}^+$ $[M+Na]^+$ 634.0683; found: 634.0695.

**2-(2-Chloroacetamido)-3,4,6-tri-*O*-benzoyl-2-deoxy-α-D-glucopyranose 1-dicyanoethyl-phosphate (23)**

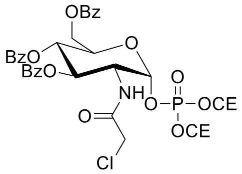

Compound **22** (200 mg, 352 µmol, 1.00 eq) was suspended in anh. ACN (5 ml). A tetrazole solution in ACN (0.45 mM, 1.57 ml, 704 µmol, 2.00 eq) and bis(2-cyanoethyl)-*N*,*N*-diisopropyl phosphoramidite (191 mg, 186 µl, 704 µmol, 2.00 eq) were added dropwise. The mixture was heated to 40°C and the solution went clear. It was stirred until full conversion of the substrate (1-2 h, TLC EtOAc / cyclohexane 1 /1). Diethyl ether (100 ml) was added, and the organic layer was washed with sat. $NaHCO_3$ (100 ml). The aqueous layer was extracted with diethyl ether (2 x 100 ml), combined organic layers were dried over $MgSO_4$ and filtered. The solvent was removed under reduced pressure and the residue was dissolved in anh. THF (16 ml) under an argon atmosphere.

For the oxidation of the intermediate a modified literature procedure was used (18). The mixture was cooled to -78°C and $H_2O_2$ (30%, 3.20 ml) was added. The mixture was stirred for 1.5 h and slowly warmed to RT. Diethyl ether was added (200 ml) and it was washed with $Na_2S_2O_3$ (1 M, 100 ml), $NaHCO_3$ (sat., 100 ml) and $H_2O$ (100 ml). The organic phase was dried over $MgSO_4$, filtered and the solvent was removed under reduced pressure. The product was purified by silica gel chromatography (EtOAc / cyclohexane; first 1 / 1, then 4 / 1). Compound **23** was isolated as a white foam (159 mg, 210 µmol, 60%).

**$^1$H NMR** (500 MHz, Chloroform-*d*) δ 8.05 – 8.01 (m, 2H), 7.92 (m, 4H), 7.60 – 7.55 (m, 1H), 7.53 – 7.47 (m, 2H), 7.46 – 7.41 (m, 2H), 7.39 – 7.32 (m, 4H), 7.21 (d, J = 8.2 Hz, 0.7H), 7.13 (d, J = 8.3 Hz, 0.3H)$^{Br}$, 5.97 (dd, J = 5.8, 3.3 Hz, 0.7H), 5.95 (dd, J = 5.5, 3.3 Hz, 0.3H)$^{Br}$, 5.82 – 5.73 (m, 2H), 4.67 (dd, J = 12.2, 3.0 Hz, 1H), 4.65 – 4.57 (m, 2H), 4.52 – 4.46 (m, 1H), 4.44 – 4.27 (m, 4H), 4.04 (d, J = 15.1 Hz, 0.7H), 3.90 (d, J = 15.1 Hz, 0.7H), 3.83 (d, J = 13.1 Hz, 0.3H)$^{Br}$, 3.71 (d, J = 13.1 Hz, 0.3H)$^{Br}$, 2.81 – 2.77 (m, 2H), 2.77 – 2.72 (m, 2H).

**$^{13}$C NMR** (126 MHz, Chloroform-*d*) δ 166.99, 166.93$^{Br}$, 166.76, 166.61$^{Br}$, 165.97, 165.07, 133.81, 133.76$^{Br}$, 133.68, 133.66$^{Br}$, 133.40, 129.96, 129.86, 129.73, 129.37, 128.54, 128.50, 128.47, 128.33, 128.27, 116.50$^{Br}$, 116.37$^{Br}$, 116.34, 116.27, 96.40 (d, J = 6.1 Hz), 70.46, 69.94, 69.88$^{Br}$, 68.37$^{Br}$, 68.27, 62.99 (d, J = 5.0 Hz), 62.82 (d, J = 4.9 Hz), 62.33, 53.14 (d, J = 8.5 Hz), 42.25, 28.17$^{Br}$, 19.75 (d, J = 7.1 Hz), 19.68 (d, J = 7.2 Hz).

**$^{31}$P NMR** (202 MHz, Chloroform-*d*) δ -3.36, -3.38$^{Br}$.

**HRMS** (ESI, positive): Calculated for $C_{35}H_{33}ClN_3NaO_{12}P^+$ $[M+Na]^+$ 776.1383; found: 776.1387.

**HRMS** (ESI, positive): Calculated for $C_{35}H_{33}BrN_3NaO_{12}P^+$ $[M+Na]^+$ 820.0877; found: 820.0869.

**2-(Chloroacetylamino)-3,4,6-tri-*O*-benzoyl-2-deoxy-α-D-glucopyranose 1-phosphate (24)**

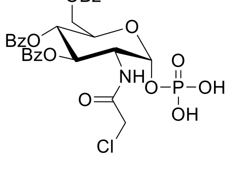

Compound **24** was synthesized using a modified literature procedure (14). Compound **23** (44.4 mg, 58.9 µmol, 1.00 eq) was dissolved in anh. ACN (10 ml) under an argon atmosphere. The mixture was cooled to 0°C and DBU (27 mg, 26 µl, 176 µmol, 3 eq) was added. BSA (270 mg, 324 µl, 1.32 mmol, 22.5 eq) was added dropwise, and the mixture was stirred at RT for

1 h. It was cooled to 0°C and a mixture of TFA in MeOH (1 / 3; 1 ml) was added dropwise to quench the reaction. It was stirred for 10 min at RT before buffer A (100 mM, 10 ml) and Celite was added. The solvent was removed under reduced pressure and the crude product was purified using a 12 g PF-30C18AQ (interchim) column and 10 mM buffer A / B as mobile phase (gradient: 5% buffer B for 10 min, to 80% buffer B in 50 min, flow rate: 15 ml/min). Compound **24** was isolated as mono(triethylammonium) salt and was a white foam (17.1 mg, 22.8 µmol, 39%).

**$^1$H NMR** (500 MHz, Methanol-$d_4$) δ 8.06 – 8.00 (m, 2H), 7.90 – 7.83 (m, 4H), 7.60 – 7.56 (m, 1H), 7.55 – 7.49 (m, 2H), 7.46 – 7.41 (m, 2H), 7.40 – 7.34 (m, 4H), 5.90 – 5.82 (m, 1H), 5.73 – 5.65 (m, 2H), 4.73 – 4.67 (m, 1H), 4.62 – 4.56 (m, 2H), 4.50 – 4.43 (m, 1H), 4.00 (d, J = 13.1 Hz, 1H), 3.93 (d, J = 13.1 Hz, 1H), 3.21 (q, J = 7.3 Hz, 3H)#, 1.31 (t, J = 7.2 Hz, 7H)#.

**$^{13}$C NMR** (126 MHz, Methanol-$d_4$) δ 169.79, 167.63, 167.54, 166.76, 134.59, 134.38, 134.26, 131.07, 130.78, 130.75, 130.72, 130.60, 130.41, 129.55, 129.51, 129.43, 95.20 (d, J = 5.9 Hz), 73.05, 71.15, 69.91, 64.02, 54.00 (d, J = 8.1 Hz), 47.91#, 42.98, 9.21#.

**$^{31}$P NMR** (202 MHz, Methanol-$d_4$) δ -1.36.

**HRMS** (ESI, negative): Calculated for $C_{29}H_{26}ClNO_{12}P^-$ $[M-H]^-$ 646.0887; found: 646.0920.

**2'-*O*,3'-*O*,*N*$^2$-triisobutyrylguanosine (25)**

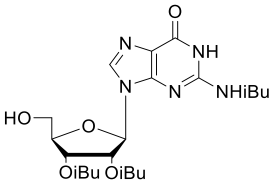

2'-*O*,3'-*O*,*N*$^2$-triisobutyrylguanosine **25** was synthesized using a modified literature procedure (19). 5'-O-DMT-*N*$^2$-isobutyrylguanosine (3.00 g, 4.58 mmol, 1.00 eq) was dissolved in anh. pyridine (40 ml) and cooled to 0°C. Isobutyric anhydride (7.21 g, 7.59 ml, 7.24 g, 45.8 mmol, 10 eq) was added dropwise and the mixture was stirred 30 min at 0°C. It was allowed to warm to RT and stirred overnight. The mixture was cooled to 0°C, MeOH (40 ml) was added, and the mixture was warmed to RT and stirred for another 30 min. The solvents were evaporated under reduced pressure, the residue was dissolved in DCM (100 ml) and washed with $H_2O$ (100 ml), sat. $NaHCO_3$ (100 ml) and brine (100 ml). The organic layer was dried over $MgSO_4$, filtered, and evaporated under reduced pressure.

The resulting foam was dissolved in DCM (50 ml) and cooled to 0°C. DCM (50 ml) containing 6% TFA and 6% MeOH was added dropwise at 0°C while stirring vigorously. It was stirred at RT until the intermediate product was completely converted (TLC, 5% MeOH in DCM). The mixture was cooled again to 0°C and sat. $NaHCO_3$ (100 ml) was added dropwise while stirring vigorously. After the orange color disappeared, phases were separated, and the organic layer was washed with brine (100 ml). The organic layer was dried over $MgSO_4$, filtered, and removed under reduced pressure. The residue was purified by reverse phase chromatography using two C18 23 g flash columns (Telos) (gradient: 10 % ACN for 5 min, to 60% ACN in 60 min, flow rate: 15 ml/min). Compound **25** was obtained as a white foam (2.08 g, 4.21 mmol, 91%).

**$^1$H NMR** (500 MHz, Chloroform-*d*) δ 12.17 (s, 1H), 9.15 (s, 1H), 7.73 (s, 1H), 5.94 (dd, J = 6.8, 5.4 Hz, 1H), 5.89 (d, J = 6.8 Hz, 1H), 5.68 (dd, J = 5.4, 2.8 Hz, 1H), 4.94 (d, J = 9.7 Hz, 1H), 4.27 – 4.24 (m, 1H), 3.98 – 3.94 (m, 1H), 3.84 – 3.78 (m, 1H), 2.73 (hept, J = 6.9 Hz, 1H), 2.61 (hept, J = 7.0 Hz, 1H), 2.50 (hept, J = 7.0 Hz, 1H), 1.26 (d, J = 6.9 Hz, 3H), 1.25 (d, J = 6.9 Hz, 3H), 1.20 (d, J = 7.0 Hz, 6H), 1.11 (d, J = 7.0 Hz, 3H), 1.08 (d, J = 7.0 Hz, 3H).

**$^{13}$C NMR** (126 MHz, Chloroform-*d*) δ 179.19, 176.07, 175.50, 155.29, 148.13, 147.32, 138.98, 122.84, 87.99, 84.94, 72.39, 71.46, 62.21, 36.50, 34.00, 33.71, 19.04, 19.02, 18.96, 18.89, 18.80.

**HRMS** (ESI, positive): Calculated for $C_{22}H_{31}N_5NaO_8^+$ $[M+Na]^+$ 516.2065; found: 516.2073.

**Protected UpG building block (26)**

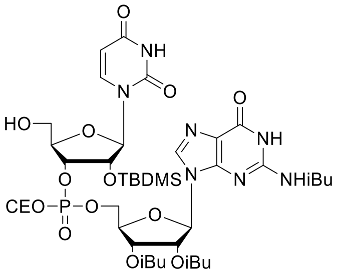

The protected UpG building block **26** was synthesized using a modified literature procedure (20). Compound **25** (0.53 g, 1.28 mmol, 1.10 eq) and DMT-2′*O*-TBDMS-rU-phosphoramidite (1.00 g, 1.16 mmol, 1.00 eq) were placed in a schlenk flask under an argon atmosphere. The substrates were dissolved in anh. ACN (10 ml) and a solution of tetrazole in anh. ACN (12.9 ml, 0.45 M, 5.81 mmol, 5.00 eq) was added. The reaction was stirred at RT until the phosphoramidite was fully converted according to TLC (MeOH / DCM; 1 / 9). An $I_2$ solution (0.1 M) in THF / pyridine / $H_2O$ (2 / 2 / 1) was added dropwise until the brownish color persisted. Excess iodide was quenched by the addition of a $Na_2S_2O_3$ solution (1 M, 100 ml), and the aqueous layer was extracted with DCM (3 x 100 ml). Combined organic layers were dried over $MgSO_4$, filtered and the solvent was removed under reduced pressure.

The resulting foam was dissolved in DCM (50 ml), cooled to 0°C and a solution of 6% MeOH and 6% TFA in DCM (50 ml) was added dropwise. The reaction was stirred at RT until the intermediate product was fully converted according to TLC (MeOH / DCM; 1 / 9). The reaction was cooled to 0°C and sat. $NaHCO_3$ (100 ml) was added dropwise under vigorous stirring. Phases were separated and the organic layer was washed with brine (100 ml). The organic layer was dried over $MgSO_4$, filtered and the solvent was removed under reduced pressure. The residue was purified by reverse phase chromatography using two C18 23 g flash columns (Telos) and $H_2O$ / ACN as mobile phase (gradient: 5 % ACN for 5 min, to 90% ACN in 85 min, flow rate: 15 ml/min). Compound **26** was isolated as a white foam (885.60 mg, 0.91 mmol, 78%, 1 : 1 mixture of diastereomers).

**$^1$H NMR** (500 MHz, Methanol-$d_4$) δ 8.24 (s, 0.5H), 8.23 (s, 0.5H), 7.99 (d, J = 8.2 Hz, 0.5H), 7.98 (d, J = 8.2 Hz, 0.5H), 6.22 – 6.17 (m, 1H), 6.03 – 5.98 (m, 1H), 5.95 (t, J = 6.4 Hz, 1H), 5.75 (d, J = 8.1 Hz, 0.5H), 5.72 (d, J = 8.1 Hz, 0.5H), 5.67 – 5.64 (m, 1H), 4.87 – 4.80 (m, 1H)*, 4.59 – 4.46 (m, 4H), 4.39 – 4.27 (m, 3H), 3.75 – 3.63 (m, 2H), 2.94 – 2.86 (m, 2H), 2.81 – 2.75 (m, 1H), 2.72 – 2.64 (m, 1H), 2.62 – 2.52 (m, 1H), 1.25 – 1.21 (m, 12H), 1.15 – 1.08 (m, 6H), 0.88 (s, 4.5H), 0.82 (s, 4.5H), 0.11 – 0.01 (m, 6H).

**$^{13}$C NMR** (126 MHz, Methanol-$d_4$) δ 179.72, 179.71, 175.11, 175.07, 174.83, 163.67, 163.64, 154.99, 154.98, 150.21, 150.20, 148.07, 148.00, 147.93, 140.06, 139.99, 138.27*, 119.56, 116.35, 116.22, 101.31, 101.23, 86.85, 86.83, 86.23, 86.16, 82.95 (d, J = 9.3 Hz), 82.93 (d, J = 9.4 Hz), 80.56 (d, J = 8.2 Hz), 80.38 (d, J = 8.2 Hz), 77.46 (d, J = 5.4 Hz), 77.38 (d, J = 5.8 Hz), 73.50 (d, J = 5.4 Hz), 73.49 (d, J = 5.3 Hz), 71.68, 71.63, 69.70, 69.44, 66.85 (d, J = 6.2 Hz), 66.56 (d, J = 6.1 Hz), 62.51 (d, J = 5.4 Hz), 62.44 (d, J = 5.4 Hz), 59.84, 59.80, 34.82, 32.89, 32.70, 24.06, 24.03, 18.00 (d, J = 7.9 Hz), 17.96 (d, J = 7.9 Hz), 17.34, 17.32, 17.18, 17.16, 17.15, 17.04, 17.03, 17.02, 17.00, 16.91, 16.85, 16.77, -6.75, -6.85, -7.11, -7.12.

**$^{31}$P NMR** (202 MHz, Methanol-$d_4$) δ -3.62, -3.74.

**HRMS** (ESI, positive): Calculated for $C_{40}H_{59}N_8NaO_{16}PSi^+$ $[M+Na]^+$ 989.3448; found: 989.3495.

### 4 Biochemical procedures

#### 4.1 RNA preparation

##### 4.1.1 Template preparation

Short templates were purchased as complementary ssDNA and annealed by heating a mixture of both (50 µM) at 90°C for 5 min followed by slow cooling to RT. Longer templates were prepared by extending the overlap of two ssDNA strands by Taq PCR.

##### 4.1.2 PCR

To fill up two partially complementary ssDNA strands, Taq PCR was conducted using ssDNA (2.5 µM), GenTherm buffer without $MgCl_2$ (1x, Rapidozym, Berlin, Germany), $MgCl_2$ (3 mM), dNTP mix (0.5 mM), DMSO (5%) and Taq polymerase (0.05 mg/ml, lab prepared stock). The template was purified using a QIAquick PCR purification kit (Qiagen, Venlo, Netherlands).

##### 4.1.3 RNA 5' pyrophosphohydrolase reaction

To create 5' monophosphate RNA for ligation, 3 µg 68mer RNA was incubated with NEBuffer 2 and RppH (0.5 U/µl) (NEB, Ipswich, MA, USA) for 30 min at 37°C in a final volume of 20 µl. RNA was purified using NEB Monarch nucleic acid purification kit.

##### 4.1.4 Ligation of capped-11mer RNA to monophosphate-68mer RNA

UDP-sugar-capped-11mer RNA (5 pmol) was mixed with 5' monophosphate-68mer RNA (2 pmol) and a complementary DNA splint (4 pmol) in a volume of 16 µl. The mixture was heated to 85°C for 5 min and slowly cooled to 30°C. T4 DNA Ligase buffer (2 µl, 10x) and T4 DNA Ligase (2 µl, 30 Weiss U/µl) (Thermo Fischer Scientific, Waltham, MA, USA) was added and the mixture was incubated for 4 h at 30°C. For negative controls, Milli-Q water was added instead of the T4 ligase. The reaction was terminated by the addition of denaturing loading dye (2x TBE, 12 mM EDTA, BPB, XC in formamide) and analyzed by 10% PAGE.

##### 4.1.5 DNAzyme cleavage

For DNAzyme cleavage reactions, RNA (5 pmol) and DNAzyme (10 pmol, PAGE purified) were mixed in a total volume of 16 µl. The mixture was heated at 85°C for 2 min before slowly cooling to 37°C. 5x DNAzyme reaction buffer (4 µl, 1 M NaCl, 50 mM $MgCl_2$, 250 mM HEPES (pH 8)). was added and it was incubated for 1 h at 37°C. It was heated again at 85°C for 2 min, slowly cooled to 37°C and incubated for another 1 h at 37°C. Denaturing loading dye (2x TBE, 12 mM EDTA, BPB, XC in formamide) was added and reactions were analyzed by 20% denaturing PAGE. Capping efficiencies have been calculated using band intensities of capped ($I_{capped}$) and uncapped ($I_{uncapped}$) fragments according to the following equation:

$$Capping\ efficiency = \frac{I_{capped}}{I_{capped} + I_{uncapped}} * 100\%$$

##### 4.1.6 Streptavidin shift assay

Biotinylated RNA and streptavidin (0.25 mg/ml) were mixed in a volume of 6 µl and incubated for 20 min at 37°C. Negative controls did not contain streptavidin. Afterwards, 6 µl strepshift loading dye (40 mM NaCl, 2x TBE, XC, BPB in 50% glycerol) was added and samples were analyzed using 10% native PAGE. The gel was developed in a cold room at 4°C. Capping efficiencies have been calculated using band intensities of shifted ($I_{shifted}$) and non-shifted ($I_{non-shifted}$) RNA according to the following equation:

$$Capping\ efficiency = \frac{I_{shifted}}{I_{shifted} + I_{non-shifted}} * 100\%$$

#### 4.2 Protein preparation

##### 4.2.1 Cloning of MurC

The MurC gene was amplified from *E. coli* K12 genomic DNA using primers specified in **Table S1** and Phusion HF DNA polymerase (Thermo Fischer Scientific, Waltham, MA, USA). The amplified gene was digested with NcoI and XhoI (Thermo Fischer Scientific, Waltham, MA, USA) and ligated to Pet-28c(+), digested with the same restriction enzymes, by T4 DNA ligase

(Thermo Fischer Scientific, Waltham, MA, USA). The ligation mixture was transformed into chemically competent *E. coli* DH5α and isolated vectors containing the right insert were transformed into chemically competent BL21 Star (DE3).

###### 4.2.2 Growth of *E. coli* BL21(DE3)

Proteins were expressed in *E. coli* BL21 Star (DE3) containing Pet-28c(+) vectors with the respective genes shown in **Table S3**. The overnight culture of *E. coli* BL21(DE3) was grown in lysogeny broth (LB) medium containing 1 mM kanamycin at 37°C, 650 rpm. 1 L LB medium containing 1 mM kanamycin was inoculated with an overnight culture to an $OD_{600}$ of 0.1. Bacteria were grown until a final $OD_{600}$ of 0.6-0.8 at 37°C, 650 rpm and protein expression was induced by the addition of 1 mM IPTG. It was incubated at 37°C 650 rpm for an additional 3-4 h. For harvesting, cells were transferred to 1 L centrifugation tubes and centrifuged at 4°C, 4000 rpm for 15 min. The supernatant was discarded, the cell pellet was resuspended in 10 ml PBS and transferred to a 50 ml Falcon tube. It was centrifuged at 4°C, 4000 rpm for 15 min, the supernatant was discarded, and the pellet was stored at -80° C overnight.

###### 4.2.3 Lysis

The cell pellet was resuspended in 10 ml protein buffer A (50 mM Tris-HCl (pH 7.5), 300 mM NaCl, 5 mM β-mercaptoethanol, 1 mM imidazole, 5 mM $MgSO_4$, 5% (v/v) glycerol, complete EDTA free protease inhibitor cocktail (Roche, Basel, Switzerland)) and lysed by sonication using a Sonopuls HD 2070 sonicator (Bandilin electronic, Berlin, Germany) while cooling on ice. Cell debris was pelleted through centrifugation at 4°C, 37500 g for 30 min. The supernatant was transferred to a falcon tube and filtered using a 0.45 µm syringe filter.

###### 4.2.4 Immobilized metal affinity chromatography (IMAC)

Proteins were purified from cell lysate using a 1 ml HisTrap HP column (Cytiva, Marlborough, MA, USA). The column was equilibrated and samples were loaded using 100% protein buffer A. Afterwards, the column was washed with 5% protein buffer B (50 mM Tris-HCl (pH 7.5), 300 mM NaCl, 5 mM β-mercaptoethanol, 300 mM imidazole, 5 mM $MgSO_4$, 5% (v/v) glycerol, complete EDTA free protease inhibitor cocktail) followed by elution of the protein of interest by gradual increasing protein buffer B content to 100%.

Fractions containing eluted protein were identified through absorbance measured at 280 nm and analyzed using 10% SDS-PAGE. Pure fractions containing the protein of interest were combined and concentrated using 10 kDa MW cut-off filters. If samples were further purified by size exclusion chromatography, the protein solution was concentrated to approximately 800 µl and further purified as described in 4.2.5. If proteins were directly used, the buffer was exchanged for protein storage buffer (50 mM Tris-HCl (pH 7.5), 300 mM NaCl, 5 mM β-mercaptoethanol, 5 mM $MgSO_4$, 5% (v/v) glycerol, complete EDTA free protease inhibitor cocktail) using 10 kDa MW cut-off filters. The protein was diluted with protein storage buffer to a final concentration of 2 mg/ml and an equal volume of glycerol was added giving a final concentration of 1 mg/ml protein.

###### 4.2.5 Size exclusion chromatography (SEC)

The enzyme hNudt5 was additionally purified by SEC using a HiPrep 26/60 Sephacryl S-300 HR column (Cytiva, Marlborough, MA, USA) and isocratic elution with protein storage buffer. Fractions containing protein were analyzed using 10% SDS-PAGE. Pure fractions were combined and concentrated to a final concentration of 2 mg/ml using 10 kDa MW cut-off filters. The protein solution was mixed with an equal volume of glycerol.

##### 4.3 Enzymatic reactions

Reaction mixtures containing UDP-sugars, capped-dinucleotides and capped-11mer RNA were separated by RP-HPLC using a VDSpher PUR 100 C18-SE 250 x 4.6 mm, 5 µm column (for mono and dinucleotides: gradient: 1% buffer B for 10 min, to 10% buffer B in 18 min, flow rate: 1 ml/min; for capped 11mers: gradient: 1.6% buffer B for 5 min, to 14% buffer B in 35 min; flow rate: 1 ml/min).

##### 4.3.1 Comparison of mono- and dinucleotide substrates of MurA

Conversion of MurA substrates was conducted in triplicates and analyzed by quantifying orthophosphate generated during the reaction using BIOMOL GREEN phosphate quantification kit (Enzo Life Sciences, Farmingdale, NY, USA). Reactions were conducted in a final volume of 1 ml. Substrates (0.05 mM), MurA (50 nM) and Tris HCl (50 mM, pH 7.8) were mixed in Milli-Q water and pre-incubated for 10 min at 37°C before reactions were started by the addition of PEP to a final concentration of 1 mM. In negative controls, no sugar-mononucleotide or sugar-dinucleotide was present. At different time points, 100 µl aliquots were drawn from the reactions and quenched by the addition of 1 ml BIOMOL GREEN reagent. Mixtures were incubated for 20-30 min at RT and absorbance was measured at 650 nm using a Cary 50 UV-Vis spectrophotometer (Varian Medical Systems, Palo Alto, CA, USA). A calibration curve was prepared according to the manufacturer's protocol and conversion was calculated according to quantified orthophosphate.

##### 4.3.2 Comparison of mono- and dinucleotide substrates of MurB

Conversion of MurB substrates was conducted in triplicates and analyzed by quantifying NADPH consumption during the reaction by measuring the decrease in absorbance at 340 nm. Reactions were conducted in a final volume of 1 ml. Substrates (0.1 mM), KCl (1.5 mM), DTT (0.5 mM), Tris HCl (50 mM, pH 7.8) and MurB (25 nM) were mixed in Milli-Q water and pre-incubated for 10 min at 25°C before reactions were started by NADPH addition to a final concentration of 0.15 mM. Negative controls were conducted in the absence of substrate and blank reactions in the absence of enzyme. Reactions were carried out in a quartz cuvette at 25°C and absorbance was measured throughout the reaction using a Cary 100 Bio UV/VIS-spectrometer (Varian Medical Systems, Palo Alto, CA, USA).

Afterward, the enzyme was separated by filtration through a 10 kDa MW cut-off filter, the filtrate was lyophilized, the residue was dissolved in 100 µl buffer A (100 mM) and substrate conversion was confirmed by RP-HPLC and ESI-MS.

##### 4.3.3 Comparison of mono- and dinucleotide substrates of MurC

Conversion of MurC substrates was conducted in triplicates and analyzed by RP-HPLC measurement. Reactions were conducted in a final volume of 1 ml. Substrate (0.5 mM), DTT (1 mM), $(NH_4)_2SO_4$ (25 mM) $MgCl_2$ (20 mM), Tris-HCl (100 mM, pH 8), MurC (1.6 µM) and alanine (1 mM) were mixed in Milli-Q water and pre-incubated for 10 min at 37°C before reactions were started by ATP addition to a final concentration of 1.5 mM.

At different time points, 100 µl aliquots were drawn from the reaction, quenched by the addition of EDTA (8 µl, 500 mM, pH 8) and diluted to 200 µl with buffer A (200 mM). Mixtures were filtered using a 10 kDa MW cut-off filter and filtrates were analyzed by RP-HPLC.

##### 4.3.4 Comparison of mono- and dinucleotide substrates of hNudt5

Reactions were conducted in triplicates. Substrates (0.625 mM), DTT (1 mM), $MgCl_2$ (10 mM), Tris HCl (50 mM, pH 8) and hNudt5 (4 µM) in a total volume of 200 µl were incubated at 37°C. At different time points, 20 µl aliquots were drawn from the reaction and diluted to 200 µl containing EDTA (1 mM), and buffer A (100 mM). The enzyme was removed by filtration through 3 kDa MW cut-off filters and the filtrate was analyzed by RP-HPLC.

##### 4.3.5 PABLO analysis

Ligation was conducted using RNA (10 pmol; hNudt5 treated UDP-sugar-11mer RNA, negative controls or 5' monophosphate-11mer RNA positive controls), 5' OH-39mer DNA (10 pmol, page purified) and a complementary DNA splint (10 pmol, page purified). The three components were mixed in a total volume of 16 µl, heated to 85°C for 5 min and slowly cooled to 30°C. T4 DNA Ligase buffer (2 µl, 10x) and T4 DNA Ligase (2 µl, 30 Weiss U/µl) (Thermo Fischer Scientific, Waltham, MA, USA) were added and the mixture was incubated for 4 h at 30°C. For negative controls, Milli-Q water was added instead of the Ligase. The reaction was terminated through the addition of denaturing loading dye (2x TBE, 12 mM EDTA, BPB, XC, Formamide) and analyzed using 10% PAGE.

##### 4.4 Azide-alkyne cycloaddition (AAC)

###### 4.4.1 Copper-catalyzed azide-alkyne cycloaddition (CuAAC)

A 2x copper click master mix was prepared containing $CuSO_4$ (2 mM), THPTA (1 mM) and sodium ascorbate (4 mM). RNA was mixed with AF647-alkyne or -azide (500 µM) and phosphate buffer (100 mM, pH 7) in a 10 µl volume. An equal volume of the copper click master mix was added and reactions were incubated for 30 min at 25°C. Reactions were diluted to 200 µl containing EDTA (10 mM) and extracted three times with aqueous phenol as well as twice with diethyl ether. RNA was precipitated with sodium acetate and ethanol.

###### 4.4.2 Strain-promoted azide-alkyne cycloaddition (SPAAC)

RNA was incubated with DBCO-AF647 (50 µM) or DBCO-Biotin (50 µM) and phosphate buffer (50 mM, pH 7) in a total volume of 20 µl for 2 h at 50°C. The reaction was diluted to 200 µl and extracted three times with aqueous phenol and twice with diethyl ether. RNA was precipitated with sodium acetate and ethanol.

#### 5 NMR spectra

##### Compound 1 $^{1}H$ NMR

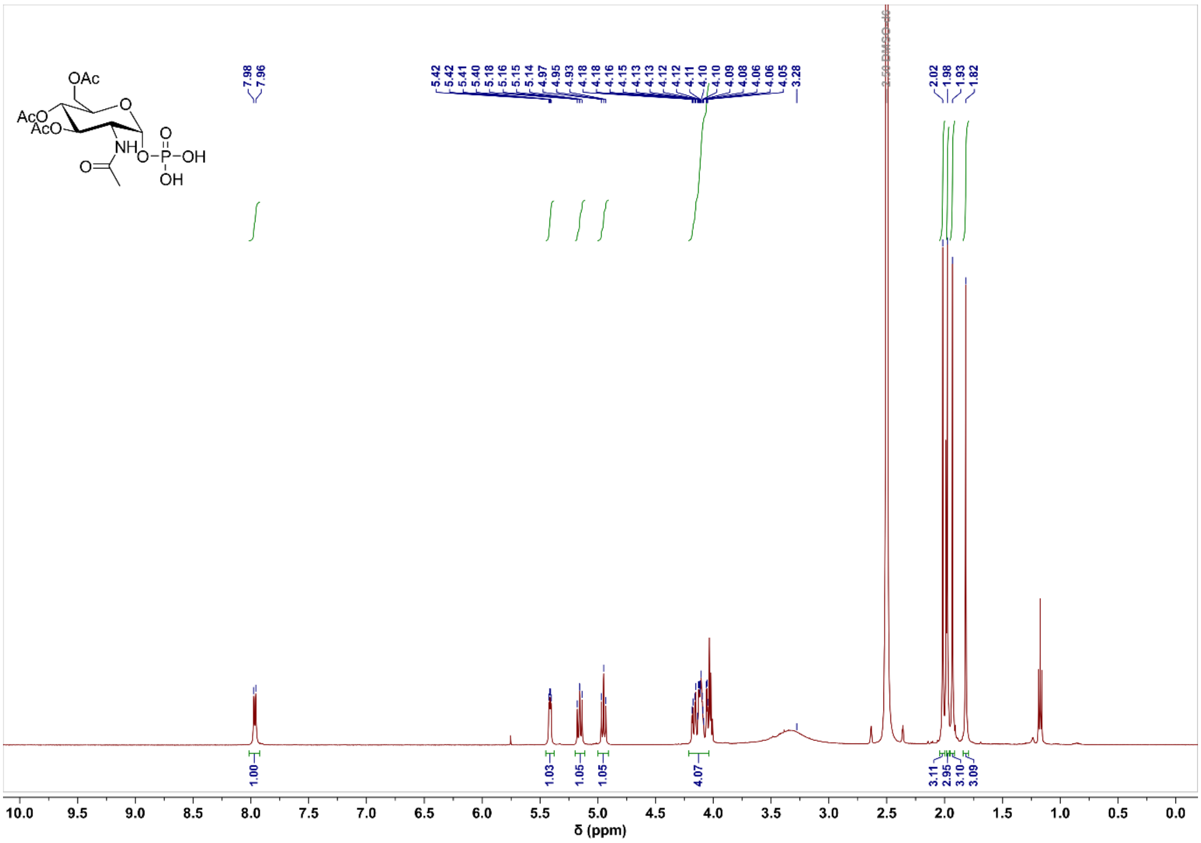

##### Compound 1 $^{13}C$ NMR

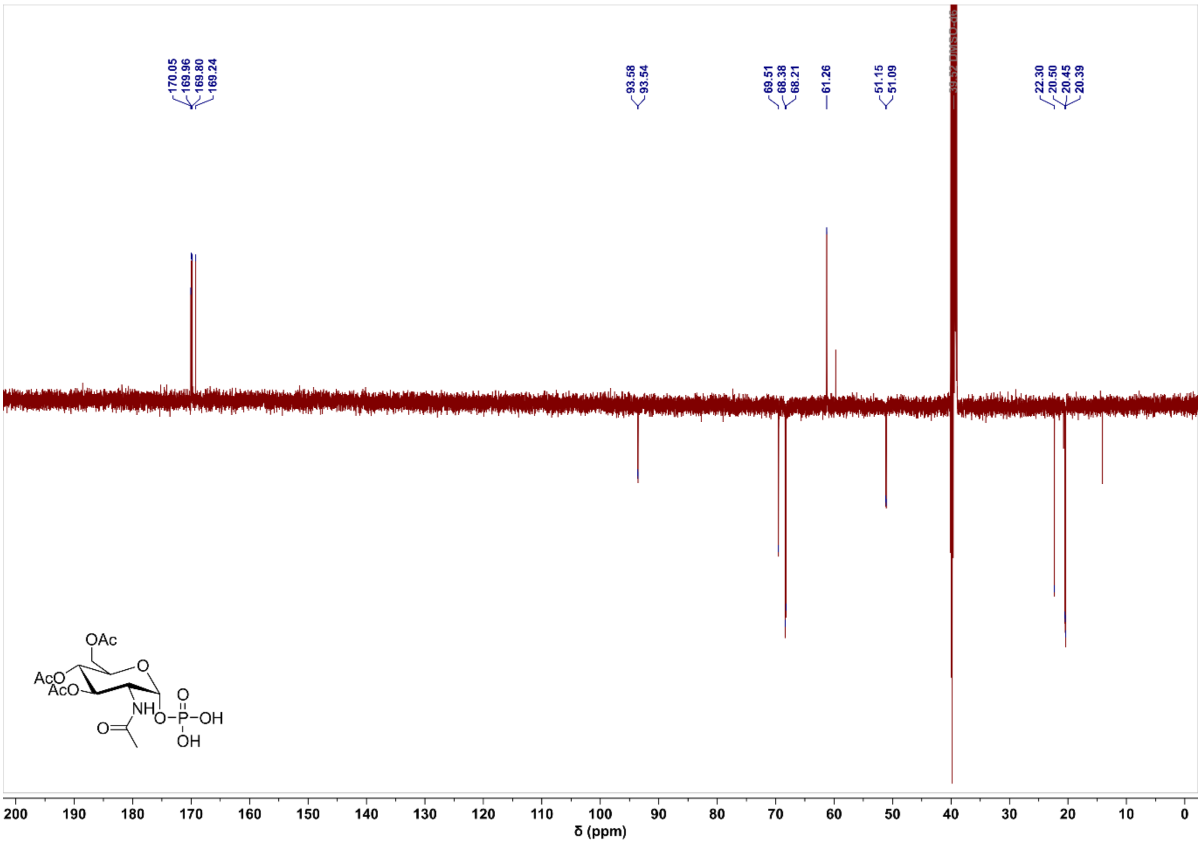

#### Compound 1 $^{31}$P NMR

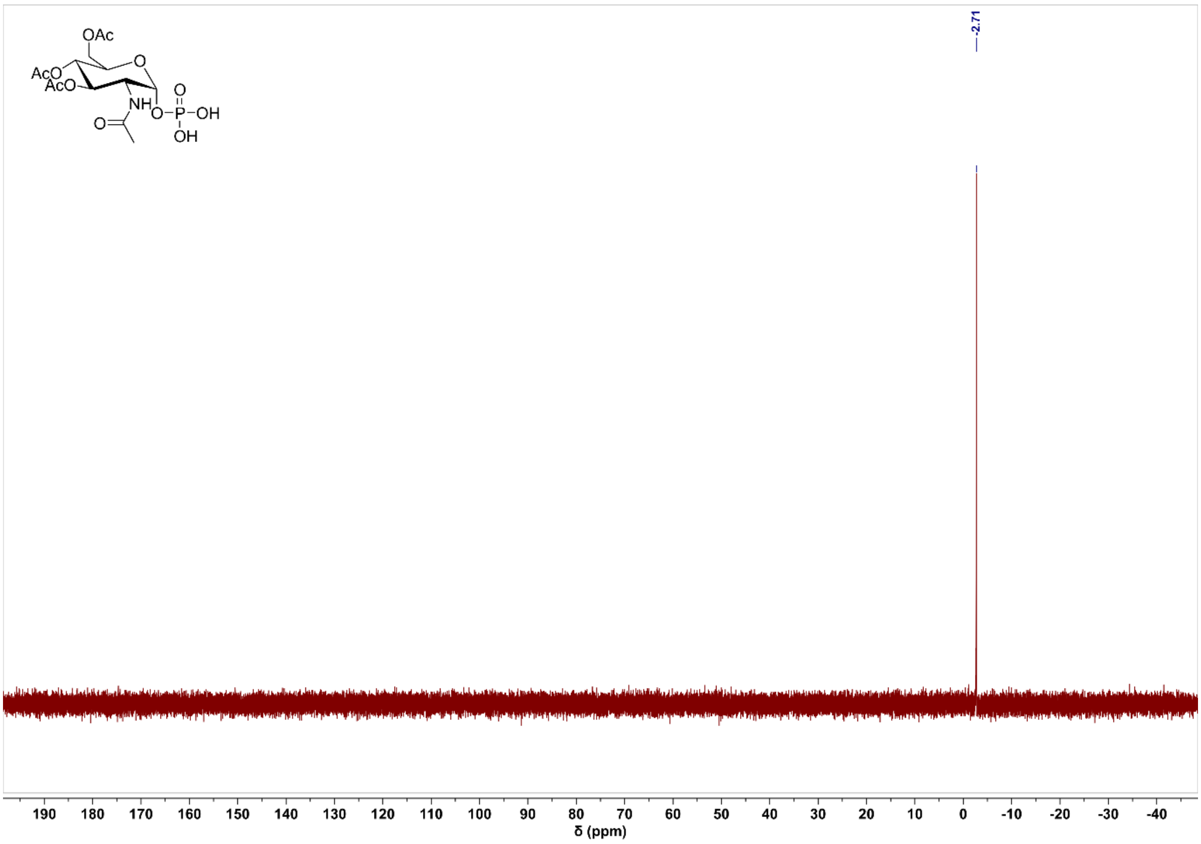

#### Compound 2 $^{1}$H NMR

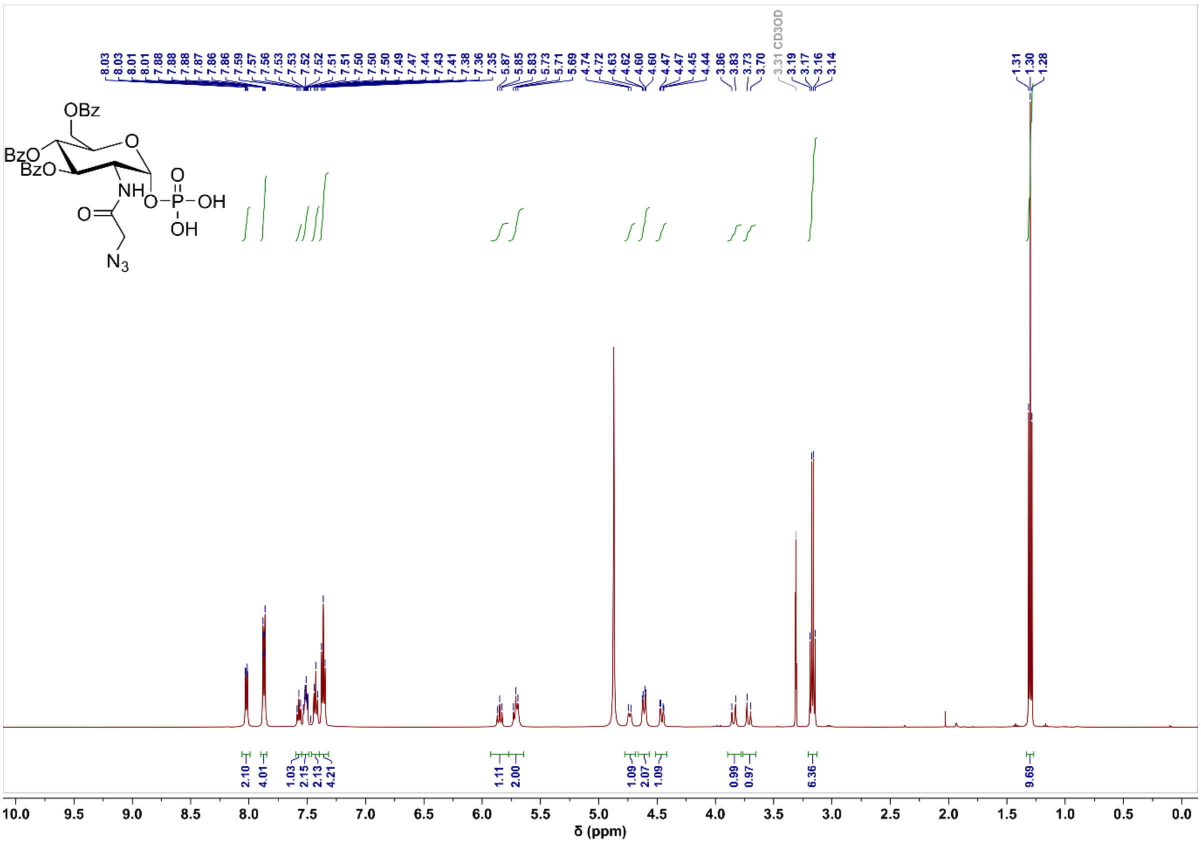

#### Compound 2 $^{13}C$ NMR

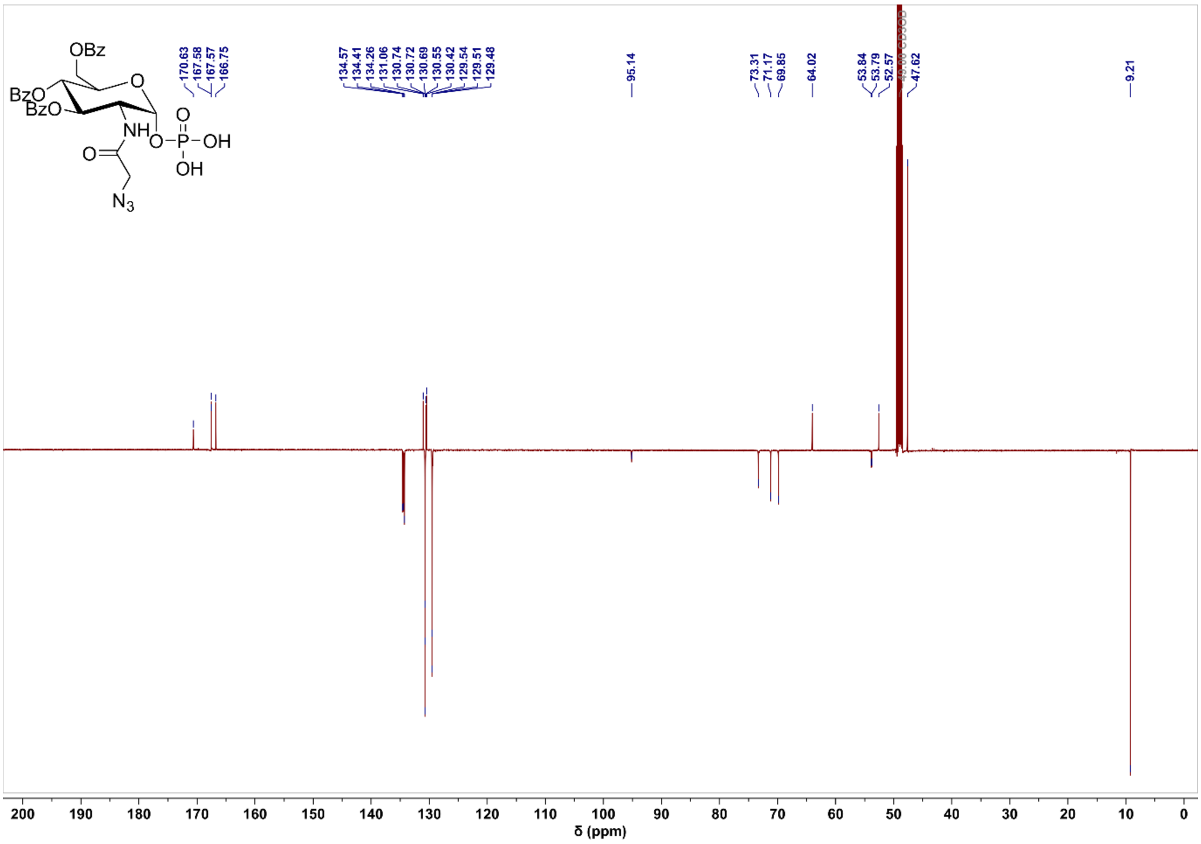

#### Compound 2 $^{31}P$ NMR

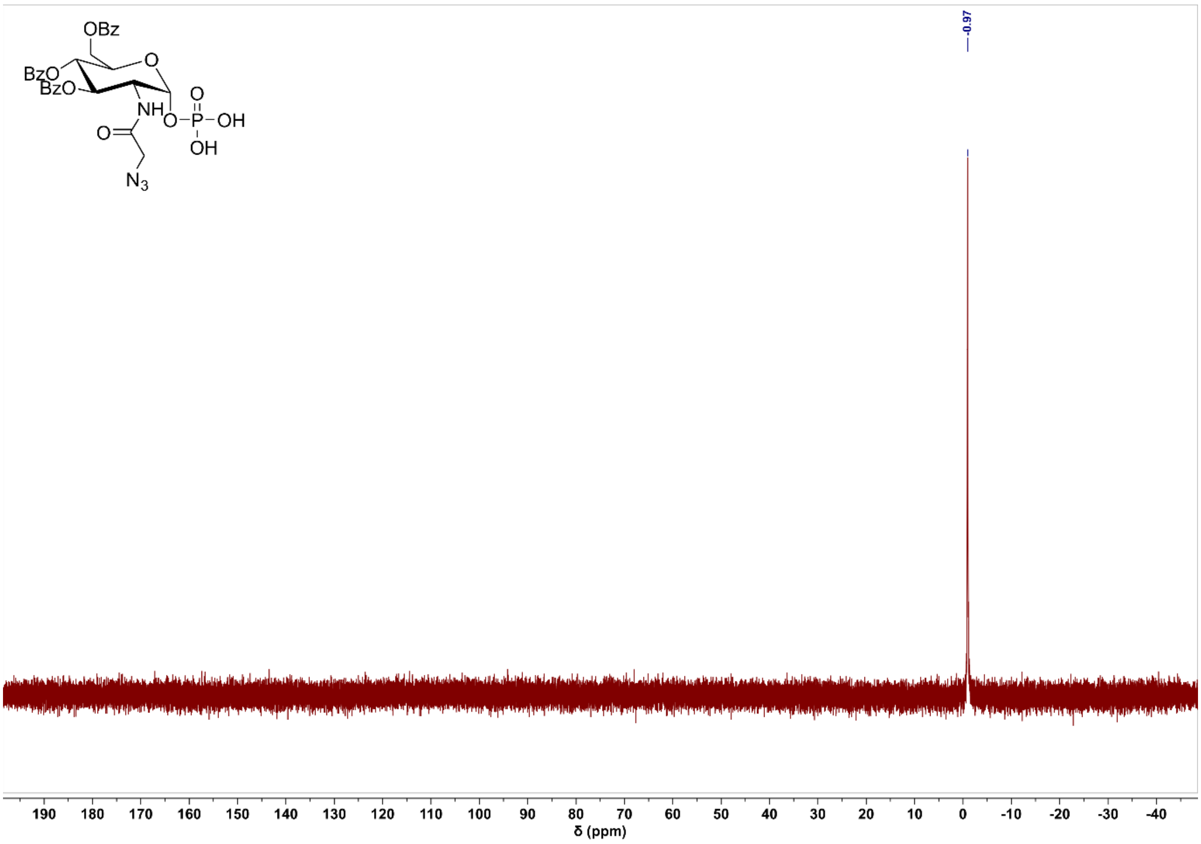

#### Compound 3 $^{1}$H NMR

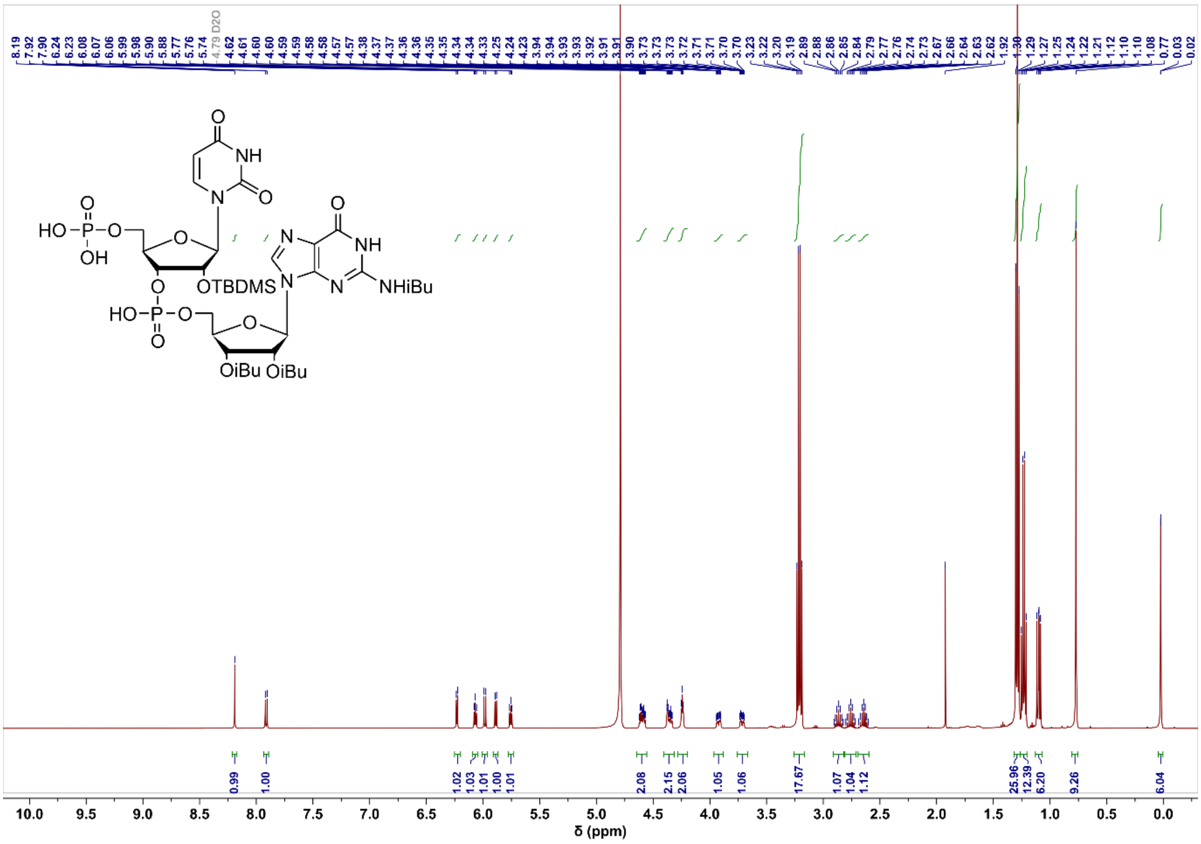

#### Compound 3 $^{13}$C NMR

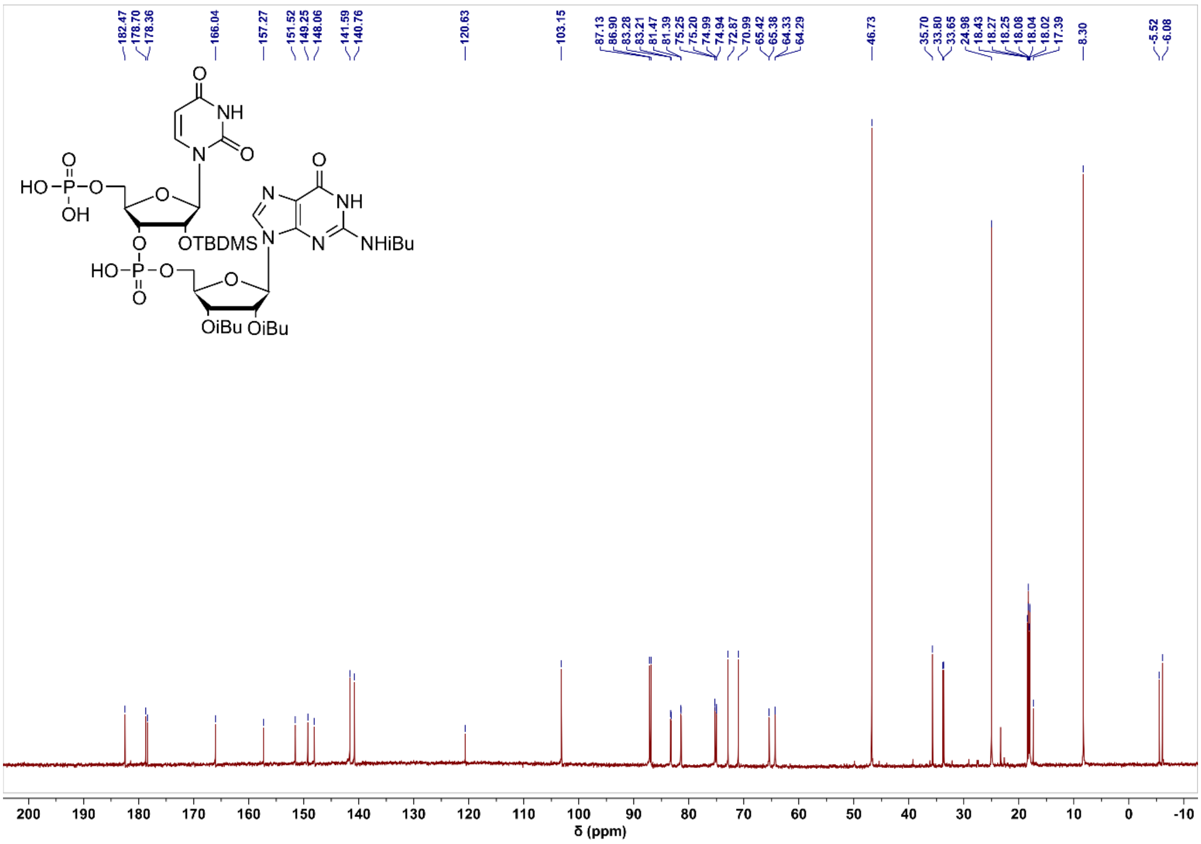

#### Compound 3 $^{31}$P NMR

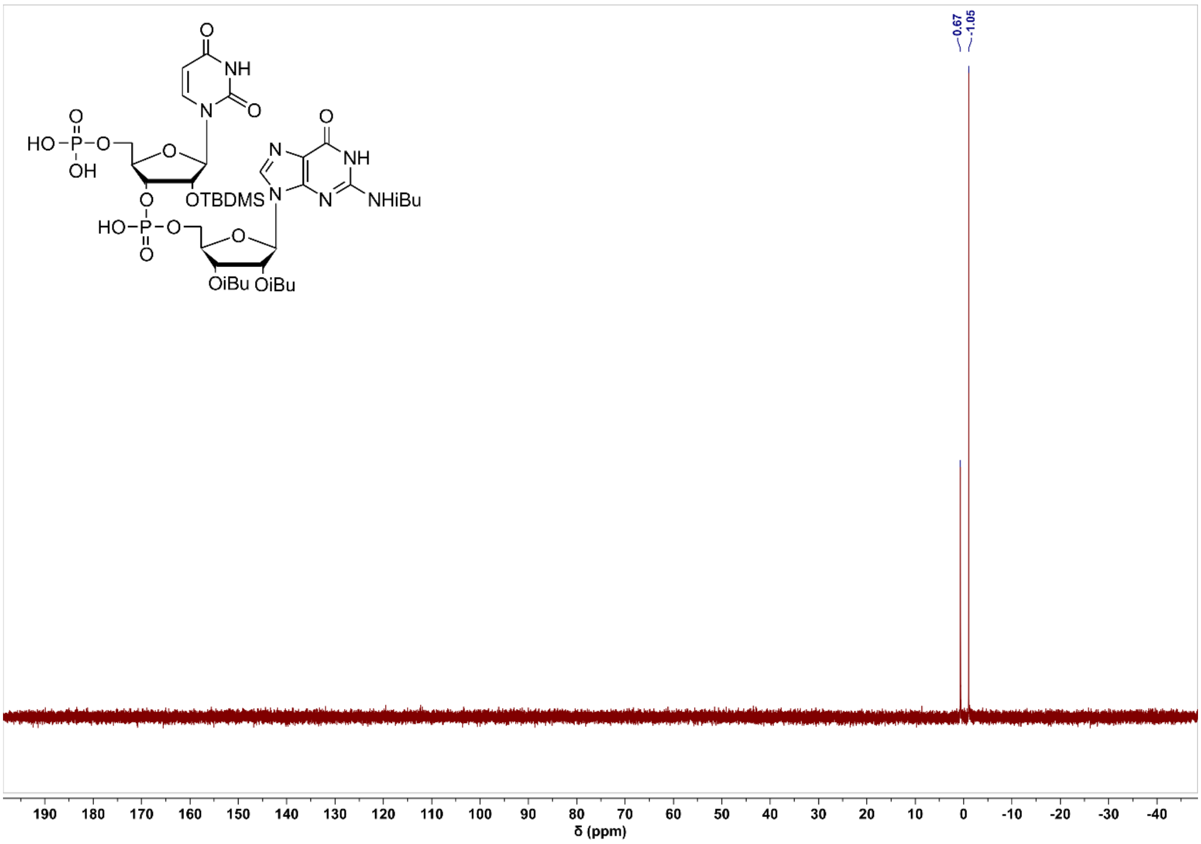

#### Compound 4 intermediate 1 $^{1}$H NMR

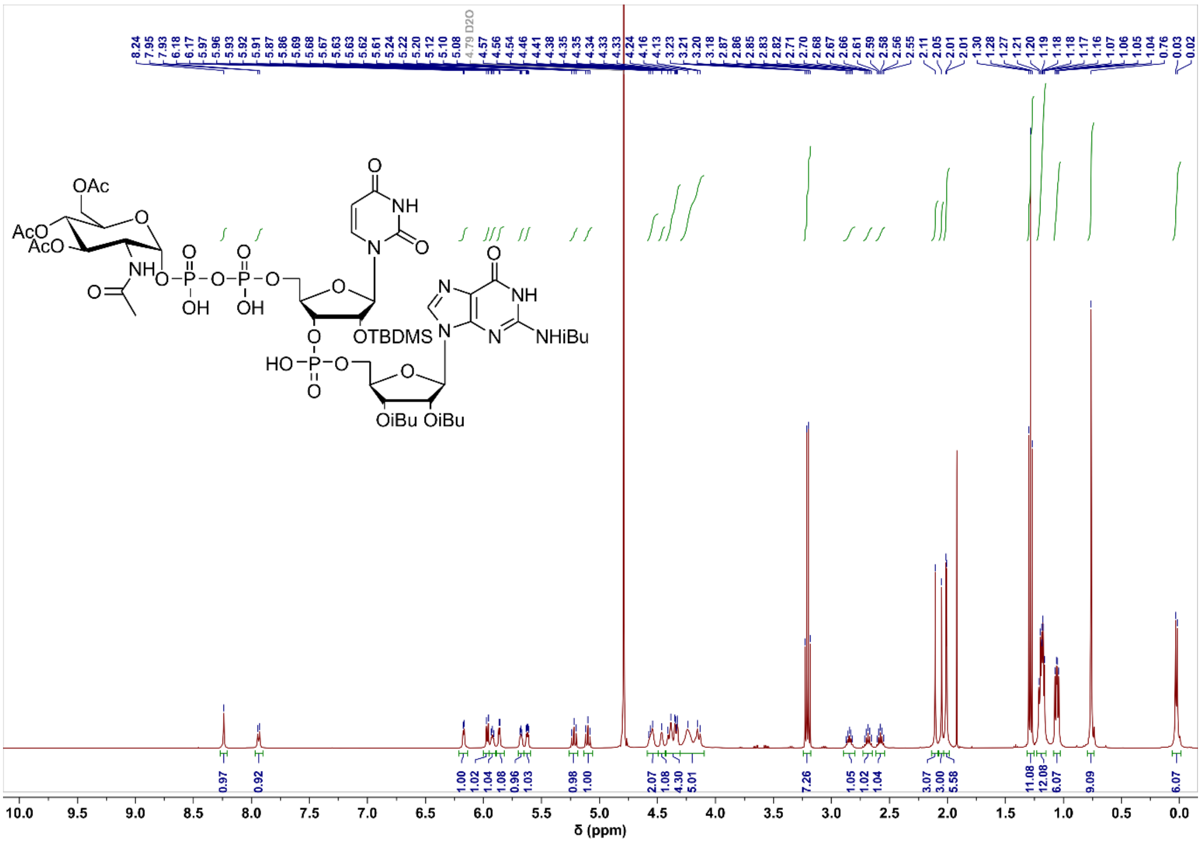

#### Compound 4 intermediate 1 $^{13}C$ NMR

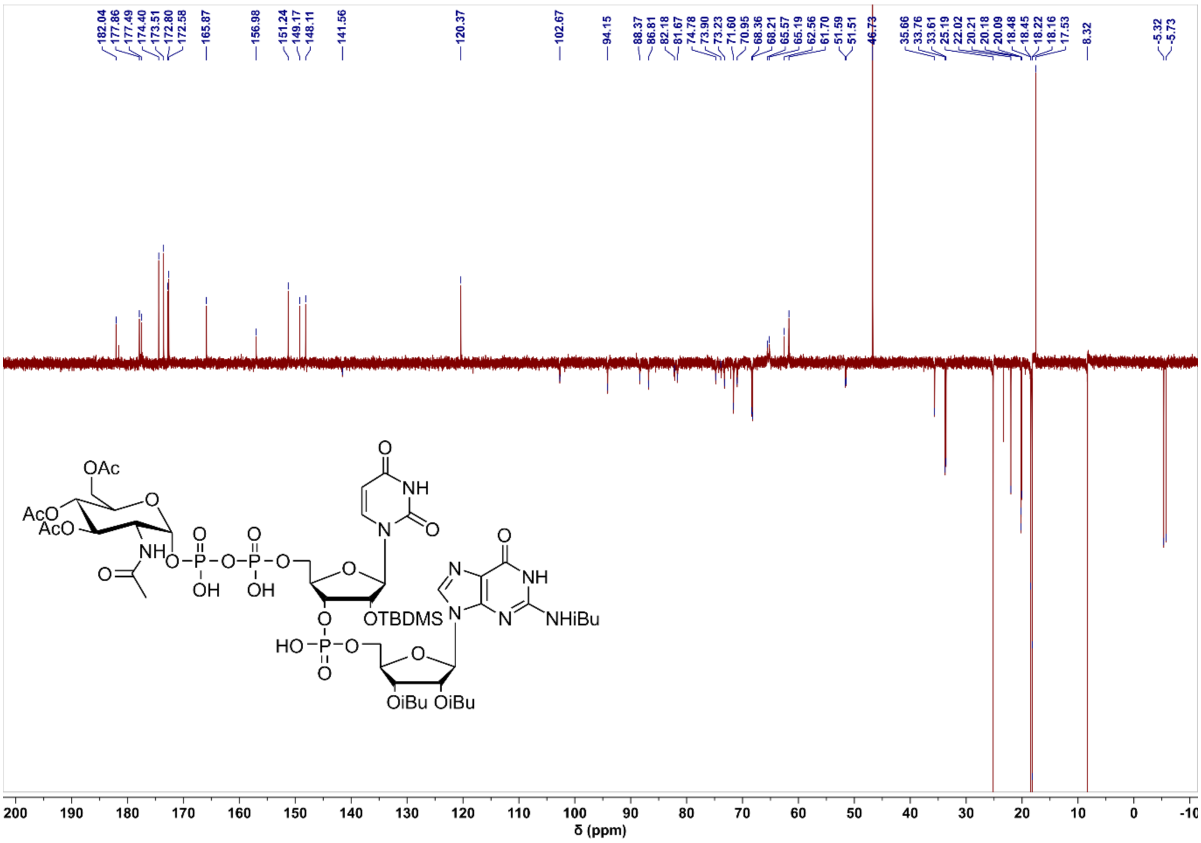

#### Compound 4 intermediate 1 $^{31}P$ NMR

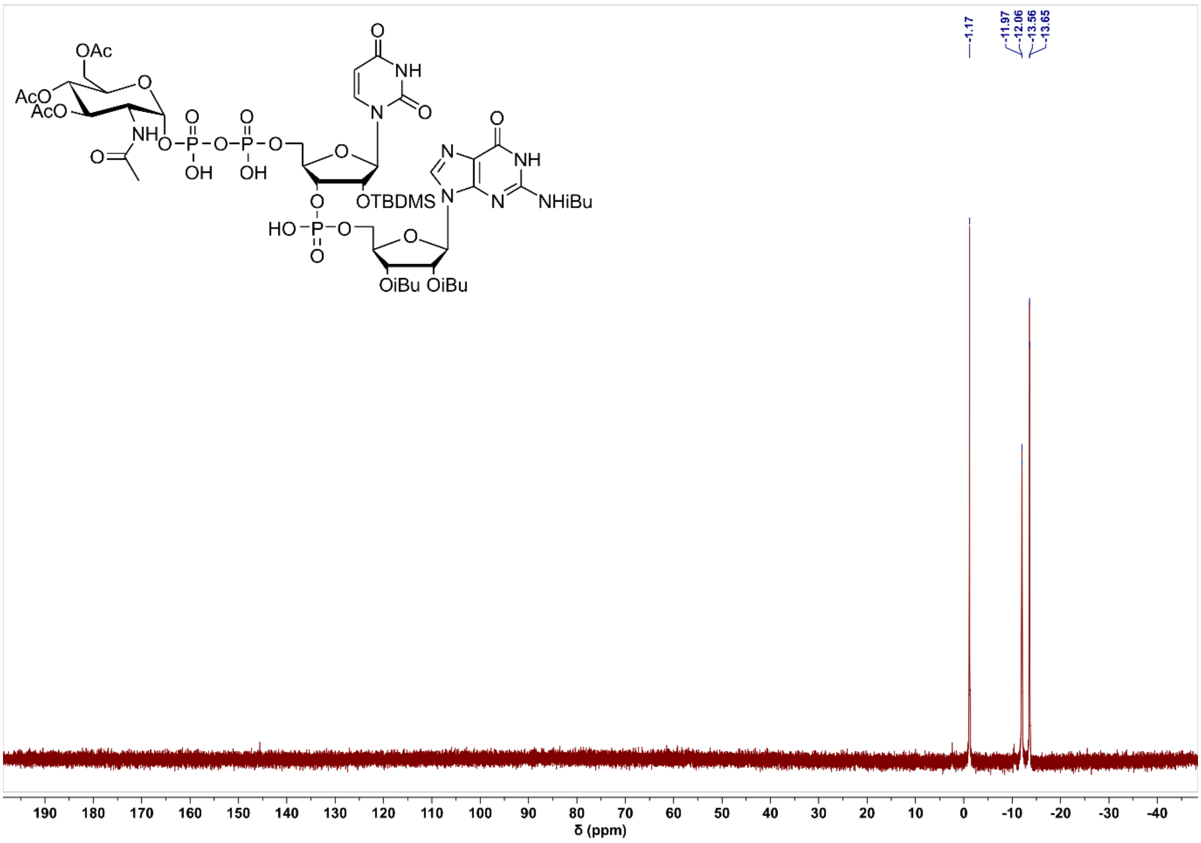

#### Compound 4 intermediate 2 $^{1}H$ NMR

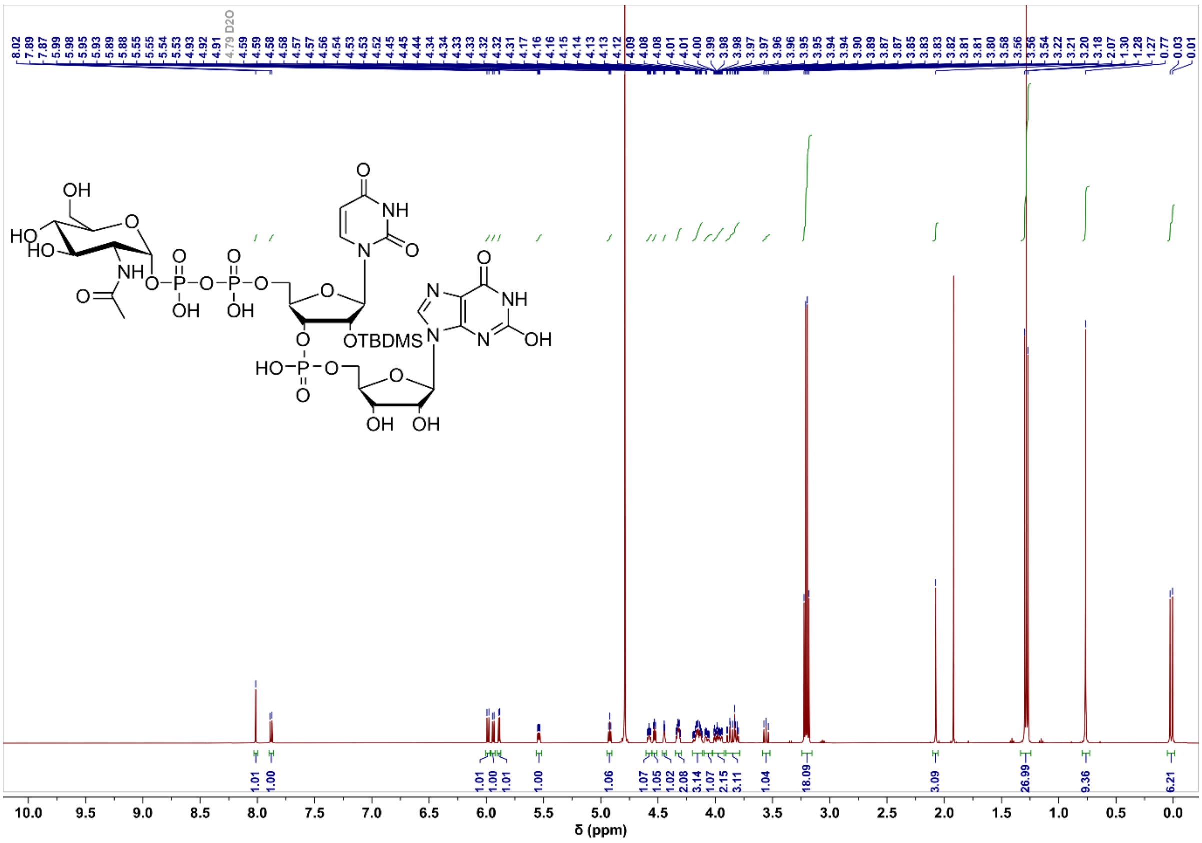

#### Compound 4 intermediate 2 $^{13}C$ NMR

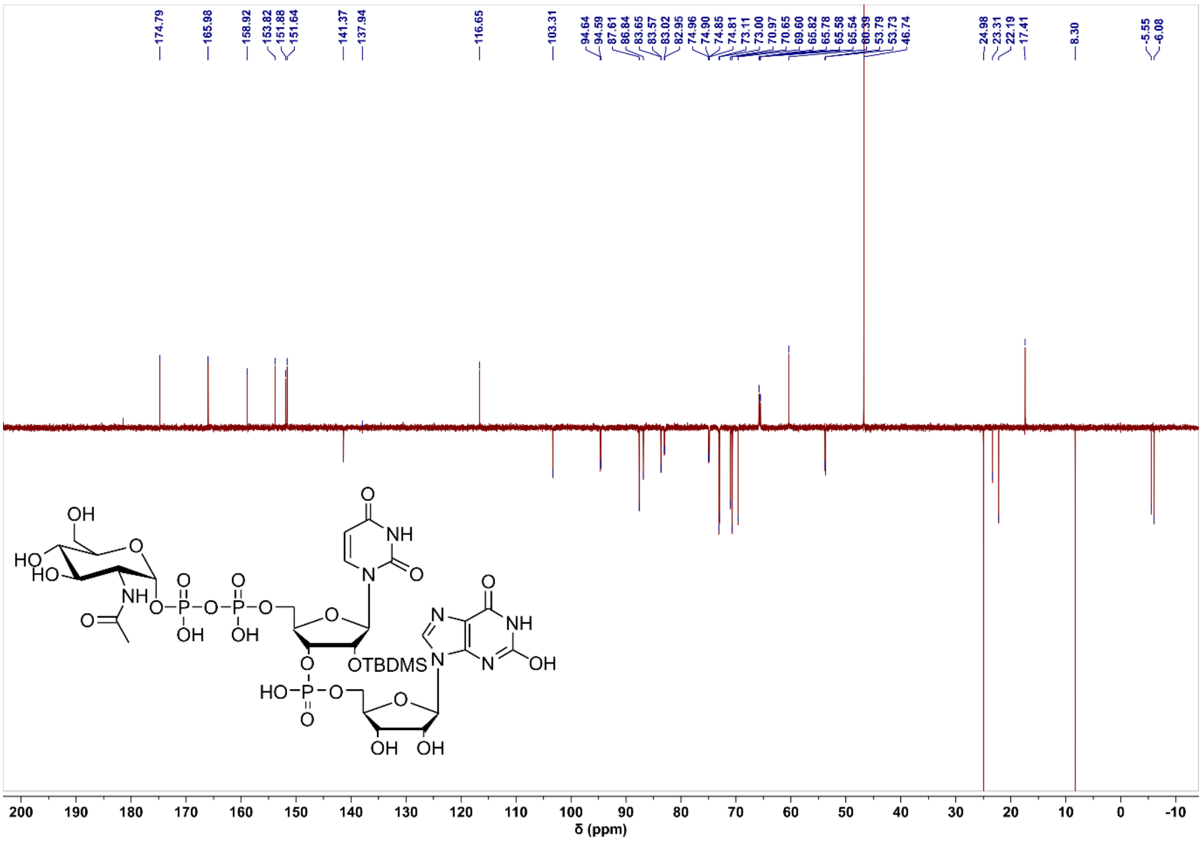

#### Compound 4 intermediate 2 ³¹P NMR

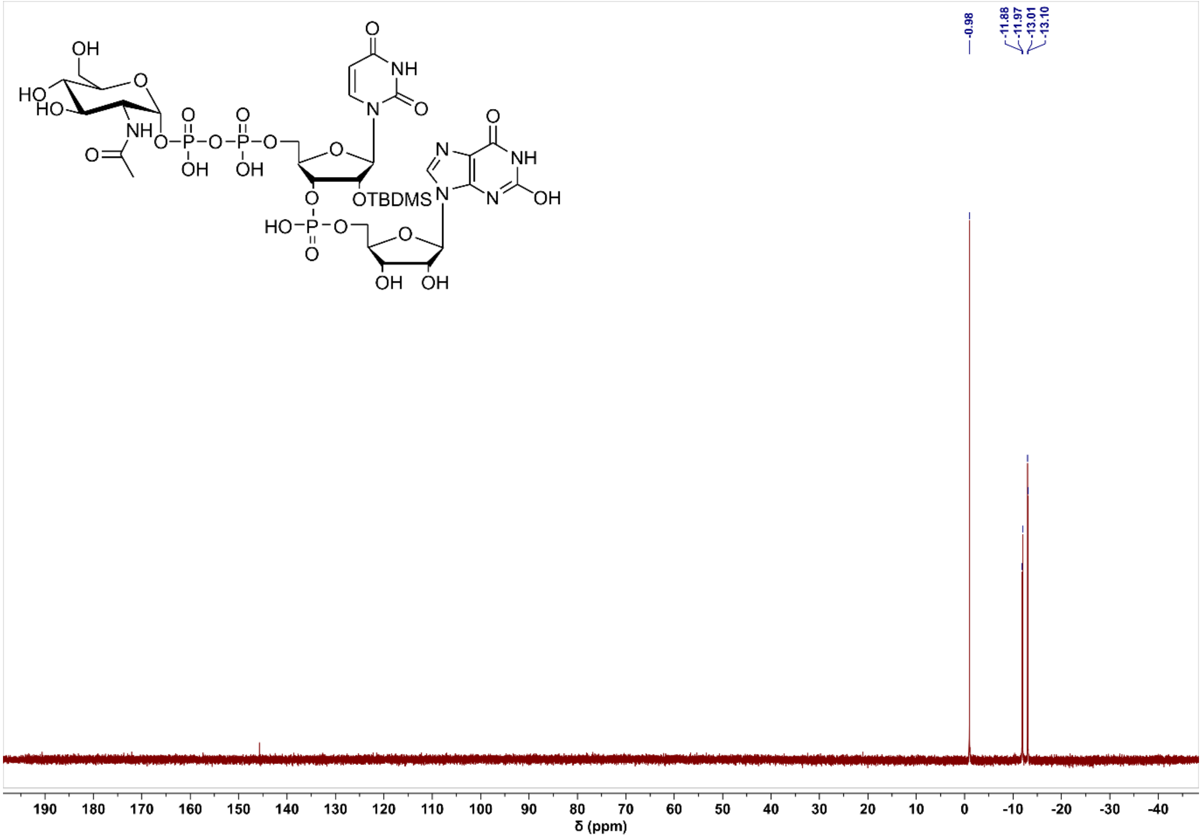

#### Compound 4 product ¹H NMR

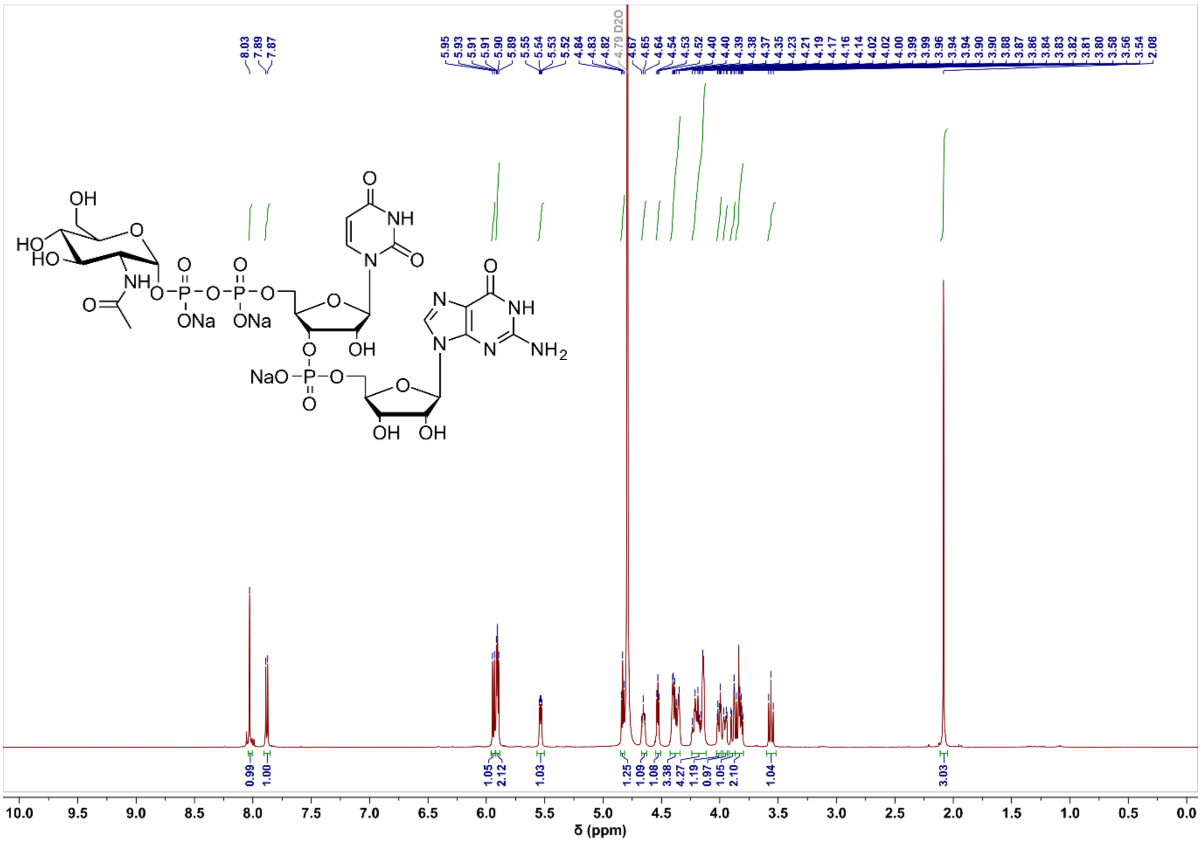

#### Compound 4 product $^{13}C$ NMR

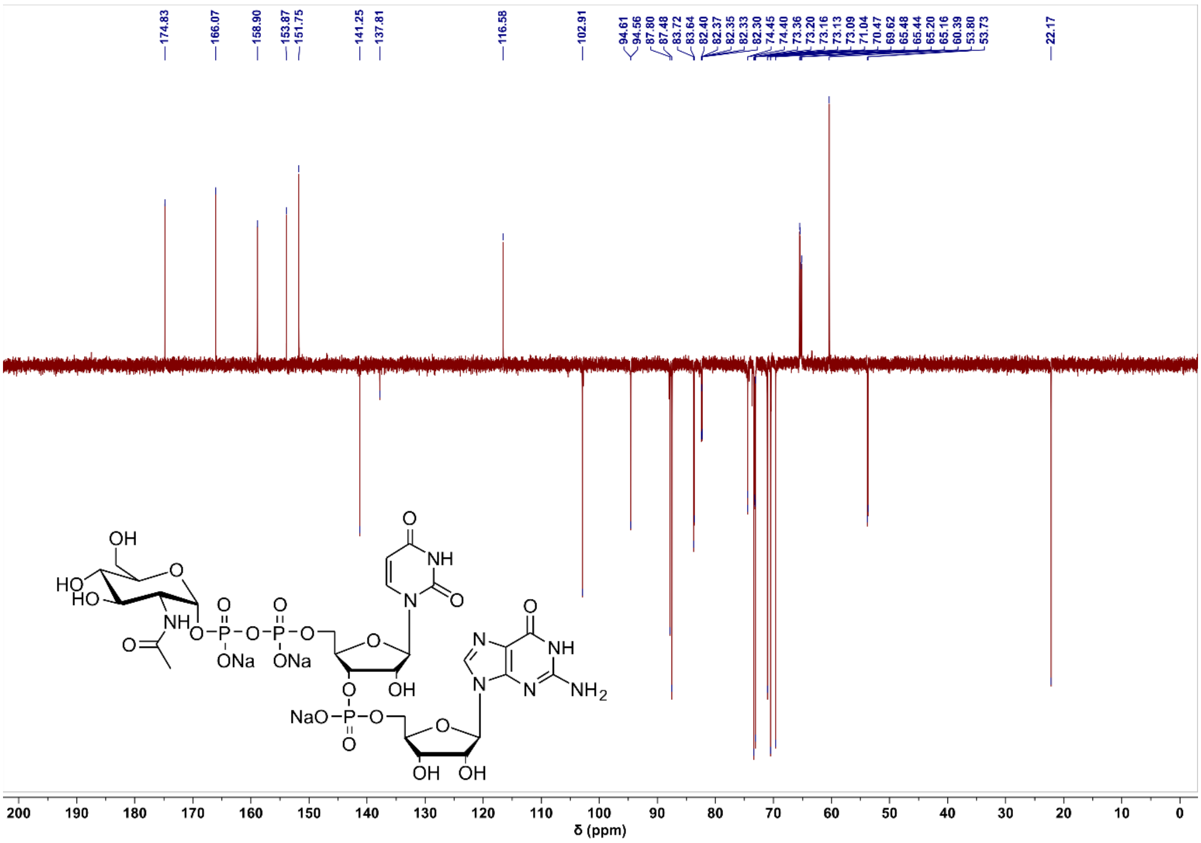

#### Compound 4 product $^{31}P$ NMR

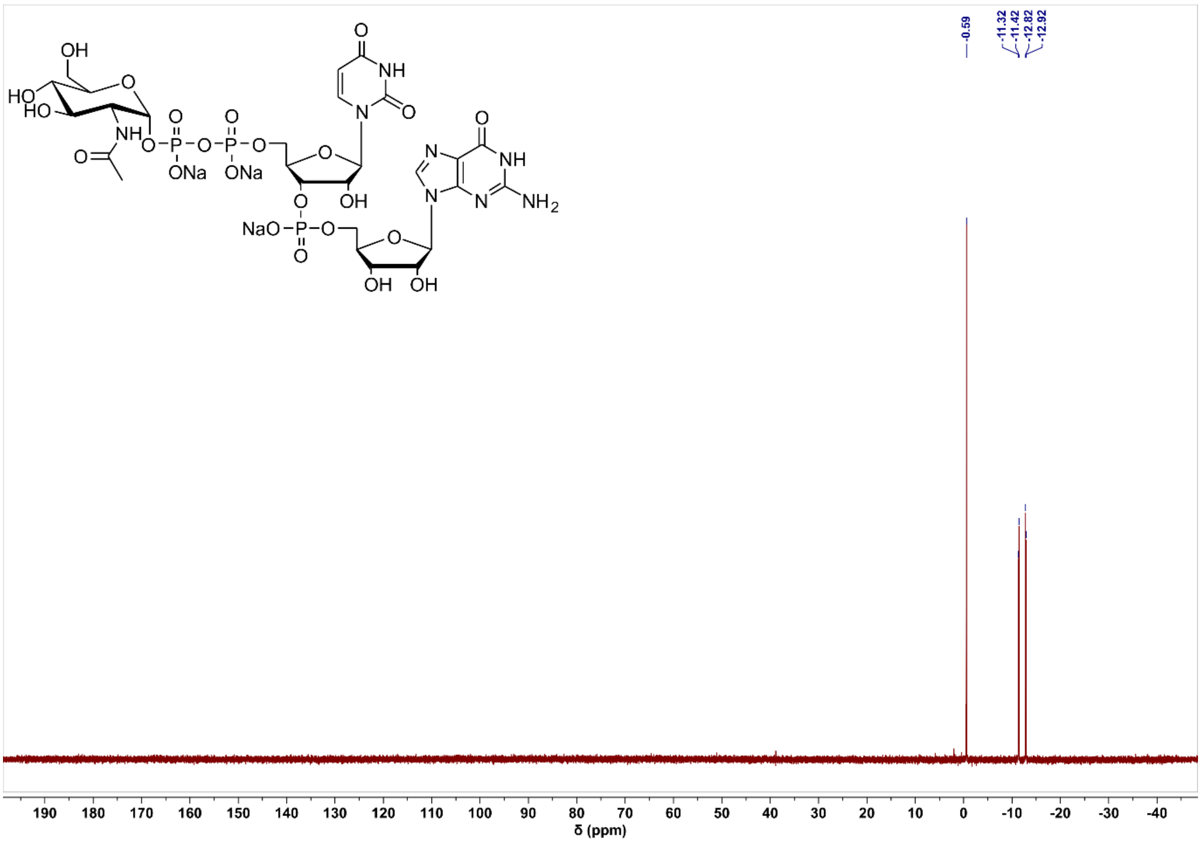

#### Compound 5 intermediate 2 $^1$H NMR

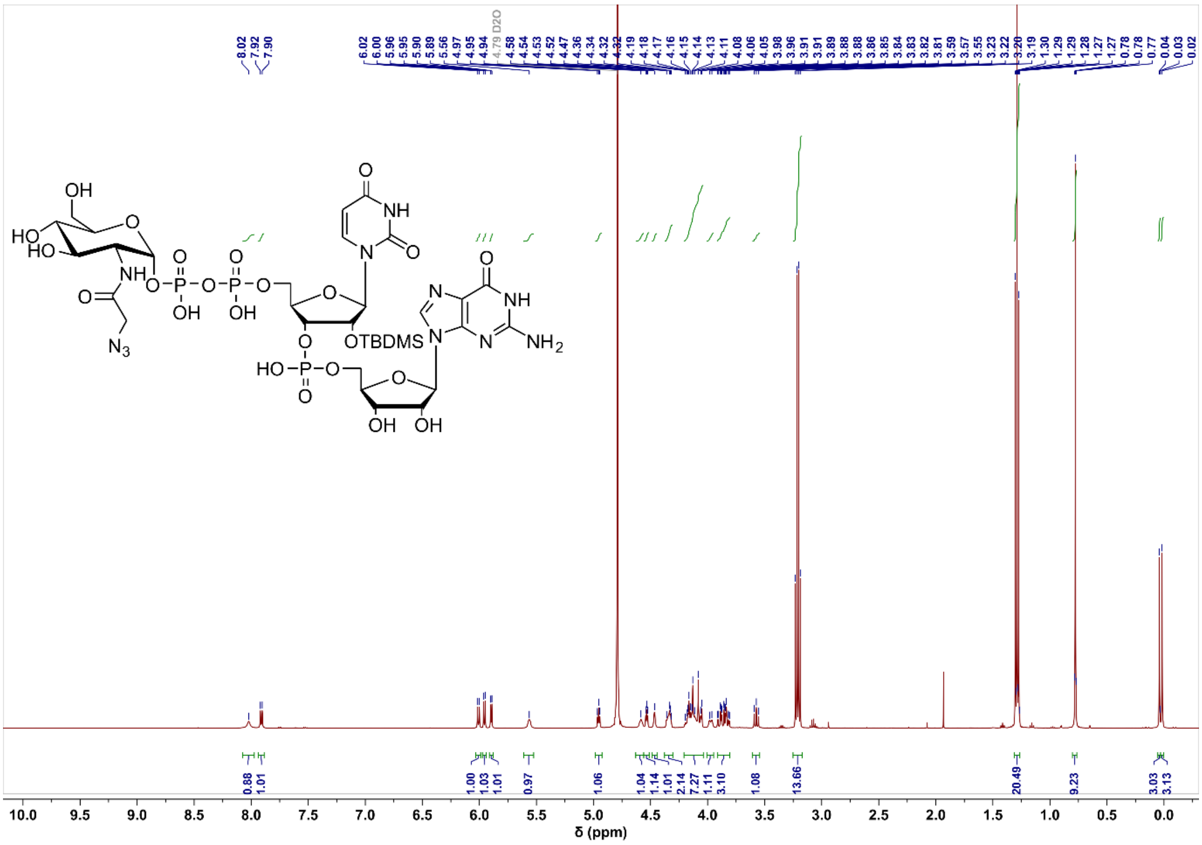

#### Compound 5 intermediate 2 $^{13}$C NMR

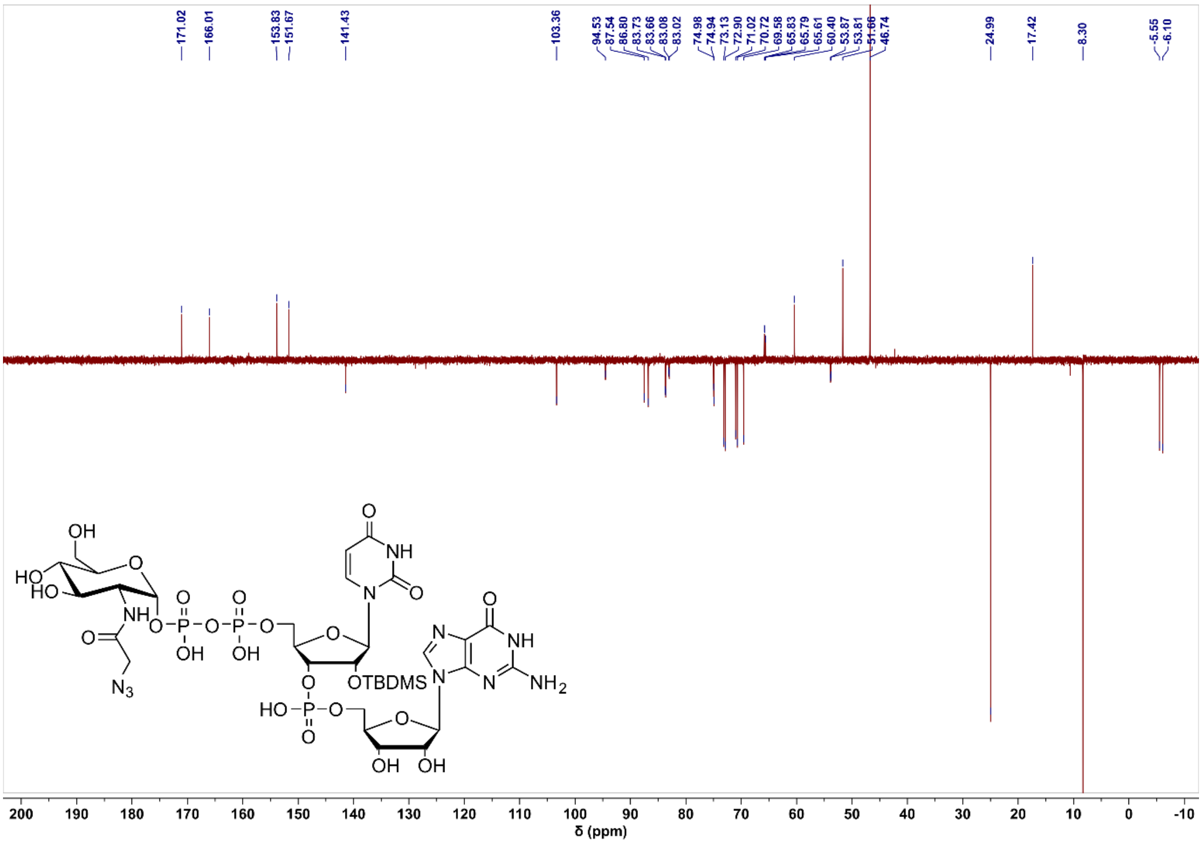

#### Compound 5 intermediate 2 $^{31}$P NMR

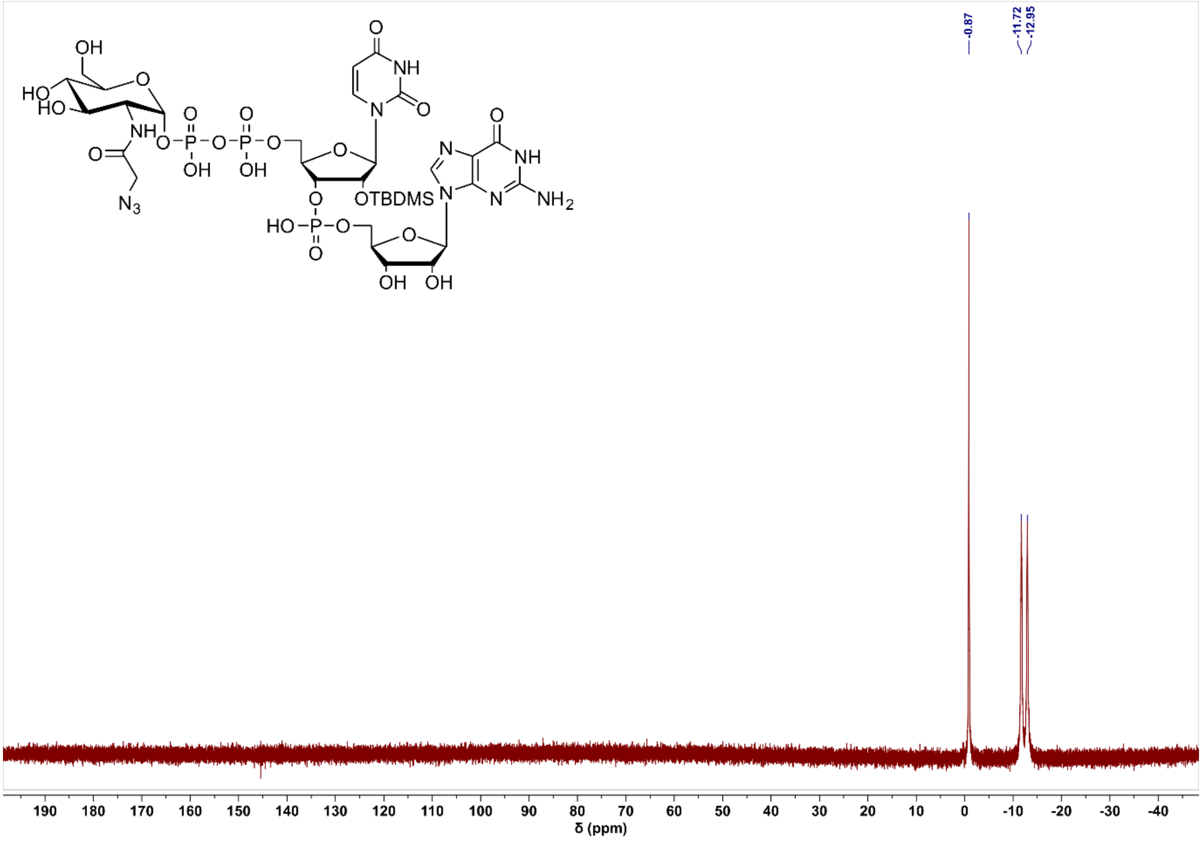

#### Compound 5 product $^1$H NMR

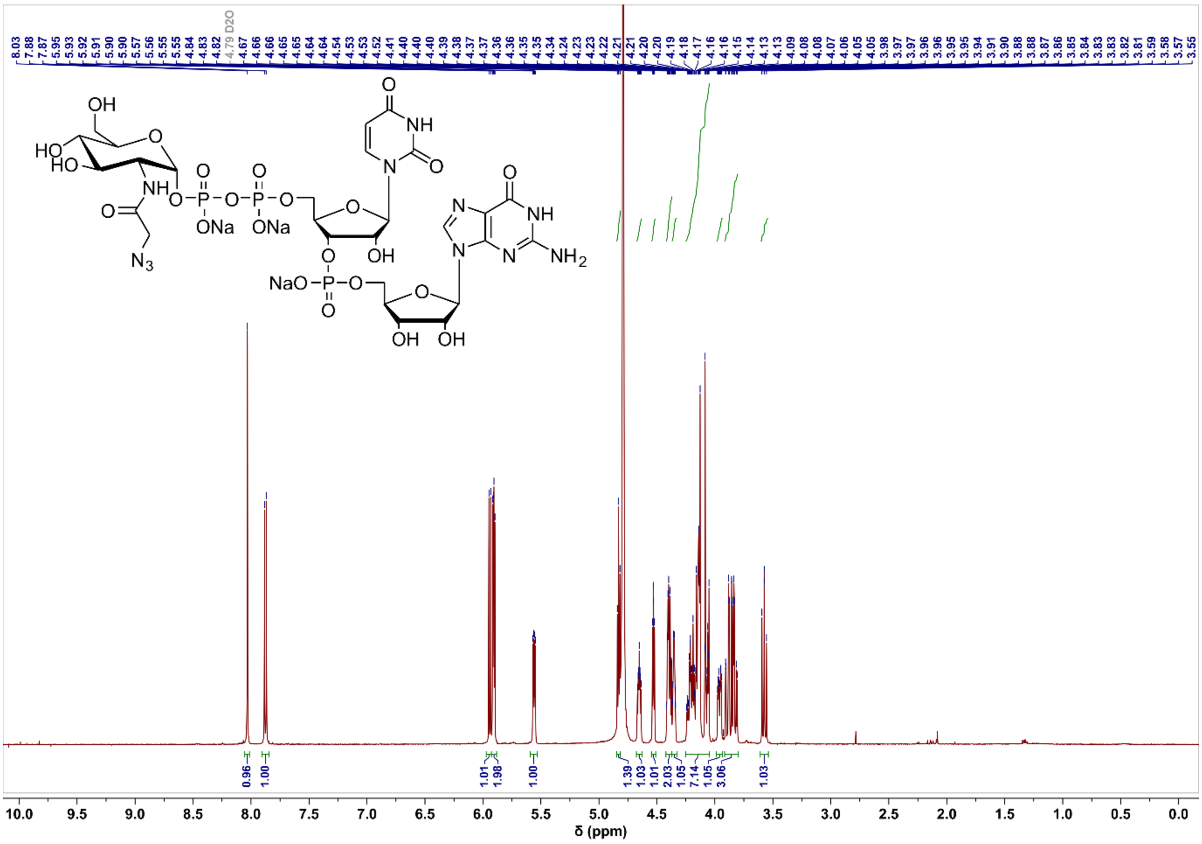

#### Compound 5 product $^{13}C$ NMR

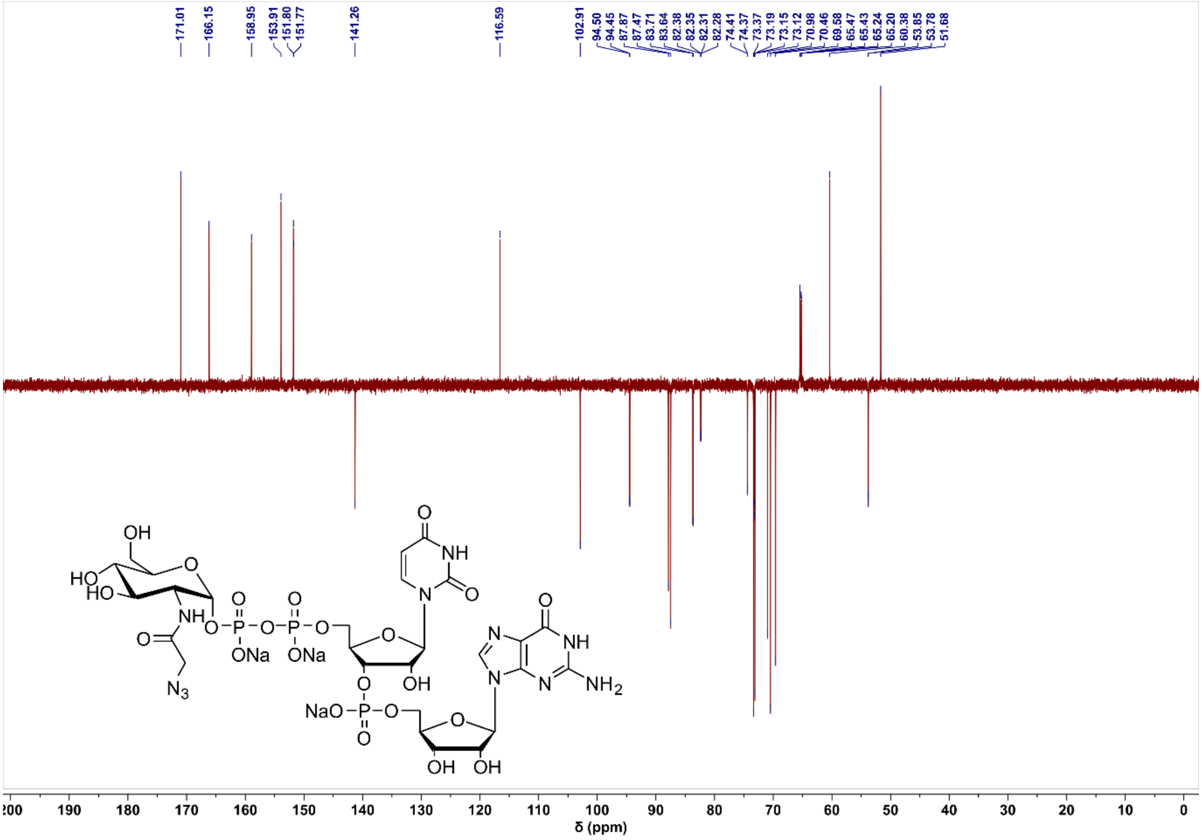

#### Compound 5 product $^{31}P$ NMR

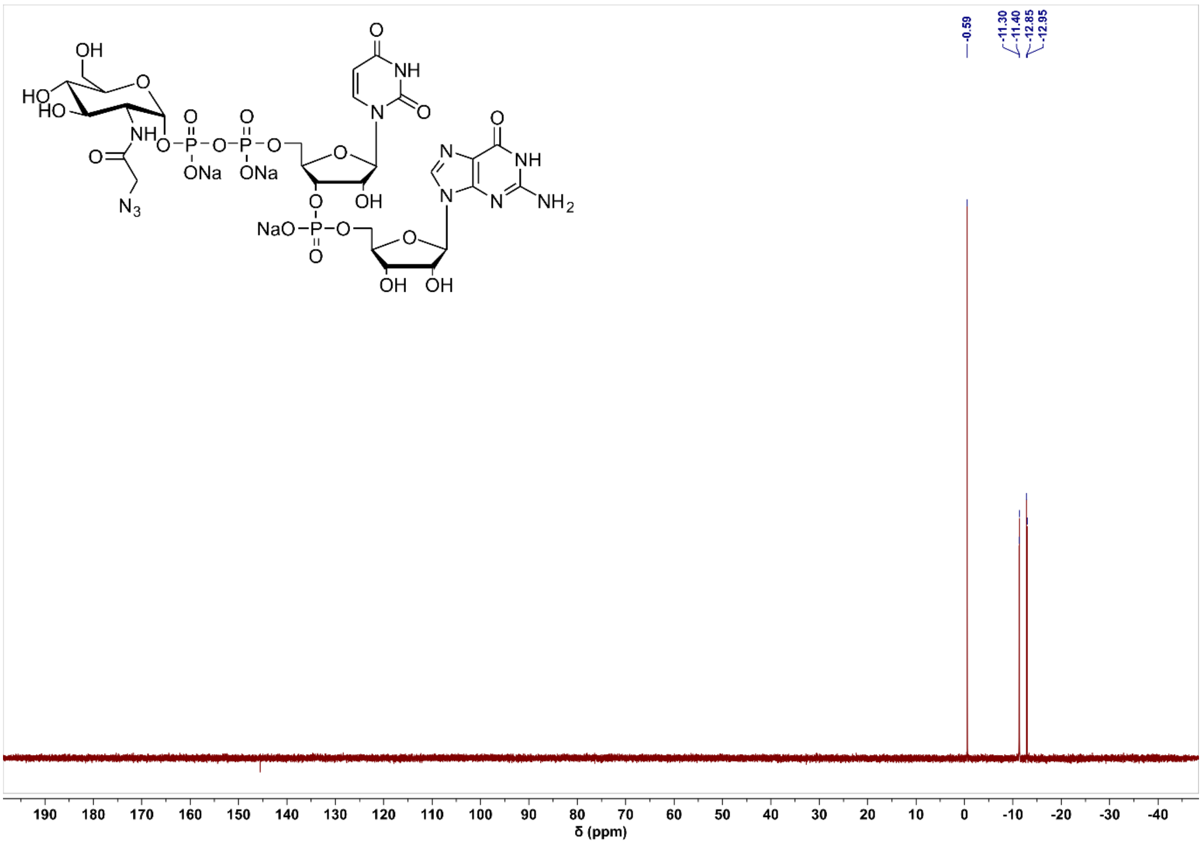

#### Compound 6 $^{1}H$ NMR

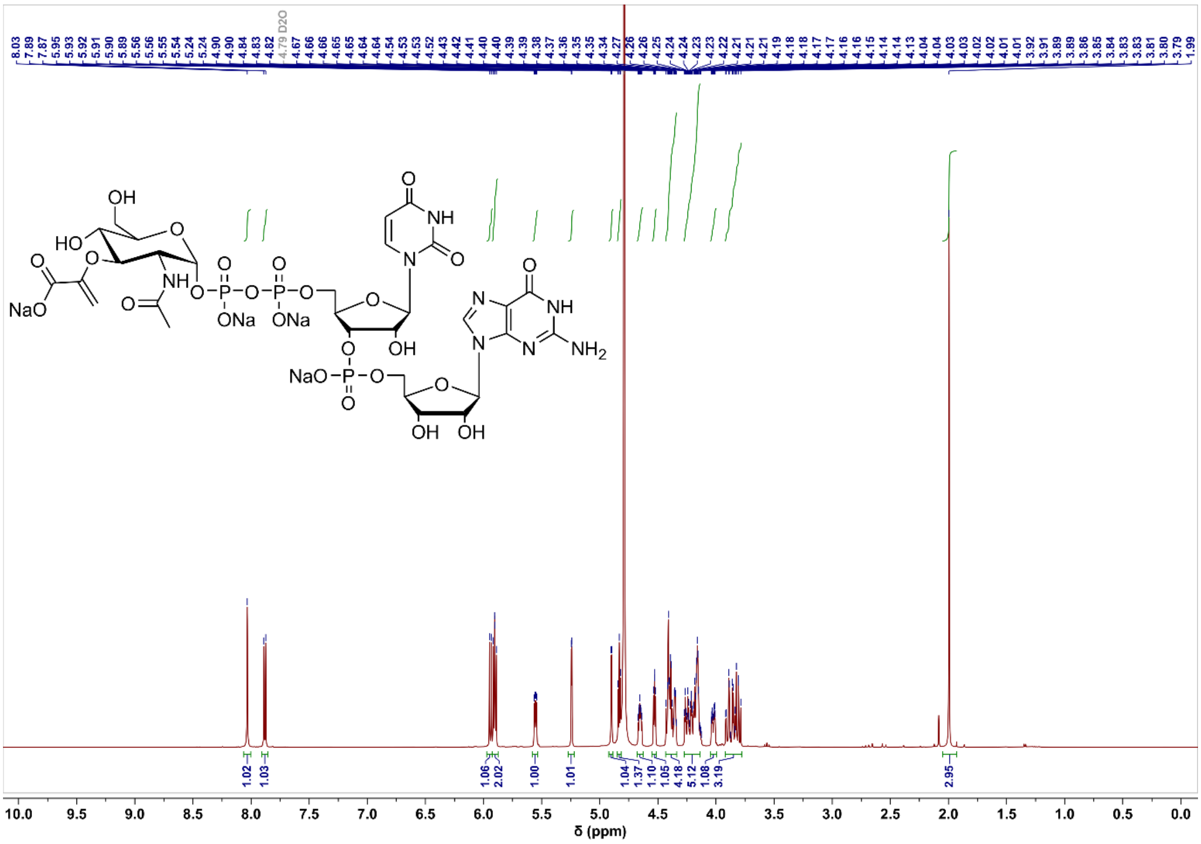

#### Compound 6 $^{13}C$ NMR

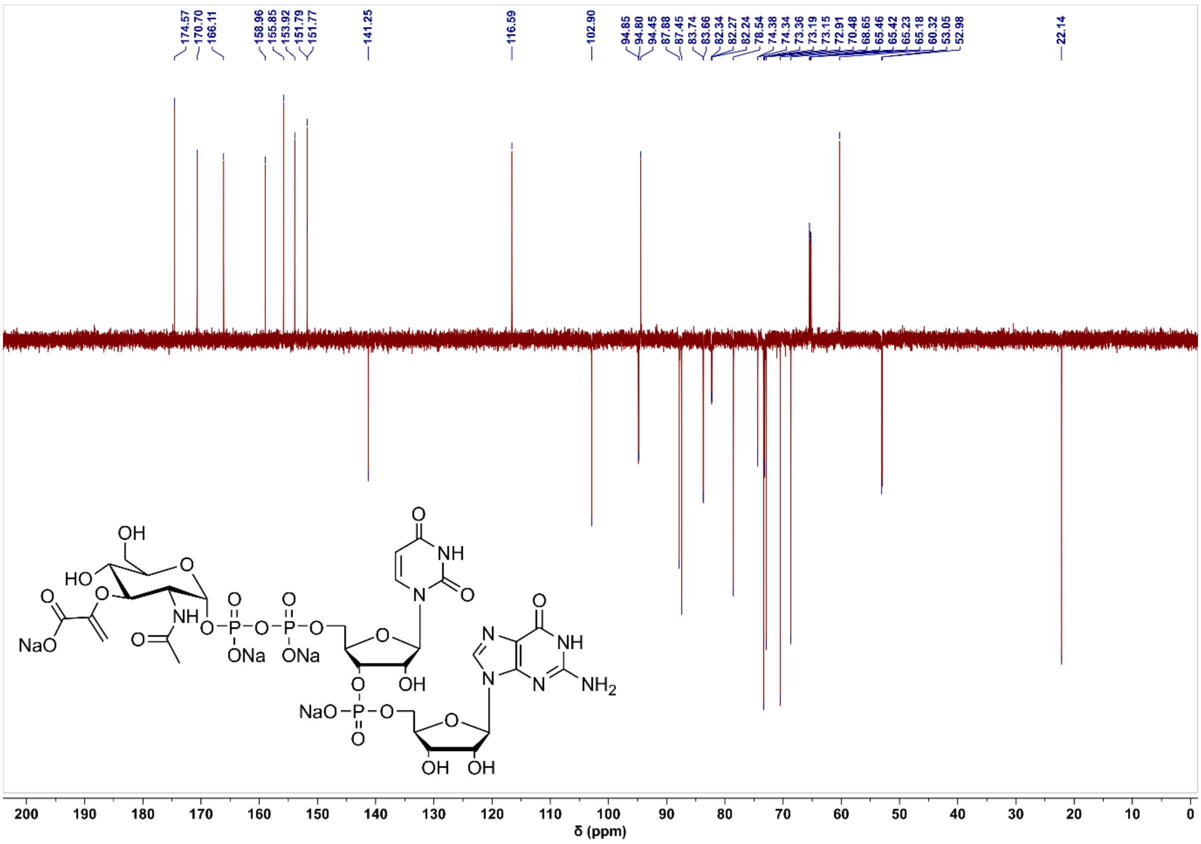

#### Compound 6 $^{31}$P NMR

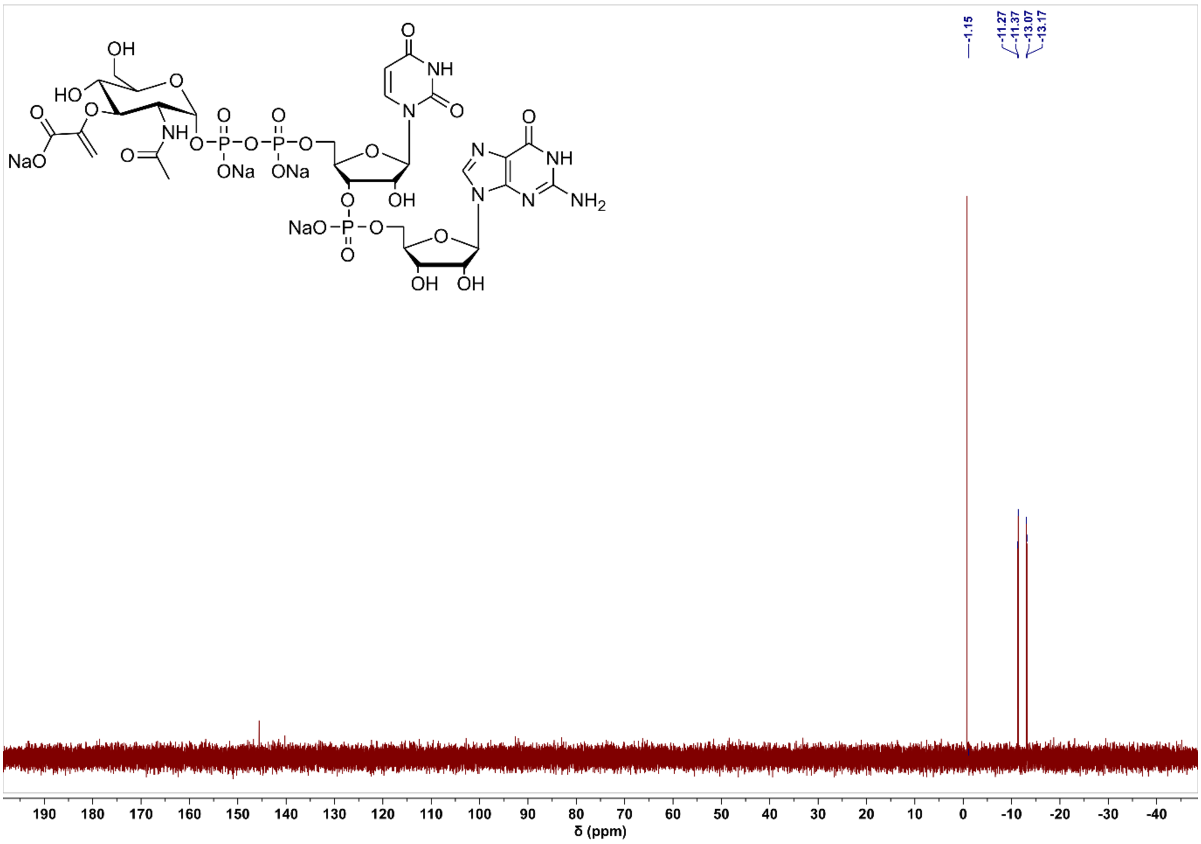

#### Compound 7 $^{1}$H NMR

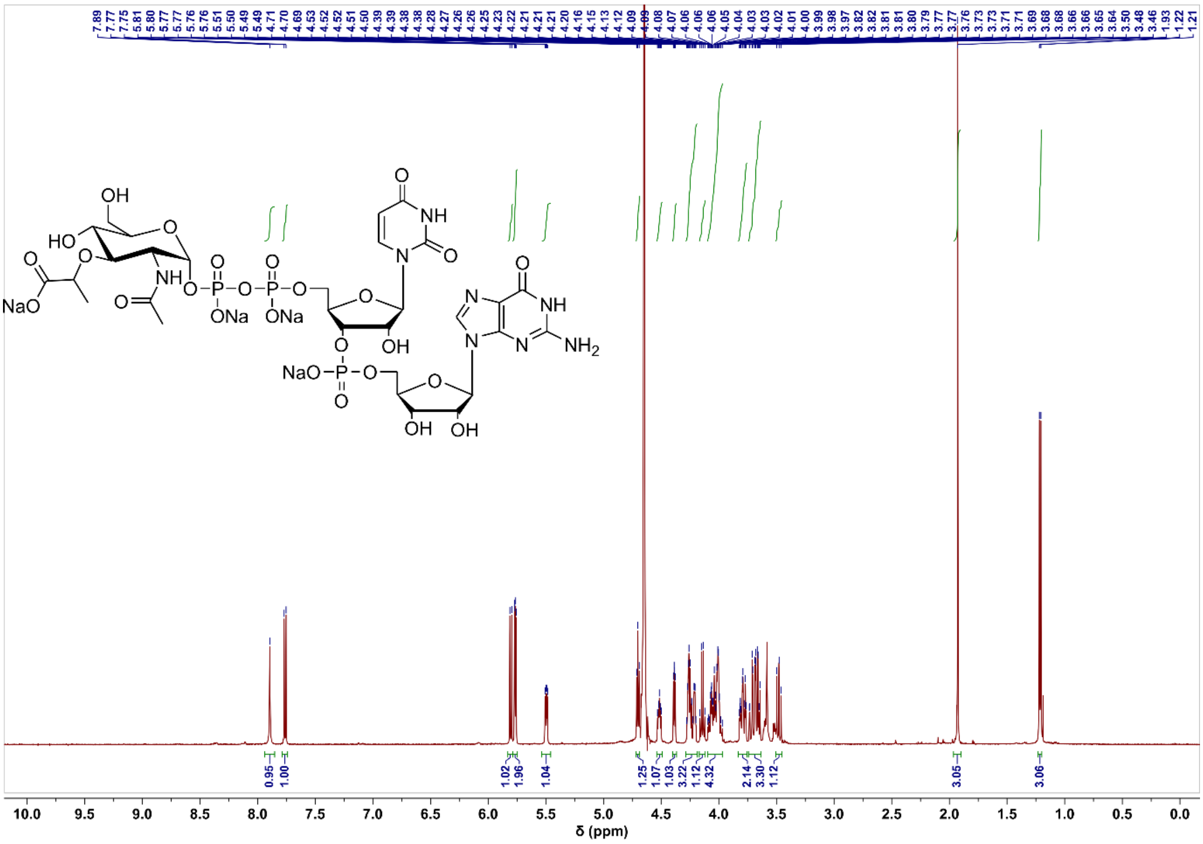

#### Compound 7 $^{13}C$ NMR

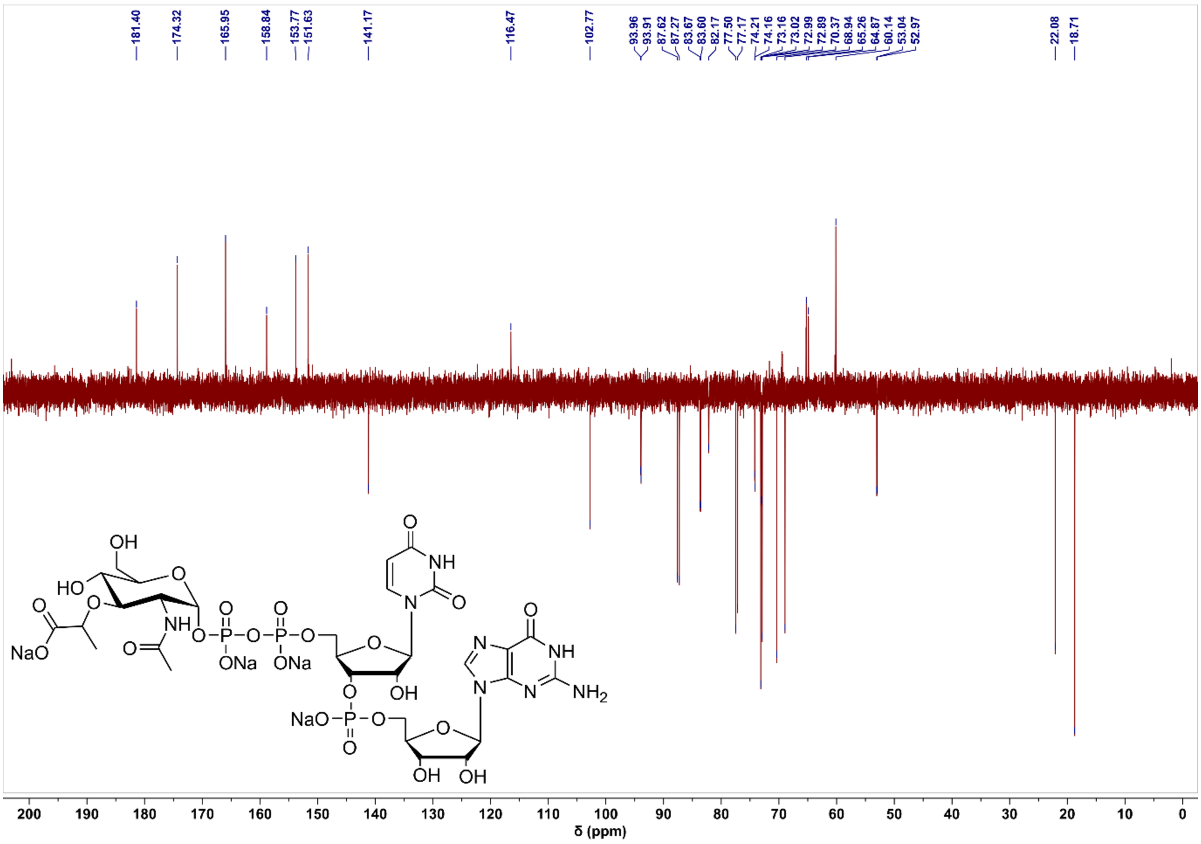

#### Compound 7 $^{31}P$ NMR

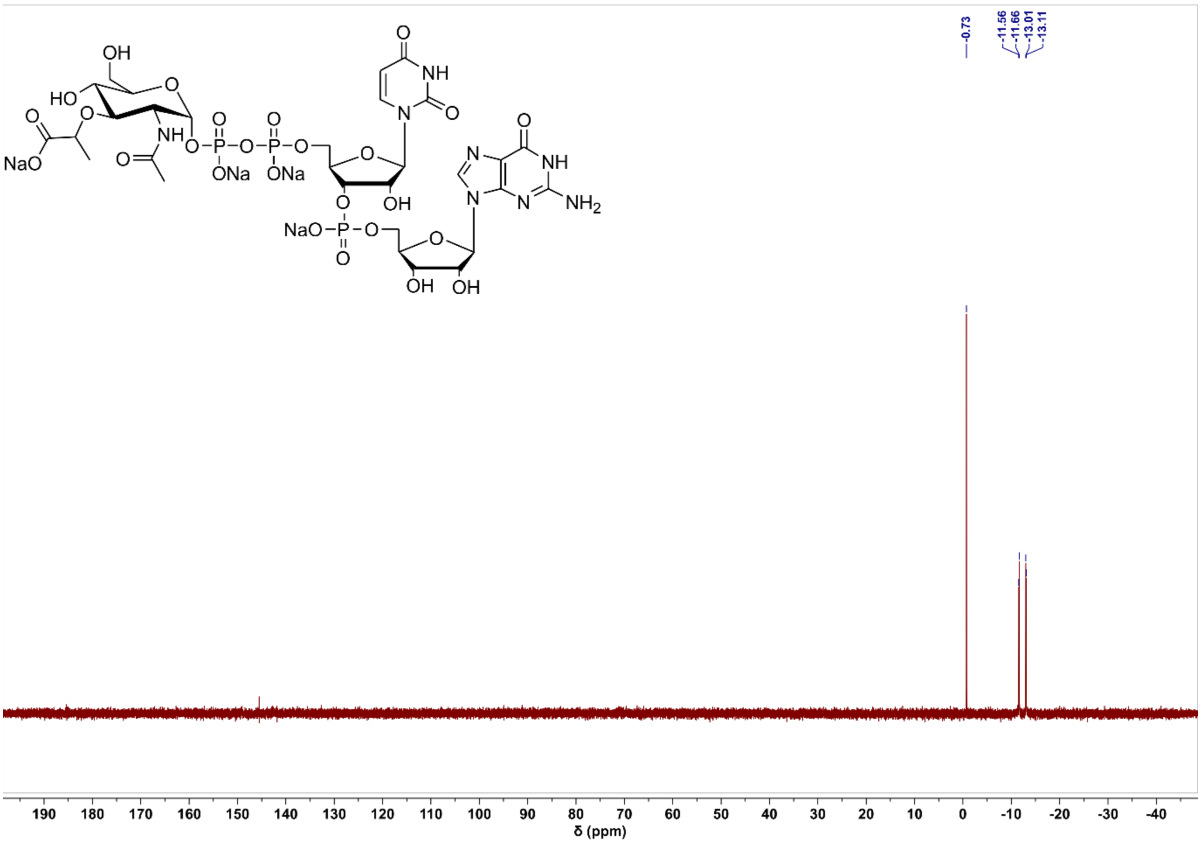

#### Compound 16 $^{1}$H NMR

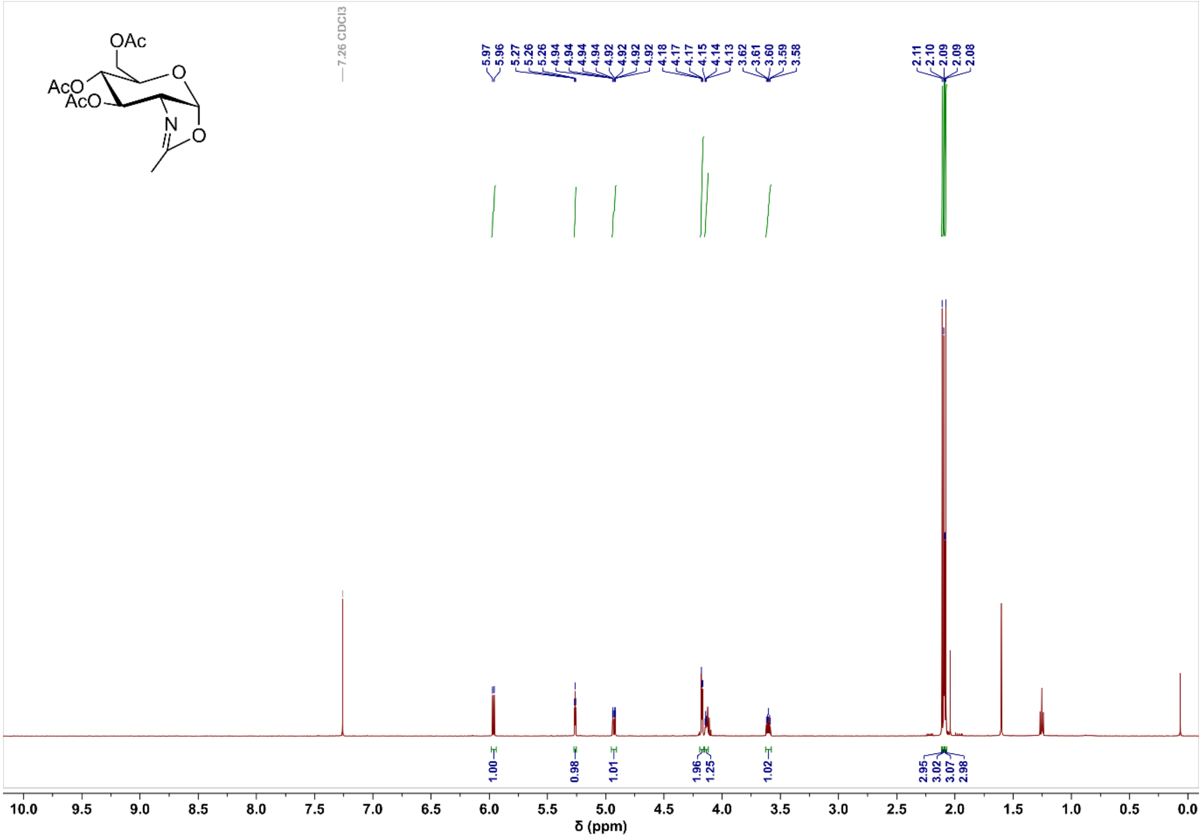

#### Compound 16 $^{13}$C NMR

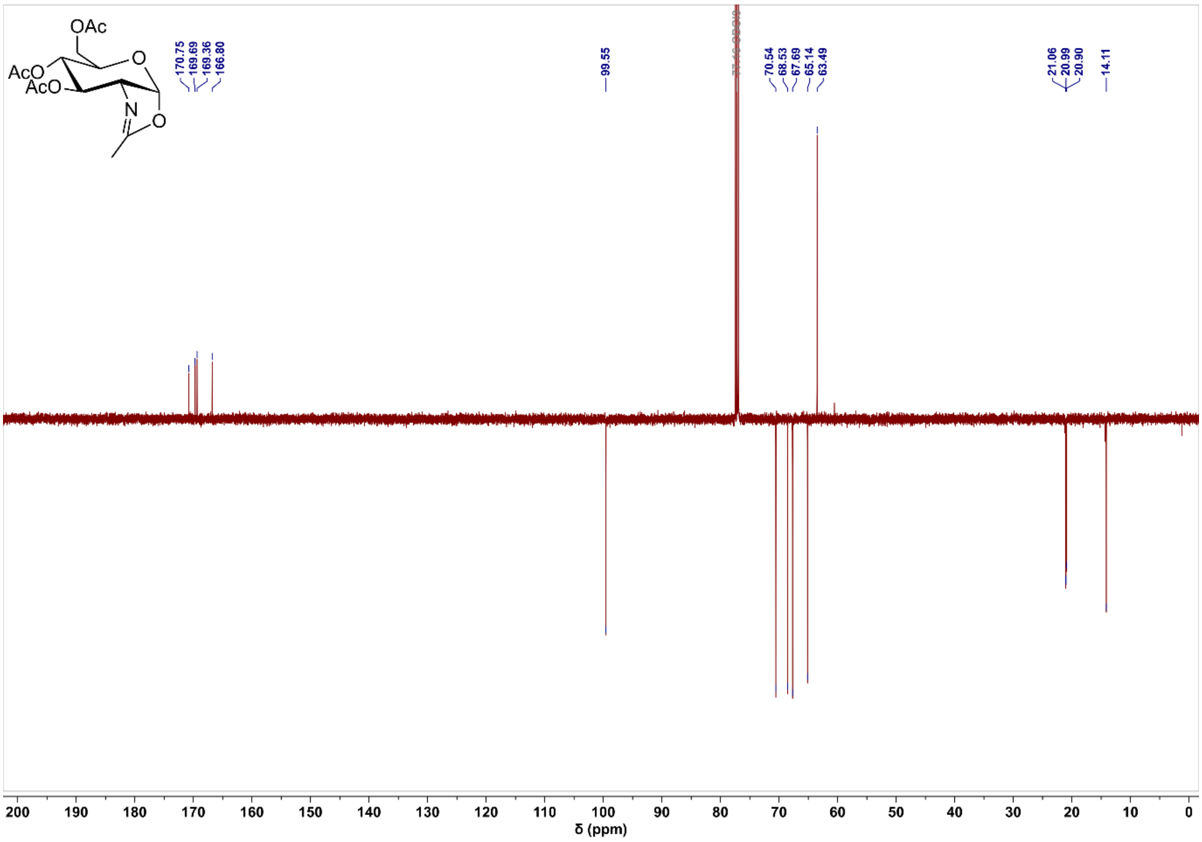

#### Compound 17 $^{1}$H NMR

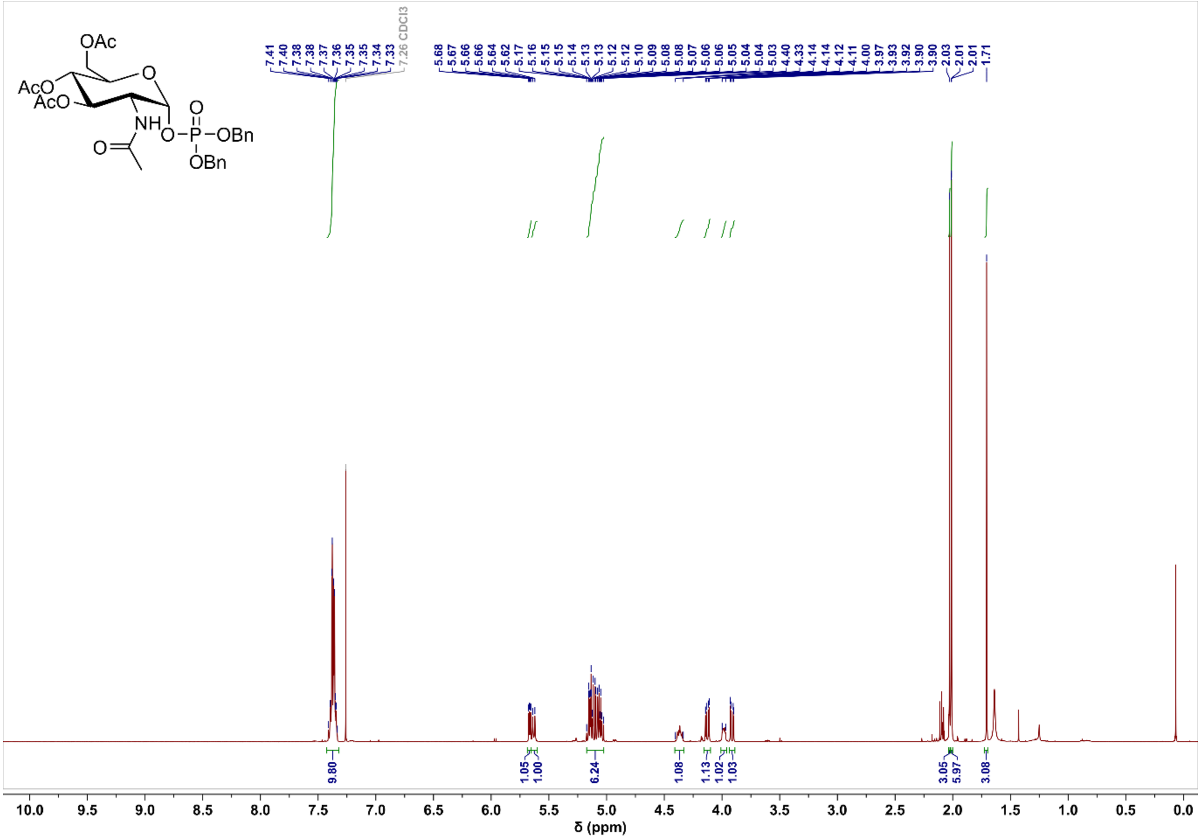

#### Compound 17 $^{13}$C NMR

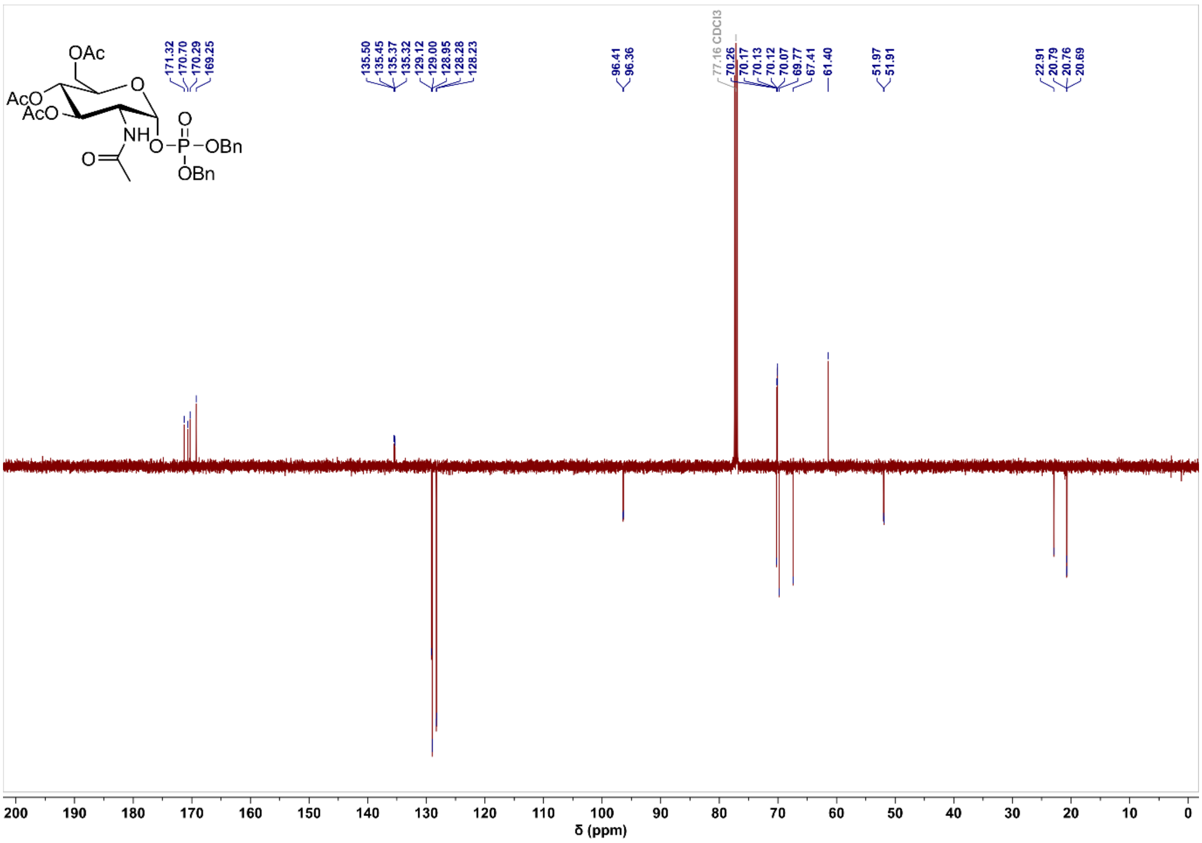

#### Compound 17 $^{31}$P NMR

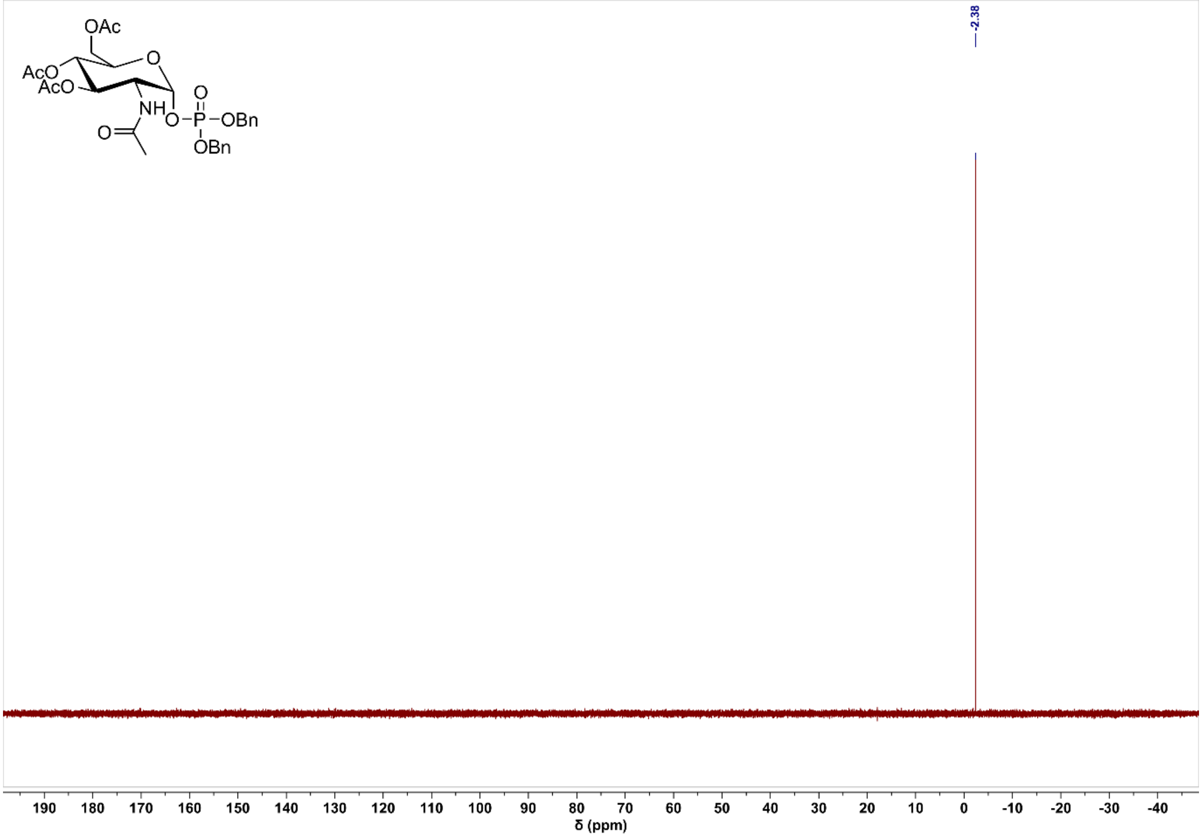

#### Compound 18 $^{1}$H NMR

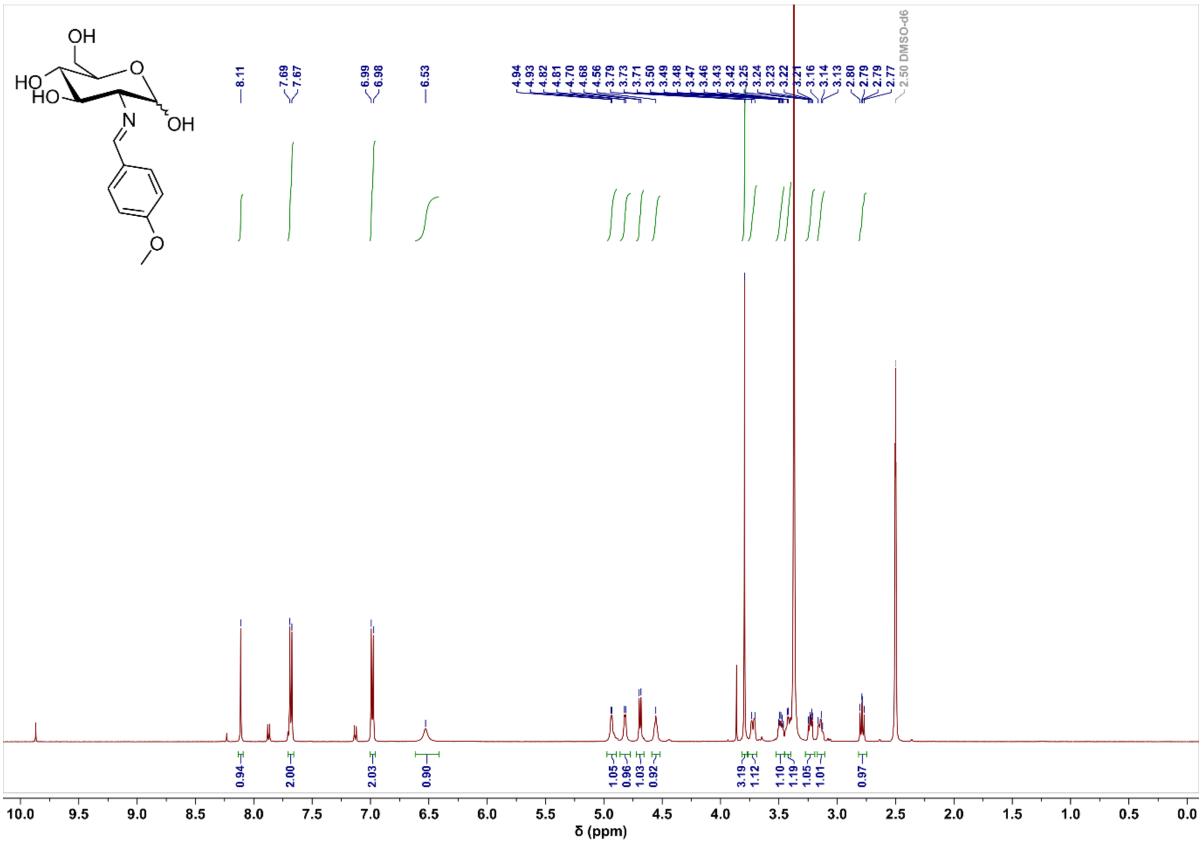

#### Compound 18 $^{13}$C NMR

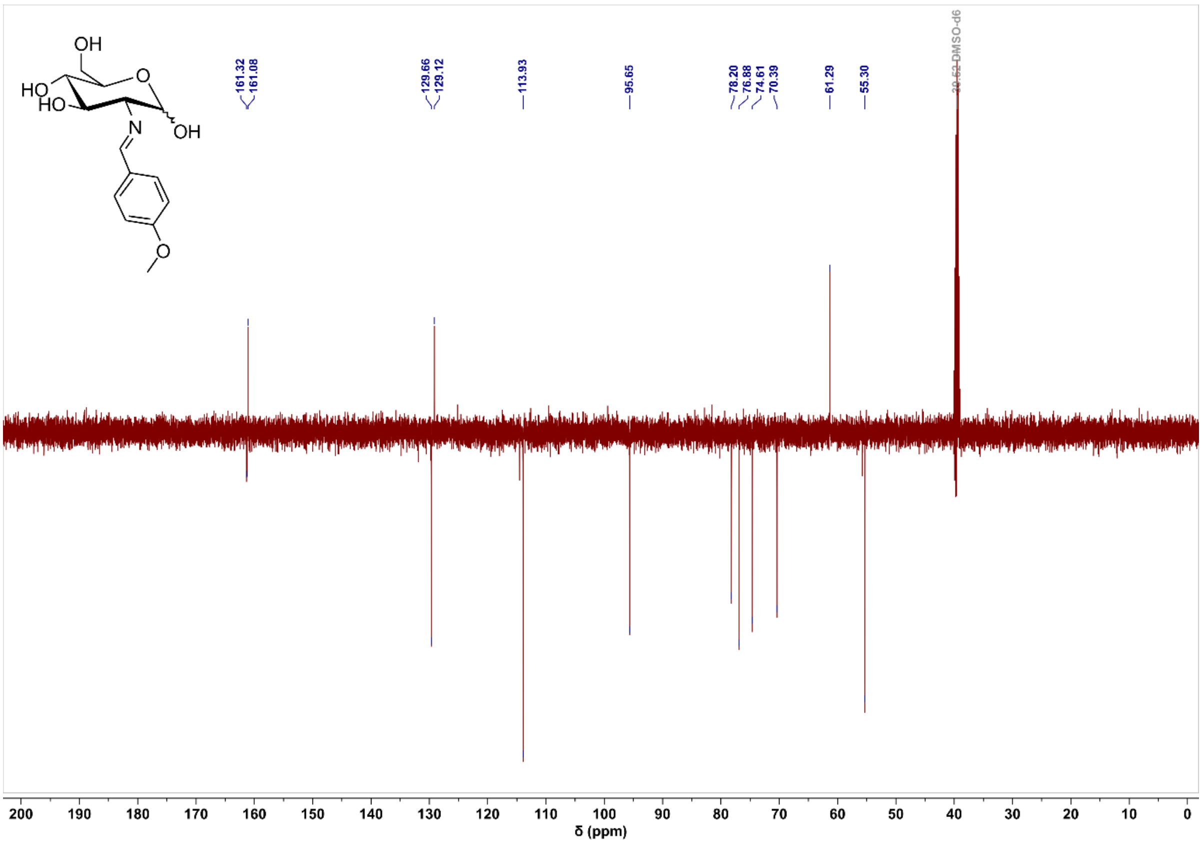

#### Compound 19 $^{1}$H NMR

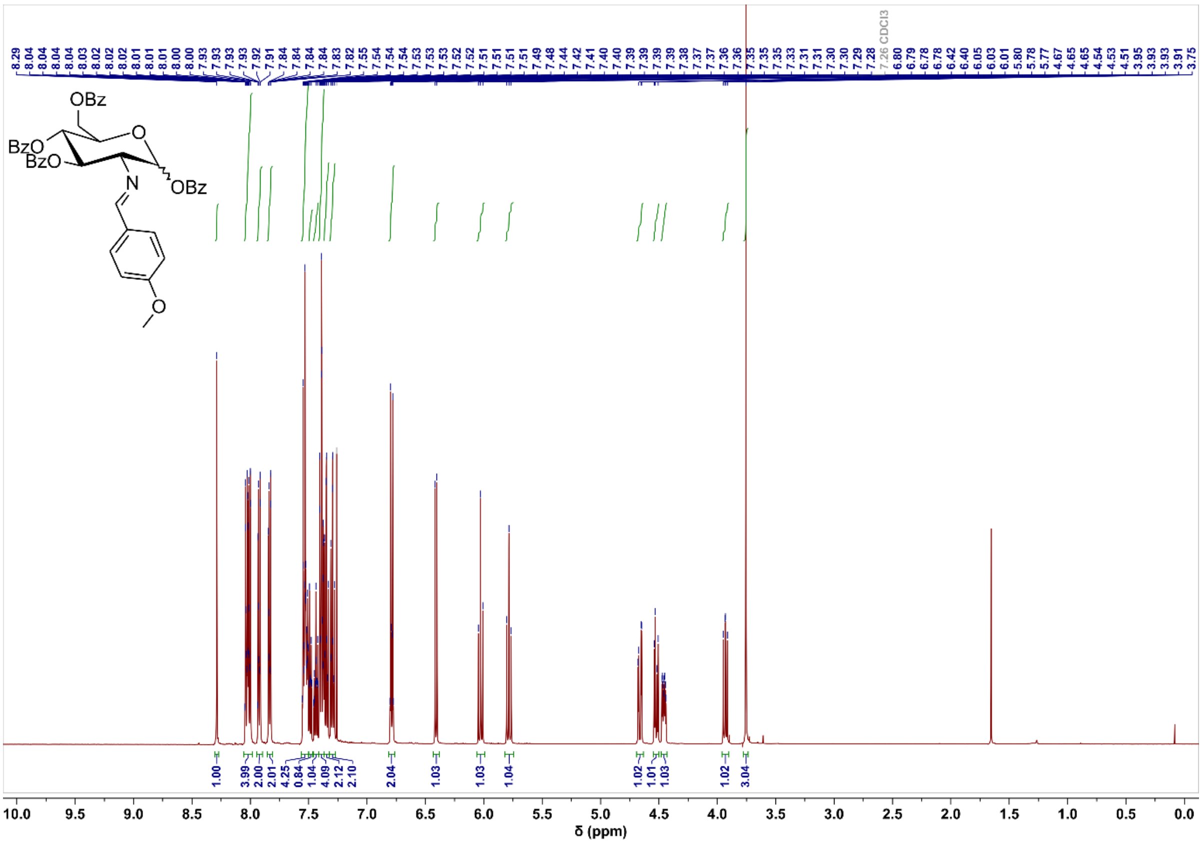

#### Compound 19 $^{13}C$ NMR

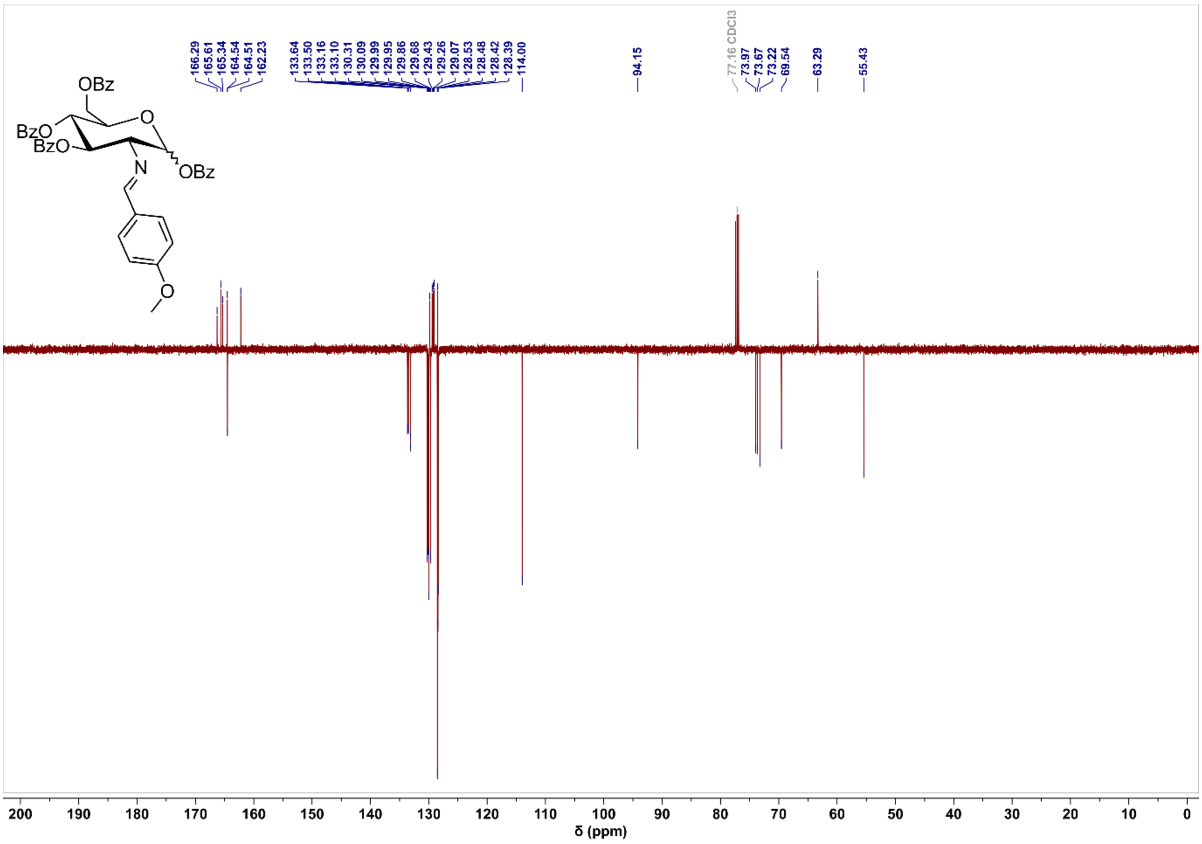

#### Compound 20 $^{1}H$ NMR

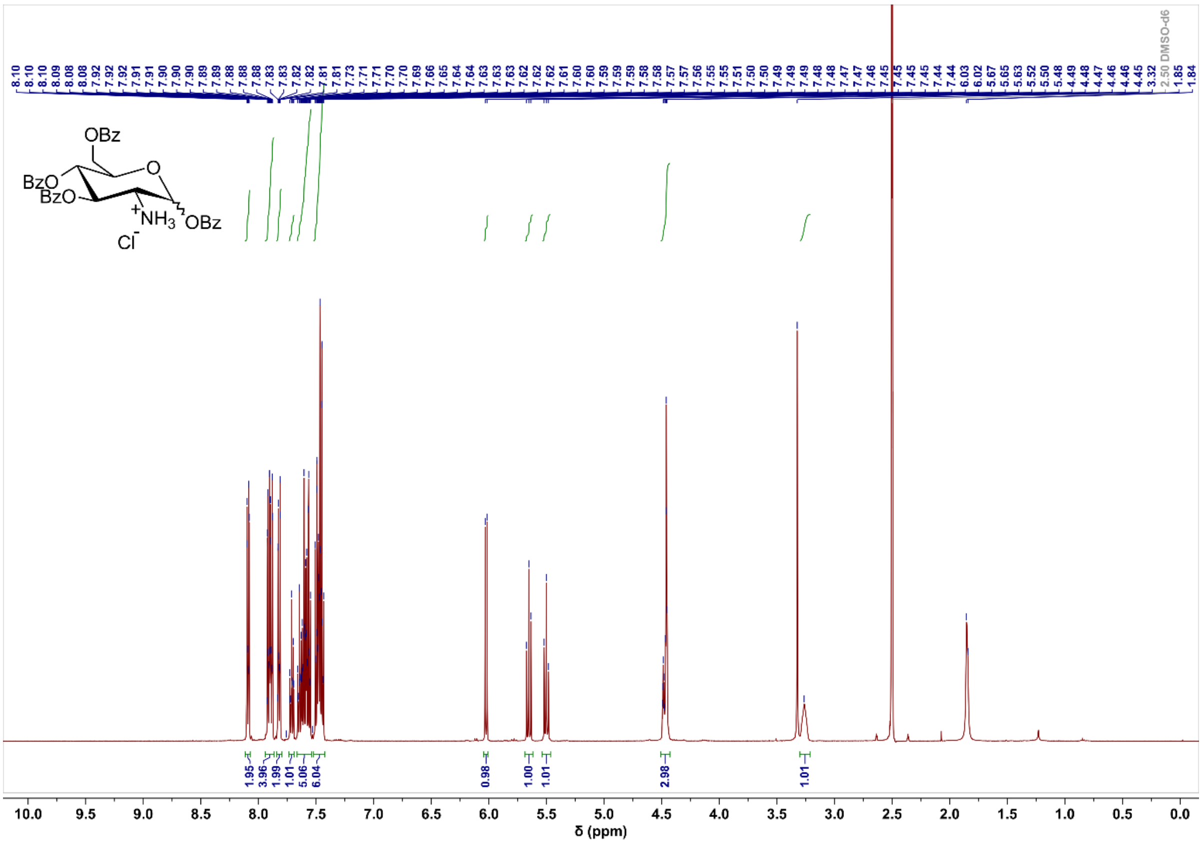

#### Compound 20 $^{13}C$ NMR

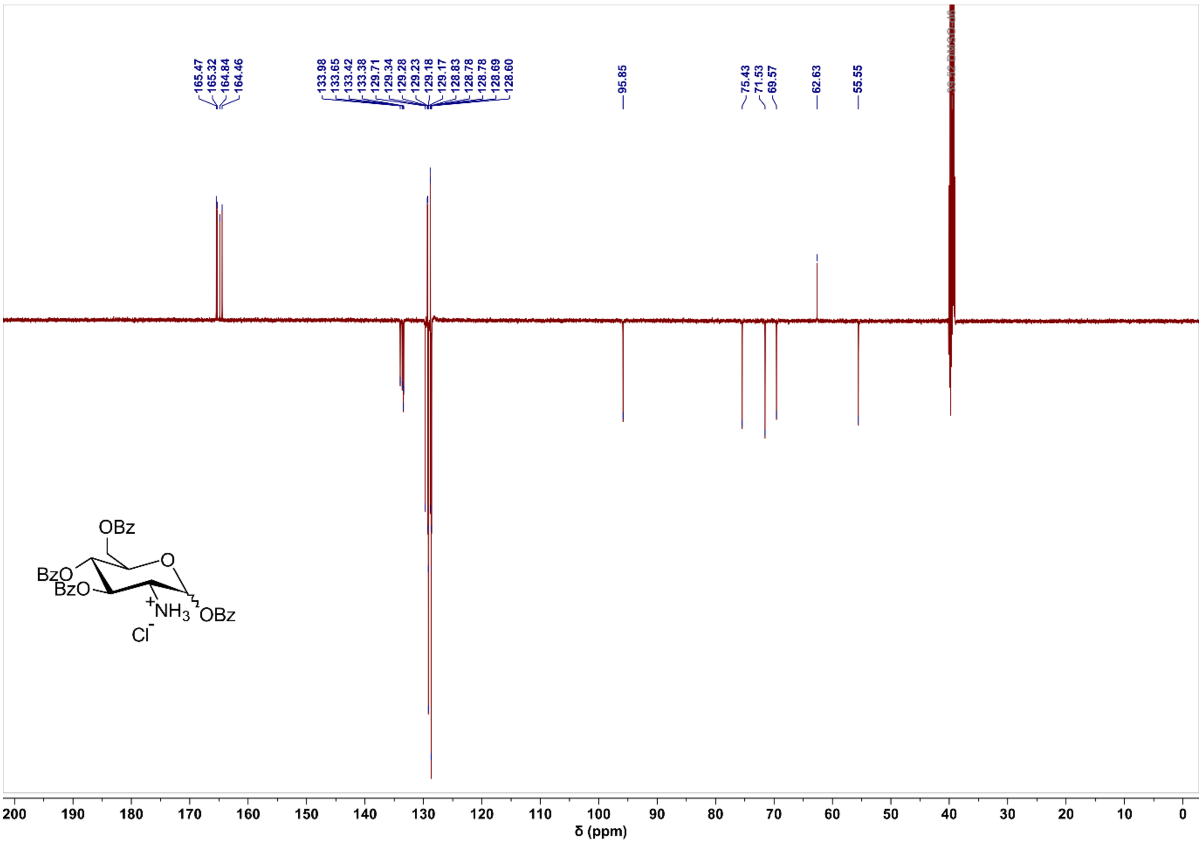

#### Compound 21 $^{1}H$ NMR

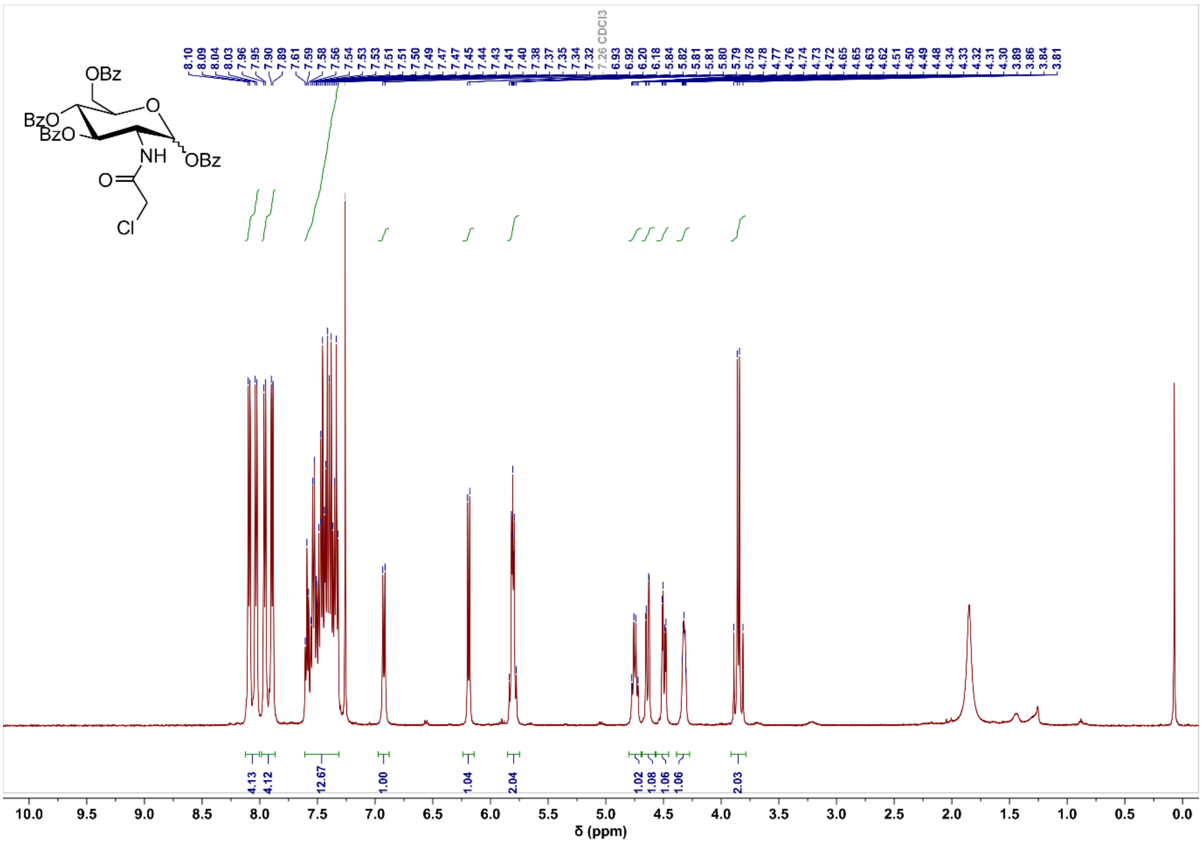

#### Compound 21 $^{13}C$ NMR

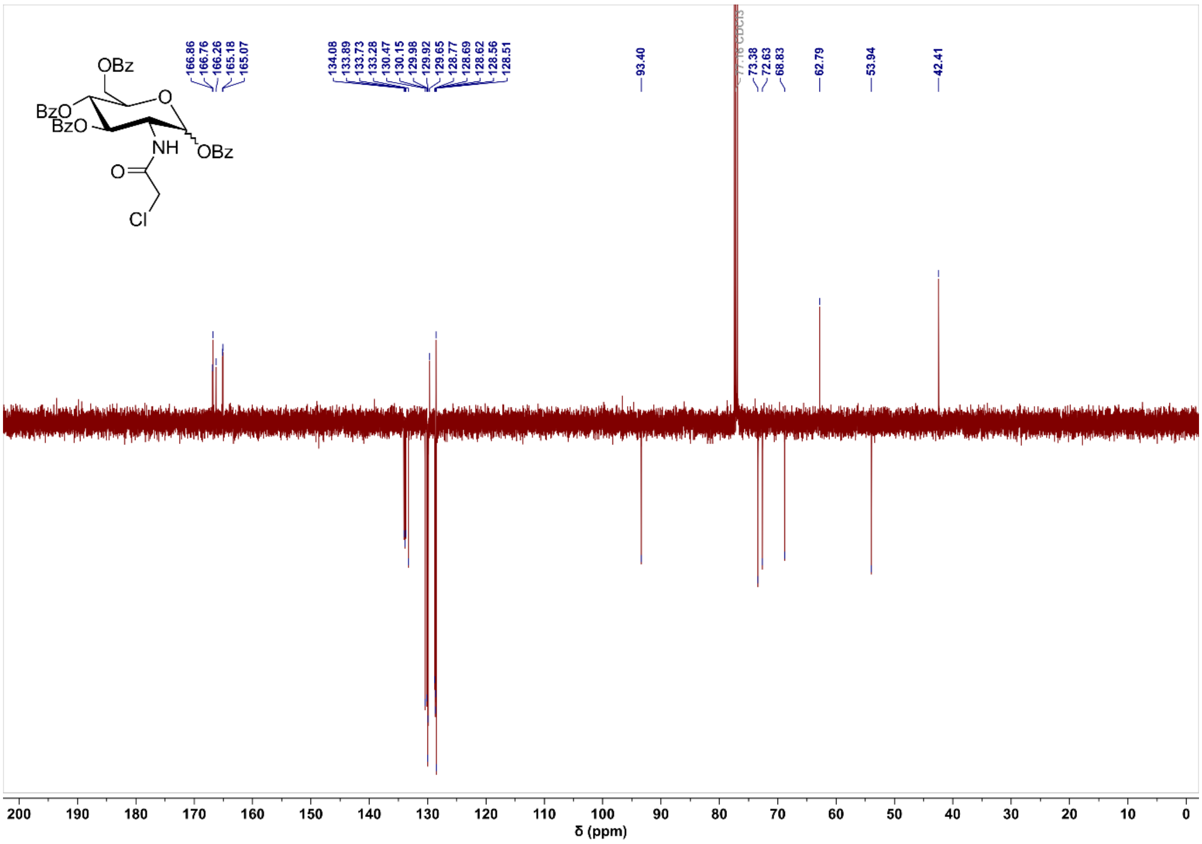

#### Compound 22 $^{1}H$ NMR

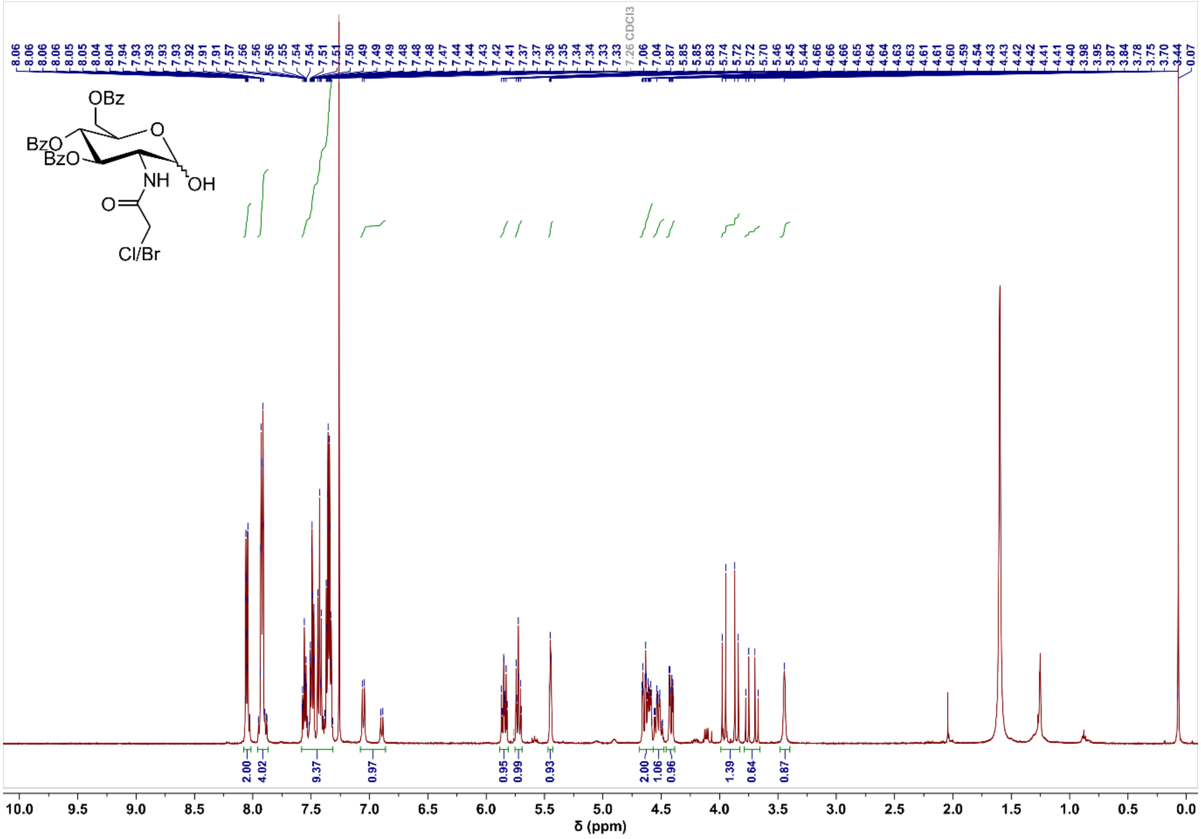

#### Compound 22 $^{13}$C NMR

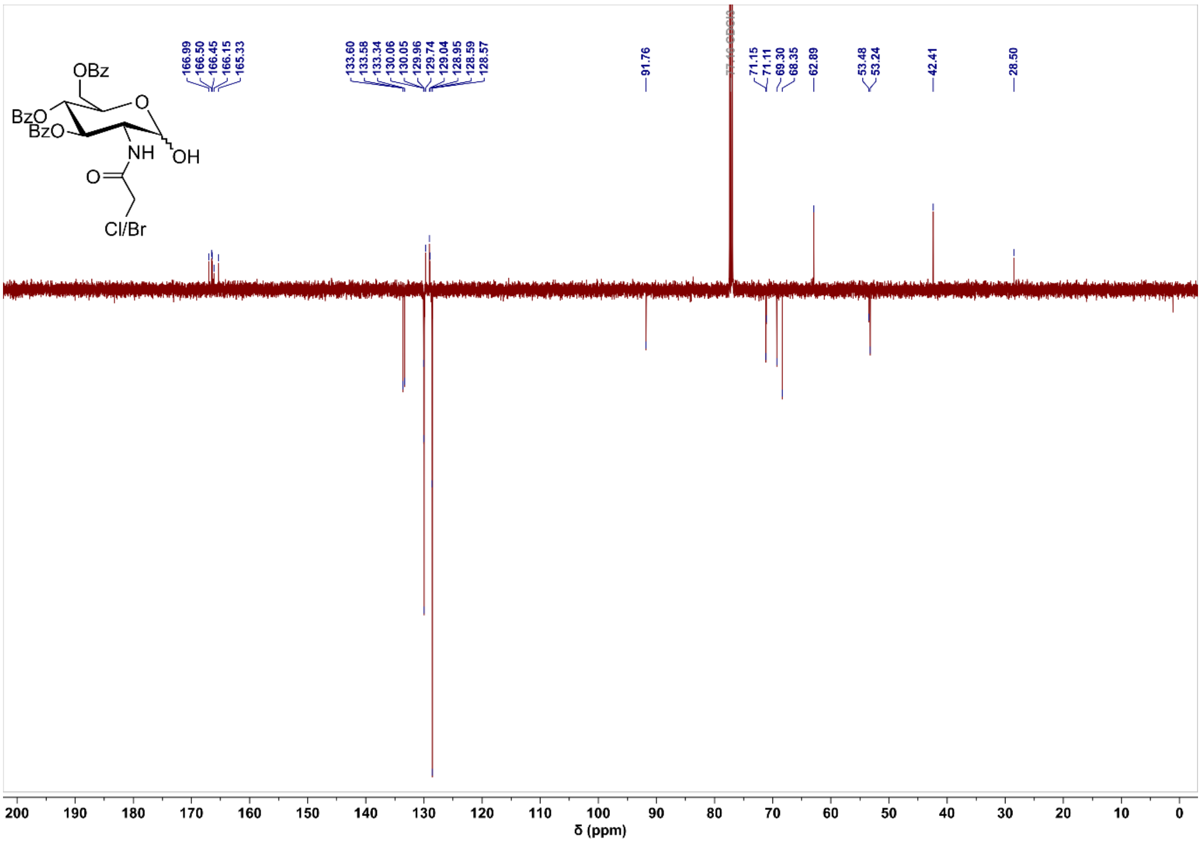

#### Compound 23 $^{1}$H NMR

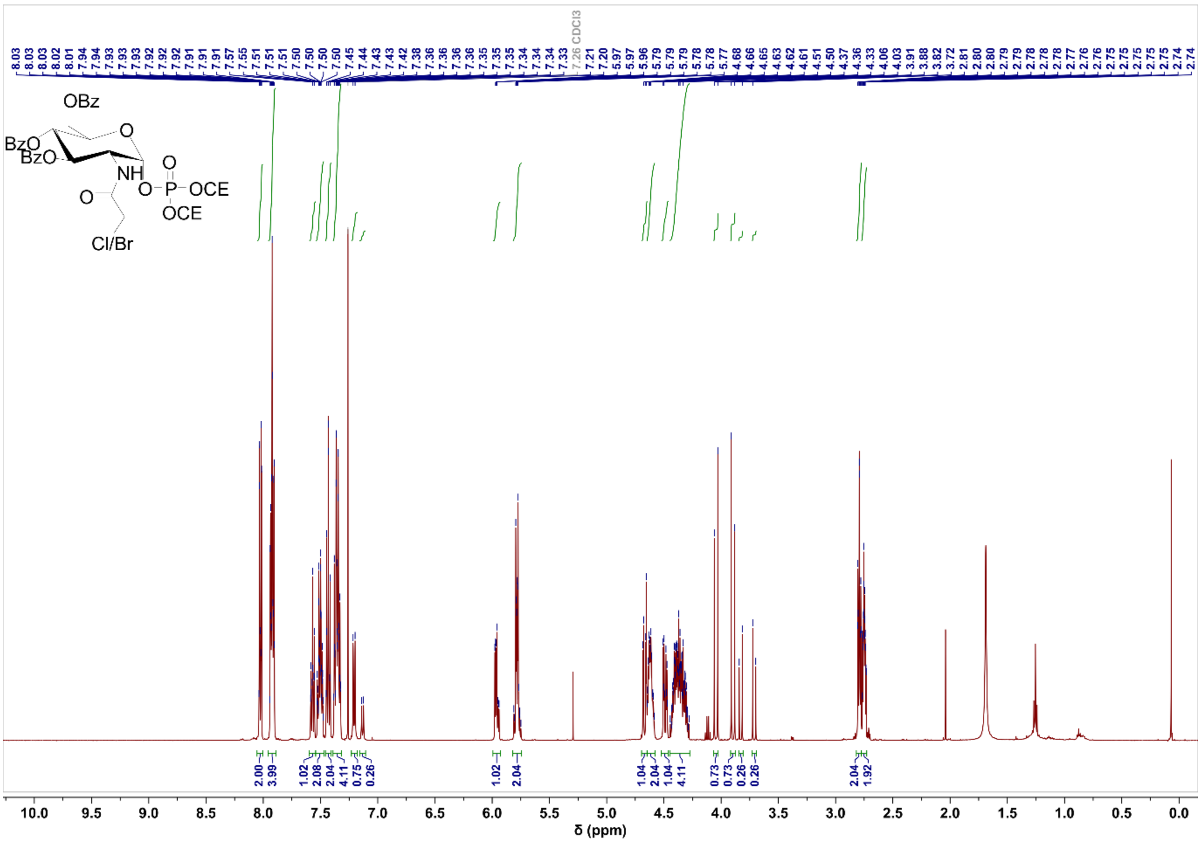

#### Compound 23 $^{13}$C NMR

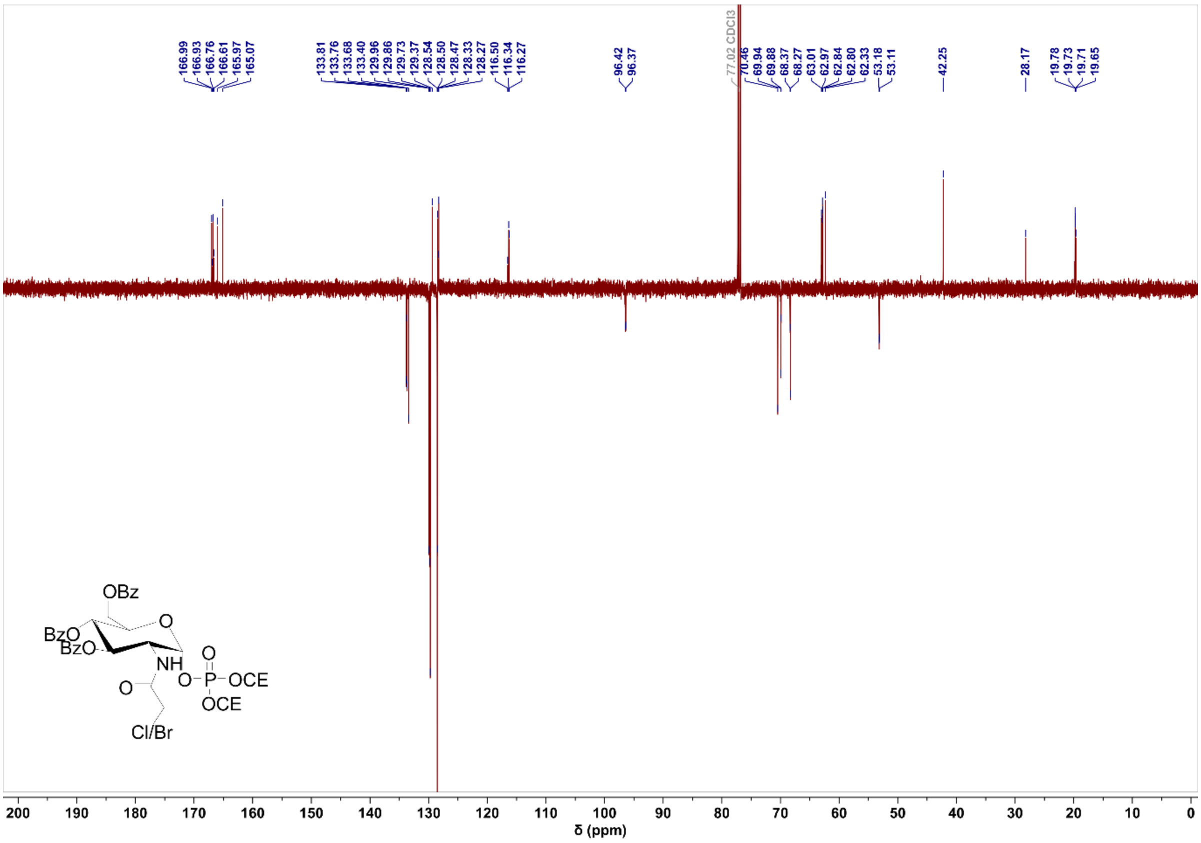

#### Compound 23 $^{31}$P NMR

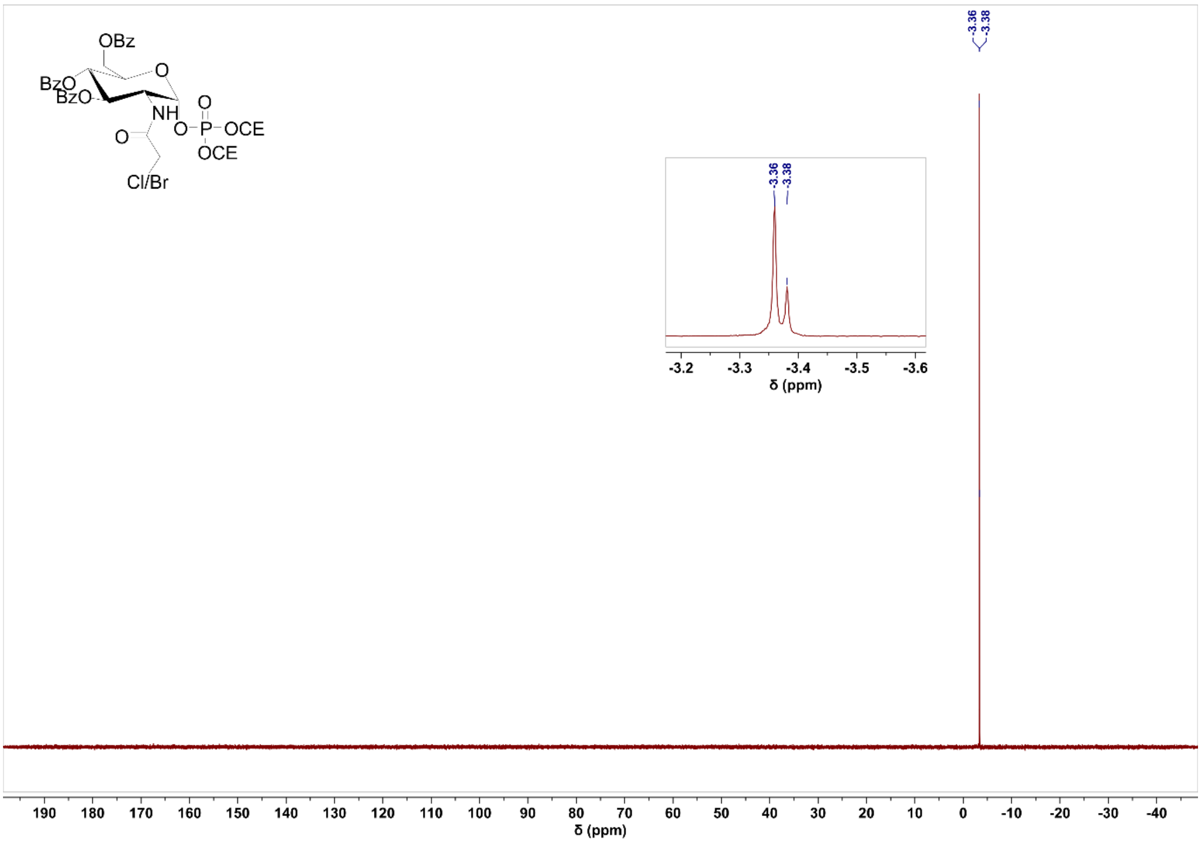

#### Compound 24 $^1$H NMR

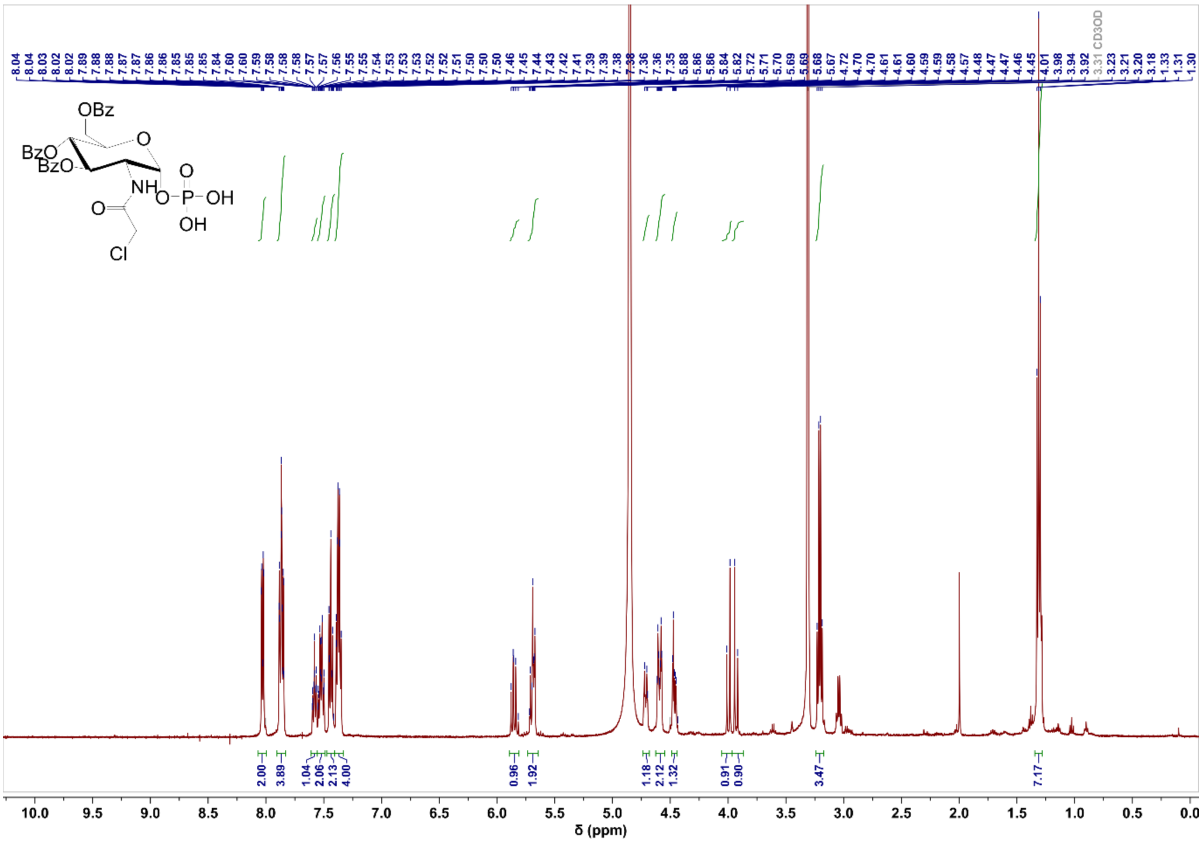

#### Compound 24 $^{13}$C NMR

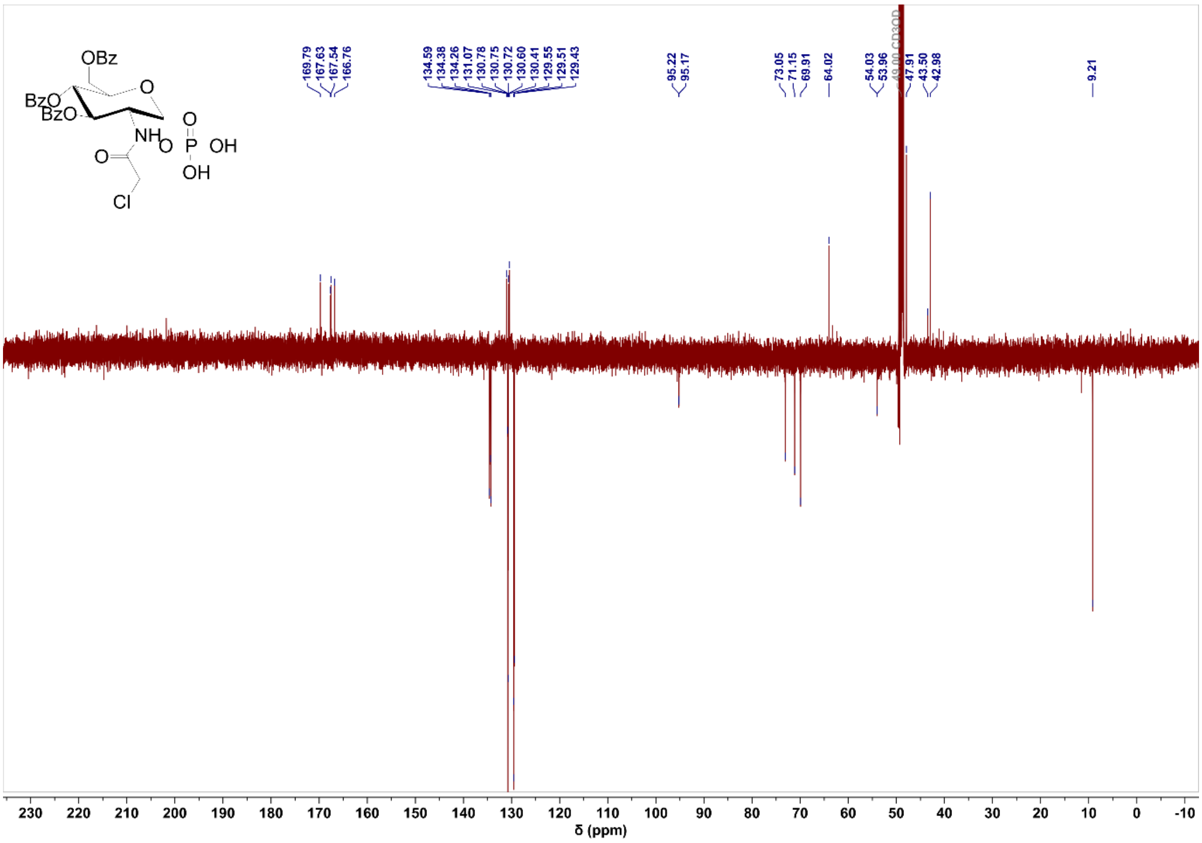

#### Compound 24 $^{31}$P NMR

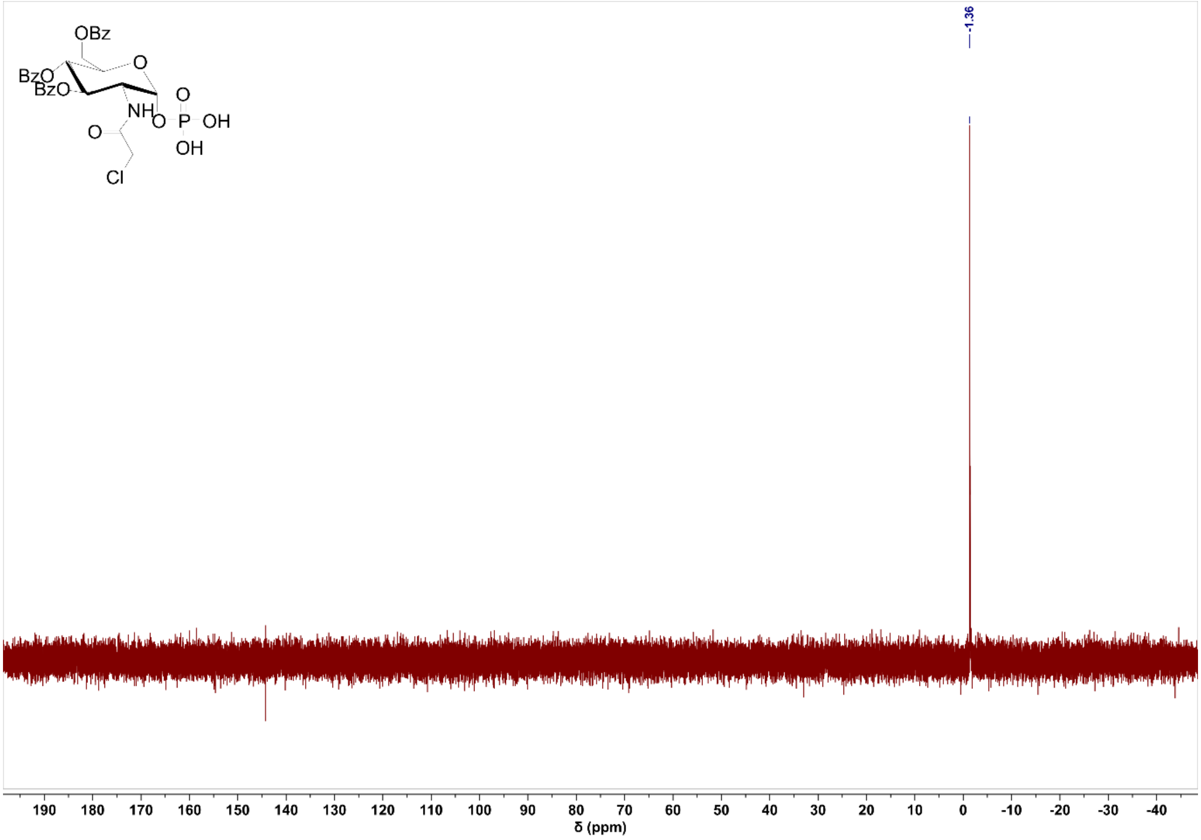

#### Compound 25 $^{1}$H NMR

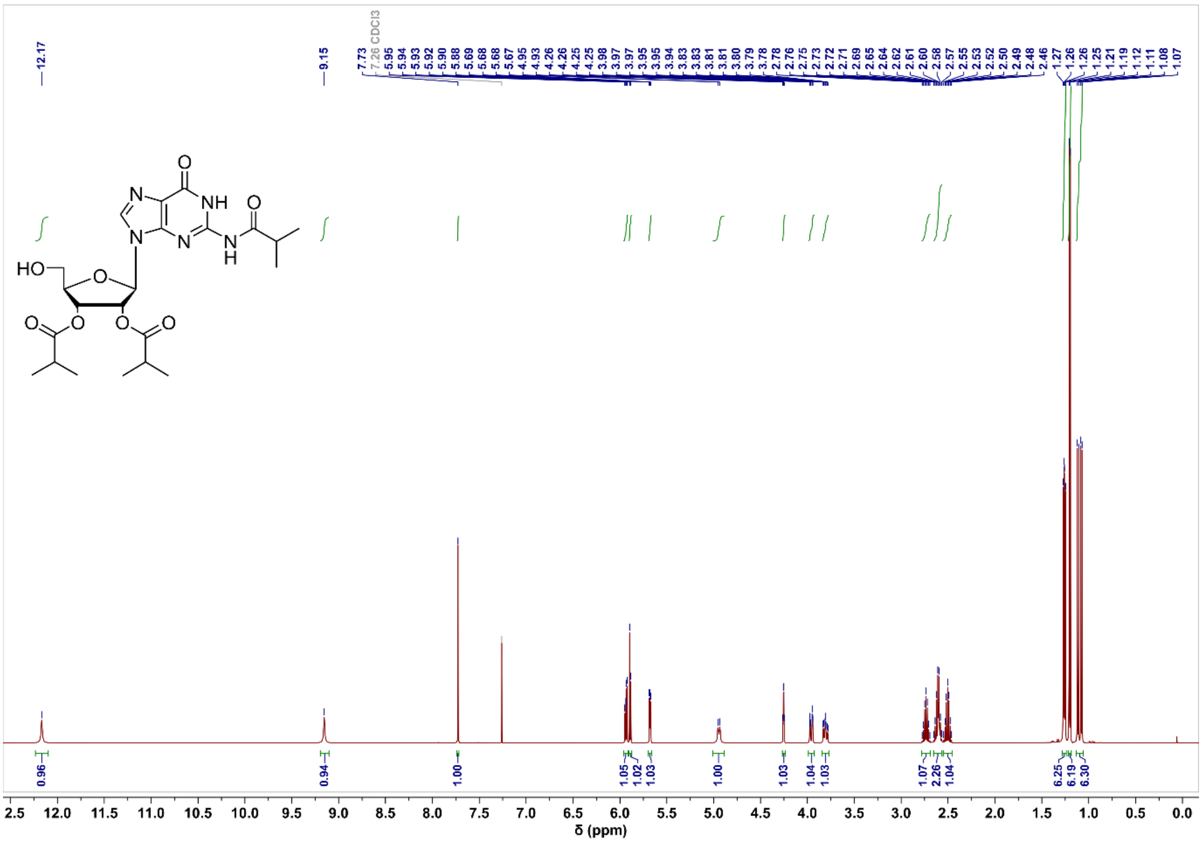

#### Compound 25 $^{13}$C NMR

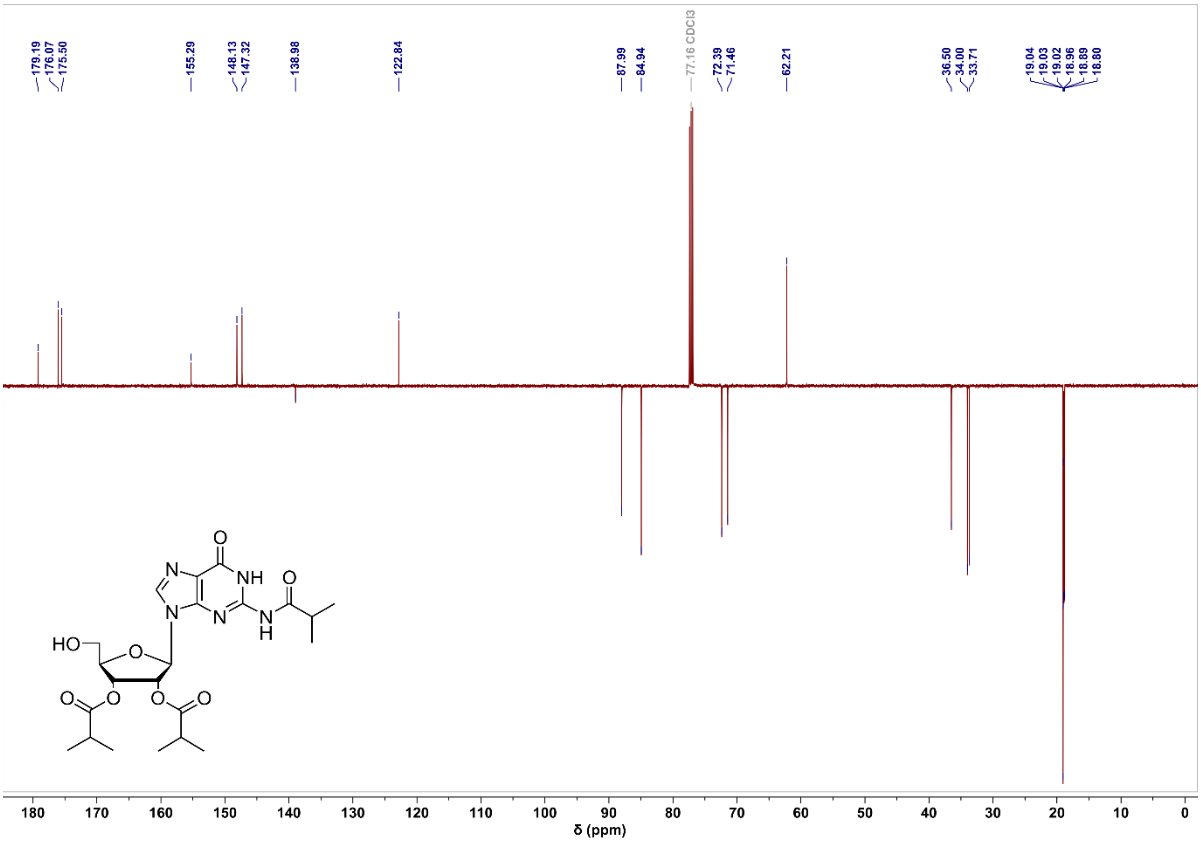

#### Compound 26 $^{1}$H NMR

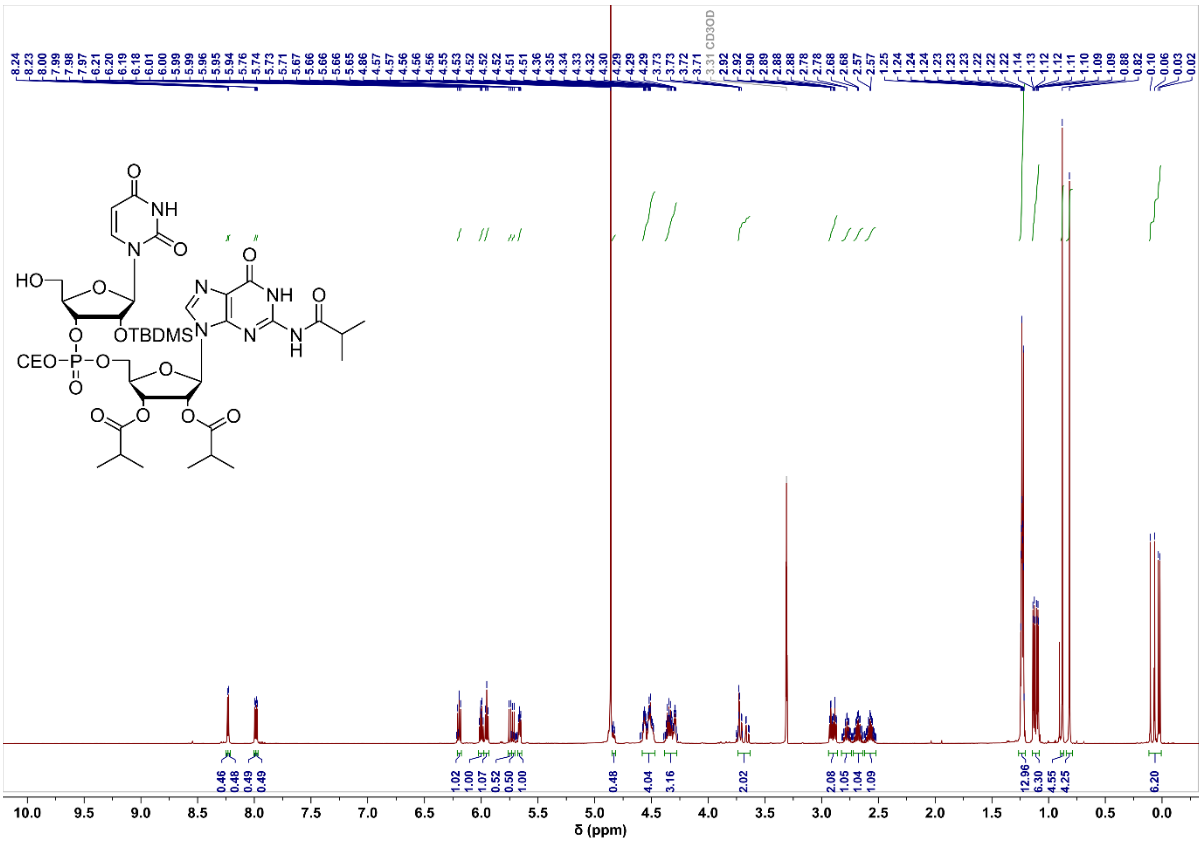

#### Compound 26 $^{13}C$ NMR

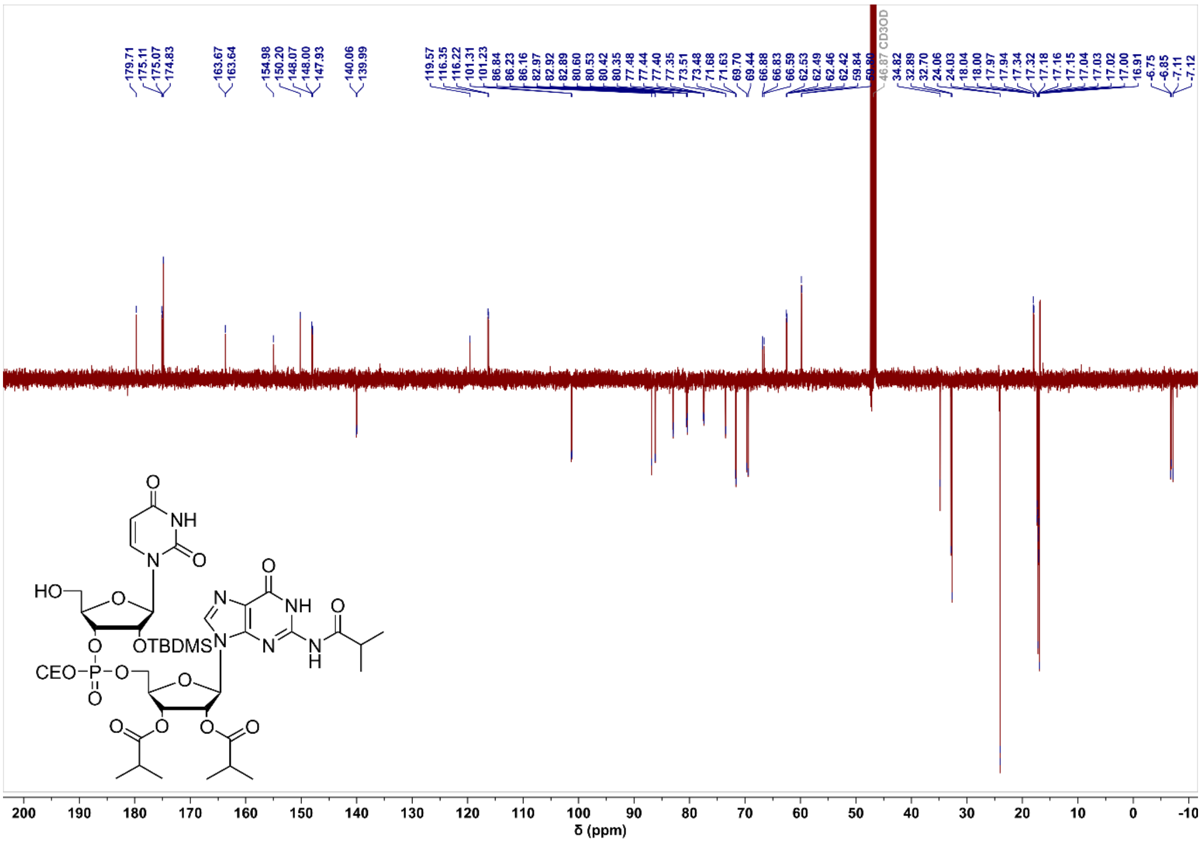

#### Compound 26 $^{31}P$ NMR

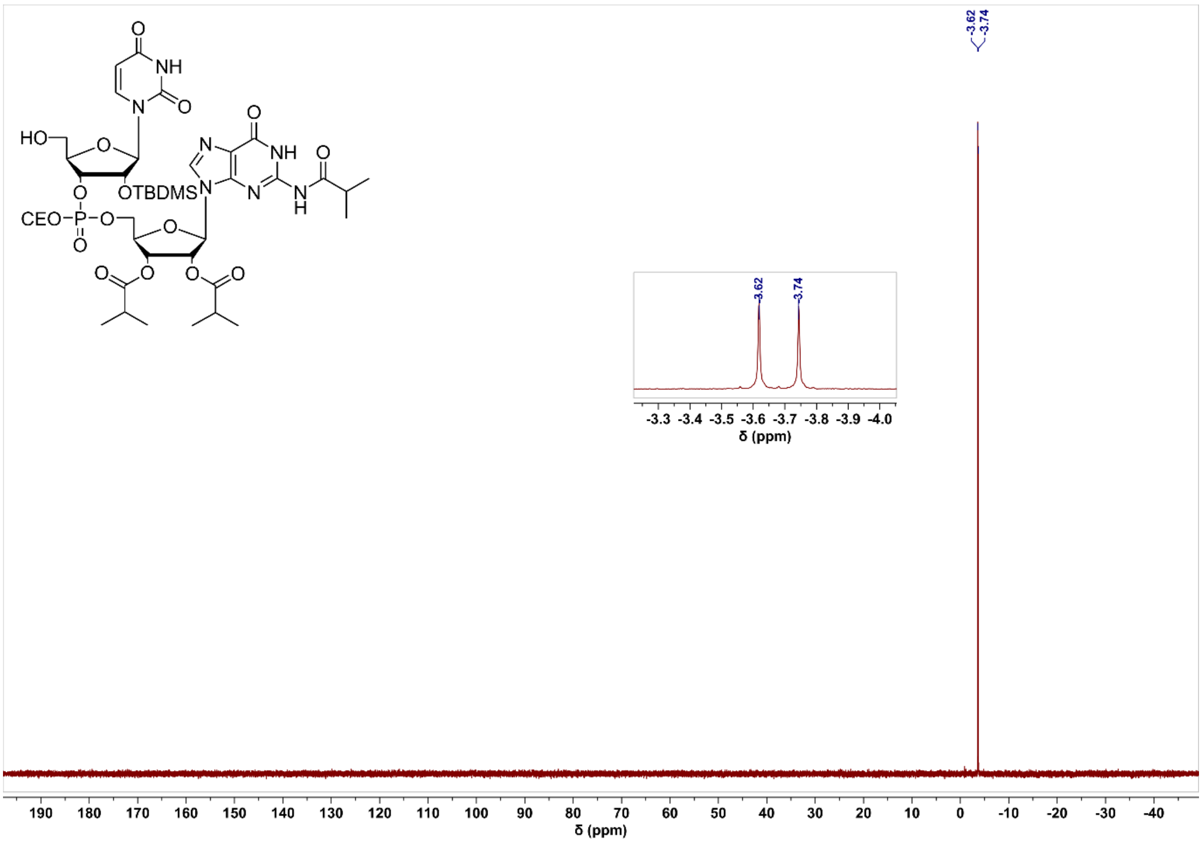

#### 6 References

1. Marquardt, J.L., Siegele, D.A., Kolter, R. and Walsh, C.T. (1992) Cloning and sequencing of *Escherichia coli murZ* and purification of its product, a UDP-*N*-acetylglucosamine enolpyruvyl transferase. *J. Bacteriol.*, **174**, 5748-5752.
2. Benson, T.E., Marquardt, J.L., Marquardt, A.C., Etzkorn, F.A. and Walsh, C.T. (1993) Overexpression, purification, and mechanistic study of UDP-*N*-acetylenolpyruvylglucosamine reductase. *Biochemistry*, **32**, 2024-2030.
3. Liger, D., Masson, A., Blanot, D., Vanheijenoort, J. and Parquet, C. (1995) Over-production, purification and properties of the uridine-diphosphate-*N*-acetylmuramate:L-alanine ligase from *Escherichia coli*. *Eur. J. Biochem.*, **230**, 80-87.
4. Dunn, C.A., O'Handley, S.F., Frick, D.N. and Bessman, M.J. (1999) Studies on the ADP-ribose pyrophosphatase subfamily of the Nudix hydrolases and tentative identification of *trgB*, a gene associated with tellurite resistance. *J. Biol. Chem.*, **274**, 32318-32324.
5. Yang, H., Slupska, M.M., Wei, Y.F., Tai, J.H., Luther, W.M., Xia, Y.R., Shih, D.M., Chiang, J.H., Baikalov, C., Fitz-Gibbon, S. *et al.* (2000) Cloning and characterization of a new member of the Nudix hydrolases from human and mouse. *J. Biol. Chem.*, **275**, 8844-8853.
6. Gasmi, L., Cartwright, J.L. and McLennan, A.G. (1999) Cloning, expression and characterization of YSA1H, a human adenosine 5 '-diphosphosugar pyrophosphatase possessing a MutT motif. *Biochem J.*, **344**, 331-337.
7. Skarzynski, T., Mistry, A., Wonacott, A., Hutchinson, S.E., Kelly, V.A. and Duncan, K. (1996) Structure of UDP-*N*-acetylglucosamine enolpyruvyl transferase, an enzyme essential for the synthesis of bacterial peptidoglycan, complexed with substrate UDP-*N*-acetylglucosamine and the drug fosfomycin. *Structure*, **4**, 1465-1474.
8. Benson, T.E., Walsh, C.T. and Hogle, J.M. (1997) X-ray crystal structures of the S229A mutant and wild-type MurB in the presence of the substrate enolpyruvyl-UDP-*N*-acetylglucosamine at 1.8-Å resolution. *Biochemistry*, **36**, 806-811.
9. Mol, C.D., Brooun, A., Dougan, D.R., Hilgers, M.T., Tari, L.W., Wijnands, R.A., Knuth, M.W., McRee, D.E. and Swanson, R.V. (2003) Crystal structures of active fully assembled substrate- and product-bound complexes of UDP-*N*-acetylmuramic acid:L-alanine ligase (MurC) from *Haemophilus influenzae*. *J. Bacteriol.*, **185**, 4152-4162.
10. Gabelli, S.B., Bianchet, M.A., Bessman, M.J. and Amzel, L.M. (2001) The structure of ADP-ribose pyrophosphatase reveals the structural basis for the versatility of the Nudix family. *Nat. Struct. Biol.*, **8**, 467-472.
11. Zha, M., Zhong, C., Peng, Y., Hu, H. and Ding, J. (2006) Crystal structures of human NUDT5 reveal insights into the structural basis of the substrate specificity. *J. Mol. Biol.*, **364**, 1021-1033.
12. Wang, S., Czuchry, D., Liu, B., Vinnikova, A.N., Gao, Y., Vlahakis, J.Z., Szarek, W.A., Wang, L., Feng, L. and Brockhausen, I. (2014) Characterization of two UDP-Gal:GalNAc-diphosphate-lipid β1,3-galactosyltransferases WbwC from *Escherichia coli* serotypes O104 and O5. *J. Bacteriol.*, **196**, 3122-3133.
13. Heidlas, J.E., Lees, W.J., Pale, P. and Whitesides, G.M. (1992) Gram-scale synthesis of uridine 5'-diphospho-*N*-acetylglucosamine: Comparison of enzymatic and chemical routes. *J. Org. Chem.*, **57**, 146-151.
14. Kadokura, M., Wada, T., Seio, K. and Sekine, M. (2000) Synthesis of 4-thiouridine, 6-thioinosine, and 6-thioguanosine 3',5'-*O*-bisphosphates as donor molecules for RNA ligation and their application to the synthesis of photoactivatable TMG-capped U1 snRNA fragments. *J. Org. Chem.*, **65**, 5104-5113.
15. Mizyed, S., Oddone, A., Byczynski, B., Hughes, D.W. and Berti, P.J. (2005) UDP-*N*-acetylmuramic acid (UDP-MurNAc) is a potent inhibitor of MurA (enolpyruvyl-UDP-GlcNAc synthase). *Biochemistry*, **44**, 4011-4017.
16. Doyle, L.M., O'Sullivan, S., Di Salvo, C., McKinney, M., McArdle, P. and Murphy, P.V. (2017) Stereoselective epimerizations of glycosyl thiols. *Org. Lett.*, **19**, 5802-5805.

17. Mbadugha, B.N.A. and Menger, F.M. (2003) Sugar/steroid/sugar conjugates: Sensitivity of lipid binding to sugar structure. *Org. Lett.*, **5**, 4041-4044.
18. Dong, Y.Y., Wang, H., Pike, A.C.W., Cochrane, S.A., Hamedzadeh, S., Wyszyński, F.J., Bushell, S.R., Royer, S.F., Widdick, D.A., Sajid, A. *et al.* (2018) Structures of DPAGT1 explain glycosylation disease mechanisms and advance TB antibiotic design. *Cell*, **175**, 1045-1058.
19. Seelig, B. and Jäschke, A. (1999) Ternary conjugates of guanosine monophosphate as initiator nucleotides for the enzymatic synthesis of 5 '-modified RNAs. *Bioconjugate Chem.*, **10**, 371-378.
20. Krause, A., Hertl, A., Muttach, F. and Jäschke, A. (2014) Phosphine-free Stille-Migita chemistry for the mild and orthogonal modification of DNA and RNA. *Chem. Eur. J.*, **20**, 16613-16619.
